# Multi-omics links microbial dysbiosis, systemic inflammation and metabolomic disruptions to SNAE risk in treated HIV

**DOI:** 10.64898/2026.04.09.717347

**Authors:** Christopher M. Basting, Jodi Anderson, Kevin Escandón, Garritt Wieking, Candace Guerrero, Jarrett Reichel, Ross Cromarty, Erik Swanson, Ty Schroeder, Elaina Creagan, Maura Barrett, Nicholas Funderburg, Peter Hunt, Melanie Graham, Santiago Avila-Rios, Gonzalo Salgado, Timothy W. Schacker, Nichole R. Klatt

**Affiliations:** Division of Surgical Outcomes and Precision Medicine Research, Department of Surgery, University of Minnesota, Minneapolis, MN, USA; Division of Infectious Diseases and International Medicine, Department of Medicine, University of Minnesota, Minneapolis, MN, USA; Masonic Cancer Center, University of Minnesota, Minneapolis, MN, USA; School of Health and Rehabilitation Sciences, Ohio State University, Columbus, OH, USA; Division of Experimental Medicine, University of California San Francisco. San Francisco, CA, USA; Pre-Clinical Research Center, Department of Surgery, University of Minnesota, Minneapolis, MN, USA; Instituto Nacional de Enfermedades Respiratorias Ismael Cosío Villegas, Centro de Investigación en Enfermedades Infecciosas, Mexico City, Mexico

## Abstract

Serious non-AIDS events (SNAEs), including non-AIDS malignancies, cardiovascular disease, and hepatic complications, remain major causes of mortality in treated HIV infection. These outcomes are driven by persistent immune activation, systemic inflammation, and metabolic dysfunction despite effective viral suppression with antiretroviral therapy (ART). To investigate mechanisms underlying SNAE pathogenesis, we performed a cross-site multi-omic analysis integrating plasma proteins, plasma metabolites, and mucosal microbiomes (ileum and rectum) in 82 ART-treated people with HIV (PWH) and 10 people without HIV (PWoH) from the United States and Mexico. Geography was the dominant source of variation, particularly across lipid classes. However, individuals at high risk for SNAEs, defined by low CD4 T cell counts and low CD4/CD8 ratios, shared a consistent signature of systemic inflammation, mitochondrial dysfunction, and microbial dysbiosis including elevated plasma IL-6, ω-oxidation products (adipic and suberic acids), and depletion of short-chain fatty acid–producing commensals in the gut mucosa, including *Akkermansia muciniphila*, *Bacteroides uniformis*, and *Ruminococcus*. Notably, *A. muciniphila* abundance correlated with lower IL-6 levels, fewer HIV RNA-producing cells in lymph nodes, and higher CD4/CD8 ratios. Together, these findings identify a shared inflammatory and metabolic phenotype in PWH and implicate *A. muciniphila* as a potential microbiome-based target to mitigate immune activation and SNAE risk in treated HIV.

## Introduction

Antiretroviral therapy (ART) has transformed HIV into a chronic condition with near-normal life expectancy, yet a gap remains largely due to serious non-AIDS events (SNAEs). These include non-AIDS malignancies, cardiovascular events (e.g., myocardial infarction and stroke), and hepatic disease, which remain major causes of morbidity and mortality in treated HIV infection despite ART^1,2^. The pathogenesis of SNAEs is multifactorial, attributed in part to chronic immune activation sustained by microbial translocation, co-infections, and persistent HIV reservoirs^3–7^. This chronic immune activation drives systemic inflammation as well as immune dysfunction characterized by altered T cell counts, immune cell senescence, and metabolic dysfunction, further contributing to SNAE pathogenesis^8,9^. T cell criteria including low CD4^+^ T cell counts and a low CD4/CD8 T cell ratio are particularly well-established correlates of immune activation and dysfunction, often used as end points in interventional studies and predictors of SNAE risk^4,10^.

Current strategies to prevent SNAEs focus on early ART initiation, treatment of co-infections such as cytomegalovirus (CMV) and Epstein-Barr virus, and cardioprotective lifestyle changes^1^, though these do not fully address the sources of immune activation and dysfunction that underlie SNAE pathogenesis. Interventional studies targeting these sources, such as fibrosis or microbial translocation, have had mixed results at improving correlates of SNAE risk^1,11–14^, highlighting the need for a better understanding of mechanisms driving SNAE pathogenesis. This is especially crucial for individuals at highest risk, such as those with <350 CD4^+^ T cells/µL after 2 years of ART, who are nearly three times more likely to have SNAE over the subsequent 5 years than those with >350 CD4^+^ T cells/ µL, with similar associations observed for low CD4/CD8 ratios^10^. Interventions that effectively reduce immune activation and restore immune homeostasis in these individuals could substantially lower SNAE incidence and improve long-term survival of people with HIV (PWH).

To address this knowledge gap, we conducted a cross-site, multi-omic analysis integrating plasma proteins (cytokines, microbial translocation and gut barrier damage markers), the mucosal microbiome (ileum and rectum), and plasma metabolome in 82 ART-treated PWH and 10 people without HIV (PWoH) enrolled from the United States (US) and Mexico. Our objectives were threefold: (1) to characterize the multi-omic differences between PWH from the two geographic cohorts; (2) to identify geography-independent biomarkers of SNAE risk that may reveal shared biological mechanisms underlying SNAE pathogenesis; and (3) to highlight potential therapeutic interventions that could mitigate the drivers of SNAE pathogenesis in high-risk individuals.

## Results

### Differences in baseline characteristics

Baseline characteristics for enrolled participants from the US and Mexico are shown in Table 1. Compared to PWH from the US, PWH from Mexico were older (p = 0.037), and had significantly lower CD4^+^ T cell counts (p = 0.006) and a lower nadir CD4^+^ T cell count (p = 0.003). When stratified by SNAE risk group (Table 2, with PWoH included as a reference), the high-risk group had significantly lower CD4^+^ T cell counts (p < 0.001), a lower CD4/CD8 ratio (p = 0.001), and lower nadir CD4^+^ T cell count (p = 0.002) compared to the low-risk group. There was no significant difference in country of enrollment (p = 0.6) or race (p = 0.8) between low- and high-risk groups. CMV IgG serostatus was assessed in all participants, of which, all but 1 were seropositive (Supplemental Tables 1 and 2). Summaries of ART regimens are provided with stratification by country (Supplemental Table 3) and SNAE risk group (Supplemental Table 4). Most enrolled participants (78%) were on an ART regimen consisting of two nucleoside/nucleotide reverse transcriptase inhibitors (NRTIs) and one integrase strand transfer inhibitor (INSTI). There was a significant difference in ART regimens between countries (p < 0.001), reflecting differences in national treatment guidelines. There was no difference in ART regimens between SNAE risk groups (p = 0.5).

**Table 1.** Baseline characteristics of HIV-positive patients by country.

|  | Mexico (n = 48) | US (n = 34) | p-value |
| --- | --- | --- | --- |
| **Characteristic (Median [Q1, Q3])** |  |  |  |
| CD4<sup>+</sup> T cell count (cells/mm<sup>3</sup>) | 478 [329, 650] | 643 [441, 787] | 0.006<sup>A</sup> |
| CD8<sup>+</sup> T cell count (cells/mm<sup>3</sup>) | 685 [445, 872] | 800 [506, 1,087] | 0.058<sup>A</sup> |
| CD4/CD8 ratio | 0.77 [0.55, 0.99] | 1.00 [0.58, 1.20] | 0.5<sup>A</sup> |
| Nadir CD4<sup>+</sup> T cell count (cells/mm<sup>3</sup>) | 109 [31, 306] | 280 [170, 434] | 0.003<sup>A</sup> |
| Time living with HIV (years) | 12 [7, 15] | 13 [4, 22] | 0.2<sup>A</sup> |
| Age (years) | 53 [48, 57] | 46 [34, 59] | 0.037<sup>A</sup> |
| Sex at birth (n [%]) |  |  | >0.9<sup>B</sup> |
| Female | 2 (4.2%) | 1 (2.9%) |  |
| Male | 46 (96%) | 33 (97%) |  |
| Race/ethnicity (n [%]) |  |  | <0.001<sup>B</sup> |
| Black | 0 (0%) | 14 (41%) |  |
| Hispanic | 48 (100%) | 4 (12%) |  |
| Other | 0 (0%) | 4 (12%) |  |
| White | 0 (0%) | 12 (35%) |  |

Clinical characteristics of patients enrolled in the study, stratified by country of enrollment.
<sup>A</sup>Welch Two Sample t-test. <sup>B</sup>Fisher's exact test

**Table 2.**
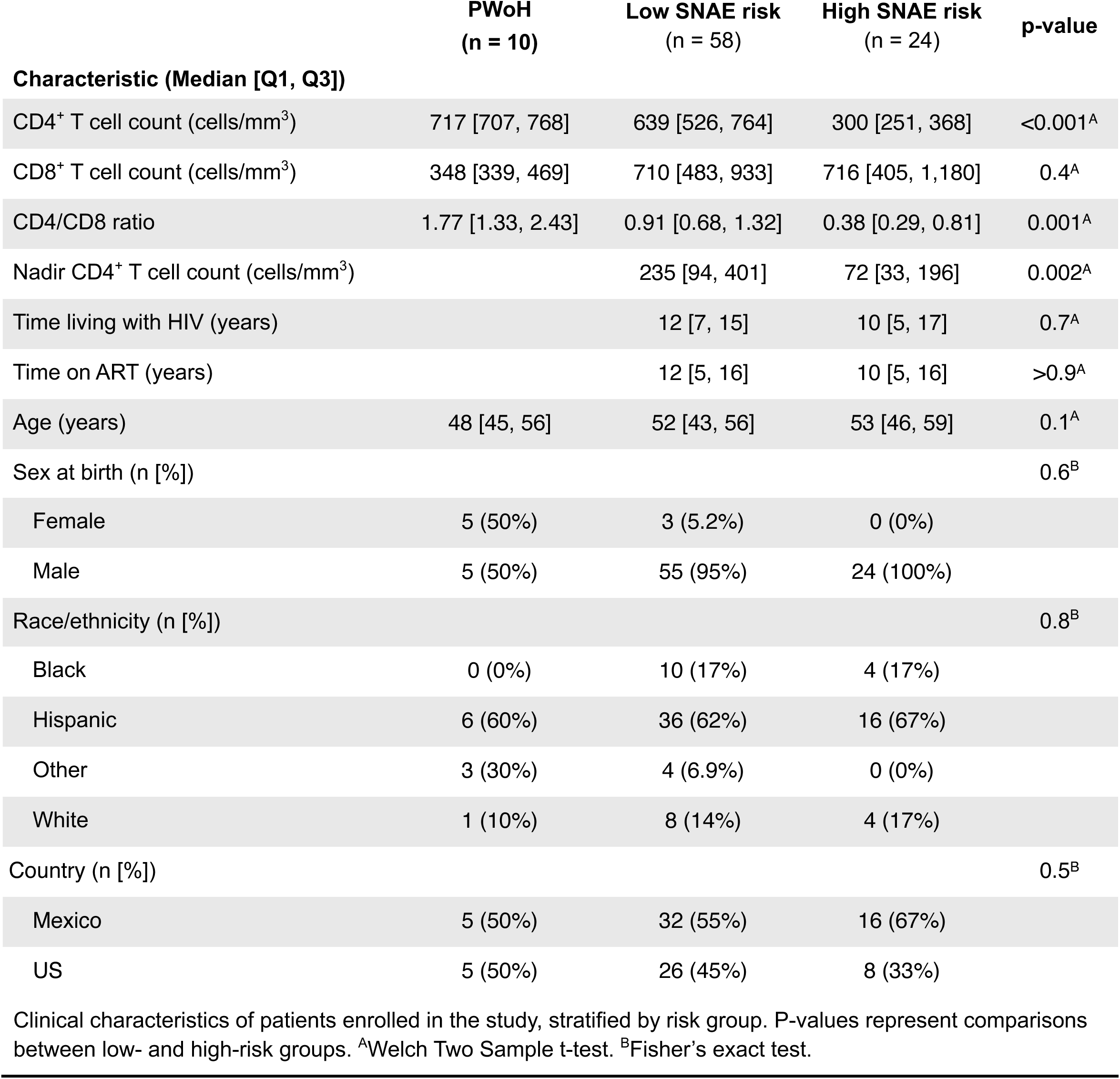
Baseline characteristics of enrolled participants stratified by risk groups.

|  | **PWoH**<br>**(n = 10)** | **Low SNAE risk**<br>**(n = 58)** | **High SNAE risk**<br>**(n = 24)** | **p-value** |
| --- | --- | --- | --- | --- |
| **Characteristic (Median [Q1, Q3])** |  |  |  |  |
| CD4<sup>+</sup> T cell count (cells/mm<sup>3</sup>) | 717 [707, 768] | 639 [526, 764] | 300 [251, 368] | <0.001<sup>A</sup> |
| CD8<sup>+</sup> T cell count (cells/mm<sup>3</sup>) | 348 [339, 469] | 710 [483, 933] | 716 [405, 1,180] | 0.4<sup>A</sup> |
| CD4/CD8 ratio | 1.77 [1.33, 2.43] | 0.91 [0.68, 1.32] | 0.38 [0.29, 0.81] | 0.001<sup>A</sup> |
| Nadir CD4<sup>+</sup> T cell count (cells/mm<sup>3</sup>) |  | 235 [94, 401] | 72 [33, 196] | 0.002<sup>A</sup> |
| Time living with HIV (years) |  | 12 [7, 15] | 10 [5, 17] | 0.7<sup>A</sup> |
| Time on ART (years) |  | 12 [5, 16] | 10 [5, 16] | >0.9<sup>A</sup> |
| Age (years) | 48 [45, 56] | 52 [43, 56] | 53 [46, 59] | 0.1<sup>A</sup> |
| Sex at birth (n [%]) |  |  |  | 0.6<sup>B</sup> |
| Female | 5 (50%) | 3 (5.2%) | 0 (0%) |  |
| Male | 5 (50%) | 55 (95%) | 24 (100%) |  |
| Race/ethnicity (n [%]) |  |  |  | 0.8<sup>B</sup> |
| Black | 0 (0%) | 10 (17%) | 4 (17%) |  |
| Hispanic | 6 (60%) | 36 (62%) | 16 (67%) |  |
| Other | 3 (30%) | 4 (6.9%) | 0 (0%) |  |
| White | 1 (10%) | 8 (14%) | 4 (17%) |  |
| Country (n [%]) |  |  |  | 0.5<sup>B</sup> |
| Mexico | 5 (50%) | 32 (55%) | 16 (67%) |  |
| US | 5 (50%) | 26 (45%) | 8 (33%) |  |

Clinical characteristics of patients enrolled in the study, stratified by risk group. P-values represent comparisons between low- and high-risk groups. <sup>A</sup>Welch Two Sample t-test. <sup>B</sup>Fisher's exact test.

### Multi-omic geographic differences in PWH on ART

Geographic and environmental factors shape the microbiome, metabolome, and immune landscape, yet most HIV omic studies have focused on cohorts from high-income countries. The global heterogeneity of the gut microbiome in PWH, and its intersection with metabolic and immune phenotypes, remains poorly characterized.

We first looked at plasma concentrations of cytokines, markers of gut epithelial barrier dysfunction, and microbial translocation including intestinal fatty acid binding protein (I-FABP), lipopolysaccharide binding protein (LBP), and soluble CD14 (sCD14) by principal component analysis (PCA). Country of enrollment was the largest, and only significant effect on the overall composition of these biomarkers when tested by PERMANOVA (R^2^ = 0.068, q = 0.020, Figure 1A). Numerous plasma proteins were differentially abundant between participants in different countries (Figure 1B). Compared to PWH from the US, those from Mexico had significantly higher concentrations of I-FABP (p < 0.001, q < 0.001) and sCD14 (p = 0.018, q = 0.1637). Cytokines involved in antigen presentation^15^, mucosal function^16^, and proliferation^17^ were also elevated in patients from Mexico including interleukin (IL)-12p70 (p < 0.001, q = 0.0028), IL-17A (p = 0.002, q = 0.0341), and IL-2 (p = 0.003, q = 0.0341). PWH from the US were characterized by increased concentrations of macrophage inflammatory protein (MIP)-1β (p < 0.001, q = 0.0086), IL-7 (p = 0.009, q = 0.0916), and IL-8 (p = 0.003, q = 0.0341).

**Figure 1.**
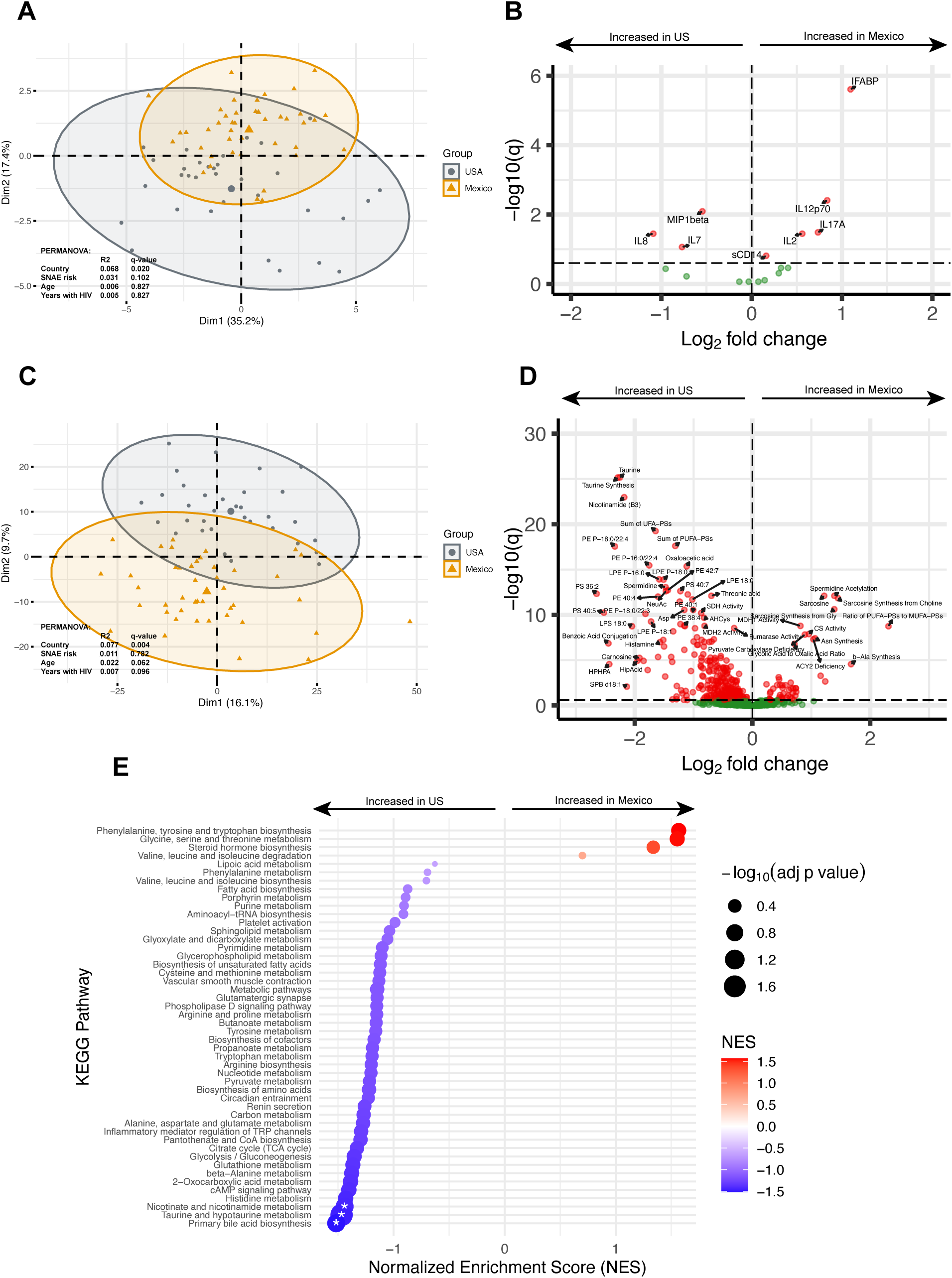
Differences in plasma proteins and metabolome between PWH enrolled in the US and Mexico. Principal component analysis of plasma protein concentrations, colored by country of enrollment and tested by PERMANOVA (A). Volcano plot of cytokines and gut barrier integrity markers differentially associated with country of enrollment (B). Principal component analysis of plasma metabolomic features, colored by country of enrollment and tested by PERMANOVA (C). Volcano plot of metabolomic features differentially associated with country of enrollment (D). GSEA of metabolomic data comparing KEGG pathway enrichment between PWH in US and Mexico, asterisks denoting FDR adjusted q-value ≤ 0.25.

We next evaluated the plasma metabolome of PWH from the US or Mexico using a comprehensive targeted LC-MS/MS panel of 1006 metabolites, including 751 lipids, 255 small molecules, and 273 sums or ratios for a total of 1,279 metabolomic features after filtering. Once again, PCA showed that country was the largest and only significant effect on the overall metabolomic profile composition (R^2^ = 0.077, q = 0.004, Figure 1C), followed by age. Differential analysis revealed 357 metabolomic features that were significantly different between PWH in US and Mexico at an adjusted p-value threshold of ≤0.25, and 236 at a threshold of ≤0.05, most of which were elevated in PWH from the US (Figure 1D, Supplemental Data 1). When looking at the classes of significantly different metabolites (q ≤ 0.25, Supplemental Figure 1), phospholipids were especially higher in PWH from the US, including phosphatidylethanolamines (PEs), phosphatidylcholines (PCs), phosphatidylglycerols (PGs), and phosphatidylserines (PSs), which represented 39.6% of the significantly different metabolites. The lyso-forms of these lipid classes were also elevated in PWH from the US including lyso-PEs and lyso-PCs. Sphingomyelins (SMs) were also exclusively elevated in PWH from the US. The primary metabolite class that was most represented by PWH in Mexico was acylcarnitines (ACs). When looking at individual metabolites, taurine, nicotinamide, oxaloacetic acid, spermidine, and acetylneuraminic acid were some of the top US-associated metabolites (p < 0.001 and q < 0.001 for all metabolites) while sarcosine (p < 0.001, q < 0.001), biotin (p < 0.001, q < 0.001), glycolic acid (p < 0.001, q = 0.001), octanoylcarnitine (p < 0.001, q = 0.012), and decanoylcarnitine (p < 0.001, q = 0.012) were some of the top Mexico-associated. Gene set enrichment analysis (GSEA) of the metabolite data showed that the KEGG pathways for primary bile acid biosynthesis, taurine/hypotaurine metabolism, and nicotinate/nicotinamide metabolism were all enriched in PWH from the US (q < 0.25, Figure 1E).

At the phylum level, ileal and rectal biopsies were primarily composed of Bacillota (formerly Firmicutes) and Bacteroidota (Supplemental Figure 2A). We did not observe any difference in alpha diversity metrics for both rectal or ileal biopsies between the US and Mexico (Figures 2A and 2B, respectively). Overall microbial composition of the rectal biopsies (Figure 2C), but not ileal biopsies (Figure 2D), was significantly associated with country of enrollment (PERMANOVA: R^2^ = 0.076, q = 0.02). Differential abundance analysis performed at the genus and species level showed increased abundances of *Ruminococcus, Eubacterium eligens group, Enterococcus, Veillonella, Escherichia-Shigella, Blautia stercoris,* and *Gemmiger formicilis* in the rectal biopsies of PWH from Mexico and increased abundances of *Agathobaculum butyriciproducens, Flavonifractor, Flavonifractor plautii, Enterocloster, Parabacteroides distasonis, and Parabacteroides* in the PWH from the US (q < 0.25, Figure 2E). When comparing the ileal biopsies, we observed increased abundance of *Eubacterium eligens group, Escherichia-Shigella*, Oscillospiraceae *NK4A214 group*, and *Haemophilus* in PWH from Mexico, whereas *Akkermansia*, *Akkermansia muciniphila*, *Desulfovibrio, Phascolarctobacterium faecium, Bacteroides uniformis, Parabacteroides distasonis, and Phascolarctobacterium* were all increased in PWH from the US (q < 0.25, Figure 2F).

**Figure 2.**
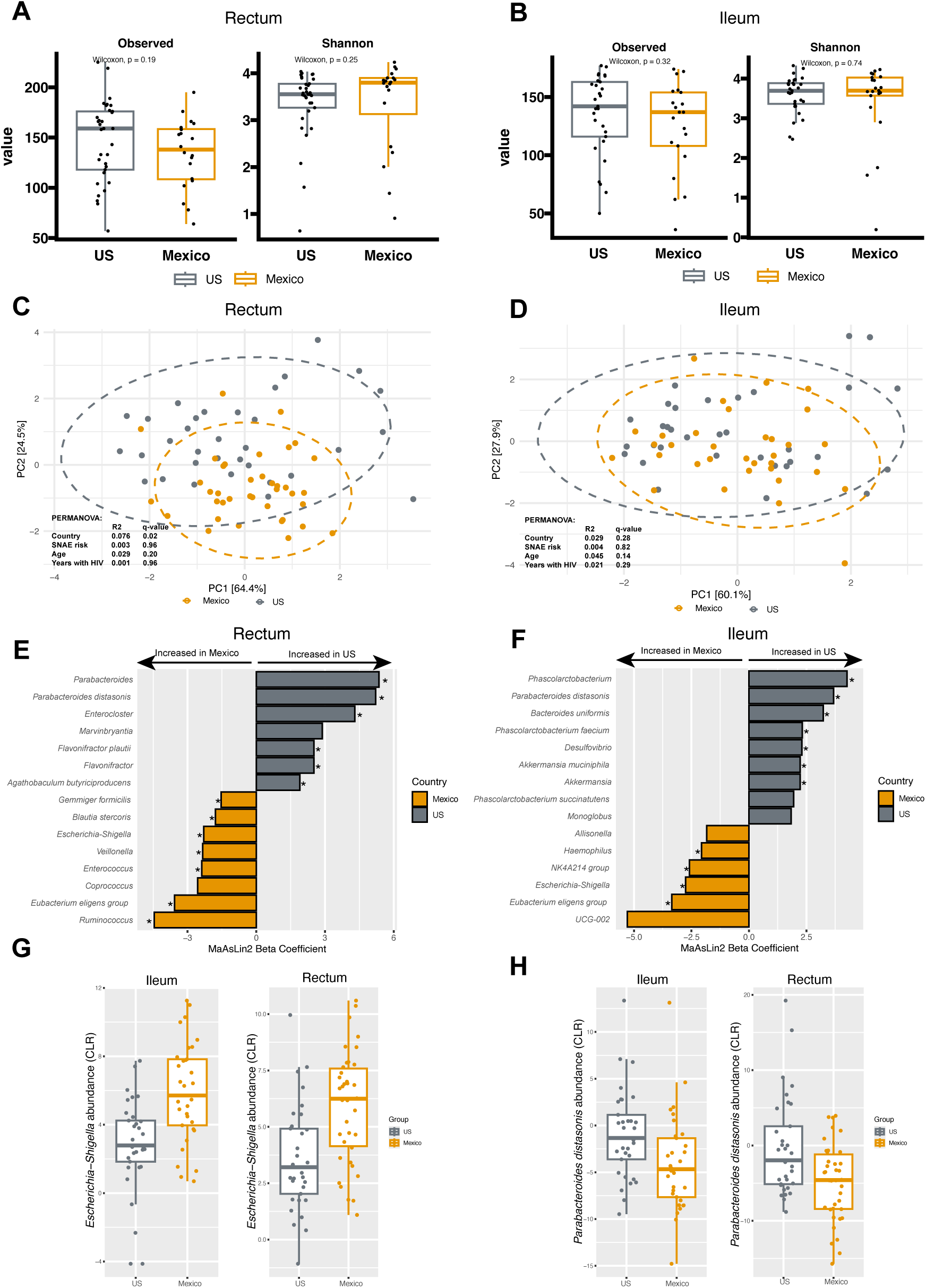
Differences in the ileal and rectal microbiomes of PWH enrolled from the US and Mexico. Alpha diversity metrics for rectum (A) and ileum (B) biopsies between PWH from the US and Mexico. Beta diversity of rectum (C) and ileum (D) biopsies, tested by PERMANOVA using the robust Aitchison distance of genus level taxa. Differential abundance of genus and species level taxa between the US and Mexico, tested by MaAsLin2 for rectum (E) and ileum (F) biopsies, showing the top 15 differentially abundant taxa with asterisks denoting FDR adjusted q-values ≤ 0.25. Representative box plots of *Escherichia-Shigella* (G) and *Parabacteroides distasonis* (F) in both the ileum and rectum.

Overall, some of the most striking differences between countries, were the consistent increased abundance of *Escherichia-Shigella* (Figure 2G) and decreased abundance of *Parabacteroides distasonis* (Figure 2H) in PWH from Mexico versus the US in both the ileal and rectal biopsies. We did not see any significant correlations between *Escherichia-Shigella* abundance in PWH from Mexico with mucosal cytokines or markers of gut barrier damage (Supplemental Figure 3).

### Systemic inflammation and metabolomic disruptions in PWH at high risk of SNAEs

Given the geographic differences between participants from the US and Mexico, we next examined whether there were biomarkers distinguishing PWH at low versus high risk for SNAEs, defined by CD4^+^ T cell counts and the CD4/CD8 ratio, that were independent of geographic effects. Such biomarkers could provide important insights into the processes driving SNAE risk. We first compared concentrations of plasma cytokines and markers of gut epithelial dysfunction. In contrast to country of enrollment, SNAE risk group did not have a significant effect on the overall plasma protein profile when tested by PERMANOVA (R^2^ = 0.031, q = 0.102, Figure 3A). When looking at individual markers, IL-6 was the only plasma protein we measured that was significantly increased in PWH at high risk for SNAEs (p = 0.021, q = 0.159, Figure 3B). Although not statistically significant after p-value adjustment, IL-10 was also elevated in the high-risk group (p = 0.039, q = 0.264). Of note, when stratified by country, IL-6 was non-significantly (q > 0.25) increased in PWH at high-risk of SNAEs (Supplemental Figure 4), and only reached statistical significance when the cohorts were combined.

**Figure 3.**
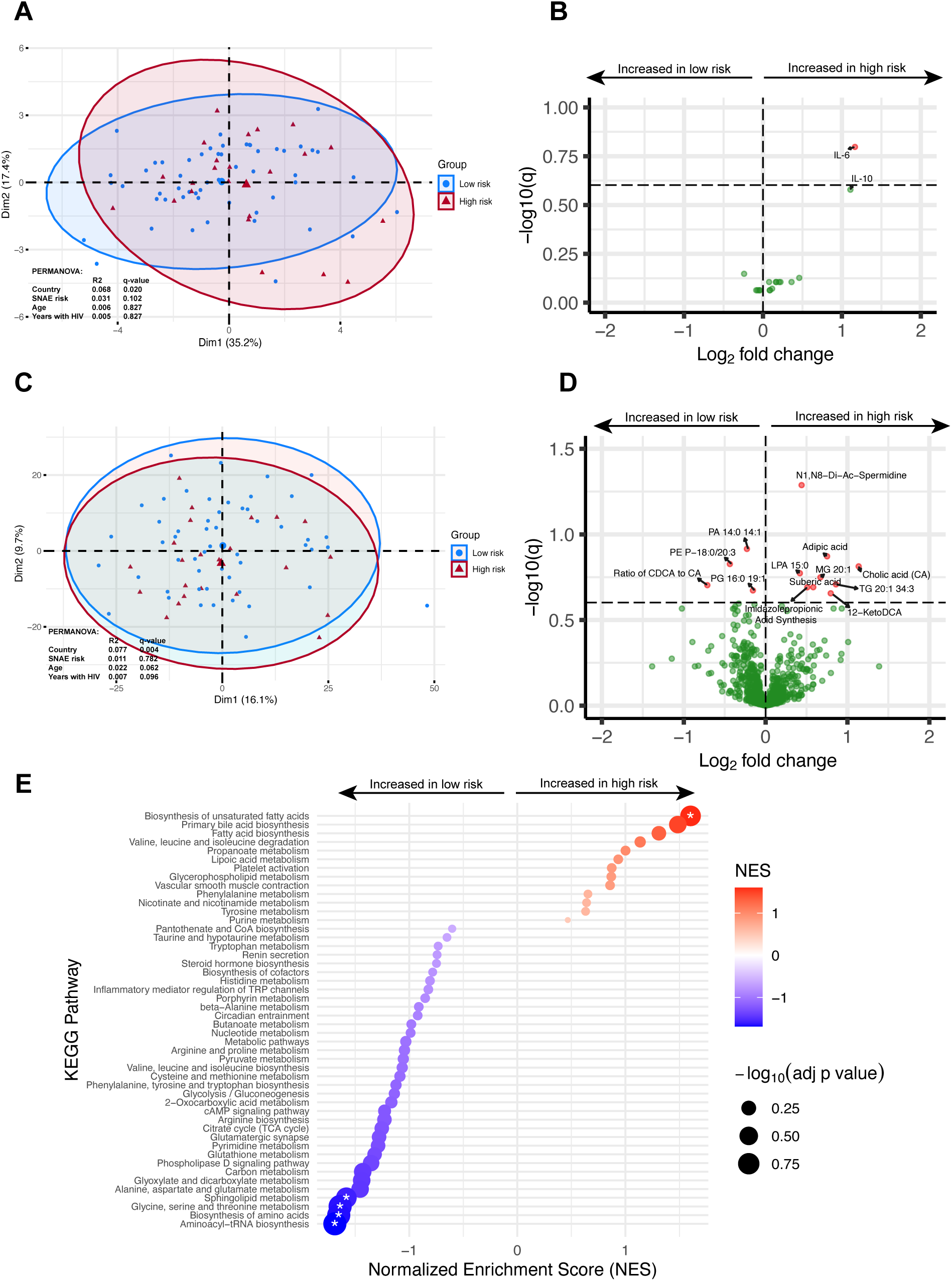
Differences in plasma proteins and metabolome between PWH at low or high risk of SNAEs. Principal component analysis of plasma protein concentrations, colored by SNAE risk group, tested by PERMANOVA (A). Volcano plot of cytokines and gut barrier integrity markers differentially associated with SNAE risk group (B). Principal component analysis of plasma metabolomic features, colored by SNAE risk group and tested by PERMANOVA (C). Volcano plot of metabolomic features differentially associated with SNAE risk group (D). GSEA of metabolomic data comparing KEGG pathway enrichment between low and high SNAE risk groups, asterisks denoting FDR adjusted q-value ≤ 0.25.

We next compared the plasma metabolome of individuals at low and high risk of SNAE development. Risk group did not have a significant effect on the overall profile of metabolomic features when tested by PERMANOVA (R^2^ = 0.011, q = 0.782, Figure 3C). Overall, we found 13 metabolomic features that were significantly different between SNAE risk groups (q < 0.25, Figure 3D). Individuals with high risk of SNAEs had increased plasma concentrations of dicarboxylic acids (adipic acid and suberic acid), bile acids (cholic acid and 12-ketodeoxycholic acid), triglycerides (TG 20:1 34:3), polyamines (N1,N8-Di-Ac-spermidine), monoglycerides (MG 20:1), and lysophosphatidic acids (LPA 15:0), whereas those at low risk had higher concentrations of specific phosphatidic acids (PA 14:0 14:1), PGs (PG 16:0 19:1), and PEs (PE P-18:0/20:3). Individuals at high risk also had a reduced ratio of chenodeoxycholic acid (CDCA) to cholic acid (CA), and higher activity of imidazole propionic acid synthesis as measured by the ratio of imidazole propionic acid to histidine. GSEA showed enrichment of KEGG pathways for aminoacyl-tRNA biosynthesis, biosynthesis of amino acids, glycine/serine/threonine metabolism, and sphingolipid metabolism in patients with low risk, and enrichment of biosynthesis of unsaturated fatty acids in patients at high SNAE risk (q < 0.25, Figure 3E).

### Reduced commensal bacteria in gut biopsies of PWH at high risk of SNAEs

We did not see any significant differences in alpha or beta diversity metrics in both the rectum and ileum of PWH with low versus high risk for SNAEs (Figure 4A-D). However, differential abundance analysis revealed several genus and species level bacterial taxa that were associated with SNAE risk group. In the rectal biopsies, *Ruminococcus gauvreauii group* was increased in PWH at high risk for SNAEs, while *Ruminococcus* and *Bacteroides uniformis* were both decreased (q < 0.25, Figure 4E). In the ileal biopsies, *Intestinibacter, Intestinibacter bartlettii,* Oscillospiraceae *NK4A214 group*, *and Collinsella bouchesdurhonensis* were increased in PWH at high risk of SNAEs, while *Akkermansia,* and *Akkermansia muciniphila* were decreased (q < 0.25, Figure 4F).

**Figure 4.**
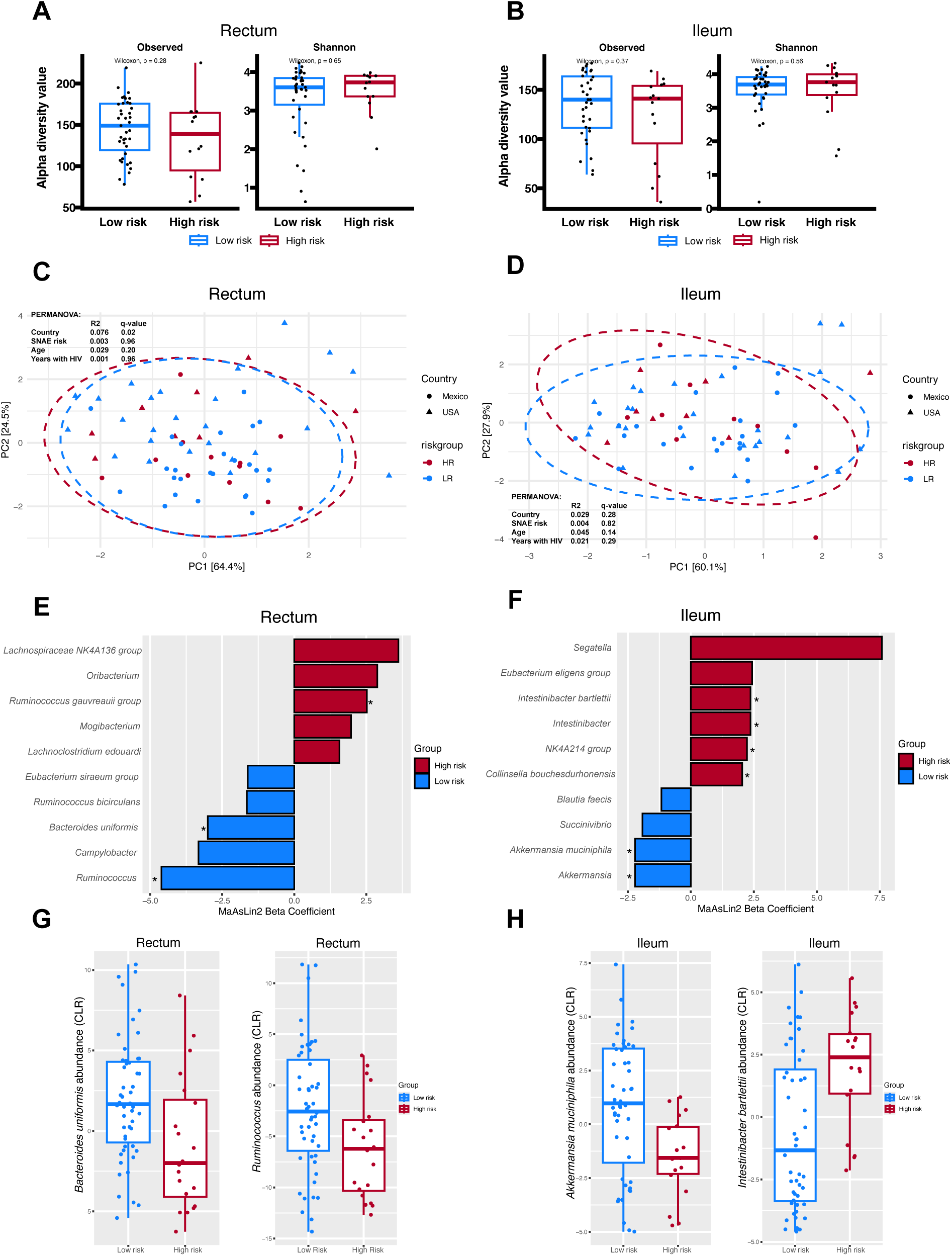
Differences in the ileal and rectal microbiomes of PWH at low or high risk of SNAEs. Alpha diversity metrics for rectum (A) and ileum (B) biopsies between SNAE risk groups. Beta diversity of rectum (C) and ileum (D) biopsies, tested by PERMANOVA using the robust Aitchison distance of genus level taxa. Differential abundance of genus and species level taxa between SNAE risk groups, tested by MaAsLin2 for rectum (E) and ileum (F) biopsies, showing the top 10 differentially abundant taxa with asterisks denoting FDR adjusted q-values ≤ 0.25. Representative box plots of differentially abundant taxa in the rectum (G) and ileum (F) biopsies.

Notably, *Segatella* (formerly part of the *Prevotella* genus) was increased in the ileal biopsies of the high risk SNAE group, though it did not reach statistical significance after adjusting for multiple comparisons (p = 0.016, q = 0.289, Figure 4F).

### Multi-omic machine learning discriminates between SNAE risk groups

We next applied supervised machine learning methods from the mixOmics R package to evaluate how well each ome (i.e., proteins, metabolome, ileum, rectum) classified SNAE risk, identify key discriminative variables, and examined cross-omic relationships. For single-ome models we used partial least squares discriminant analysis (PLS-DA) or sparse (s)PLS-DA when there were more variables than samples (Figure 5A). To integrate all omes, we used Data Integration Analysis for Biomarker discovery using Latent variable approaches for Omics studies (DIABLO, Figure 5B).

**Figure 5.**
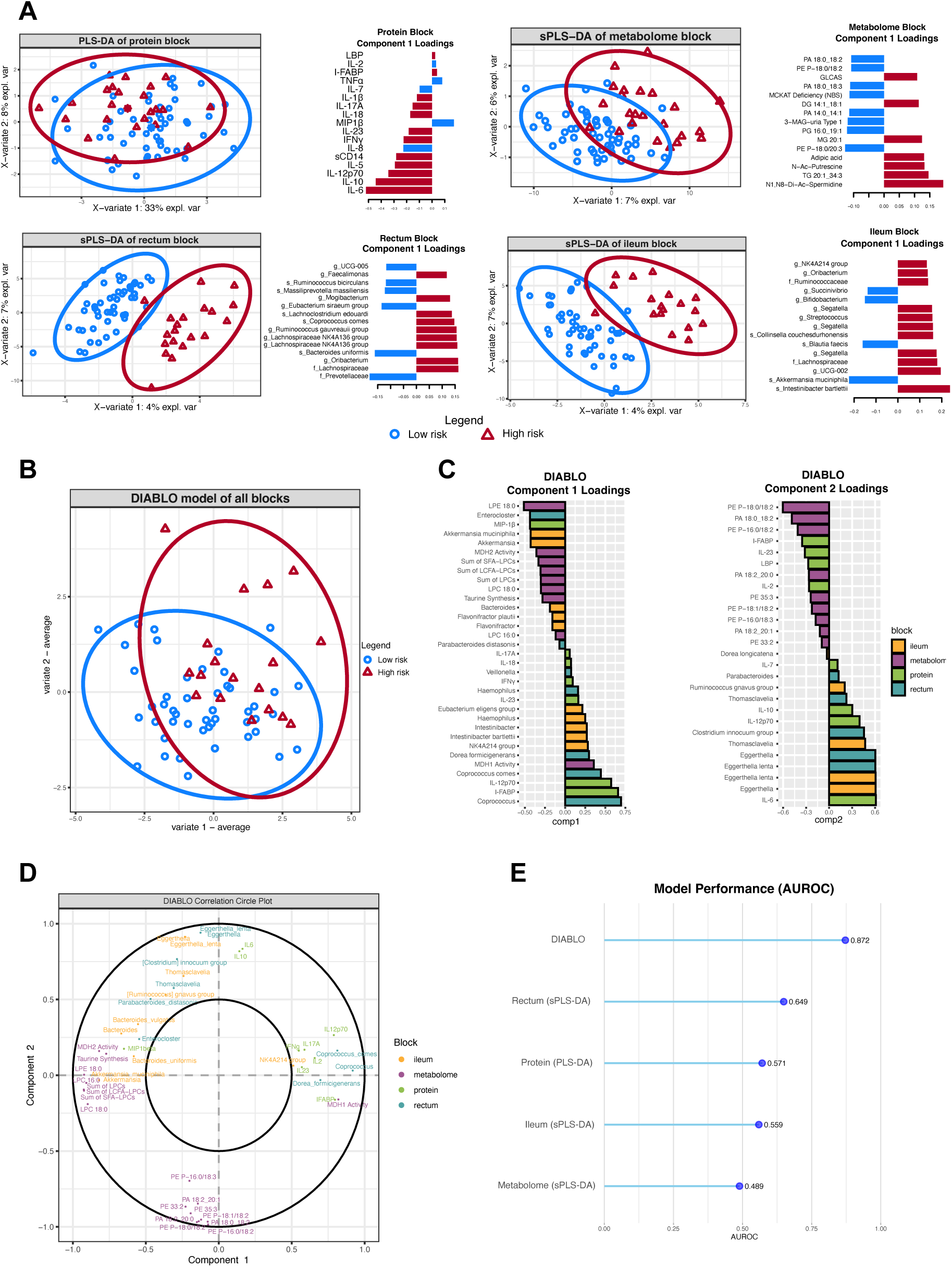
Supervised machine learning of SNAE risk groups. Scores plots and the top loading vectors on component one for the protein block PLS-DA, metabolome block (s)PLS-DA, rectum block (s)PLS-DA, and ileum block (s)PLS-DA (A). Loading vectors are colored by the group with the highest mean value. Scores plot of the integrated DIABLO model, combining all four data blocks, and showing the weighted average of components one and two according to their correlation with SNAE risk group (B). Top loading scores on components one and two for each block in the DIABLO model, bars are colored by the corresponding block the variable belongs to (C). Correlation circle plot of the DIABLO model, showing variables with correlations (>0.5) on components one and two (D). A variable’s location depicts both its correlation on each component and its correlation with other variables in proximity. Performance of each model in predicting SNAE risk based on AUROC, determined by 5-fold cross-validation repeated 50 times (E).

Many of the top discriminatory features for each model were also identified in our differential analysis. For example, IL-6 and N1,N8-Di-Ac-Spermidine were the top features for component one in the protein and metabolome models, both discriminatory for the high risk SNAE group. Similarly, *Akkermansia muciniphila* and *Intestinibacter bartlettii* were the top features in the ileum model, in the direction of low- and high-risk SNAE groups respectively. In the integrated DIABLO model, top features in the direction of PWH at high risk included rectum abundance of *Coprococcus* and *Coprococcus comes* as well as I-FAPB, IL-12p70 and MDH1 activity as measured by the ratio of malic to oxaloacetic acid (Figure 5C). Top features in the direction of PWH at low risk included LPE 18:0, rectum *Enterocloster* abundance, MIP1-β, and ileum abundances of *Akkermansia* and *Akkermansia muciniphila*. On component two, top features in the direction of PWH at high risk included IL-6 and both ileum and rectum abundances of *Eggerthella* and *Eggerthella lenta*, whereas features in the direction of PWH at low risk included the lipids PE P-18:0/18:2, PA 18:0/18:2, and PA P-16:0/18:2.

The ability of each model to discriminate between low- and high-risk groups was estimated by the mean AUROC from 5-fold cross-validation repeated 50 times. Individually, blocks performed modestly, with the rectal microbiome having the relative best performance (AUROC = 0.649, Figure 5E). In contrast, the integrated DIABLO model combining all omes achieved the top performance (AUROC = 0.872), indicating a strong ability to discriminate between low- and high-risk SNAE groups.

### Individual associations between omic features and T cell criteria

As our high-risk SNAE group included criteria for both CD4^+^ T cell counts and the CD4/CD8 ratio, we next sought to understand how each biomarker was correlated with individual T cell criteria. We performed partial Spearman correlations between each biomarker and CD4^+^ T cell counts, CD8^+^ T cell counts, and the CD4/CD8 ratio adjusting for the effect of country, age, and years living with HIV (Figure 6A). To reduce the number of comparisons, we focused only on biomarkers that were significantly different between our low- and high-risk groups. Notably, CD4^+^ T cell counts were negatively correlated with the ileum abundance of both *Intestinibacter bartlettii* (r = -0.405, p = 0.001, q = 0.011) and NK4A214 group (r = -0.41, p < 0.001, q = 0.011). These taxa were similarly negatively correlated with the CD4/CD8 ratio (r = -0.312, p = 0.014, q = 0.15 and r = -0.305, p = 0.016, q = 0.15 respectively). In the opposite direction, ileum abundance of *Akkermansia muciniphila* and rectum abundance of *Bacteroides uniformis* were positively correlated with the CD4/CD8 ratio (r = 0.264, p = 0.038, q = 0.164 and r = 0.248, p = 0.043, q = 0.164, respectively). In the metabolomic data, CD4^+^ T cell counts were negatively correlated with TG 20:1 34:4 (r = -0.359, p = 0.002, q = 0.012), imidazolepropionic acid synthesis (r = -0.269, p = 0.021, q = 0.089), and cholic acid (r = -0.253, p = 0.031, q = 0.107), while positively correlated with PA 14:0 14:1 (r = 0.274, p = 0.019, q = 0.089). The CD4/CD8 ratio was positively correlated with PE P-18:0/20:3 (r = 0.269, p = 0.021, q = 0.15). We did not observe any significant correlations with CD8^+^ T cell counts.

**Figure 6.**
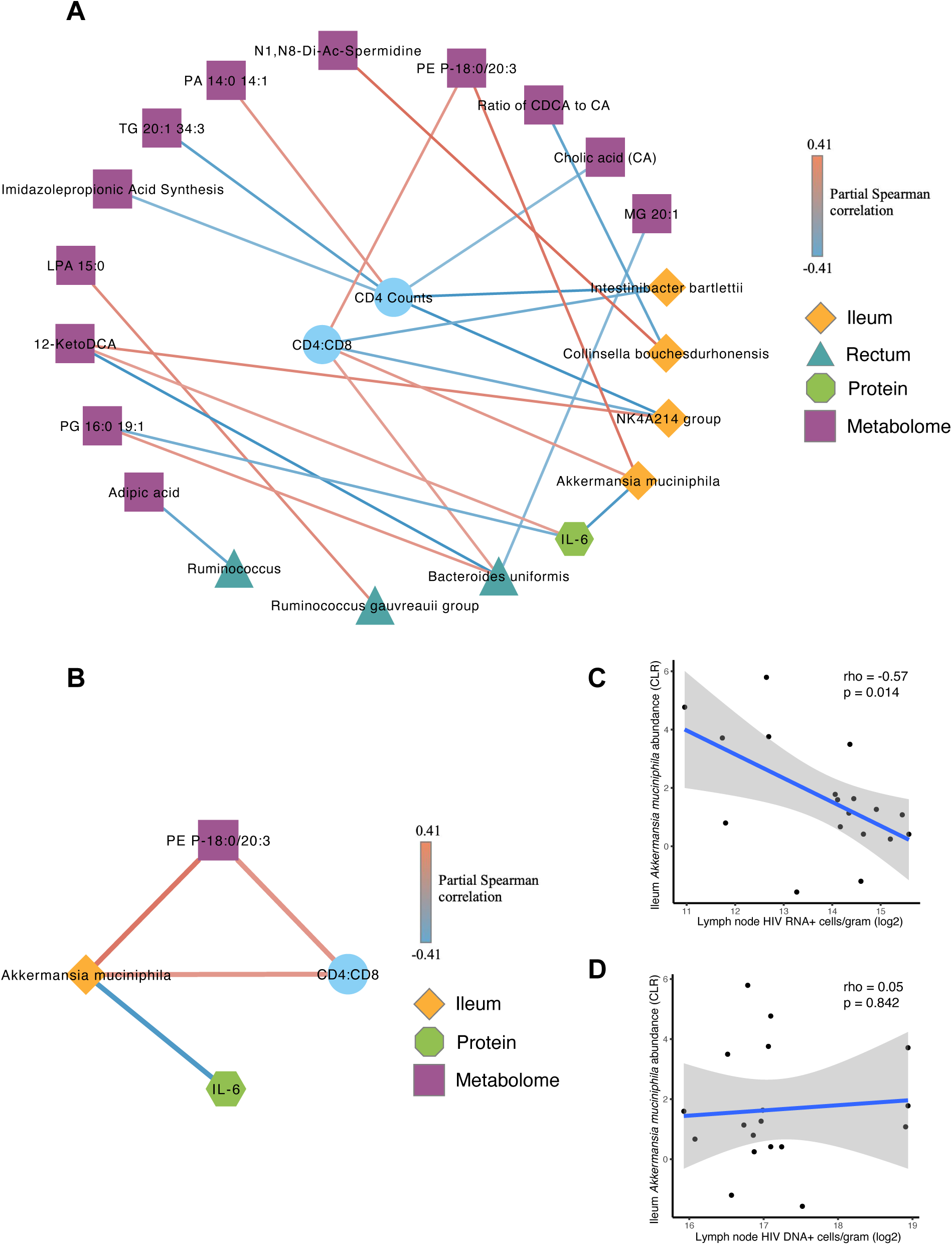
Partial correlations between variables of interest, CD4 counts, and the CD4/CD8 ratio. Network of partial Spearman correlations between variables that were significantly different between SNAE risk groups, adjusted for country, age, and years living with HIV (A). Selected correlations between bacterial taxa and the CD4/CD8 ratio with all connected nodes, highlighting potential mechanisms involved in driving immune activation (B). All correlations with an FDR adjusted q-value ≤ 0.25 and nominal p-value ≤ 0.05 are shown. Spearman correlations between ileum *Akkermansia muciniphila* abundance and HIV RNA^+^ (C) and DNA^+^ (D) cells/gram in lymph node tissue.

### Correlation analysis highlights potential role of *Akkermansia muciniphila* in reducing immune activation via IL-6 and the active HIV reservoir

We next performed partial correlations between these omic layers to understand how the relationships between them may be driving SNAE risk (Figures 6A and 6B). Interestingly, we observed a negative correlation between ileum *Akkermansia muciniphila* abundance and plasma IL-6 concentrations (r = -0.388, p = 0.002, q = 0.007), with the same pattern observed when stratified by country (Supplemental Figure 5). IL-6 was further positively correlated with the secondary bile acid 12-ketodeoxycholic acid (r = 0.247, p = 0.035, q = 0.23) and negatively correlated with PG 16:0 19:1 (r = - 0.302, p = 0.009, q = 0.121). Rectum *Bacteroides uniformis* abundance was similarly positively correlated with PG 16:0 19:1 (r = 0.269, p = 0.028, q = 0.242) and negatively correlated with 12-ketodeoxycholic acid (r = -0.415, p < 0.001, q = 0.019). Other notable correlations included a positive correlation between ileum *Collinsella bouchesdurhonensis* abundance and N1, N8-Di-Ac-Spermidine (r = 0.375, p = 0.003, q = 0.118), and a positive correlation between *Akkermansia muciniphila* and PE P-18:0/20:3 (r = 0.354, p = 0.005, q = 0.112).

Given the negative correlation we observed between *Akkermansia muciniphila* and plasma IL-6, we next asked whether ileal *Akkermansia muciniphila* abundance was associated with the HIV reservoir. For a subset (n = 20) of the US participants, we were able to obtain lymph node tissue and measured the quantity of HIV RNA-producing cells and proviral DNA^+^ cells as we have done previously^13,18^. Interestingly, ileal *Akkermansia muciniphila* abundance was negatively correlated with the number of HIV RNA^+^ cells/gram of lymph node tissue (Spearman, r = -0.57, p = 0.014, Figure 6C), though was not correlated with the number of HIV DNA^+^ cells/gram (Spearman, r = 0.05, p = 0.842, Figure 6D). Further, we did not observe a significant correlation between either IL-6 or IL-10 plasma concentrations and lymph node HIV reservoir measures (Supplemental Figure 6). Collectively, these findings suggest that *Akkermansia muciniphila* may reduce SNAE risk through a potential mechanism involving suppression of IL-6 and HIV RNA production in latently infected cells.

### Multi-omic SNAE risk signatures contextualized in people without HIV

To assess whether the observed differences between SNAE risk groups reflected true biological signals rather than confounding between groups, we compared the significantly different variables to a group of people without HIV (n = 10) from both the US and Mexico (n = 5 per country) using linear regression adjusted for age and country. Although limited statistical power likely hindered significance in many comparisons with PWoH, nearly all variables followed the expected ordinal trend, whereby features elevated in the high-risk relative to the low-risk group were also higher than in PWoH and vice versa. For example, IL-6 was non-significantly elevated in the high-risk group relative to PWoH (β = 0.85, p = 0.272) and non-significantly reduced in the low-risk group (β = -0.31, p = 0.657, Figure 7A). Among metabolomic features, N1,N8-Di-Ac-Spermidine, TG 20:1 34:3, and imidazole propionic acid synthesis were each significantly higher in the high-risk group compared to PWoH (p < 0.05, Figure 7B).

**Figure 7.**
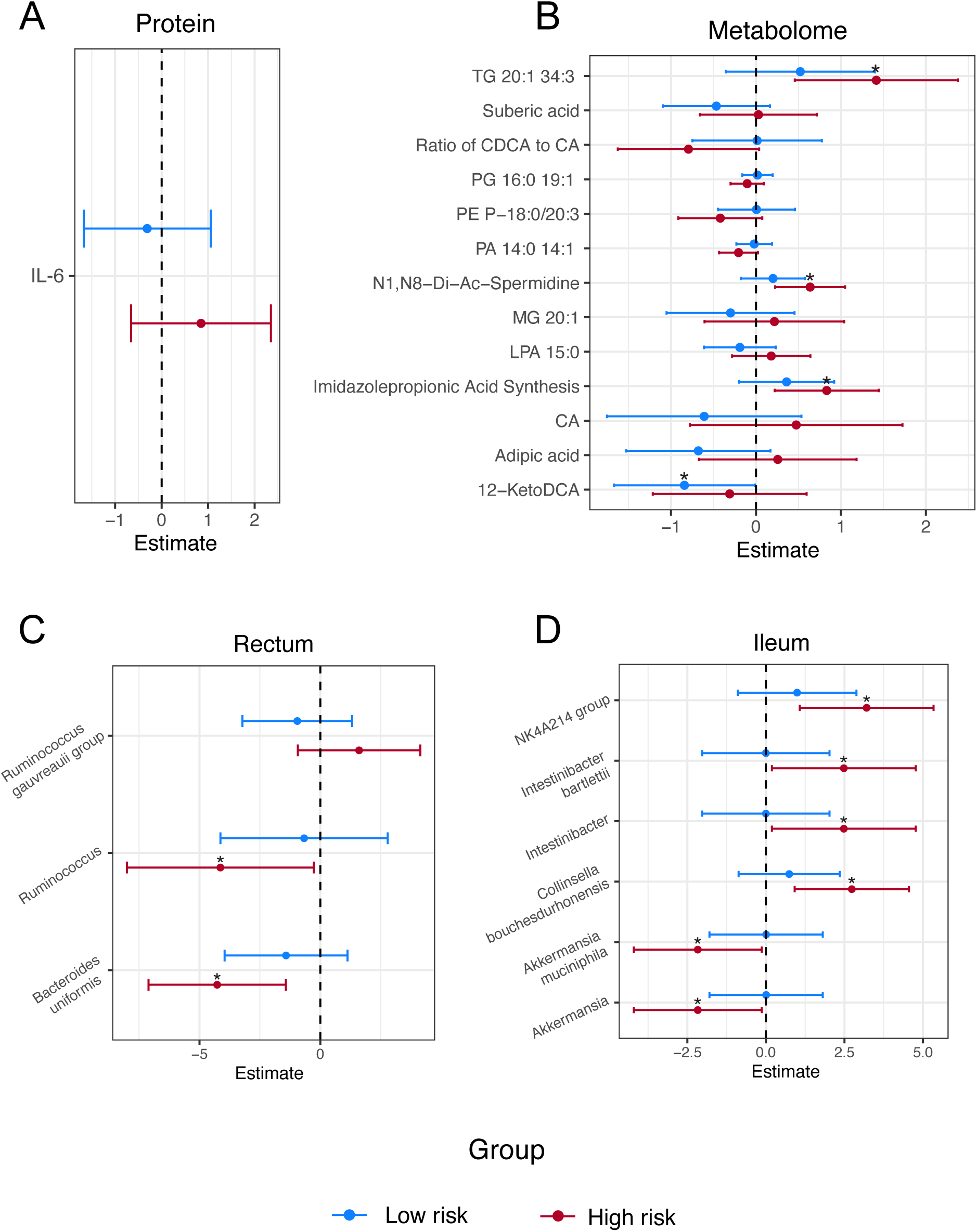
Comparison of SNAE risk groups to people without HIV. Variables that were significantly different between SNAE risk groups (q < 0.25) were compared against a PWoH reference group (n = 10, n = 5 per country) using linear regression adjusted for age and country. Panels display the linear model estimate and 95% confidence interval for each SNAE risk group relative to PWoH for log2-transformed plasma IL-6 (A), log2-transformed metabolomic features (B), and CLR-transformed rectal (C) and ileal (D) microbiome taxa abundances, with microbiome estimates derived from MaAsLin2. Asterisks (*) denote nominally significant differences from PWoH (p < 0.05).

Microbiome trends were similarly conserved, with *Ruminococcus* and *Bacteroides uniformis* significantly reduced in the high-risk group relative to PWoH (p < 0.05, Figure 7C). In the ileum, *Akkermansia* and *Akkermansia muciniphila* were likewise reduced in the high-risk group, while *NK4A214 group*, *Intestinibacter*, *Intestinibacter bartlettii*, and *Collinsella bouchesdurhonensis* were all significantly increased compared to PWoH (p < 0.05, Figure 7D). The only exception to these patterns was 12-ketodeoxycholic acid, which was lowest in the low-risk group, followed by high risk and PWoH. Collectively, the significant differences observed between the high-risk group and PWoH in key metabolomic features and microbiome taxa, combined with the directional consistency of nearly all remaining variables, support the biological plausibility of these signatures as indicators of SNAE risk.

## Discussion

In this multi-site analysis, we first examined country-level differences within our HIV cohort. Country of enrollment was the largest determinant of variation across plasma proteins, the plasma metabolome, and the gastrointestinal microbiome. Participants from Mexico exhibited higher plasma cytokines involved in adaptive and mucosal immunity (IL-12p70, IL-17A) and markers of gut epithelial injury and microbial translocation (I-FABP, sCD14), collectively suggesting greater mucosal immune activation in this cohort. IL-12p70 and IL-17A support Th1-driven antiviral responses and gut barrier integrity^19,20^, functions that are impaired in HIV infection. These immune features coincided with substantially greater mucosal abundance of *Escherichia–Shigella,* a common intestinal pathogen in developing countries^21^, in both ileal and rectal biopsies, which we suspected could be leading to increased mucosal inflammation in PWH from Mexico, although this taxon did not directly correlate with these plasma markers. In contrast, US participants had higher levels of MIP-1β, IL-8, and IL-7, indicating distinct patterns of immune activation between countries that may reflect differences in HIV care, environmental exposures or co-infections, however CMV serostatus did not differ between them.

The ileal and rectal microbiomes were dominated by Bacillota and Bacteroidota, consistent with previous reports^22^. Rectal biopsy composition differed most strongly by country, although alpha diversity metrics did not vary. Given differences in baseline characteristics between countries (e.g., race, sex) and unmeasured effects such as diet, it is not surprising that the gastrointestinal microbiomes of these countries would significantly differ. Some of the most notable differences we observed were the increased abundance of *Escherichia–Shigella* and *Eubacterium eligens group* in participants from Mexico in both biopsy sites. In comparison, US participants displayed increased *Parabacteroides distasonis* in both biopsy sites. *Escherichia-Shigella* is, whereas *Eubacterium eligens group* has been linked to reduced visceral fat accumulation and protective for pulmonary arterial hypertension^23,24^.

Country effects were most pronounced in the plasma metabolome. US participants had significantly higher concentrations of numerous lipid classes. Compared to Mexican men living in Mexico, Mexican men living in the US are at a greater risk of obesity, abdominal obesity, and diabetes, which may be reflective of the lipid changes we see^25^. Consistent with this, sphingomyelins, phosphatidylcholines, and lyso-phosphatidylcholines, all previously linked to obesity^26^, were enriched in our US cohort. These differences could potentially reflect differences in BMI between countries.

However, BMI was unavailable as a covariate in our analysis and is a limitation in interpreting these findings. US-associated metabolites in our study, such as taurine and oxaloacetic acid, also aligned with a previous study comparing the plasma metabolome of US-born versus foreign-born (primarily from Mexico) Hispanic men^27^, attributed to dietary differences. Mexico participants exhibited higher levels of medium-chain acylcarnitines (C8, C10, C10:1, C12:1); several of which have been shown to increase following moderate-intensity exercise, promote lipid oxidation, and decrease with oral food intake^28^, potentially reflecting differences in physical activity, energy metabolism, and dietary practices between cohorts.

Understanding how geography influences multi-omic profiles in PWH is essential for capturing the global impact of HIV. We therefore examined whether PWH from the US and Mexico who were at high risk for SNAEs, based on low CD4^+^ T cell counts and CD4/CD8 ratios, shared consistent biomarkers. Although SNAE risk explained far less variance than country of enrollment, a clear pattern emerged involving systemic inflammation, microbial dysbiosis, and metabolomic alterations suggestive of mitochondrial dysfunction.

First, we observed increased plasma IL-6 concentrations in participants at high risk for SNAEs, which was consistent when stratified by country, in agreement with previous studies^29,30^, and followed the expected ordinal trend relative to PWoH, collectively indicating an elevated level of systemic inflammation in these individuals. Further, IL-6 was a topmost discriminatory variable in the supervised models built to discriminate between low- and high-risk SNAE groups. Similarly, other cytokines including IL-10 and IL-12p70 were non-significantly elevated in the high-risk group but were top cytokines in contributing to class separation. Given the known increase of IL-10 with HIV progression^31^ and its dual role in moderating inflammation^32,33^ and supporting reservoir persistence^34^, this may reflect a compensatory yet potentially pathogenic response.

Among plasma metabolites, dicarboxylic acids, especially suberic and adipic acid, were markedly increased in the high-risk SNAE group. These products of ω-oxidation accumulate when mitochondrial β-oxidation is impaired or overloaded, as observed in fatty-acid oxidation disorders such as medium-chain acyl-Coenzyme A(CoA) dehydrogenase deficiency^35^. Because β-oxidation normally fuels ATP generation via the electron transport chain, diversion toward ω-oxidation and the buildup of dicarboxylic acids indicate mitochondrial stress^36,37^. Thus, higher adipic and suberic acids in high-risk PWH support the presence of disrupted β-oxidation and mitochondrial dysfunction. Several mechanisms could contribute to this dysfunction. Carnitine deficiency, which is more common in PWH^38^, can restrict β-oxidation by limiting fatty acid transport into mitochondria. Inflammation may further impair β-oxidation by suppressing PGC-1α, a key regulator of mitochondrial biogenesis and metabolic gene expression^39^. Notably, the HIV Tat protein has been shown to downregulate PGC-1α, linking viral activity to mitochondrial defects observed in HIV-associated neurocognitive disorder^40^. Elevated adipic and suberic acids may also exert direct cellular effects by inducing DNA damage in leukocytes^35^, a process associated with CD4⁺ T cell apoptosis, immune senescence, and the heightened inflammaging characteristic of PWH^41^. Importantly, these dicarboxylic acids are increased even in treatment-naïve PWH compared to healthy controls^42^, suggesting that their accumulation is not driven by ART.

Additional metabolites elevated in high-risk PWH included N1,N8-diacetylspermidine and primary bile acid CA. N1,N8-diacetylspermidine has been identified as a tumor biomarker associated with poor cancer prognosis^43,44^, potentially reflecting the increased malignancy risk in this group. CA and the CA/CDCA ratio, both linked to liver fibrosis and inflammation^45,46^, were also higher in high-risk participants, and CA negatively correlated with CD4^+^ T cell counts, suggesting a link between primary bile acids and T cell reconstitution. Further, imidazole propionic acid, an exclusive microbially derived metabolite derived from histidine, has been linked to dysbiosis and increased risk of cardiovascular disease^47^ and type 2 diabetes^48^. Here, imidazole propionic acid synthesis (ratio of imidazole propionic acid to histidine) was increased in the high-risk group and was also negatively correlated with both CD4^+^ T cell counts.

Notably, N1,N8-diacetylspermidine, TG 20:1 34:3, and imidazole propionic acid synthesis were also significantly elevated in the high-risk group relative to PWoH, further supporting their relevance as markers of heightened SNAE risk. Several lipids were elevated in the low-risk group, most notably PE P-18:0/20:3, which positively correlated with both the CD4/CD8 ratio and ileal *Akkermansia muciniphila* abundance. This finding is of particular interest given that a separate PE lipid derived from the *A. muciniphila* cell membrane (PE a15:0-i15:0) was recently shown to promote homeostatic immune responses through low-level TLR2-TLR1 signaling^49^, raising the possibility that other PE lipids associated with *A. muciniphila* have similar immunomodulatory effects.

The ileal and rectal microbiomes were similar between low- and high-risk SNAE groups, with no significant differences in alpha or beta diversity. However, several taxa differed in abundance, with notable reductions in commensal, short-chain fatty acid (SCFA) producers in the high-risk group, including *A. muciniphila* in the ileum and *Bacteroides uniformis* and *Ruminococcus* in the rectum, both of which were also significantly reduced relative to PWoH. These organisms are associated with anti-inflammatory and beneficial effects, with of *A. muciniphila* and *B. uniformis*, are being developed as next-generation probiotics^50,51^. Both these taxa positively correlated with the CD4/CD8 ratio, whereas ileal *Intestinibacter bartlettii*, previously linked to irritable bowel syndrome^52^, was negatively correlated with CD4^+^ counts and CD4/CD8 ratios. Notably, *A. muciniphila* abundance was also inversely associated with plasma IL-6 and HIV RNA-producing cells in lymph nodes, suggesting a potential role in limiting systemic inflammation and the active HIV reservoir. The lack of a correlation between *A. muciniphila* abundance and proviral DNA+ cells in lymph nodes suggests that the inverse correlation with HIV RNA-producing cells is related to an effect on latency reversal rather than the size of the latent reservoir.

Probiotic or prebiotic interventions, especially in PWH at high risk of SNAEs, represents a potential strategy to improve immune recovery and mitigate comorbidity development. The use of *A. muciniphila* is particularly promising, given its link to a wide range of beneficial host effects relevant to HIV-associated inflammation including mitochondrial dysfunction, hepatic steatosis, and diabetes, in addition to a growing body of mechanistic evidence supporting these effects^53–60^. No human studies have directly tested *A. muciniphila* supplementation in PWH, though a recently completed clinical trial (NCT04058392) utilized a prebiotic specifically meant to increase *A. muciniphila* abundance in PWH with a CD4/CD8 ratio <1, with the goal of reducing inflammation in these individuals. Supporting our observations in PWH, *A. muciniphila* has repeatedly been shown to inversely correlate with IL-6^61–63^, possibly via a process that inhibits IL-6/JAK/STAT signaling^64,65^. JAK/STAT inhibitors such as tofacitinib and ruxolitinib have been shown to reduce HIV replication in latently infected cell lines^66,67^, thus it is conceivable that an *A. muciniphila-*mediated effect on JAK/STAT signaling could explain the negative correlation we observed with HIV RNA-producing cells in the lymph nodes. An alternative explanation is that increased *A. muciniphila* abundance reflects enhanced gut barrier integrity and reduced microbial translocation, however we did not observe differences in LBP, sCD14, or I-FABP between low- and high-risk groups. Together with our findings, these data suggest *A. muciniphila* as a compelling candidate for a microbiome-based intervention to mitigate systemic inflammation and lower SNAE risk in PWH, supporting the need for clinical evaluation.

This study should be interpreted in the context of its limitations. First, the lack of metadata on covariates including sexual orientation, BMI, diet, co-infections (beyond CMV), medication use, and comorbidities, represents a major constraint on the interpretation of the multi-omic readouts we measured, particularly given the well-established influence of diet on both metabolomic and microbiome profiles. Second, although most participants received similar ART regimens, the remaining individuals were sparsely distributed across many different ART combinations, preventing meaningful assessment of ART-specific effects. Further, while CMV serostatus was positive for nearly all patients, the absence of quantitative CMV replication data (e.g., IgG titers or CMV DNA) and other markers of systemic inflammation (e.g., C-Reactive protein) limits our ability to fully evaluate inflammatory drivers, including the contribution of CMV to IL-6. Our SNAE risk groups were also limited by a modest sample size composed almost entirely of men, reducing generalizability. Given the exploratory nature of this work, we also opted to use a relaxed p-value threshold for highly dimensional analyses, requiring an FDR adjusted q-value ≤ 0.25 to prioritize hypothesis generation. Finally, our composite SNAE risk definition used CD4^+^ T cell counts and CD4/CD8 ratios as surrogate markers in the absence of longitudinally observed SNAE data. Future studies may benefit from investigating the biomarkers we observed with longitudinal SNAE data to associate them with specific health outcomes. These T cell criteria additionally represent bulk rather than antigen-specific populations, limiting our ability to disentangle more granular associations between immune cells and inflammatory biomarkers. Notwithstanding these limitations, many of our findings are consistent with previous research and support biologically plausible conclusions.

In summary, this cross-site multi-omic analysis revealed geography as the dominant driver of variation among PWH, with distinct inflammatory, metabolic, and microbial signatures observed between cohorts from the US and Mexico. Despite these differences, individuals at high risk for SNAEs shared a consistent pattern of systemic inflammation, mitochondrial dysfunction, and loss of key commensal taxa. Elevated plasma IL-6, increased ω-oxidation products (adipic and suberic acids), and accumulation of bile acid and polyamine metabolites indicated persistent immune activation, hepatic stress, and dysfunctional mitochondrial β-oxidation. Along with reductions in mucosal SCFA producers *A. muciniphila*, *B. uniformis*, and *Ruminococcus* suggests that microbiome disruption may contribute to this phenotype. In particular, *A. muciniphila* stands out as a mechanistically validated commensal that promotes mucosal integrity, enhances mitochondrial metabolism, and attenuates hepatic and systemic inflammation in experimental models^53,54,68^. The depletion of *A. muciniphila* and its negative association with IL-6 and the active HIV reservoir underscores its potential therapeutic relevance. Exploring *A. muciniphila*-based therapeutics may represent a promising strategy to reduce inflammation and improve long-term health outcomes in PWH.

## Methods

### Sex as a biological variable

Sex was considered a biological variable. The study protocol permitted enrollment of all eligible participants regardless of sex, gender, race, or ethnicity. However, the cohort enrolled in this study was predominantly male.

### Participant enrollment

This analysis included 82 PWH from Mexico (n = 48) and the US (n = 34) enrolled in a study led by the University of Minnesota (UMN) between July 2021 and December 2023. Participants were enrolled and had their sample collection procedures performed at either the Center for Research in Infectious Disease (CIENI) of the National Institute of Respiratory Diseases (INER) in Mexico City, Mexico, or UMN in Minneapolis, US. Participants were required to be ≥ 18 years of age, laboratory-confirmed for HIV-1 infection per medical records, stable on an ART regimen for over 12 months, with plasma HIV RNA <48 copies/mL (isolated single blips up to 200 copies/mL were allowed if preceded and followed by undetectable viral load determinations), and within institutional normal ranges for screening tests (complete blood cell count and metabolic panel). Exclusion criteria included pregnancy or breastfeeding, having a BMI ≥ 30 kg/m^2^, and not being a suitable candidate for an inguinal LN biopsy (e.g., current use of anticoagulants, ≥3 LN biopsies in the past). No participants were taking antibiotics at enrollment. An additional reference group of PWoH (n = 10; n = 5 per country) was recruited from the same sites and matched by country of enrollment for contextual comparison of SNAE risk-associated signatures. PWoH were required to meet the same age and general health eligibility criteria but had no laboratory-confirmed HIV infection and were not on ART.

### SNAE risk group definition

The CD4^+^ T cell count and CD4/CD8 ratio have been shown to independently predict the risk of SNAEs^10^, with lower CD4+ counts and CD4/CD8 ratios being associated with higher SNAE risk. In our analysis, PWH were grouped as having a high risk or low risk for SNAEs based on these T cell criteria. High SNAE risk was defined as having a CD4^+^ T cell count <350 cells/µL or a CD4/CD8 ratio <0.4 regardless of CD4+ count after 2 years of ART, which are criteria associated with a nearly threefold increased risk of SNAE over the next 5 years. Any participants who did not meet these criteria were defined as having low SNAE risk.

### Sample collection

Blood was collected in EDTA tubes prior to either colonoscopy or lymph node biopsy with instructions to fast. Within 30 minutes of collection, plasma was isolated by centrifugation and stored at -80°C. Colonoscopies were conducted at the Endoscopy Center of the UMN Medical Center (UMMC) and at CIENI-INER under mild-to-moderate sedation and after overnight bowel preparation. Pinch biopsies were done in the ileum and rectum (Radial Jaw 4 Biopsy Forceps, Jumbo with Needle, 3.2 mm; Boston Scientific, Marlborough, MA, USA), placed in Qiagen Allprotect solution, and stored at - 80°C. Excisional inguinal LN biopsies were completed for a subset of US participants by UMN surgeons. The groin area was scrubbed with an antiseptic solution, and local anesthetics were injected to numb the inguinal area. A 1-inch skin incision was made over the lymph node. After the lymph node was removed, the incision was closed with dissolvable stitches. Participants were observed by the study team for a minimum of 2 hours post-procedure before discharge. All procedures and sample collections happened within a 2-week period.

### Plasma protein measurements

Concentrations of cytokines were measured in EDTA plasma using a 13-plex (IL-1β, IL-2, IL-5, IL-6, IL-7, IL-8, IL-10, IL-12p70, IL-17A, IL-23, TNFα, IFNγ, and MIP-1β) Luminex panel (Cat. HSTCMAG-28SK, MILLIPLEX) and by ELISA (IL-18) (Cat. DY318-05, R&D systems, MN, USA). Biomarkers for gut barrier damage and microbial translocation were measured in EDTA plasma by ELISA including I-FABP (Cat. DFBP20, R&D systems, MN, USA), sCD14 (Cat. QK383, R&D systems, MN, USA), and LBP (Cat. Ab279407, Abcam, Cambridge, UK). All processing was performed according to the kit manufacturer’s instructions. Values below the limit of detection for each analyte were replaced with the lowest detectable concentration divided by two. CMV IgG serostatus was determined qualitatively in plasma using a commercially available ELISA according to manufacturer’s instructions (Cat. KA6698, Abnova, Taipei City, Taiwan).

### Plasma metabolomics

The plasma metabolome was characterized in EDTA plasma using the commercially available MxP Quant 1000 kit (biocrates, Austria). Lipids were measured by flow injection analysis-tandem mass spectrometry (FIA-MS/MS) using a 5500 QTRAP instrument (AB Sciex, Darmstadt, Germany) with an electrospray ionization (ESI) source, and small molecules were measured by liquid chromatography-tandem mass spectrometry (LC-MS/MS) using a 5500+ Triple Quad™ instrument (AB Sciex, Darmstadt, Germany). Briefly, samples were loaded onto 96-well plates containing inserts impregnated with internal standards. For small molecule determination, derivatization was performed using a phenyl isothiocyanate or 3-nitrophenylhydrazine solution, followed by extraction with an organic solvent. Extracts were then analyzed by LC-MS/MS using multiple reaction monitoring. All concentrations were quantified using the biocrates WebIDQ software. Metabolites that were present in less than 50% of samples were removed from the analysis, and any values that were below the limit of detection were replaced with the lowest detectable concentration divided by two. A comprehensive list of the metabolites used in this analysis are provided (Supplemental Data 2), as well as any sum or ratio features (Supplemental Data 3).

### HIV reservoir quantification

The HIV reservoir was quantified in 4% paraformaldehyde-fixed lymph node biopsies as previously described^13,69,70^. Briefly, Five-to-ten 5 µm sections separated by 20 µm were analyzed by RNAscope 2.5 (Advanced Cell Diagnostics) using in situ hybridization with HIV specific probes from Advanced Cell Diagnostics to identify Clade B viruses (RNA anti-sense Cat. 416111 and DNA sense Cat. 425531). Quantitative image analysis was then used to determine the number of HIV RNA^+^ and DNA^+^ cells per tissue area.

### Microbiome analysis

DNA was extracted from ileal and rectal biopsies using the Qiagen PowerSoil Pro DNA extraction kit. The V4 rRNA region was amplified from template DNA using the 515F-806R primer set described in the earth microbiome project^71^, using a 30-cycle polymerase chain reaction (PCR). PCR products were then subjected to a second 10-cycle PCR to attach Illumina sequencing primer-compatible DNA regions as well as individual barcodes for each sample. Samples were all uniquely dual-indexed, as previously described^72^. Sequencing libraries were loaded onto an Illumina NextSeq using a 2x300 P2 flow cell and sequenced to an average depth of 79,756 reads per sample.

Variable region primers were removed from the demultiplexed sequences using cutadapt^73^ and then quality filtered, trimmed, denoised, and merged in R (version 4.4.1) using the dada2^74^ package. Merged sequences were further filtered to only include those within the expected base-pair length of the V4 amplicon and then used to generate an Amplicon Sequence Variant (ASV) table. Taxonomy was assigned using the SILVA v138.2 database^75^. Unassigned ASVs or those belonging to Archaea, Eukaryota, Mitochondria, Chloroplast, or Limnobacter were discarded. The decontam^76^ R package was then used to filter out any ASVs that were possible contaminants based on their prevalence in reagent-only negative control samples. ASVs were further filtered requiring a prevalence of at least 50 reads (∼0.001% median read depth) in 10% of samples, followed by removal of any samples with less than 1,000 total filtered reads. The filtered ASV tables for rectal and ileal biopsies were used for all downstream analyses.

For alpha diversity assessment, samples were first normalized by scaling with ranked subsampling (SRS)^77^ using the SRS.shiny.app for the determination of Cmin. Metrics including Shannon and unique taxa (i.e. Observed) were compared at the ASV level between groups using Wilcox tests with an alpha of 0.05 to determine significance. Beta diversity was tested by PERMANOVA with the adonis function in the vegan R package^78^, using the Euclidean distance of the robust centered log-ratio (CLR) transformed^79^ counts (i.e. robust Aitchison distance) for genus level taxa. Effects included in the PERMANOVA model were evaluated individually using the “margin” argument and included SNAE risk group, country, age, and years living with HIV, using 999 permutations and an alpha of 0.05 after adjustment with the Benjamini-Hochberg (FDR) method^80^. Differential abundance of taxa was determined using the MaAsLin2 R package^81^ at the genus and species level using CLR transformed counts and the default FDR threshold of 0.25 for determining significance.

### Supervised machine learning analysis

Supervised machine learning algorithms including PLS-DA (plasma proteins), sPLS-DA (ileum, rectum, and metabolome), and DIABLO (all omes) were implemented using the mixOmics R package^82^ to characterize the prediction capability of each omic data type and to identify the top discriminatory features for SNAE risk. Models were tuned for the number of components and number of features to keep as appropriate for each model, using 5-fold cross-validation repeated 50 times. Performance of each model was assessed by the area under the receiver operator curve (AUROC) again by 5-fold cross validation repeated 50 times. For the DIABLO model, a design matrix with a moderate correlation of 0.5 was used between each data type.

### Statistics

Statistical analysis was performed using R. Serological biomarkers (e.g., cytokines, metabolites) were log transformed prior to downstream analysis. Multiple linear regression was used to evaluate the effect of SNAE risk group (i.e., low risk versus high risk) and country on biomarker concentrations adjusting for age, and years living with HIV. Partial Spearman correlations between biomarkers and T cell criteria (CD4^+^ T cell count, CD8^+^ T cell count, and CD4/CD8 ratio) were performed using the ppcor R package^83^, adjusting for the effects of country, age, and years living with HIV. Comparisons and correlations were FDR adjusted separately for each ome and considered significant with both a nominal p-value ≤ 0.05 and an adjusted q-value ≤ 0.25. GSEA was performed using HMDB matched identifiers for each available metabolite with the multiGSEA and fgsea R packages^84,85^. Metabolites were supplied as a pre-ranked list of the p-value multiplied by the metabolite’s foldchange from the multiple linear regression models and matched to gene sets from the KEGG database, requiring a minimum gene set size of 3, and an adjusted p-value ≤ 0.25 to determine significance. Differences in the overall composition of the plasma metabolome or proteins were tested by PERMANOVA, similar to the microbiome analysis, using Euclidean distances of the respective log transformed biomarkers. Differences in baseline characteristics were tested using Welch t-tests or chi-squared tests without p-value adjustment.

### Study approval

All participants gave informed consent using IRB-approved forms. The UMN IRB approved the study (STUDY00009216) in the US and the INER Ethics and Research Committees approved the study (C71-18) in Mexico. All participants gave written informed consent using IRB-approved forms.

### Data availability

Sequencing data for the 16S rRNA analysis is available at NCBI under BioProject ID PRJNA1405126. Code used for this analysis available from the corresponding upon reasonable request. Data values for all applicable figures can be found in the Supplemental Data Values file.

### Competing interests

The authors declare no competing interests

### Funding

Funding for this project was provided by the National Institute of Health (grants AI140923, DK143536, DK091538, AI177584, and AI147912) and UMN’s Department of Surgery funds to Dr. Nichole Klatt. CIENI-INER is supported by the Mexican Government (Programa Presupuestal P016; Anexo 13 del Decreto del Presupuesto de Egresos de la Federación).

## Contributions

Conceptualization: NRK, TWS, GS, CMB

Supervision: NRK, TWS, GS, SAR, JA, CMB

Funding acquisition: NRK, TWS, GS

Investigation: CMB, JA, KE, GW, CG, JR, RC, TS, EC, SAR

Formal analysis: CMB

Visualization: CMB

Writing – original draft: CMB

Writing – review & editing: NRK, ES, MB, NF, PH, MG, KE, SAR, TWS

## Acknowledgements

We are deeply grateful to the participants of this study, Dr. Alexander Khoruts who performed the colonoscopies, and Dr. Greg Beilman and Dr. Jeffrey Chipman who performed the lymph node biopsies. We would also like to thank biocrates for generating the metabolomic data, UMN Preclinical Research Center for generating the cytokine data, and the University of Minnesota Genomics Center for generating the 16S rRNA data.

## Abbreviations

SNAE: Serious non-AIDS Event
HIV: Human Immunodeficiency Virus
PWH: People with HIV
ART: Antiretroviral Therapy
US: United States
IRB: Institutional Review Board
IL: Interleukin
TNF: Tumor Necrosis Factor
IFN: Interferon
MIP: Macrophage inflammatory protein
LBP: Lipopolysaccharide Binding Protein
sCD14: Soluble Cluster of Differentiation 14
I-FABP: Intestinal Fatty Acid Binding Protein
LC: Liquid Chromatography
MS: Mass Spectrometry
PCA: Principal component analysis
PE: Phosphatidylethanolamines
PC: Phosphatidylcholines
PG: Phosphatidylglycerols
PS: Phosphatidylserines
SM: Sphingomyelins
AC: Acylcarnitines
TG: Triglyceride
MG: Monoglyceride
LPA: Lysophosphatidic Acid
CDCA: Chenodeoxycholic Acid
CA: Cholic Acid
GSEA: Gene set enrichment analysis
KEGG: Kyoto Encyclopedia of Genes and Genomes
PLS-DA: Partial Least Squares Discriminant Analysis
sPLS-DA: Sparse Partial Least Squares Discriminant Analysis
DIABLO: Data Integration Analysis for Biomarker Discovery using Latent Variable Approaches for Omics Studies
AUROC: Area Under the Receiver Operator Curve
SIV: Simian Immunodeficiency Virus
CoA: Coenzyme A
ASV: Amplicon Sequence Variant
FDR: False Discovery Rate
FIA: Flow Injection Analysis
ESI: Electrospray Ionization
PCR: Polymerase Chain Reaction
CLR: Centered Log-Ratio
SRS: Scaling with Ranked Subscaling

## Supplemental Material

**Supplemental Table 1.**
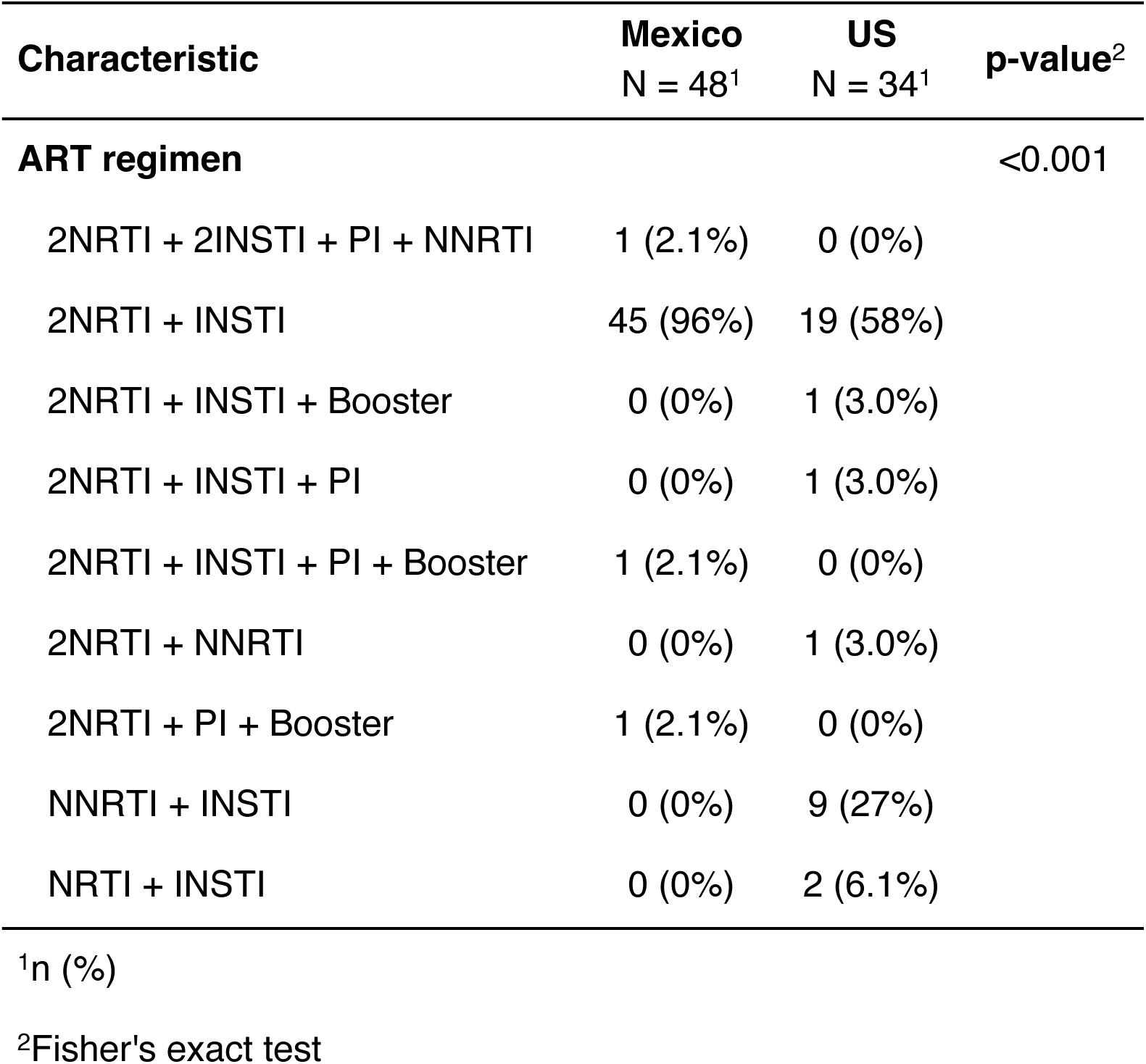
ART regimens by country of enrollment.

**Supplemental Table 2.**
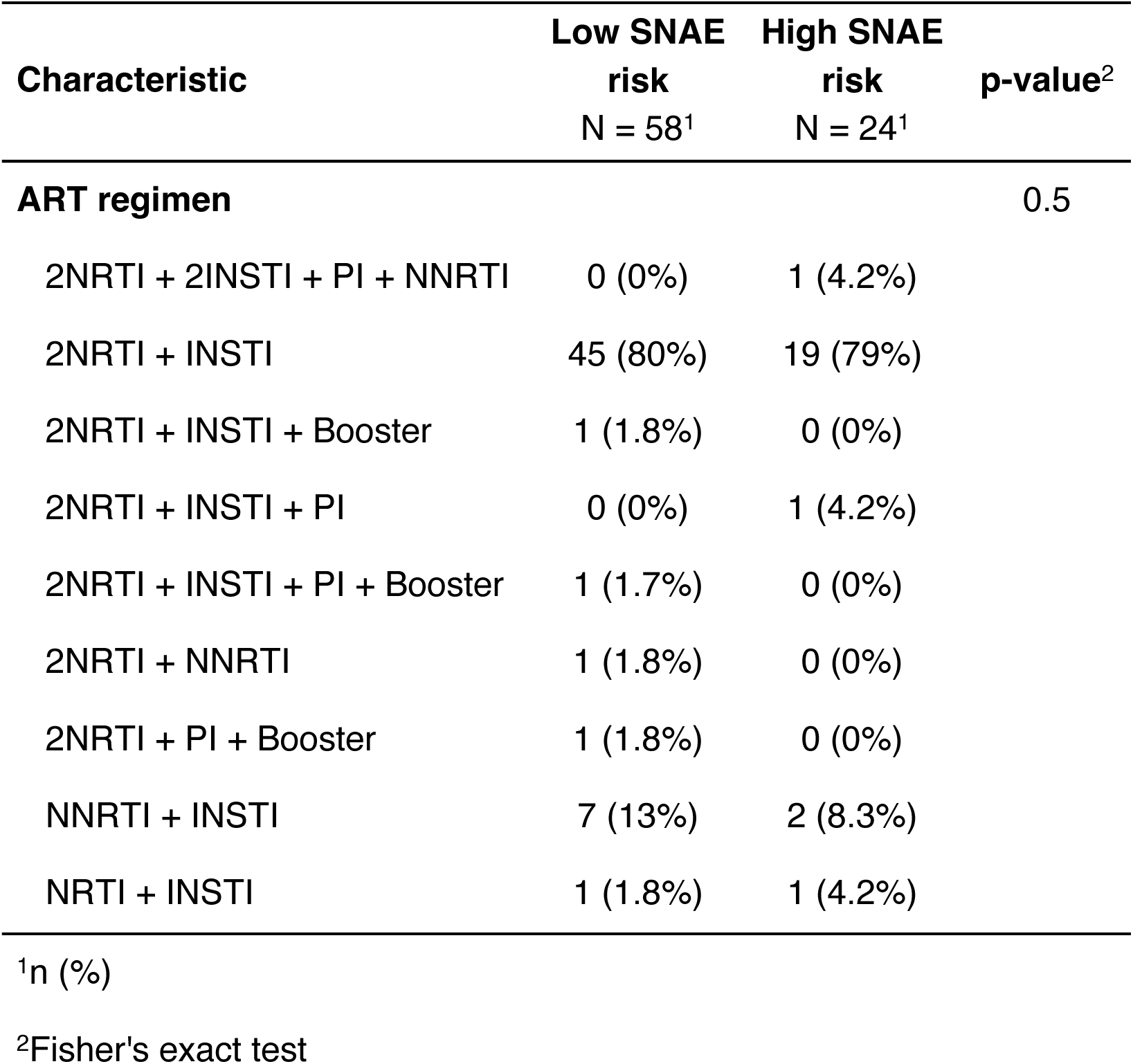
ART regimens by serious non-AIDs event risk group.

**Supplemental Figure 1.**
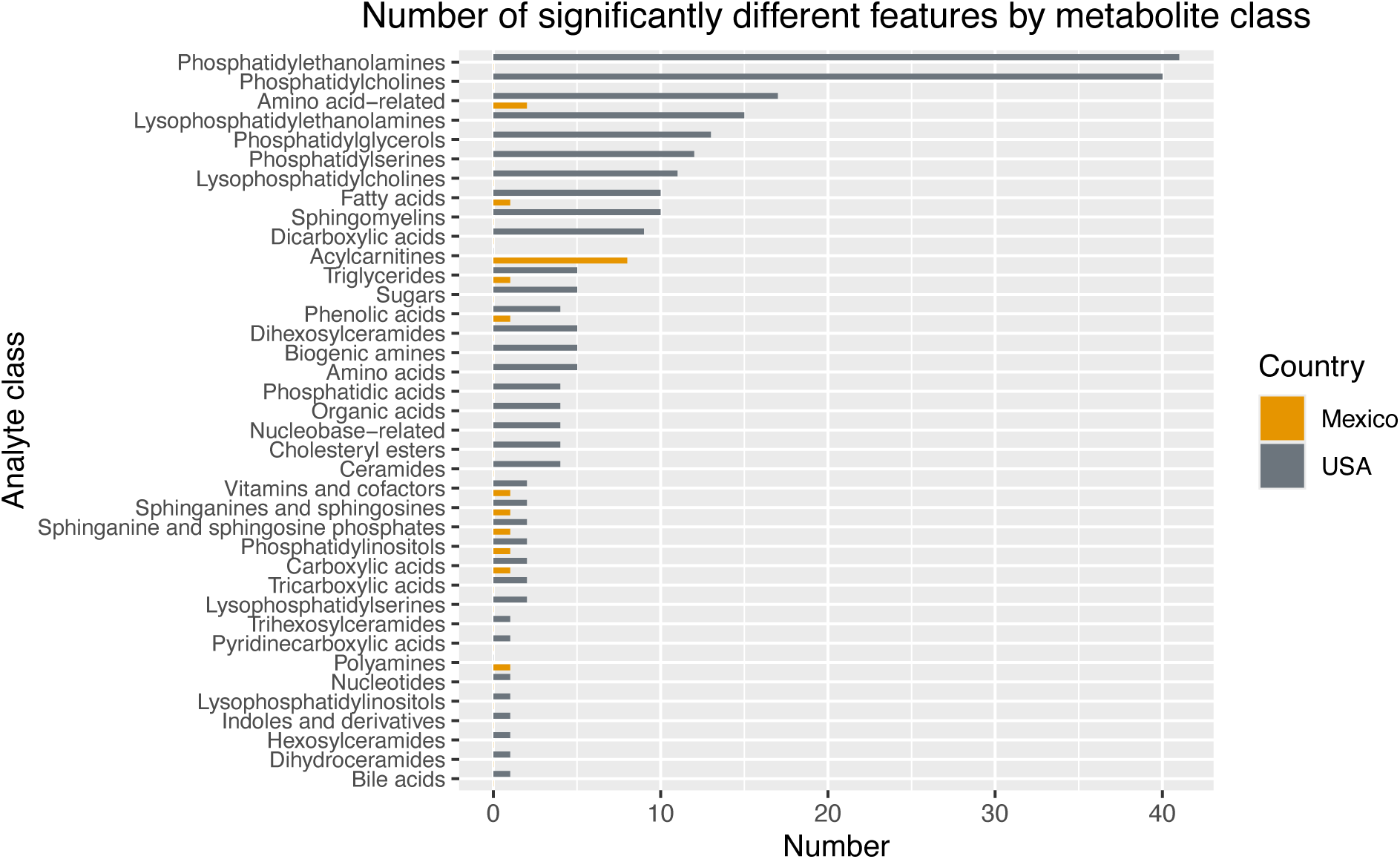
Bar plot representing the number of significantly different metabolites (q-value < 0.25) per each analyte class and whether they were more represented in participants from the US or Mexico (determined by fold change).

**Supplemental Figure 2.**
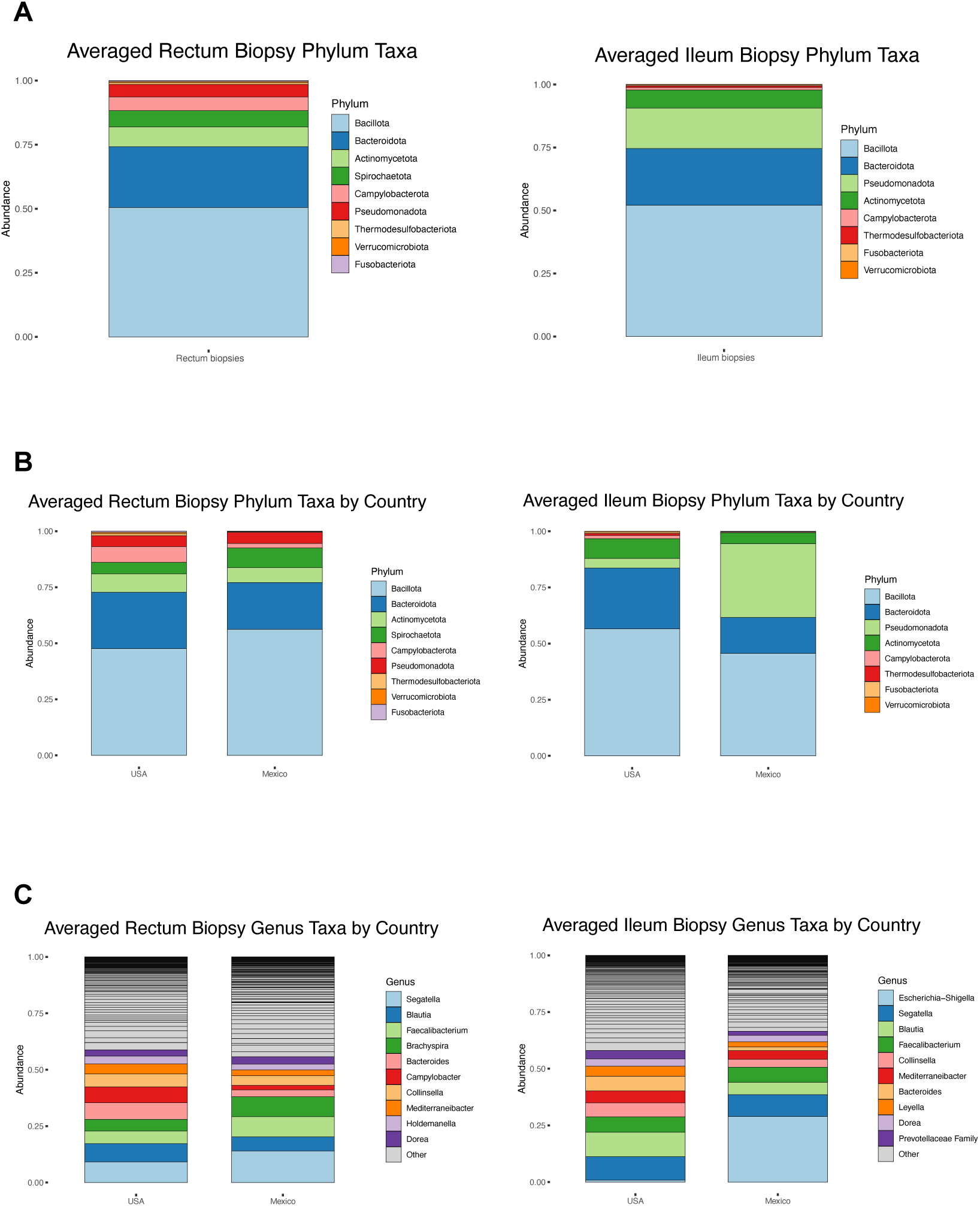
Stacked bar plots showing the average relative abundance of bacterial taxa at the phylum level in rectal and ileal biopsies (A), stratified by country (B), and at the genus level (C).

**Supplemental Figure 3.**
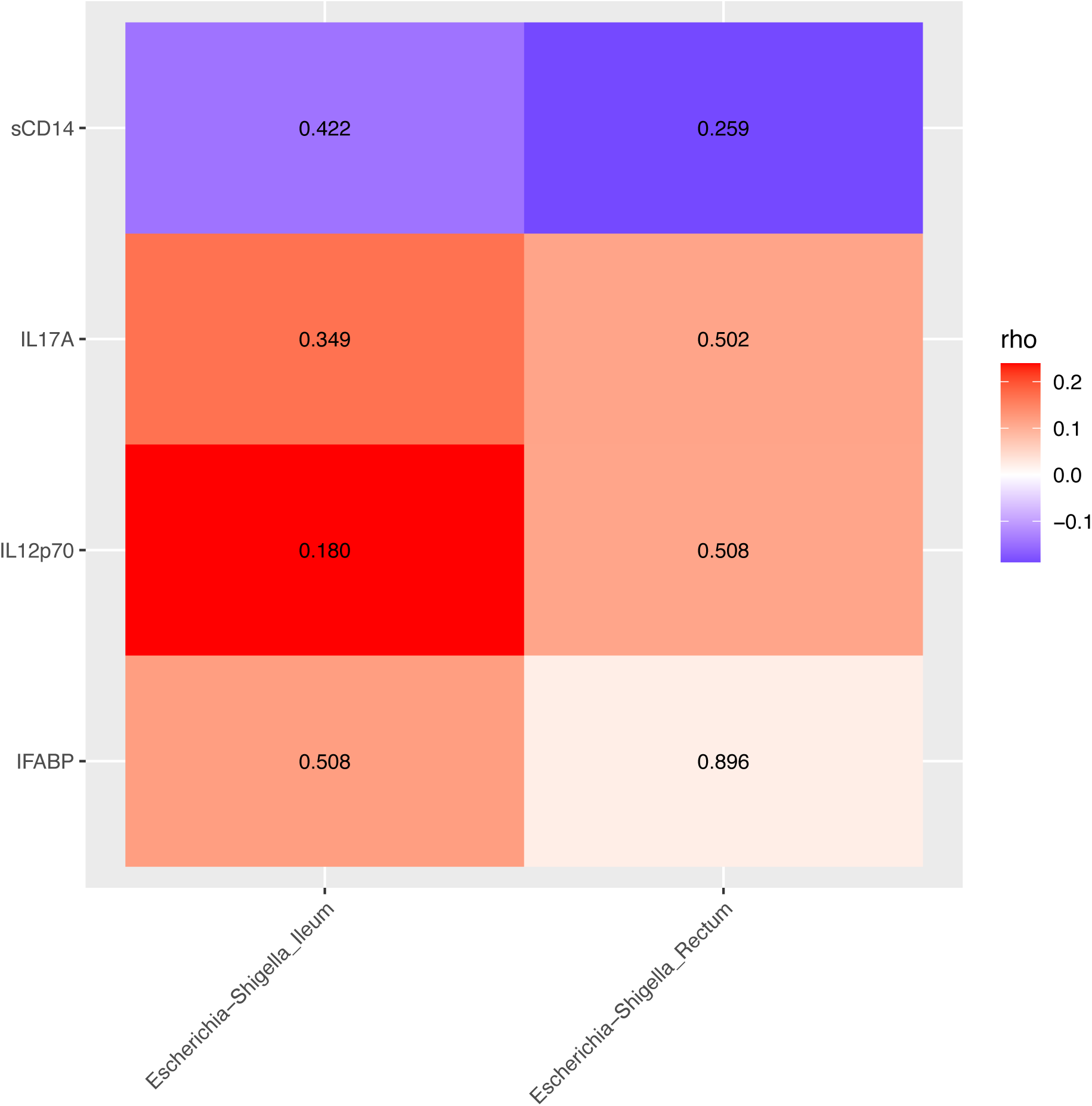
Spearman correlations of *Escherichia-Shigella* abundance (CLR transformed) in the ileum and rectum biopsies of PWH enrolled from Mexico with mucosal cytokines and gut barrier damage markers. Values in each cell are the nominal p-values for each correlation.

**Supplemental Figure 4.**
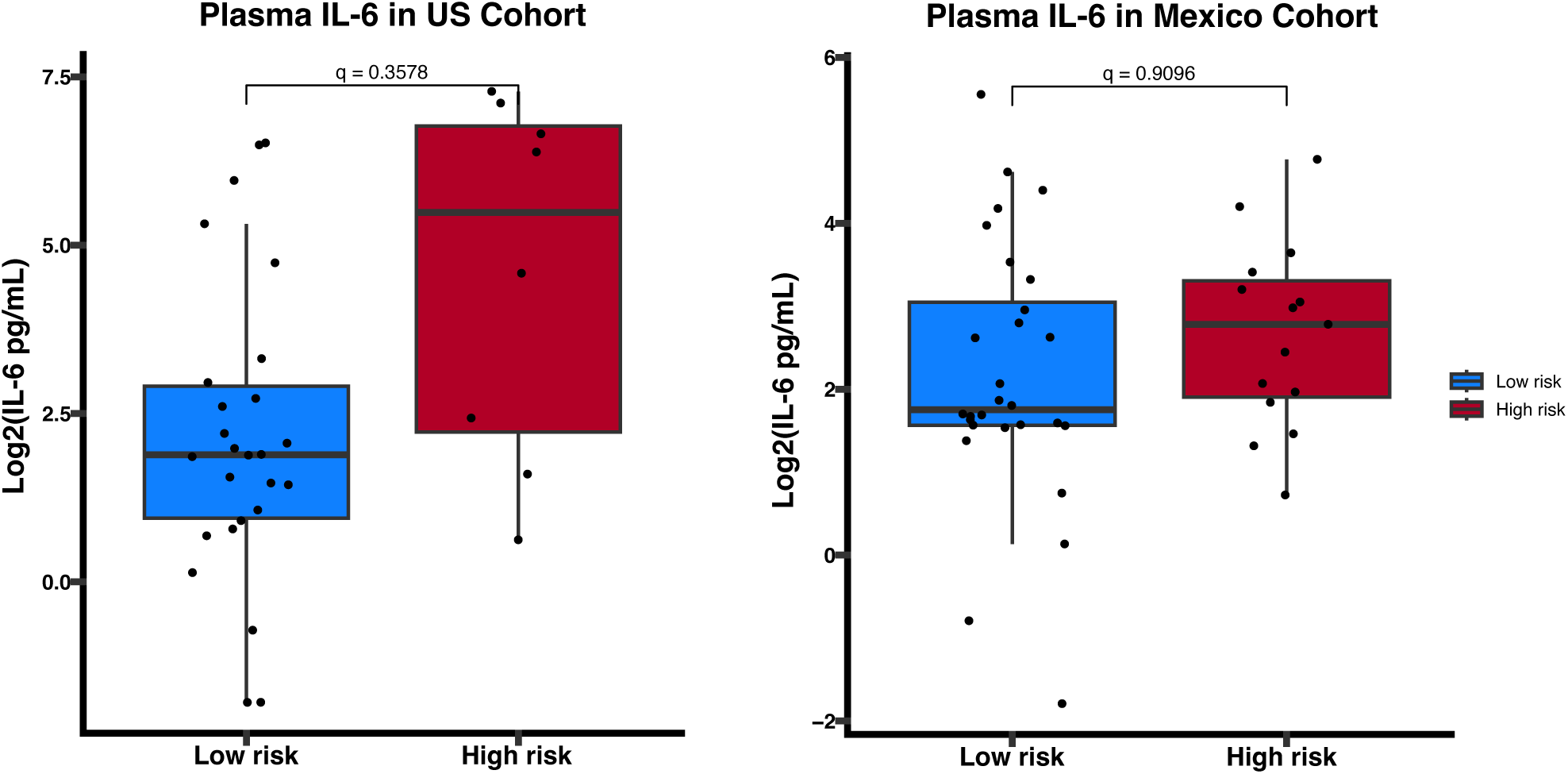
Comparison of log2 transformed plasma IL-6 concentrations between low and high risk SNAE groups when stratified by country, performed using multiple linear regression adjusting for age and years living with HIV. IL-6 concentrations are non-significantly (q > 0.25) higher in the high-risk group within both cohorts of PWH and only becomes significant when combined.

**Supplemental Figure 5.**
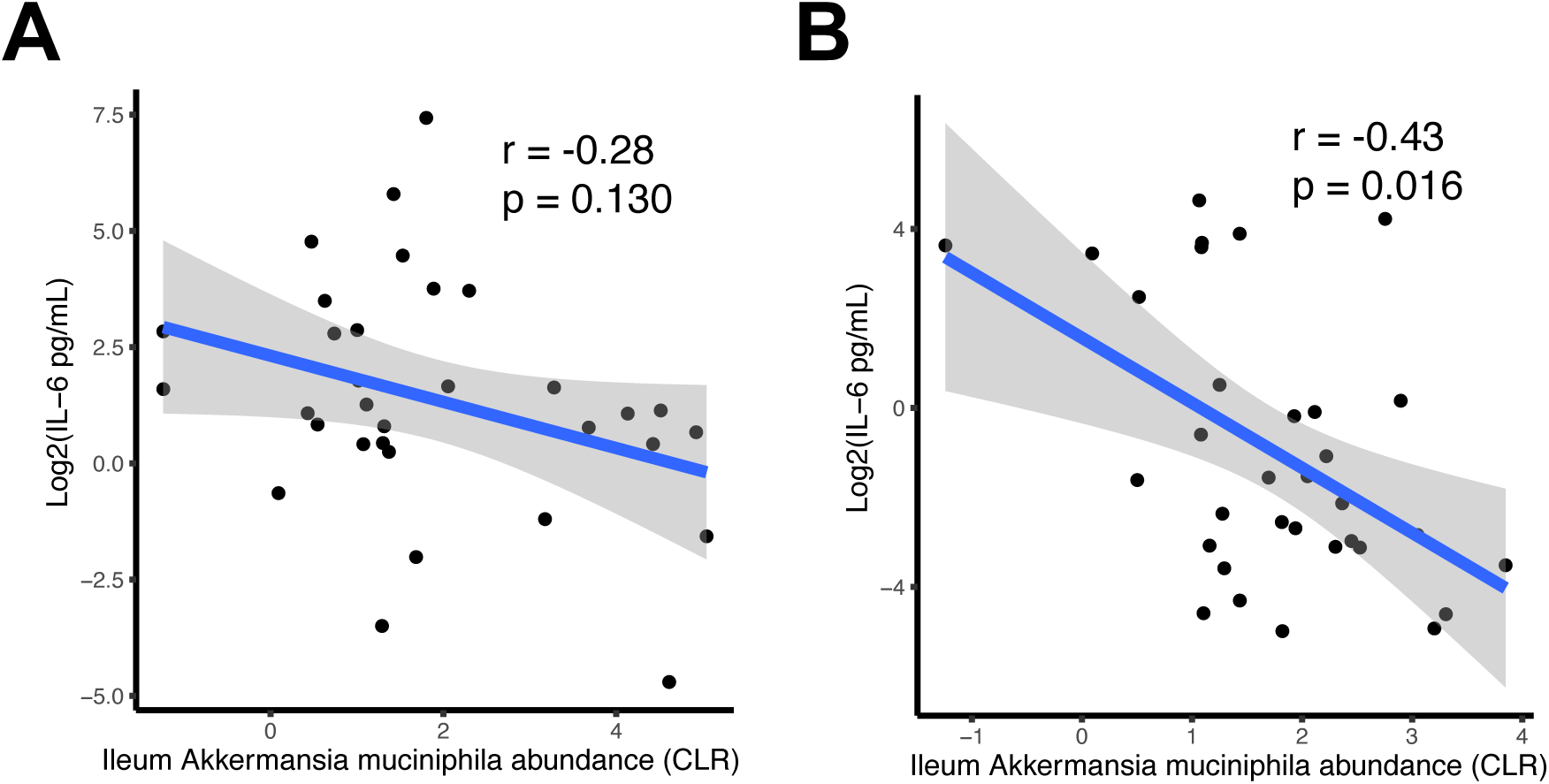
Partial Spearman correlations (adjusted for age and years living with HIV) of log2 transformed plasma IL-6 concentrations with CLR transformed abundance of *Akkermansia muciniphila* in the ileum, stratified by US patients (A) and Mexico patients (B). A negative correlation is seen in both cohorts, though only statistically significant (p < 0.05) in the Mexico cohort.

**Supplemental Figure 6.**
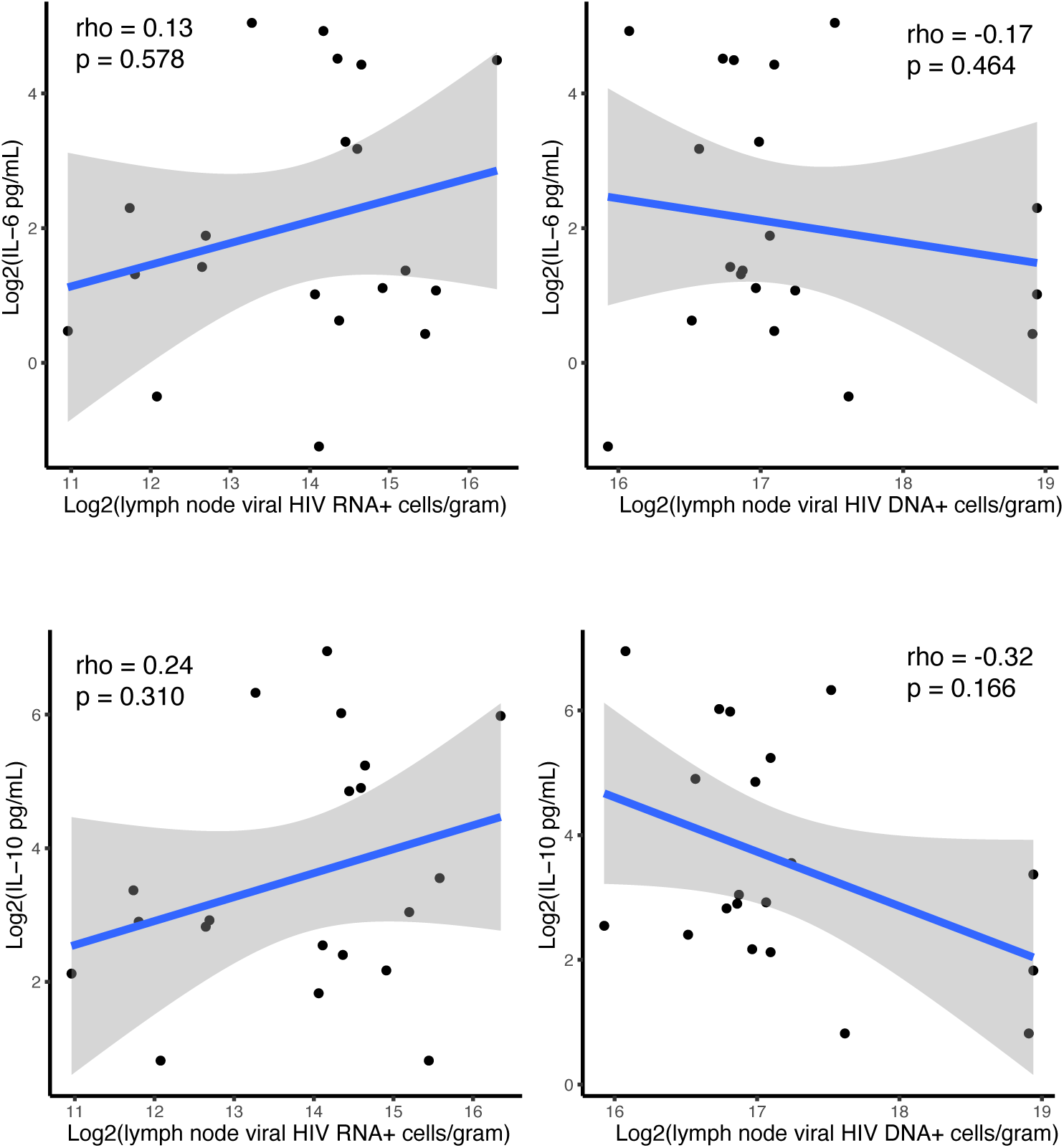
Spearman correlations of log2 transformed plasma IL-6 and IL-10 concentrations with log2 transformed HIV reservoir measures in lymph nodes (RNA^+^ and DNA^+^ cells/gram).

## Notes

### Competing Interest Statement

The authors have declared no competing interest.

