## Supplemental Materials for "Multi-omics links microbial dysbiosis, systemic inflammation and metabolomic disruptions to SNAE risk in treated HIV"

**Supplemental Material:**

***Supplemental Table 1. ART regimens by country of enrollment***

| **Characteristic** | **Mexico** N = 48[1] | **US** N = 34[1] | **p-value**[2] |
|---|---|---|---|
| **ART regimen** | | | <0.001 |
| 2NRTI + 2INSTI + PI + NNRTI | 1 (2.1%) | 0 (0%) | |
| 2NRTI + INSTI | 45 (96%) | 19 (58%) | |
| 2NRTI + INSTI + Booster | 0 (0%) | 1 (3.0%) | |
| 2NRTI + INSTI + PI | 0 (0%) | 1 (3.0%) | |
| 2NRTI + INSTI + PI + Booster | 1 (2.1%) | 0 (0%) | |
| 2NRTI + NNRTI | 0 (0%) | 1 (3.0%) | |
| 2NRTI + PI + Booster | 1 (2.1%) | 0 (0%) | |
| NNRTI + INSTI | 0 (0%) | 9 (27%) | |
| NRTI + INSTI | 0 (0%) | 2 (6.1%) | |

[1]n (%)

[2]Fisher's exact test

*Supplemental Table 2. ART regimens by serious non-AIDs event risk group*

| **Characteristic** | **Low SNAE risk** N = 58[1] | **High SNAE risk** N = 24[1] | **p-value**[2] |
|---|---|---|---|
| **ART regimen** | | | 0.5 |
| 2NRTI + 2INSTI + PI + NNRTI | 0 (0%) | 1 (4.2%) | |
| 2NRTI + INSTI | 45 (80%) | 19 (79%) | |
| 2NRTI + INSTI + Booster | 1 (1.8%) | 0 (0%) | |
| 2NRTI + INSTI + PI | 0 (0%) | 1 (4.2%) | |
| 2NRTI + INSTI + PI + Booster | 1 (1.7%) | 0 (0%) | |
| 2NRTI + NNRTI | 1 (1.8%) | 0 (0%) | |
| 2NRTI + PI + Booster | 1 (1.8%) | 0 (0%) | |
| NNRTI + INSTI | 7 (13%) | 2 (8.3%) | |
| NRTI + INSTI | 1 (1.8%) | 1 (4.2%) | |

[1]n (%)

[2]Fisher's exact test

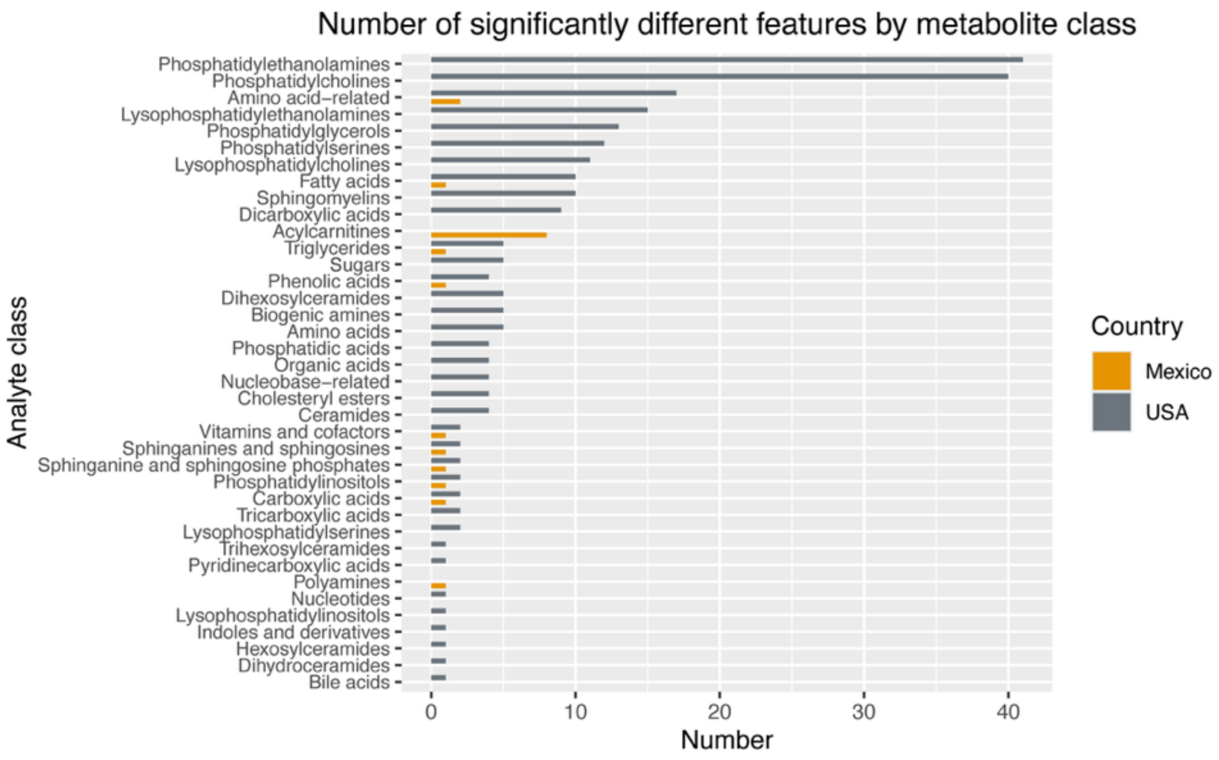


**Supplemental Figure 1** Bar plot representing the number of significantly different metabolites (q-value < 0.25) per each analyte class and whether they were more represented in participants from the US or Mexico (determined by fold change).

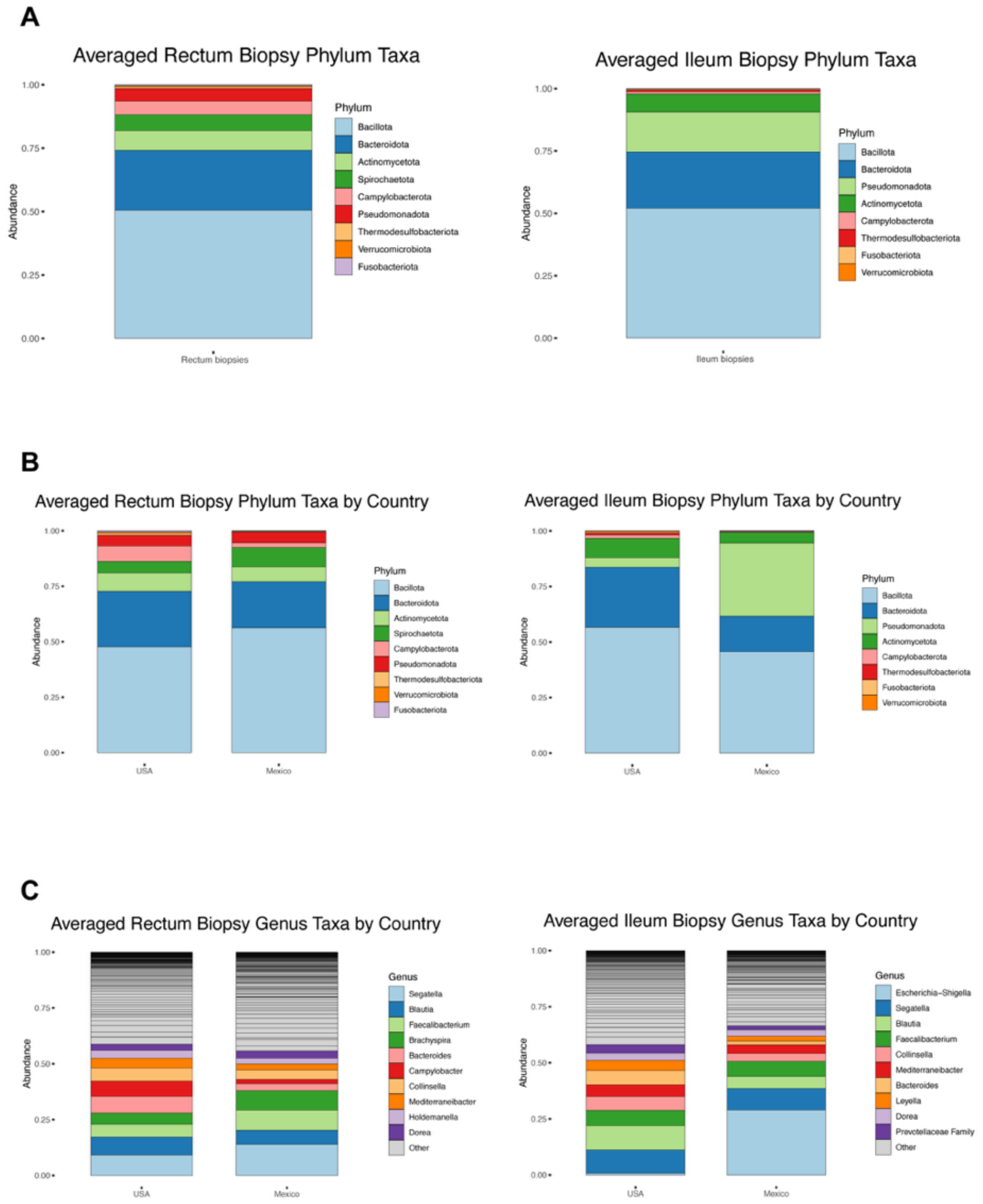


**Supplemental Figure 2** Stacked bar plots showing the average relative abundance of bacterial taxa at the phylum level in rectal and ileal biopsies (A), stratified by country (B), and at the genus level (C).

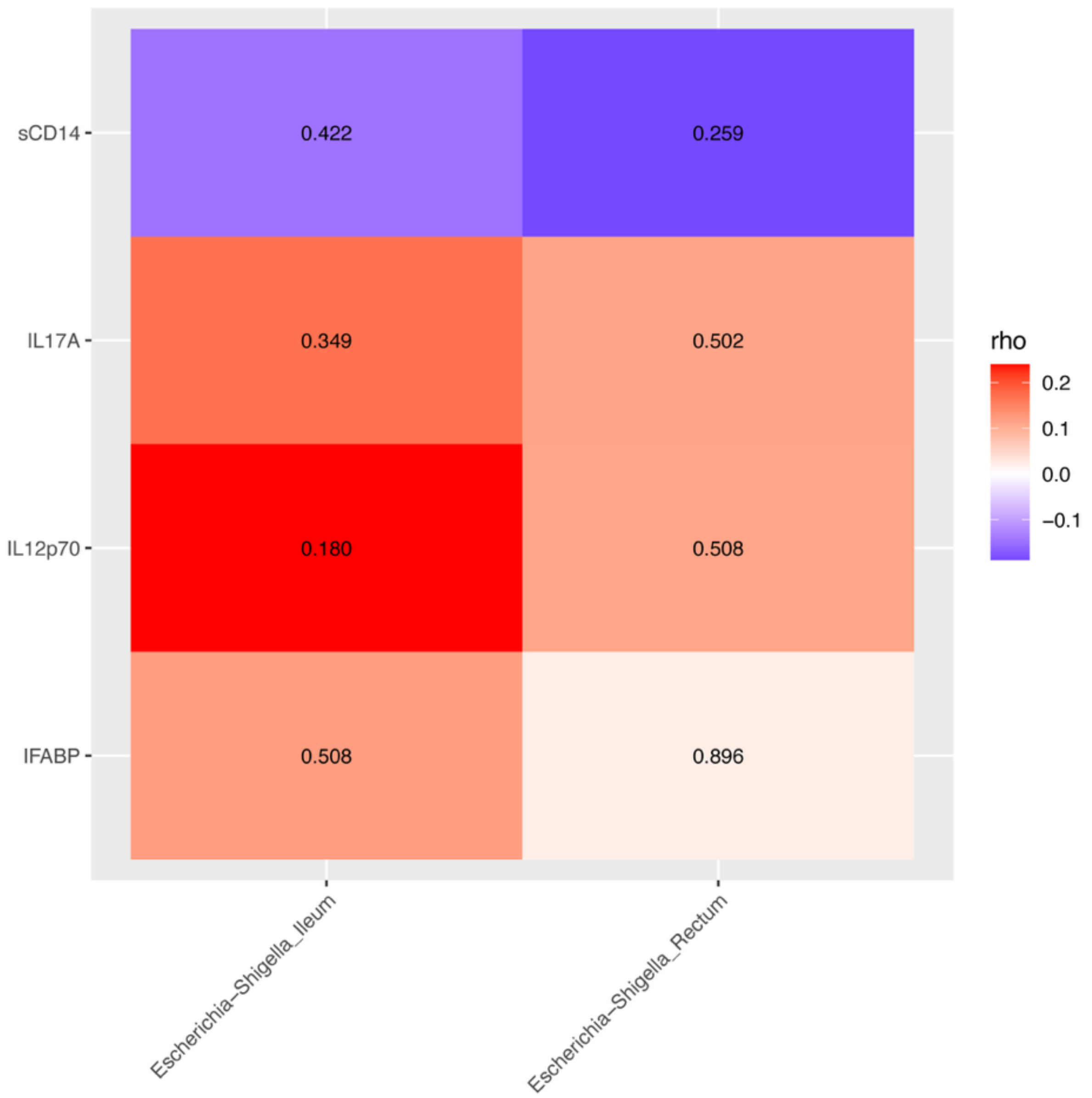


**Supplemental Figure 3** Spearman correlations of *Escherichia-Shigella* abundance (CLR transformed) in the ileum and rectum biopsies of PWH enrolled from Mexico with mucosal cytokines and gut barrier damage markers. Values in each cell are the nominal p-values for each correlation.

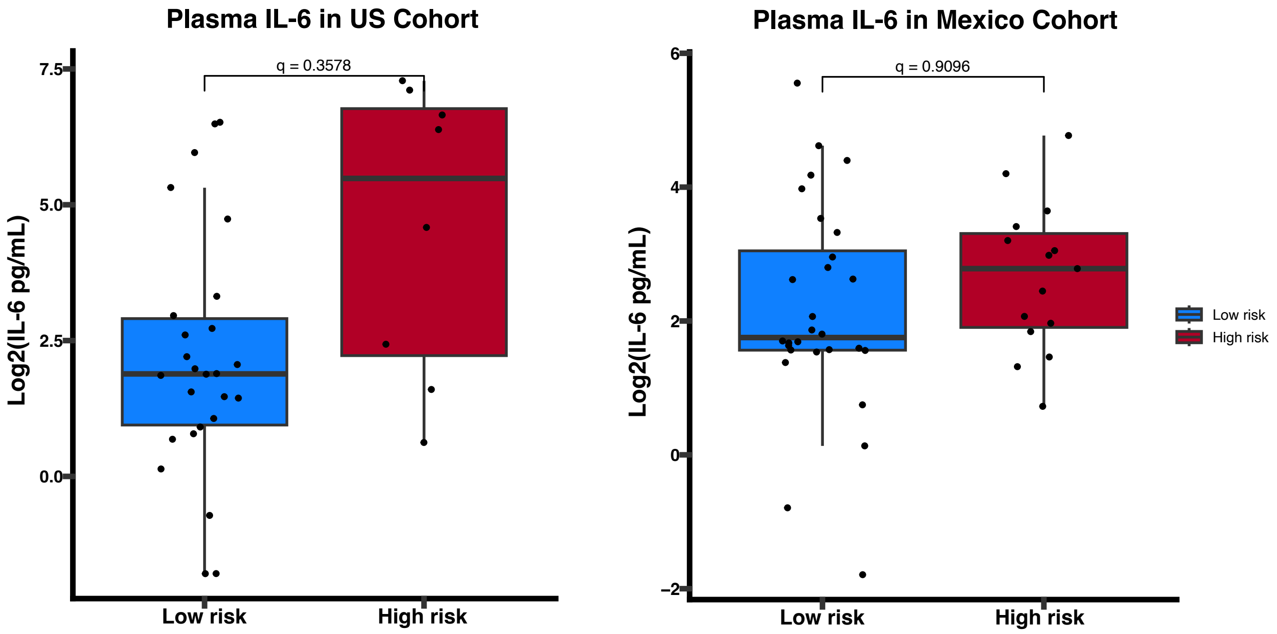


**Supplemental Figure 4** Comparison of log2 transformed plasma IL-6 concentrations between low and high risk SNAE groups when stratified by country, performed using multiple linear regression adjusting for age and years living with HIV. IL-6 concentrations are non-significantly ($q > 0.25$) higher in the high-risk group within both cohorts of PWH and only becomes significant when combined.

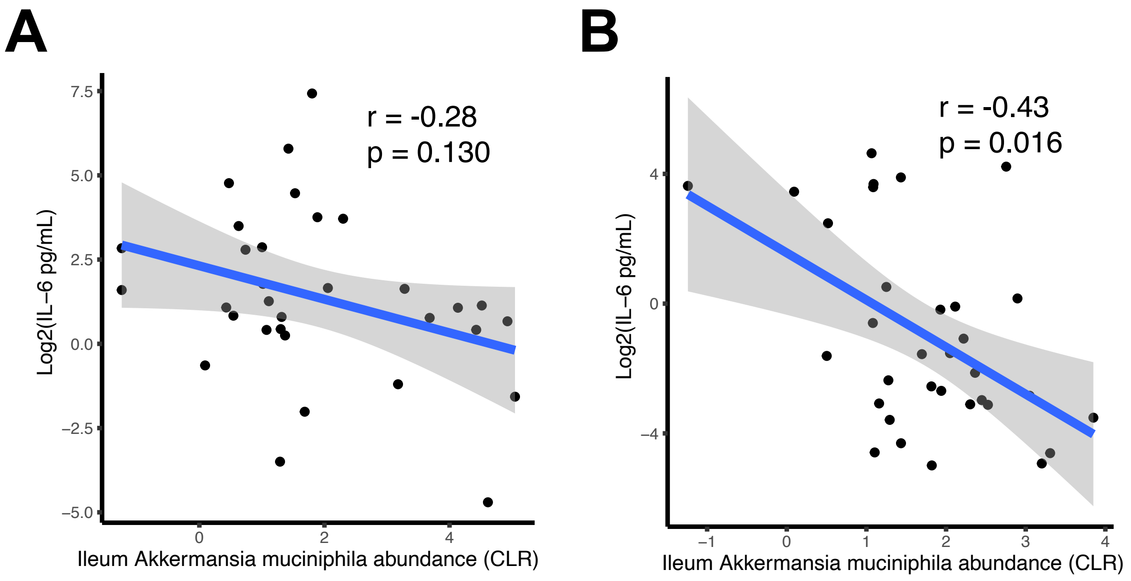


**Supplemental Figure 5** Partial Spearman correlations (adjusted for age and years living with HIV) of log2 transformed plasma IL-6 concentrations with CLR transformed abundance of *Akkermansia muciniphila* in the ileum, stratified by US patients (A) and Mexico patients (B). A negative correlation is seen in both cohorts, though only statistically significant ($p < 0.05$) in the Mexico cohort.

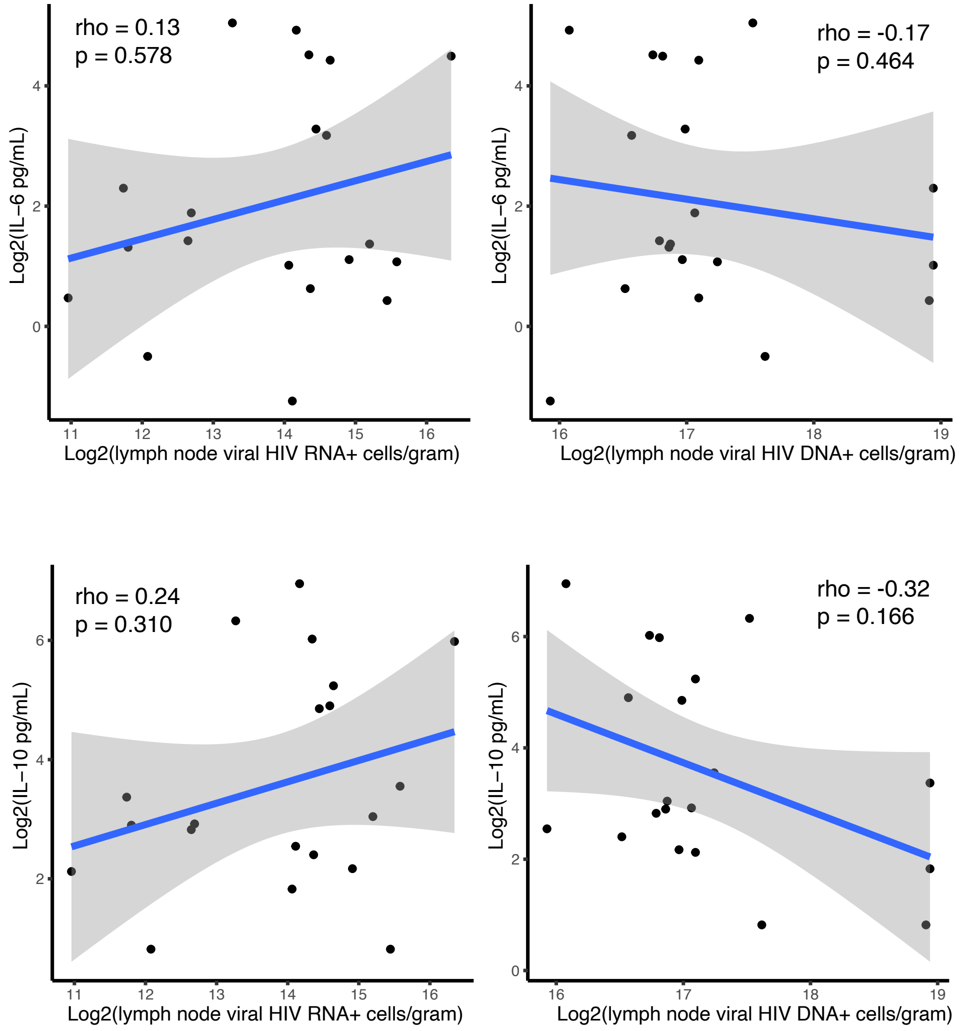


**Supplemental Figure 6** Spearman correlations of log2 transformed plasma IL-6 and IL-10 concentrations with log2 transformed HIV reservoir measures in lymph nodes (RNA$^{+}$ and DNA$^{+}$ cells/gram).

**Supplemental Data 1:** List of country specific effects on metabolomic features

**Supplemental Data 2:** List of metabolites and their shorthand names

**Supplemental Data 3:** List of formulas used for sums and ratios calculated from available metabolites

**Supplemental Data 4:** List of analyte classes and number of molecules per class used in this analysis

**SUPPLEMENTAL DATA 1**

| term | estimate | std.error | statistic | R2 | Variable | q.value |
|---|---|---|---|---|---|---|
| CountryMexico | -2.259 | 0.118 | -19.214 | 0.864 | Taurine | 2.60E-26 |
| CountryMexico | -2.287 | 0.120 | -19.122 | 0.862 | Taurine Synthesis | 2.60E-26 |
| CountryMexico | -2.185 | 0.126 | -17.400 | 0.840 | Nicotinamide (B3) | 4.40E-24 |
| CountryMexico | -3.620 | 0.236 | -15.359 | 0.806 | PS 38:4 | 3.70E-21 |
| CountryMexico | -1.649 | 0.112 | -14.769 | 0.788 | Sum of UFA-PSs | 2.48E-20 |
| CountryMexico | -1.316 | 0.096 | -13.666 | 0.758 | Sum of PUFA-PSs | 1.24E-18 |
| CountryMexico | -2.342 | 0.172 | -13.612 | 0.765 | PE P-18:0/22:4 | 1.31E-18 |
| CountryMexico | -1.761 | 0.143 | -12.307 | 0.726 | PE P-16:0/22:4 | 1.79E-16 |
| CountryMexico | -1.118 | 0.091 | -12.219 | 0.748 | Oxaloacetic acid | 2.26E-16 |
| CountryMexico | -3.039 | 0.266 | -11.415 | 0.709 | PS 40:4 | 5.10E-15 |
| CountryMexico | -3.617 | 0.318 | -11.380 | 0.723 | Sum of MUFA-PSs | 5.36E-15 |
| CountryMexico | -1.600 | 0.143 | -11.165 | 0.721 | LPE P-16:0 | 1.14E-14 |
| CountryMexico | -1.525 | 0.137 | -11.154 | 0.692 | LPE P-18:0 | 1.14E-14 |
| CountryMexico | -1.489 | 0.139 | -10.700 | 0.656 | Spermidine | 6.82E-14 |
| CountryMexico | -1.449 | 0.138 | -10.516 | 0.628 | PE 42:7 | 1.36E-13 |
| CountryMexico | -1.226 | 0.117 | -10.491 | 0.638 | PS 40:7 | 1.42E-13 |
| CountryMexico | -1.467 | 0.140 | -10.460 | 0.628 | NeuAc | 1.52E-13 |
| CountryMexico | -2.646 | 0.257 | -10.296 | 0.681 | PS 36:2 | 2.83E-13 |
| CountryMexico | -1.107 | 0.108 | -10.226 | 0.660 | 2-OH-Glutaric acid | 3.59E-13 |
| CountryMexico | 1.220 | 0.120 | 10.148 | 0.620 | Sarcosine | 4.73E-13 |
| CountryMexico | 1.403 | 0.139 | 10.093 | 0.653 | Spermidine Acetylation | 5.66E-13 |
| CountryMexico | -1.614 | 0.160 | -10.065 | 0.613 | PE 40:4 | 6.07E-13 |
| CountryMexico | -0.693 | 0.070 | -9.943 | 0.608 | Threonic acid | 9.71E-13 |
| CountryMexico | -1.031 | 0.105 | -9.815 | 0.635 | LPE 18:0 | 1.59E-12 |
| CountryMexico | 1.469 | 0.150 | 9.797 | 0.579 | Sarcosine Synthesis from Choline | 1.65E-12 |
| CountryMexico | -1.242 | 0.128 | -9.731 | 0.646 | 2-OH-Glutarate Synthesis | 2.09E-12 |
| CountryMexico | -4.205 | 0.433 | -9.705 | 0.663 | PS 36:1 | 2.25E-12 |
| CountryMexico | -3.460 | 0.364 | -9.497 | 0.627 | Serotonin | 5.21E-12 |
| CountryMexico | 3.479 | 0.370 | 9.410 | 0.640 | Kynurenine to Serotonin Ratio | 7.29E-12 |
| CountryMexico | -3.627 | 0.389 | -9.322 | 0.620 | Serotonin Synthesis | 1.02E-11 |
| CountryMexico | -1.165 | 0.127 | -9.177 | 0.592 | Asp | 1.77E-11 |
| CountryMexico | -1.009 | 0.110 | -9.179 | 0.589 | PE 40:1 | 1.77E-11 |
| CountryMexico | 1.384 | 0.151 | 9.157 | 0.566 | Sarcosine Synthesis from Gly | 1.87E-11 |
| CountryMexico | -2.527 | 0.281 | -9.001 | 0.584 | PS 40:5 | 3.52E-11 |
| CountryMexico | -1.826 | 0.205 | -8.923 | 0.547 | PE P-18:0/22:3 | 4.77E-11 |
| CountryMexico | -0.884 | 0.100 | -8.831 | 0.608 | SDH Activity | 6.85E-11 |
| CountryMexico | -0.832 | 0.097 | -8.615 | 0.515 | AHCys | 1.63E-10 |
| CountryMexico | 3.235 | 0.375 | 8.618 | 0.605 | Serotonin Catabolism | 1.63E-10 |
| CountryMexico | -1.736 | 0.206 | -8.420 | 0.584 | LPE P-18:1 | 3.65E-10 |

| | | | | | | |
|---|---|---|---|---|---|---|
| CountryMexico | -1.228 | 0.148 | -8.276 | 0.541 | PE P-18:0/22:5 | 6.58E-10 |
| CountryMexico | -1.167 | 0.143 | -8.172 | 0.496 | PE 38:4 | 1.00E-09 |
| CountryMexico | -1.148 | 0.141 | -8.120 | 0.532 | PE P-20:0/20:4 | 1.22E-09 |
| CountryMexico | -2.066 | 0.255 | -8.092 | 0.525 | LPS 18:0 | 1.34E-09 |
| CountryMexico | 2.301 | 0.286 | 8.052 | 0.596 | Ratio of PUFA-PSs to MUFA-PSs | 1.56E-09 |
| CountryMexico | 0.808 | 0.101 | 8.004 | 0.600 | MDH1 Activity | 1.83E-09 |
| CountryMexico | -0.808 | 0.101 | -8.004 | 0.600 | MDH2 Activity | 1.83E-09 |
| CountryMexico | -0.979 | 0.123 | -7.981 | 0.479 | PG 16:1_20:4 | 1.98E-09 |
| CountryMexico | -0.314 | 0.040 | -7.951 | 0.560 | Fumarase Activity | 2.20E-09 |
| CountryMexico | -1.356 | 0.174 | -7.785 | 0.536 | PS 40:6 | 4.37E-09 |
| CountryMexico | 0.962 | 0.126 | 7.621 | 0.578 | CS Activity | 8.62E-09 |
| CountryMexico | 0.894 | 0.118 | 7.587 | 0.552 | Pyruvate Carboxylase Deficiency | 9.78E-09 |
| CountryMexico | 1.067 | 0.145 | 7.352 | 0.508 | Asn Synthesis | 2.61E-08 |
| CountryMexico | -0.936 | 0.128 | -7.305 | 0.493 | Hex2Cer d18:1/22:0 | 3.13E-08 |
| CountryMexico | 1.041 | 0.143 | 7.285 | 0.535 | ACY2 Deficiency | 3.33E-08 |
| CountryMexico | -0.662 | 0.091 | -7.270 | 0.464 | Aconitase Activity | 3.48E-08 |
| CountryMexico | -1.516 | 0.210 | -7.219 | 0.449 | Adenosine | 4.24E-08 |
| CountryMexico | -0.879 | 0.123 | -7.121 | 0.463 | Suc | 6.21E-08 |
| CountryMexico | -0.696 | 0.098 | -7.123 | 0.471 | PS 38:5 | 6.21E-08 |
| CountryMexico | -0.736 | 0.104 | -7.095 | 0.457 | PE 38:2 | 6.82E-08 |
| CountryMexico | -1.014 | 0.143 | -7.091 | 0.479 | Suc to a-Ketoglutarate Ratio | 6.82E-08 |
| CountryMexico | -0.818 | 0.115 | -7.084 | 0.440 | Isocitric acid | 6.91E-08 |
| CountryMexico | -2.456 | 0.351 | -6.999 | 0.428 | Benzoic Acid Conjugation | 9.72E-08 |
| CountryMexico | 0.717 | 0.103 | 6.966 | 0.444 | Glycolic Acid to Oxalic Acid Ratio | 1.10E-07 |
| CountryMexico | -0.932 | 0.135 | -6.910 | 0.412 | PE 42:8 | 1.37E-07 |
| CountryMexico | -0.942 | 0.137 | -6.895 | 0.501 | Spermidine Synthesis | 1.44E-07 |
| CountryMexico | -1.611 | 0.235 | -6.841 | 0.458 | Histamine | 1.78E-07 |
| CountryMexico | -0.833 | 0.123 | -6.784 | 0.404 | AHCys to Leu Ratio | 2.23E-07 |
| CountryMexico | -1.537 | 0.232 | -6.632 | 0.437 | Histamine Synthesis | 4.15E-07 |
| CountryMexico | -1.163 | 0.177 | -6.583 | 0.437 | PA 18:2_18:3 | 5.02E-07 |
| CountryMexico | -0.840 | 0.133 | -6.304 | 0.417 | PE P-16:0/22:5 | 1.58E-06 |
| CountryMexico | -0.979 | 0.158 | -6.189 | 0.370 | Hex2Cer d18:1/24:0 | 2.50E-06 |
| CountryMexico | -0.538 | 0.087 | -6.172 | 0.390 | PE 44:11 | 2.65E-06 |
| CountryMexico | -1.601 | 0.263 | -6.095 | 0.356 | 4-OH-Phenylacetic acid | 3.58E-06 |
| CountryMexico | -1.376 | 0.226 | -6.081 | 0.419 | PS 34:1 | 3.73E-06 |
| CountryMexico | -0.716 | 0.118 | -6.072 | 0.394 | LPC 17:0 | 3.82E-06 |
| CountryMexico | -1.927 | 0.319 | -6.048 | 0.384 | Carnosine | 4.16E-06 |
| CountryMexico | -0.548 | 0.091 | -6.016 | 0.346 | Aconitate to Citrate Ratio | 4.69E-06 |
| CountryMexico | -0.861 | 0.143 | -6.013 | 0.390 | LPC 20:4 | 4.69E-06 |
| CountryMexico | -0.636 | 0.106 | -6.002 | 0.366 | PC O-40:1 | 4.85E-06 |
| CountryMexico | -0.648 | 0.109 | -5.941 | 0.362 | LPE 17:0 | 6.13E-06 |

| | | | | | | |
|---|---|---|---|---|---|---|
| CountryMexico | -1.963 | 0.335 | -5.867 | 0.353 | HipAcid | 8.17E-06 |
| CountryMexico | -0.726 | 0.124 | -5.850 | 0.364 | Homovanillic acid | 8.54E-06 |
| CountryMexico | -0.774 | 0.132 | -5.851 | 0.329 | PE 40:7 | 8.54E-06 |
| CountryMexico | -0.875 | 0.150 | -5.841 | 0.433 | Glu to a-Ketoglutarate Ratio | 8.75E-06 |
| CountryMexico | -1.853 | 0.318 | -5.825 | 0.370 | Carnosine Synthesis | 9.24E-06 |
| CountryMexico | -0.621 | 0.107 | -5.809 | 0.335 | PC O-34:0 | 9.76E-06 |
| CountryMexico | -0.733 | 0.126 | -5.803 | 0.325 | PE 40:8 | 9.86E-06 |
| CountryMexico | -1.382 | 0.240 | -5.755 | 0.362 | 4-OH-HipAcid | 1.18E-05 |
| CountryMexico | -0.437 | 0.076 | -5.741 | 0.347 | PC O-40:4 | 1.24E-05 |
| CountryMexico | 1.675 | 0.300 | 5.592 | 0.365 | b-Ala Synthesis | 2.23E-05 |
| CountryMexico | -0.705 | 0.127 | -5.547 | 0.321 | AconAcid | 2.64E-05 |
| CountryMexico | -0.892 | 0.162 | -5.508 | 0.303 | PE 40:5 | 3.03E-05 |
| CountryMexico | -0.723 | 0.131 | -5.507 | 0.302 | Sum of PUFA-PEs | 3.03E-05 |
| CountryMexico | -2.403 | 0.439 | -5.475 | 0.331 | HPHPA | 3.40E-05 |
| CountryMexico | -0.591 | 0.108 | -5.454 | 0.331 | LPE 20:4 | 3.65E-05 |
| CountryMexico | -0.612 | 0.113 | -5.426 | 0.381 | LPC 18:0 | 4.04E-05 |
| CountryMexico | -0.367 | 0.068 | -5.414 | 0.316 | PC 32:0 | 4.19E-05 |
| CountryMexico | -0.875 | 0.162 | -5.402 | 0.351 | PE P-18:0/20:4 | 4.36E-05 |
| CountryMexico | -0.501 | 0.093 | -5.385 | 0.365 | Sum of SFA-LPCs | 4.61E-05 |
| CountryMexico | -0.448 | 0.084 | -5.337 | 0.386 | Cit | 5.52E-05 |
| CountryMexico | -0.574 | 0.108 | -5.324 | 0.353 | LPE 16:0 | 5.75E-05 |
| CountryMexico | -0.501 | 0.094 | -5.310 | 0.366 | Sum of LPCs | 6.00E-05 |
| CountryMexico | -0.503 | 0.095 | -5.307 | 0.366 | Sum of LCFA-LPCs | 6.02E-05 |
| CountryMexico | -0.843 | 0.159 | -5.297 | 0.332 | PA 20:0_20:4 | 6.20E-05 |
| CountryMexico | -0.453 | 0.086 | -5.255 | 0.308 | PC O-36:0 | 7.24E-05 |
| CountryMexico | -1.798 | 0.346 | -5.190 | 0.306 | HipAcid Synthesis | 9.23E-05 |
| CountryMexico | -0.745 | 0.145 | -5.148 | 0.282 | AHCY Deficiency | 0.000107659 |
| CountryMexico | -0.478 | 0.093 | -5.125 | 0.341 | LPC 16:0 | 0.000116767 |
| CountryMexico | -0.613 | 0.120 | -5.118 | 0.324 | PE 38:1 | 0.00011877 |
| CountryMexico | -0.665 | 0.130 | -5.107 | 0.272 | 3-HMGA | 0.00012266 |
| CountryMexico | -1.047 | 0.206 | -5.093 | 0.285 | cAMP | 0.000128769 |
| CountryMexico | -0.418 | 0.082 | -5.090 | 0.308 | PC O-42:1 | 0.000128779 |
| CountryMexico | -0.740 | 0.146 | -5.064 | 0.342 | Glu | 0.000141123 |
| CountryMexico | -0.458 | 0.092 | -4.992 | 0.330 | CPS Deficiency (NBS) | 0.000184839 |
| CountryMexico | -0.511 | 0.103 | -4.948 | 0.339 | 5-Oxo-Pro | 0.000216076 |
| CountryMexico | -0.233 | 0.048 | -4.867 | 0.259 | Glyceric acid | 0.000292171 |
| CountryMexico | -0.423 | 0.087 | -4.845 | 0.252 | PC O-38:4 | 0.000314408 |
| CountryMexico | -0.434 | 0.090 | -4.821 | 0.293 | PC 28:1 | 0.000338109 |
| CountryMexico | -0.542 | 0.114 | -4.754 | 0.363 | Sum of PUFA-LPCs | 0.000431943 |
| CountryMexico | -0.871 | 0.183 | -4.747 | 0.334 | PS 36:3 | 0.000439907 |
| CountryMexico | 1.157 | 0.244 | 4.738 | 0.344 | Ethylmalonic Aciduria | 0.000451918 |

| | | | | | | |
|---|---|---|---|---|---|---|
| CountryMexico | -0.608 | 0.129 | -4.709 | 0.283 | LPE 22:4 | 0.00049858 |
| CountryMexico | -0.848 | 0.180 | -4.701 | 0.255 | Orotic acid | 0.000509054 |
| CountryMexico | -1.481 | 0.316 | -4.681 | 0.244 | PE P-18:0/16:1 | 0.000545451 |
| CountryMexico | 0.447 | 0.096 | 4.646 | 0.253 | Biotin (B7) | 0.000616769 |
| CountryMexico | -0.513 | 0.112 | -4.585 | 0.261 | PE 20:0 | 0.000766375 |
| CountryMexico | -0.496 | 0.109 | -4.549 | 0.240 | PS 38:7 | 0.000867852 |
| CountryMexico | -0.497 | 0.109 | -4.547 | 0.324 | Sum of UFA-LPCs | 0.000867852 |
| CountryMexico | -0.389 | 0.086 | -4.514 | 0.223 | PC O-42:5 | 0.000971512 |
| CountryMexico | 0.463 | 0.103 | 4.478 | 0.270 | Glycolic acid | 0.001101189 |
| CountryMexico | -0.704 | 0.157 | -4.472 | 0.302 | PE P-16:0/20:4 | 0.001114874 |
| CountryMexico | -0.342 | 0.077 | -4.467 | 0.259 | 3-Deoxyglucosone | 0.001127902 |
| CountryMexico | -0.354 | 0.080 | -4.398 | 0.220 | PC O-40:5 | 0.001439422 |
| CountryMexico | -0.265 | 0.060 | -4.394 | 0.251 | Ratio of PI 18:0_20:4 to PIs | 0.001446244 |
| CountryMexico | -0.751 | 0.172 | -4.376 | 0.228 | N-Ac-Arg | 0.001533001 |
| CountryMexico | -0.675 | 0.155 | -4.368 | 0.225 | 2-OH-2-Met-butyric acid | 0.00156578 |
| CountryMexico | -0.291 | 0.067 | -4.341 | 0.242 | Sum of MUFA-PCs O | 0.001714008 |
| CountryMexico | -0.460 | 0.106 | -4.336 | 0.223 | 2-OH-Phenylacetic acid | 0.001734352 |
| CountryMexico | -0.547 | 0.127 | -4.316 | 0.233 | Putrescine | 0.00184871 |
| CountryMexico | -0.550 | 0.128 | -4.304 | 0.212 | PE 38:7 | 0.00191397 |
| CountryMexico | -0.584 | 0.136 | -4.301 | 0.223 | 3-OH-Glutaric acid | 0.001919427 |
| CountryMexico | -0.276 | 0.065 | -4.263 | 0.258 | N-Ac-Ser | 0.002183872 |
| CountryMexico | -0.608 | 0.143 | -4.250 | 0.205 | Hex2Cer d18:1/14:0 | 0.00227444 |
| CountryMexico | -0.601 | 0.142 | -4.228 | 0.209 | PE 38:5 | 0.002443686 |
| CountryMexico | -0.457 | 0.108 | -4.224 | 0.204 | PS 40:8 | 0.002463375 |
| CountryMexico | -0.586 | 0.140 | -4.197 | 0.203 | PE 36:4 | 0.002692178 |
| CountryMexico | 1.215 | 0.290 | 4.182 | 0.316 | Ratio of CDCA to CA | 0.002815856 |
| CountryMexico | 0.642 | 0.156 | 4.121 | 0.221 | MCAD Deficiency (NBS) | 0.003466826 |
| CountryMexico | -0.450 | 0.109 | -4.114 | 0.208 | PC O-44:6 | 0.003519807 |
| CountryMexico | -0.348 | 0.085 | -4.073 | 0.211 | PC O-32:1 | 0.004039965 |
| CountryMexico | -0.655 | 0.162 | -4.036 | 0.188 | PE 32:0 | 0.004503952 |
| CountryMexico | -0.324 | 0.080 | -4.039 | 0.250 | PC O-36:1 | 0.004503952 |
| CountryMexico | 0.479 | 0.119 | 4.037 | 0.265 | Betaine Synthesis | 0.004503952 |
| CountryMexico | 0.236 | 0.059 | 4.018 | 0.255 | SPBP d17:0 | 0.004734638 |
| CountryMexico | 0.575 | 0.143 | 4.019 | 0.250 | Phenylacetate to PAGln Ratio | 0.004734638 |
| CountryMexico | -0.623 | 0.156 | -4.004 | 0.268 | Glutaminase Activity | 0.004930193 |
| CountryMexico | -0.762 | 0.190 | -4.001 | 0.237 | PE P-20:0/20:5 | 0.004959638 |
| CountryMexico | -0.424 | 0.107 | -3.966 | 0.217 | PC O-30:0 | 0.005555707 |
| CountryMexico | -0.875 | 0.221 | -3.963 | 0.238 | Fructose | 0.005581785 |
| CountryMexico | -0.498 | 0.126 | -3.958 | 0.206 | PG 20:4_22:3 | 0.005631842 |
| CountryMexico | 0.504 | 0.127 | 3.953 | 0.273 | Ratio of Pro to Cit | 0.00569505 |
| CountryMexico | -1.316 | 0.334 | -3.940 | 0.208 | SPB d18:0 | 0.005921107 |

| | | | | | | |
|---|---|---|---|---|---|---|
| CountryMexico | -0.881 | 0.224 | -3.937 | 0.239 | Fructose to Glucose Ratio | 0.00594052 |
| CountryMexico | -0.277 | 0.071 | -3.929 | 0.200 | FA 20:0 | 0.006069419 |
| CountryMexico | -2.176 | 0.558 | -3.897 | 0.259 | SPB d18:1 | 0.006700937 |
| CountryMexico | -0.588 | 0.153 | -3.858 | 0.242 | Homovanillate to Vanillylmandelate Ratio | 0.007597339 |
| CountryMexico | 0.408 | 0.106 | 3.841 | 0.190 | FA 8:0 to FA 10:0 Ratio | 0.007984715 |
| CountryMexico | -0.402 | 0.106 | -3.783 | 0.214 | PC O-28:1 | 0.00942312 |
| CountryMexico | -0.472 | 0.125 | -3.773 | 0.184 | PG 20:3_20:4 | 0.009706144 |
| CountryMexico | -0.652 | 0.174 | -3.757 | 0.176 | PE 36:1 | 0.010166085 |
| CountryMexico | 0.512 | 0.136 | 3.755 | 0.226 | Carnitine Uptake Defect (NBS) | 0.010166085 |
| CountryMexico | -0.909 | 0.243 | -3.749 | 0.272 | LPE P-20:0 | 0.010334332 |
| CountryMexico | 0.694 | 0.185 | 3.741 | 0.254 | C10:1 | 0.010527096 |
| CountryMexico | 0.737 | 0.199 | 3.701 | 0.223 | C10 | 0.011964854 |
| CountryMexico | -0.455 | 0.123 | -3.691 | 0.309 | LPC 18:2 | 0.012316522 |
| CountryMexico | 0.730 | 0.198 | 3.689 | 0.212 | C8 | 0.012328755 |
| CountryMexico | -0.364 | 0.099 | -3.683 | 0.174 | SM 35:1 | 0.012467943 |
| CountryMexico | -0.409 | 0.111 | -3.681 | 0.216 | Sum of MUFA-LPCs | 0.012498794 |
| CountryMexico | 0.651 | 0.178 | 3.662 | 0.242 | Glu Acetylation | 0.013240178 |
| CountryMexico | -0.702 | 0.192 | -3.655 | 0.166 | FA 20:5n-3 (EPA) | 0.013359103 |
| CountryMexico | -0.415 | 0.114 | -3.654 | 0.221 | LPC 18:1 | 0.013359103 |
| CountryMexico | -0.246 | 0.067 | -3.656 | 0.166 | PG 16:0_20:4 | 0.013359103 |
| CountryMexico | -0.592 | 0.162 | -3.645 | 0.188 | PE 30:0 | 0.013687122 |
| CountryMexico | -0.354 | 0.097 | -3.642 | 0.167 | SM 36:1 | 0.013748118 |
| CountryMexico | -0.625 | 0.175 | -3.577 | 0.168 | FA 20:3n-9 | 0.016803702 |
| CountryMexico | -0.735 | 0.206 | -3.565 | 0.161 | FA 14:1n-5 | 0.017248456 |
| CountryMexico | 0.638 | 0.179 | 3.564 | 0.251 | VLCAD Deficiency (NBS) | 0.017248456 |
| CountryMexico | -0.720 | 0.202 | -3.560 | 0.176 | 3-IAA | 0.017418926 |
| CountryMexico | -0.309 | 0.087 | -3.546 | 0.160 | PC 36:4 | 0.018118459 |
| CountryMexico | -1.228 | 0.347 | -3.544 | 0.188 | PE 34:0 | 0.018134033 |
| CountryMexico | -0.403 | 0.114 | -3.540 | 0.160 | N-Ac-Met | 0.01820223 |
| CountryMexico | 0.344 | 0.097 | 3.534 | 0.187 | Uric Acid to Creatinine Ratio | 0.018466998 |
| CountryMexico | -0.464 | 0.132 | -3.516 | 0.190 | PG 20:4_22:4 | 0.019438871 |
| CountryMexico | -0.601 | 0.172 | -3.499 | 0.188 | Cer d16:1/22:0 | 0.020373295 |
| CountryMexico | -0.828 | 0.237 | -3.493 | 0.170 | CE 20:5 | 0.02062523 |
| CountryMexico | -0.516 | 0.148 | -3.482 | 0.201 | LPC 20:3 | 0.021297211 |
| CountryMexico | -0.310 | 0.089 | -3.464 | 0.160 | Malic acid | 0.022391923 |
| CountryMexico | -0.489 | 0.142 | -3.452 | 0.158 | FA 11:0 | 0.023147786 |
| CountryMexico | -0.374 | 0.108 | -3.448 | 0.168 | PC 38:4 | 0.023327319 |
| CountryMexico | -0.350 | 0.102 | -3.442 | 0.178 | PC O-44:4 | 0.023692629 |
| CountryMexico | -0.397 | 0.115 | -3.434 | 0.179 | PG 16:2_18:1 | 0.024180578 |
| CountryMexico | -0.354 | 0.103 | -3.431 | 0.167 | SM 33:1 | 0.024298122 |
| CountryMexico | -0.211 | 0.062 | -3.405 | 0.251 | N-Ac-Asn | 0.026215225 |

| | | | | | | |
|---|---|---|---|---|---|---|
| CountryMexico | -0.582 | 0.171 | -3.397 | 0.219 | CE 17:1 | 0.026719155 |
| CountryMexico | -0.239 | 0.070 | -3.396 | 0.161 | Sum of SMs | 0.026719155 |
| CountryMexico | -0.300 | 0.089 | -3.391 | 0.195 | PC O-30:2 | 0.026818291 |
| CountryMexico | -0.383 | 0.113 | -3.394 | 0.271 | Ribose | 0.026818291 |
| CountryMexico | -0.445 | 0.131 | -3.385 | 0.177 | N-Ac-Gln | 0.02716437 |
| CountryMexico | -0.264 | 0.078 | -3.366 | 0.180 | Maleic acid | 0.028688578 |
| CountryMexico | 0.744 | 0.222 | 3.349 | 0.189 | Asp Methylation | 0.030142725 |
| CountryMexico | -0.306 | 0.092 | -3.339 | 0.259 | Cit Synthesis | 0.030635396 |
| CountryMexico | 0.306 | 0.092 | 3.339 | 0.259 | OTC Deficiency (NBS) | 0.030635396 |
| CountryMexico | -0.230 | 0.069 | -3.334 | 0.159 | Sum of EC-FA SMs | 0.031027974 |
| CountryMexico | -1.019 | 0.306 | -3.327 | 0.219 | LPE P-17:0 | 0.031469371 |
| CountryMexico | -0.255 | 0.077 | -3.325 | 0.213 | Sum of VLCFA-SMs | 0.031469371 |
| CountryMexico | -0.262 | 0.079 | -3.317 | 0.159 | PE 30:1 | 0.032098097 |
| CountryMexico | -0.312 | 0.094 | -3.309 | 0.210 | SM 42:1 | 0.032550216 |
| CountryMexico | -0.159 | 0.048 | -3.309 | 0.238 | Ratio of His to HCys+Phe+Sar | 0.032550216 |
| CountryMexico | -0.486 | 0.147 | -3.305 | 0.174 | Hypoxanthine | 0.032639435 |
| CountryMexico | -0.232 | 0.070 | -3.298 | 0.149 | Sum of UFA-PCs O | 0.03316137 |
| CountryMexico | -0.400 | 0.122 | -3.263 | 0.140 | Cer d18:1/20:0 | 0.036451251 |
| CountryMexico | -0.316 | 0.097 | -3.265 | 0.149 | Sum of OC-FA SMs | 0.036451251 |
| CountryMexico | -0.231 | 0.071 | -3.252 | 0.140 | Sum of LCFA-SMs | 0.037447885 |
| CountryMexico | -1.095 | 0.337 | -3.245 | 0.138 | UDCA Synthesis from CDCA | 0.038123927 |
| CountryMexico | 0.686 | 0.212 | 3.233 | 0.244 | C12:1 | 0.039390813 |
| CountryMexico | -0.310 | 0.096 | -3.215 | 0.165 | PC O-42:4 | 0.041178681 |
| CountryMexico | -0.298 | 0.093 | -3.210 | 0.133 | PI 18:0_20:4 | 0.041690882 |
| CountryMexico | 0.534 | 0.166 | 3.208 | 0.230 | GABA Synthesis | 0.041807462 |
| CountryMexico | -0.650 | 0.203 | -3.200 | 0.173 | 3-Met-adipic acid | 0.04243752 |
| CountryMexico | -0.245 | 0.076 | -3.201 | 0.284 | Citrullinemia | 0.04243752 |
| CountryMexico | -0.550 | 0.173 | -3.175 | 0.143 | Hex-Cer d18:1/14:0 | 0.045381038 |
| CountryMexico | -0.380 | 0.120 | -3.172 | 0.140 | PG 20:4_20:4 | 0.045602398 |
| CountryMexico | -1.182 | 0.375 | -3.154 | 0.136 | Phenylacetylglutamine | 0.047966853 |
| CountryMexico | -0.427 | 0.135 | -3.153 | 0.143 | PE 35:1 | 0.047966853 |
| CountryMexico | -0.629 | 0.200 | -3.141 | 0.191 | FA 5:0-3M | 0.049255356 |
| CountryMexico | -0.248 | 0.079 | -3.137 | 0.139 | Sum of PUFA-PCs O | 0.049579085 |
| CountryMexico | -0.245 | 0.079 | -3.118 | 0.138 | PC O-40:3 | 0.051565481 |
| CountryMexico | -0.374 | 0.120 | -3.120 | 0.123 | PC O-44:5 | 0.051565481 |
| CountryMexico | -0.666 | 0.215 | -3.092 | 0.161 | Quinaldic acid | 0.055288358 |
| CountryMexico | 0.291 | 0.094 | 3.080 | 0.169 | a-Ketoglutarate to Citrate Ratio | 0.05665583 |
| CountryMexico | -0.320 | 0.105 | -3.066 | 0.142 | PC 42:1 | 0.058534305 |
| CountryMexico | -0.513 | 0.168 | -3.056 | 0.133 | CE 20:4 | 0.059419095 |
| CountryMexico | -0.283 | 0.093 | -3.052 | 0.166 | PC 36:1 | 0.059883523 |
| CountryMexico | 0.472 | 0.156 | 3.034 | 0.274 | HCit | 0.062928557 |

| | | | | | | |
|---|---|---|---|---|---|---|
| CountryMexico | -0.335 | 0.111 | -3.021 | 0.133 | PC O-36:5 | 0.065230085 |
| CountryMexico | -0.396 | 0.132 | -3.010 | 0.177 | PE P-16:0/18:1 | 0.066739606 |
| CountryMexico | -0.523 | 0.175 | -2.992 | 0.178 | Cer d16:1/24:0 | 0.070108224 |
| CountryMexico | -0.506 | 0.169 | -2.987 | 0.119 | Hex2Cer d18:1/18:0 | 0.071010092 |
| CountryMexico | -0.402 | 0.136 | -2.954 | 0.189 | LPE 20:5 | 0.076739809 |
| CountryMexico | -0.347 | 0.117 | -2.956 | 0.141 | LPE 22:5 | 0.076739809 |
| CountryMexico | 0.609 | 0.207 | 2.939 | 0.200 | FA 5:0-4M | 0.078865473 |
| CountryMexico | 0.734 | 0.250 | 2.936 | 0.163 | C14:2 | 0.079103526 |
| CountryMexico | -0.324 | 0.110 | -2.937 | 0.132 | LPC 24:0 | 0.079103526 |
| CountryMexico | -0.623 | 0.213 | -2.929 | 0.112 | TG 20:4_36:4 | 0.08037427 |
| CountryMexico | -0.734 | 0.251 | -2.925 | 0.111 | TG 18:0_38:6 | 0.081022178 |
| CountryMexico | -0.346 | 0.118 | -2.920 | 0.175 | LPC 26:1 | 0.081999863 |
| CountryMexico | -0.692 | 0.238 | -2.905 | 0.143 | PE P-18:0/20:5 | 0.085100007 |
| CountryMexico | -0.303 | 0.105 | -2.894 | 0.133 | SM 41:2 | 0.086696016 |
| CountryMexico | -0.263 | 0.091 | -2.889 | 0.201 | Hex2Cer d18:1/24:1 | 0.087792435 |
| CountryMexico | -0.230 | 0.080 | -2.879 | 0.171 | SM 42:2 | 0.08964354 |
| CountryMexico | -0.568 | 0.198 | -2.875 | 0.142 | 2-OH-Isobutyric acid | 0.090374294 |
| CountryMexico | -0.357 | 0.125 | -2.865 | 0.112 | PC 42:0 | 0.092207633 |
| CountryMexico | -0.238 | 0.084 | -2.838 | 0.110 | PC O-40:2 | 0.098299909 |
| CountryMexico | -0.466 | 0.164 | -2.837 | 0.174 | PA 18:0_18:3 | 0.098319904 |
| CountryMexico | -1.090 | 0.385 | -2.828 | 0.126 | Cinnamoyl-Gly | 0.099806131 |
| CountryMexico | -0.381 | 0.135 | -2.827 | 0.126 | PG 16:1_18:0 | 0.099806131 |
| CountryMexico | -0.209 | 0.074 | -2.826 | 0.116 | SM 34:1 | 0.099806131 |
| CountryMexico | -0.440 | 0.157 | -2.810 | 0.142 | PE P-18:1/20:4 | 0.103202262 |
| CountryMexico | -0.274 | 0.098 | -2.803 | 0.108 | LPI 22:1 | 0.104836644 |
| CountryMexico | -0.207 | 0.074 | -2.788 | 0.155 | Thr | 0.108016596 |
| CountryMexico | -0.106 | 0.038 | -2.788 | 0.142 | Sum of Non-Essential AAs | 0.108016596 |
| CountryMexico | -0.159 | 0.057 | -2.780 | 0.134 | GSH Constituents | 0.109152854 |
| CountryMexico | -0.561 | 0.202 | -2.775 | 0.126 | FA 20:4n-6 (AA) | 0.11044385 |
| CountryMexico | -0.197 | 0.071 | -2.772 | 0.117 | PG 18:1_20:0 | 0.111018107 |
| CountryMexico | -0.264 | 0.096 | -2.764 | 0.106 | Sum of VLCFA-LPIs | 0.112706714 |
| CountryMexico | 0.651 | 0.237 | 2.752 | 0.183 | C14:1 | 0.115662643 |
| CountryMexico | -0.250 | 0.091 | -2.749 | 0.102 | N-Ac-Val | 0.115662643 |
| CountryMexico | -0.370 | 0.134 | -2.750 | 0.142 | LPE 18:1 | 0.115662643 |
| CountryMexico | -0.434 | 0.158 | -2.750 | 0.109 | PC O-38:2 | 0.115662643 |
| CountryMexico | -0.102 | 0.037 | -2.748 | 0.143 | Sum of Solely Glucogenic AAs | 0.115862544 |
| CountryMexico | -0.517 | 0.188 | -2.746 | 0.118 | N2-Ac-Lys | 0.115895196 |
| CountryMexico | -0.238 | 0.087 | -2.744 | 0.108 | PC O-42:2 | 0.11631983 |
| CountryMexico | -0.405 | 0.148 | -2.740 | 0.118 | Putrescine Synthesis | 0.116945075 |
| CountryMexico | 0.468 | 0.171 | 2.735 | 0.215 | Lys Carbamylation | 0.118113224 |
| CountryMexico | -0.776 | 0.284 | -2.729 | 0.129 | Cyclic Nucleotides Ratio | 0.119093074 |

| | | | | | | |
|---|---|---|---|---|---|---|
| CountryMexico | -0.255 | 0.094 | -2.723 | 0.156 | Oxalic acid | 0.120401096 |
| CountryMexico | -0.285 | 0.105 | -2.713 | 0.145 | PG 16:0_16:1 | 0.122658953 |
| CountryMexico | -0.244 | 0.090 | -2.709 | 0.124 | PC O-32:2 | 0.123291026 |
| CountryMexico | -0.162 | 0.060 | -2.705 | 0.152 | N-Ac-Ala | 0.124121311 |
| CountryMexico | -0.554 | 0.206 | -2.698 | 0.132 | Cer d16:1/20:0 | 0.125785351 |
| CountryMexico | -0.282 | 0.105 | -2.697 | 0.130 | PE 44:7 | 0.125785351 |
| CountryMexico | -0.307 | 0.114 | -2.683 | 0.113 | SM 43:1 | 0.12976206 |
| CountryMexico | -0.626 | 0.234 | -2.678 | 0.149 | LPE 17:1 | 0.130720386 |
| CountryMexico | -0.221 | 0.082 | -2.678 | 0.158 | PA 18:0_18:1 | 0.130720386 |
| CountryMexico | -0.815 | 0.305 | -2.676 | 0.120 | LPE 14:0 | 0.130874318 |
| CountryMexico | -0.306 | 0.114 | -2.675 | 0.110 | PC O-44:3 | 0.130874318 |
| CountryMexico | 0.493 | 0.185 | 2.672 | 0.138 | Benzoic acid | 0.131537778 |
| CountryMexico | -0.286 | 0.107 | -2.666 | 0.107 | SM 41:1 | 0.133288247 |
| CountryMexico | -0.356 | 0.134 | -2.657 | 0.122 | LPC 16:1 | 0.13432796 |
| CountryMexico | -0.456 | 0.172 | -2.658 | 0.106 | PE 33:0 | 0.13432796 |
| CountryMexico | 0.582 | 0.220 | 2.652 | 0.177 | SPB d17:0 | 0.135656842 |
| CountryMexico | -0.738 | 0.279 | -2.646 | 0.162 | TG 20:5_36:2 | 0.137344146 |
| CountryMexico | -0.376 | 0.142 | -2.642 | 0.092 | FA 24:0 | 0.138407451 |
| CountryMexico | -0.502 | 0.191 | -2.630 | 0.173 | Creatinine to Bilirubin Ratio | 0.141525292 |
| CountryMexico | -0.412 | 0.157 | -2.621 | 0.125 | SM 44:1 | 0.144074775 |
| CountryMexico | -0.371 | 0.142 | -2.619 | 0.163 | LPC 14:0 | 0.14436372 |
| CountryMexico | -0.248 | 0.095 | -2.615 | 0.130 | PC O-42:3 | 0.145506878 |
| CountryMexico | -0.348 | 0.134 | -2.605 | 0.135 | N-Ac-His | 0.148680365 |
| CountryMexico | -0.232 | 0.089 | -2.602 | 0.109 | Creatinine | 0.148992586 |
| CountryMexico | -0.327 | 0.126 | -2.600 | 0.110 | PI 16:0_17:1 | 0.14904426 |
| CountryMexico | -0.224 | 0.086 | -2.594 | 0.102 | Pyruvic acid | 0.151070225 |
| CountryMexico | -0.285 | 0.110 | -2.592 | 0.107 | PC 40:4 | 0.151299009 |
| CountryMexico | -0.382 | 0.148 | -2.584 | 0.112 | PE 34:4 | 0.153497544 |
| CountryMexico | -0.216 | 0.084 | -2.585 | 0.179 | PC 42:2 | 0.153497544 |
| CountryMexico | -0.422 | 0.164 | -2.573 | 0.099 | PE 34:1 | 0.156763324 |
| CountryMexico | -0.230 | 0.090 | -2.571 | 0.109 | Pyruvate to Glucose Ratio | 0.15679762 |
| CountryMexico | 0.684 | 0.267 | 2.564 | 0.132 | ACOD1 Activity | 0.158021405 |
| CountryMexico | -0.445 | 0.175 | -2.546 | 0.111 | PC 36:5 | 0.163375019 |
| CountryMexico | -0.784 | 0.309 | -2.536 | 0.087 | PC O-38:1 | 0.165000513 |
| CountryMexico | -0.391 | 0.154 | -2.537 | 0.116 | Serotonin Pathway Activity | 0.165000513 |
| CountryMexico | 0.515 | 0.204 | 2.526 | 0.160 | C12 | 0.167665947 |
| CountryMexico | -0.327 | 0.130 | -2.516 | 0.141 | PE 38:0 | 0.171588018 |
| CountryMexico | -0.257 | 0.103 | -2.502 | 0.097 | Sum of VLCFA-LPCs | 0.176191849 |
| CountryMexico | -0.250 | 0.101 | -2.466 | 0.144 | Choline | 0.188570304 |
| CountryMexico | -0.117 | 0.048 | -2.456 | 0.094 | Gln | 0.191261662 |
| CountryMexico | -1.323 | 0.540 | -2.450 | 0.119 | SPBP d18:1 | 0.19356034 |

| | | | | | | |
|---|---|---|---|---|---|---|
| CountryMexico | -0.237 | 0.097 | -2.443 | 0.116 | PC O-36:4 | 0.195753829 |
| CountryMexico | -0.445 | 0.183 | -2.434 | 0.178 | KYNA DH | 0.197931998 |
| CountryMexico | -0.270 | 0.111 | -2.431 | 0.147 | PG 16:1_18:1 | 0.198520495 |
| CountryMexico | -1.387 | 0.573 | -2.422 | 0.124 | SPBP d18:1 to Cer d18:1/24:1 Ratio | 0.201927762 |
| CountryMexico | -0.252 | 0.104 | -2.420 | 0.123 | beta-Ala | 0.202438567 |
| CountryMexico | -0.403 | 0.167 | -2.417 | 0.115 | CE 18:3 | 0.203470175 |
| CountryMexico | -1.193 | 0.495 | -2.409 | 0.080 | TLCA Synthesis from CDCA | 0.205890574 |
| CountryMexico | -0.566 | 0.235 | -2.408 | 0.088 | PG 16:1_22:1 | 0.206161957 |
| CountryMexico | -0.530 | 0.221 | -2.402 | 0.137 | LPS 18:1 | 0.208533965 |
| CountryMexico | -0.266 | 0.111 | -2.401 | 0.077 | PG 20:4_22:1 | 0.208533965 |
| CountryMexico | -0.171 | 0.072 | -2.388 | 0.120 | N6-Ac-Lys | 0.212468615 |
| CountryMexico | -0.218 | 0.092 | -2.380 | 0.147 | PC O-38:3 | 0.215710513 |
| CountryMexico | -0.633 | 0.266 | -2.379 | 0.089 | TG 20:5_34:2 | 0.215934328 |
| CountryMexico | -0.563 | 0.237 | -2.375 | 0.132 | THDCA | 0.217260355 |
| CountryMexico | 0.279 | 0.118 | 2.373 | 0.126 | PI 18:0_20:3 | 0.21795447 |
| CountryMexico | -0.565 | 0.239 | -2.364 | 0.151 | Cer d18:0/22:0 | 0.221000389 |
| CountryMexico | -0.350 | 0.148 | -2.362 | 0.080 | PE 36:5 | 0.221984548 |
| CountryMexico | -0.388 | 0.164 | -2.359 | 0.075 | FA 14:0 | 0.22294524 |
| CountryMexico | 0.280 | 0.119 | 2.358 | 0.134 | C18:1 | 0.2230741 |
| CountryMexico | -0.262 | 0.112 | -2.349 | 0.115 | PE 44:6 | 0.226053848 |
| CountryMexico | -0.253 | 0.108 | -2.350 | 0.116 | PC 40:2 | 0.226053848 |
| CountryMexico | -0.494 | 0.211 | -2.343 | 0.084 | EMA (NBS) | 0.227740132 |
| CountryMexico | -0.674 | 0.288 | -2.340 | 0.077 | TG 20:5_36:3 | 0.228479796 |
| CountryMexico | 0.305 | 0.131 | 2.337 | 0.086 | Phenylpyruvate to Citrate Ratio | 0.229759165 |
| CountryMexico | -0.251 | 0.108 | -2.333 | 0.080 | Arg | 0.23136634 |
| CountryMexico | -0.845 | 0.363 | -2.330 | 0.180 | Hex3Cer d18:1/22:0 | 0.232187416 |
| CountryMexico | 0.719 | 0.309 | 2.326 | 0.112 | N1,N12-Di-Ac-Spermine | 0.233732297 |
| CountryMexico | -0.185 | 0.080 | -2.312 | 0.108 | PC O-34:1 | 0.239942948 |
| CountryMexico | -0.513 | 0.223 | -2.300 | 0.091 | FA 4:0-3M | 0.245022732 |
| CountryMexico | 0.478 | 0.208 | 2.299 | 0.198 | TG 18:1_36:2 | 0.245022732 |
| CountryMexico | -1.165 | 0.508 | -2.294 | 0.102 | SPBP d16:1 | 0.247792943 |
| CountryMexico | -0.729 | 0.319 | -2.288 | 0.147 | Hex-Cer d16:1/20:0 | 0.250962813 |
| CountryMexico | -0.363 | 0.159 | -2.286 | 0.090 | PE 38:3 | 0.251478949 |
| CountryMexico | -0.343 | 0.150 | -2.281 | 0.144 | Cystine | 0.253616715 |
| CountryMexico | -0.371 | 0.163 | -2.280 | 0.153 | Cystine Synthesis | 0.253845884 |
| CountryMexico | -0.166 | 0.073 | -2.278 | 0.093 | Sum of PUFA-PC (O)s | 0.254602188 |
| CountryMexico | -1.106 | 0.487 | -2.268 | 0.092 | p-Cresol glucuronide | 0.256300691 |
| CountryMexico | -0.182 | 0.080 | -2.268 | 0.098 | Gly Synthesis | 0.256300691 |
| CountryMexico | -0.256 | 0.113 | -2.260 | 0.094 | PC 30:0 | 0.2600802 |
| CountryMexico | -0.226 | 0.100 | -2.256 | 0.106 | PC 42:6 | 0.260901977 |
| CountryMexico | 0.296 | 0.131 | 2.252 | 0.105 | Carnitine to Creatinine Ratio | 0.261861322 |

| | | | | | | |
|---|---|---|---|---|---|---|
| CountryMexico | -0.207 | 0.092 | -2.248 | 0.113 | GABA | 0.263160934 |
| CountryMexico | -0.148 | 0.066 | -2.246 | 0.101 | TAT Activity | 0.263387906 |
| CountryMexico | -0.350 | 0.157 | -2.234 | 0.192 | 3-Met-glutaric acid | 0.26906513 |
| CountryMexico | -0.421 | 0.190 | -2.222 | 0.127 | N-Met-Asp | 0.271123817 |
| CountryMexico | 0.236 | 0.106 | 2.221 | 0.084 | PI 18:2_22:1 | 0.271123817 |
| CountryMexico | -0.457 | 0.206 | -2.225 | 0.102 | PC 24:0 | 0.271123817 |
| CountryMexico | -0.219 | 0.098 | -2.226 | 0.090 | PC 42:4 | 0.271123817 |
| CountryMexico | -0.468 | 0.211 | -2.218 | 0.131 | Imidazolepropionic acid | 0.271822195 |
| CountryMexico | -0.447 | 0.202 | -2.217 | 0.142 | Cer d18:2/18:0 | 0.272122345 |
| CountryMexico | -0.515 | 0.234 | -2.206 | 0.158 | Cer d16:1/23:0 | 0.276925993 |
| CountryMexico | -0.230 | 0.104 | -2.199 | 0.073 | Hex-Cer d18:1/24:1 | 0.27973733 |
| CountryMexico | -0.264 | 0.120 | -2.196 | 0.074 | PI 18:1_20:4 | 0.280207784 |
| CountryMexico | 0.130 | 0.059 | 2.193 | 0.121 | LPI 16:0 | 0.280936507 |
| CountryMexico | -0.199 | 0.091 | -2.189 | 0.071 | PS 36:4 | 0.282418394 |
| CountryMexico | -0.589 | 0.269 | -2.188 | 0.136 | TG 22:3_30:2 | 0.282418394 |
| CountryMexico | -0.392 | 0.181 | -2.169 | 0.093 | LPE 18:3 | 0.292068466 |
| CountryMexico | -0.725 | 0.336 | -2.156 | 0.079 | Met-IAA | 0.29794482 |
| CountryMexico | -0.234 | 0.109 | -2.146 | 0.073 | SM 36:2 | 0.303004764 |
| CountryMexico | 0.422 | 0.197 | 2.141 | 0.154 | Argininic acid | 0.305513472 |
| CountryMexico | 0.411 | 0.192 | 2.138 | 0.065 | Protection against MDD | 0.306211215 |
| CountryMexico | -0.233 | 0.109 | -2.128 | 0.068 | Hex-Cer d18:1/16:0 | 0.31234382 |
| CountryMexico | -0.246 | 0.116 | -2.118 | 0.120 | Guanidinoacetic acid | 0.317484346 |
| CountryMexico | 0.229 | 0.108 | 2.116 | 0.136 | Betaine | 0.317903377 |
| CountryMexico | 0.358 | 0.170 | 2.106 | 0.111 | PA 17:1_18:2 | 0.322153345 |
| CountryMexico | 0.350 | 0.167 | 2.100 | 0.165 | C16:1 | 0.32254172 |
| CountryMexico | -0.588 | 0.280 | -2.098 | 0.143 | Adipic acid | 0.32254172 |
| CountryMexico | -0.180 | 0.086 | -2.102 | 0.060 | PC O-38:5 | 0.32254172 |
| CountryMexico | -0.135 | 0.064 | -2.099 | 0.146 | PG 16:0_19:1 | 0.32254172 |
| CountryMexico | 0.155 | 0.074 | 2.093 | 0.102 | MRC Disorders | 0.324581265 |
| CountryMexico | 0.408 | 0.195 | 2.089 | 0.203 | DG 18:1_18:1 | 0.324701793 |
| CountryMexico | 0.374 | 0.180 | 2.086 | 0.139 | PA 17:1_18:1 | 0.325067814 |
| CountryMexico | 1.051 | 0.504 | 2.086 | 0.090 | TG 20:0_34:1 | 0.325067814 |
| CountryMexico | 0.799 | 0.384 | 2.081 | 0.118 | DG 16:1_18:1 | 0.327158618 |
| CountryMexico | -0.237 | 0.114 | -2.081 | 0.059 | Met Oxidation | 0.327158618 |
| CountryMexico | -0.336 | 0.162 | -2.077 | 0.083 | N-Ac-Leu | 0.328295425 |
| CountryMexico | -0.174 | 0.084 | -2.067 | 0.059 | Butyric Acid to Isobutyric Acid Ratio | 0.333773018 |
| CountryMexico | 0.285 | 0.138 | 2.066 | 0.070 | Cortisone | 0.333898916 |
| CountryMexico | -0.822 | 0.399 | -2.063 | 0.064 | SPB d16:0 | 0.334809156 |
| CountryMexico | 0.217 | 0.105 | 2.061 | 0.076 | Asymmetrical Arg Methylation | 0.334809156 |
| CountryMexico | -0.132 | 0.064 | -2.060 | 0.246 | Ratio of PUFA-LPCs to MUFA-LPCs | 0.334809156 |
| CountryMexico | -0.185 | 0.090 | -2.057 | 0.111 | PC 40:3 | 0.335712429 |

| | | | | | | |
|---|---|---|---|---|---|---|
| CountryMexico | -0.495 | 0.241 | -2.052 | 0.082 | TG 20:4_34:3 | 0.338515365 |
| CountryMexico | -0.266 | 0.130 | -2.044 | 0.103 | LPE 22:6 | 0.341999224 |
| CountryMexico | 0.166 | 0.082 | 2.040 | 0.135 | PG 18:1_22:5 | 0.344228053 |
| CountryMexico | 0.681 | 0.337 | 2.023 | 0.155 | C14:1-OH | 0.354990132 |
| CountryMexico | -0.077 | 0.038 | -2.022 | 0.099 | Sum of AAs | 0.355253984 |
| CountryMexico | -0.227 | 0.113 | -2.013 | 0.063 | Cer d18:1/18:0 | 0.359857465 |
| CountryMexico | -0.134 | 0.067 | -1.998 | 0.070 | PG 16:0_20:3 | 0.368588728 |
| CountryMexico | -0.420 | 0.210 | -1.996 | 0.070 | TG 20:4_34:2 | 0.369504941 |
| CountryMexico | -0.171 | 0.086 | -1.994 | 0.118 | LPI 18:0 | 0.370785157 |
| CountryMexico | -0.686 | 0.345 | -1.992 | 0.126 | TG 20:4_33:2 | 0.371789294 |
| CountryMexico | -0.152 | 0.076 | -1.991 | 0.090 | Sum of PUFA-PCs | 0.371789294 |
| CountryMexico | -0.340 | 0.171 | -1.982 | 0.066 | Methylmalonic acid | 0.37535574 |
| CountryMexico | -0.424 | 0.215 | -1.975 | 0.092 | FA 10:0 | 0.37945946 |
| CountryMexico | 0.260 | 0.132 | 1.969 | 0.067 | PI 18:1_18:2 | 0.380924964 |
| CountryMexico | 0.209 | 0.107 | 1.955 | 0.166 | SCD-1 index | 0.387974646 |
| CountryMexico | 0.417 | 0.214 | 1.947 | 0.131 | PA 17:2_18:1 | 0.391191077 |
| CountryMexico | -0.197 | 0.101 | -1.946 | 0.164 | NOS activity | 0.391383284 |
| CountryMexico | -0.591 | 0.304 | -1.944 | 0.055 | TLCA | 0.39248105 |
| CountryMexico | -0.091 | 0.047 | -1.943 | 0.054 | Ratio of Non-Essential to Essential AAs | 0.39248105 |
| CountryMexico | -0.234 | 0.121 | -1.932 | 0.091 | PA 18:2_20:2 | 0.398795828 |
| CountryMexico | -0.210 | 0.109 | -1.928 | 0.067 | PC O-40:6 | 0.40123095 |
| CountryMexico | -0.361 | 0.187 | -1.927 | 0.053 | TG 20:4_36:3 | 0.40139247 |
| CountryMexico | 0.221 | 0.115 | 1.926 | 0.115 | PA 18:2_22:1 | 0.401649432 |
| CountryMexico | -0.193 | 0.101 | -1.913 | 0.066 | Hex3Cer d18:1/16:0 | 0.408929815 |
| CountryMexico | -0.395 | 0.207 | -1.903 | 0.083 | BABA | 0.414574407 |
| CountryMexico | -0.142 | 0.075 | -1.900 | 0.120 | SDMA | 0.416511348 |
| CountryMexico | -0.607 | 0.319 | -1.900 | 0.088 | Phenylacetic acid | 0.416511348 |
| CountryMexico | -0.511 | 0.270 | -1.894 | 0.097 | LPS 20:5 | 0.420127059 |
| CountryMexico | -0.124 | 0.066 | -1.892 | 0.191 | N-Ac-Asp | 0.421358496 |
| CountryMexico | -0.440 | 0.234 | -1.883 | 0.091 | PE P-18:1/20:5 | 0.425307374 |
| CountryMexico | -0.394 | 0.209 | -1.882 | 0.143 | Imidazolepropionic Acid Synthesis | 0.425307374 |
| CountryMexico | -0.446 | 0.238 | -1.877 | 0.054 | TG 18:1_38:7 | 0.427248556 |
| CountryMexico | 0.143 | 0.077 | 1.867 | 0.078 | BCAT Leu Catabolism | 0.432428856 |
| CountryMexico | -0.142 | 0.076 | -1.867 | 0.047 | Orn | 0.432429327 |
| CountryMexico | -0.225 | 0.120 | -1.866 | 0.149 | 5-HIAA | 0.432429327 |
| CountryMexico | -0.165 | 0.088 | -1.859 | 0.049 | Gly | 0.43513897 |
| CountryMexico | -0.275 | 0.148 | -1.857 | 0.067 | DG 17:0_18:1 | 0.435283945 |
| CountryMexico | -0.225 | 0.121 | -1.856 | 0.066 | Hex-Cer d18:1/24:0 | 0.435298039 |
| CountryMexico | 0.243 | 0.131 | 1.855 | 0.174 | LPI 14:1 | 0.435326371 |
| CountryMexico | -0.259 | 0.140 | -1.854 | 0.078 | PI 18:0_18:3 | 0.435330836 |
| CountryMexico | 0.286 | 0.154 | 1.854 | 0.173 | PG 15:0_18:1 | 0.435330836 |

| | | | | | | |
|---|---|---|---|---|---|---|
| CountryMexico | -0.175 | 0.095 | -1.850 | 0.065 | FA 7:0 | 0.437091041 |
| CountryMexico | -0.334 | 0.181 | -1.844 | 0.056 | FA 15:0 | 0.438595627 |
| CountryMexico | -0.118 | 0.064 | -1.837 | 0.100 | PE 40:3 | 0.442316543 |
| CountryMexico | 0.304 | 0.166 | 1.836 | 0.122 | AABA Synthesis | 0.442500939 |
| CountryMexico | 0.135 | 0.073 | 1.835 | 0.165 | a-Ketoglutaric acid | 0.442616694 |
| CountryMexico | -0.156 | 0.085 | -1.830 | 0.094 | Citric acid | 0.444369342 |
| CountryMexico | -0.309 | 0.169 | -1.830 | 0.102 | TG 20:4_36:2 | 0.444369342 |
| CountryMexico | -0.524 | 0.287 | -1.830 | 0.118 | TG 20:5_34:1 | 0.444369342 |
| CountryMexico | -0.724 | 0.397 | -1.823 | 0.125 | TG 20:4_30:0 | 0.448196234 |
| CountryMexico | 0.260 | 0.143 | 1.819 | 0.123 | PA 18:0_18:2 | 0.449621157 |
| CountryMexico | -0.321 | 0.176 | -1.818 | 0.103 | Sum MCFA | 0.449621157 |
| CountryMexico | -0.185 | 0.103 | -1.804 | 0.095 | PG 18:1_22:3 | 0.458659157 |
| CountryMexico | -0.143 | 0.080 | -1.793 | 0.089 | FA 9:0 | 0.46546037 |
| CountryMexico | -0.388 | 0.217 | -1.789 | 0.060 | Mevalonic acid | 0.467581809 |
| CountryMexico | -0.232 | 0.130 | -1.787 | 0.063 | PE 44:12 | 0.469332345 |
| CountryMexico | -0.346 | 0.195 | -1.777 | 0.056 | FA 22:5n-3 (DPA) | 0.471177455 |
| CountryMexico | -0.307 | 0.173 | -1.778 | 0.089 | PE 40:6 | 0.471177455 |
| CountryMexico | -0.130 | 0.073 | -1.775 | 0.057 | Cer d18:1/16:0 | 0.47138219 |
| CountryMexico | -0.182 | 0.103 | -1.762 | 0.052 | Cer d18:1/22:0 | 0.472675477 |
| CountryMexico | -0.256 | 0.145 | -1.766 | 0.099 | PE 33:1 | 0.472675477 |
| CountryMexico | -0.117 | 0.066 | -1.762 | 0.092 | Ratio of SGA to Glucose | 0.472675477 |
| CountryMexico | 0.088 | 0.050 | 1.754 | 0.063 | MTHFR Deficiency (NBS) | 0.476925238 |
| CountryMexico | -0.183 | 0.105 | -1.750 | 0.064 | Hex-Cer d18:1/18:0 | 0.477860014 |
| CountryMexico | -0.183 | 0.105 | -1.745 | 0.059 | C4:1 | 0.478424234 |
| CountryMexico | -0.189 | 0.109 | -1.745 | 0.084 | 1-Met-His | 0.478424234 |
| CountryMexico | -0.144 | 0.082 | -1.744 | 0.060 | SM 34:2 | 0.478424234 |
| CountryMexico | -0.248 | 0.143 | -1.734 | 0.058 | PE 36:6 | 0.485320961 |
| CountryMexico | 0.390 | 0.225 | 1.734 | 0.189 | TG 18:1_34:1 | 0.485320961 |
| CountryMexico | -0.387 | 0.223 | -1.733 | 0.055 | TG 16:0_38:6 | 0.485434657 |
| CountryMexico | -0.214 | 0.124 | -1.730 | 0.068 | Sum of PUFA-CEs | 0.485991877 |
| CountryMexico | 0.510 | 0.295 | 1.727 | 0.137 | TG 16:0_38:2 | 0.487451843 |
| CountryMexico | -0.133 | 0.077 | -1.724 | 0.063 | HSer | 0.48834065 |
| CountryMexico | 0.149 | 0.087 | 1.723 | 0.110 | Phenylpyruvic acid | 0.489060918 |
| CountryMexico | -0.405 | 0.235 | -1.721 | 0.102 | PE P-16:0/20:5 | 0.489060918 |
| CountryMexico | 0.740 | 0.430 | 1.718 | 0.070 | Gly Conjugation of CA | 0.491360269 |
| CountryMexico | -0.279 | 0.163 | -1.714 | 0.057 | Indole-Lac | 0.493997658 |
| CountryMexico | -0.933 | 0.545 | -1.712 | 0.051 | p-Cresol-SO4 Synthesis | 0.495691031 |
| CountryMexico | -0.158 | 0.093 | -1.704 | 0.081 | PC 42:5 | 0.498645241 |
| CountryMexico | -0.472 | 0.278 | -1.698 | 0.043 | SM 44:2 | 0.50291887 |
| CountryMexico | -0.236 | 0.139 | -1.695 | 0.096 | PE P-18:0/22:6 | 0.505494449 |
| CountryMexico | -0.174 | 0.103 | -1.693 | 0.087 | PC 38:5 | 0.506331118 |

| | | | | | | |
|---|---|---|---|---|---|---|
| CountryMexico | -0.404 | 0.239 | -1.691 | 0.109 | Suberic acid | 0.506819519 |
| CountryMexico | -0.541 | 0.321 | -1.688 | 0.068 | Hex3Cer d18:1/18:0 | 0.507524668 |
| CountryMexico | 0.228 | 0.136 | 1.679 | 0.098 | DMG | 0.509884198 |
| CountryMexico | -0.098 | 0.058 | -1.677 | 0.069 | Asn | 0.509884198 |
| CountryMexico | -0.152 | 0.091 | -1.673 | 0.096 | Urea | 0.509884198 |
| CountryMexico | 0.178 | 0.106 | 1.674 | 0.100 | T4 | 0.509884198 |
| CountryMexico | 0.183 | 0.109 | 1.678 | 0.142 | PG 16:3_18:1 | 0.509884198 |
| CountryMexico | -0.385 | 0.230 | -1.677 | 0.040 | TG 18:2_38:6 | 0.509884198 |
| CountryMexico | -0.476 | 0.283 | -1.681 | 0.061 | TG 20:3_34:3 | 0.509884198 |
| CountryMexico | -0.271 | 0.163 | -1.666 | 0.064 | cGMP | 0.512020455 |
| CountryMexico | -0.201 | 0.121 | -1.663 | 0.040 | PI 16:1_18:0 | 0.513220124 |
| CountryMexico | -0.144 | 0.087 | -1.661 | 0.112 | Sum of MUFA-PGs | 0.513889221 |
| CountryMexico | 0.543 | 0.328 | 1.656 | 0.093 | C16:2 | 0.514137204 |
| CountryMexico | 0.254 | 0.154 | 1.653 | 0.155 | PA 16:1_18:1 | 0.514876218 |
| CountryMexico | -0.433 | 0.262 | -1.654 | 0.077 | TG 18:2_36:1 | 0.514876218 |
| CountryMexico | -0.557 | 0.337 | -1.652 | 0.106 | TG 20:3_32:2 | 0.514876218 |
| CountryMexico | -0.448 | 0.271 | -1.651 | 0.099 | TG 20:4_32:2 | 0.514876218 |
| CountryMexico | -0.208 | 0.126 | -1.648 | 0.049 | PC O-38:0 | 0.516178366 |
| CountryMexico | -0.094 | 0.057 | -1.648 | 0.101 | Sum of Dimethylated Arg | 0.516207195 |
| CountryMexico | -0.322 | 0.196 | -1.647 | 0.062 | LPE 15:0 | 0.516794868 |
| CountryMexico | 0.572 | 0.349 | 1.639 | 0.151 | TG 20:1_34:3 | 0.520958247 |
| CountryMexico | 0.300 | 0.184 | 1.628 | 0.096 | PE P-16:0/18:2 | 0.526181895 |
| CountryMexico | 0.485 | 0.298 | 1.629 | 0.144 | TG 16:1_38:4 | 0.526181895 |
| CountryMexico | 0.100 | 0.062 | 1.620 | 0.072 | PKU (NBS) | 0.530377577 |
| CountryMexico | 0.188 | 0.116 | 1.619 | 0.073 | PI 18:0_18:2 | 0.531605649 |
| CountryMexico | -0.573 | 0.355 | -1.616 | 0.061 | IsoUDCA | 0.531808872 |
| CountryMexico | -0.329 | 0.204 | -1.617 | 0.087 | LPE P-22:0 | 0.531808872 |
| CountryMexico | -0.477 | 0.295 | -1.614 | 0.090 | TG 20:4_32:1 | 0.532789484 |
| CountryMexico | 0.101 | 0.063 | 1.609 | 0.086 | PC 34:2 | 0.535583388 |
| CountryMexico | -0.819 | 0.514 | -1.595 | 0.045 | Total p-Cresol Derivatives | 0.544772388 |
| CountryMexico | -0.823 | 0.516 | -1.593 | 0.045 | p-Cresol-SO4 | 0.544806474 |
| CountryMexico | -0.319 | 0.201 | -1.590 | 0.054 | FA 12:0 | 0.54503181 |
| CountryMexico | -0.295 | 0.185 | -1.592 | 0.062 | FA 22:4n-6 | 0.54503181 |
| CountryMexico | -0.264 | 0.166 | -1.589 | 0.090 | Hex-Cer d18:2/16:0 | 0.54503181 |
| CountryMexico | -0.086 | 0.055 | -1.582 | 0.095 | Ratio of OC-FA SMs to EC-FA SMs | 0.548018874 |
| CountryMexico | 0.240 | 0.152 | 1.580 | 0.064 | PA 17:0_18:2 | 0.548820569 |
| CountryMexico | -0.157 | 0.100 | -1.580 | 0.117 | PI 17:1_18:1 | 0.548820569 |
| CountryMexico | -0.465 | 0.295 | -1.574 | 0.041 | TG 18:3_38:5 | 0.552034924 |
| CountryMexico | -0.211 | 0.134 | -1.574 | 0.176 | Pantothenic acid (B5) | 0.552034924 |
| CountryMexico | -0.181 | 0.115 | -1.574 | 0.050 | Acetic Acid to Isobutyric Acid Ratio | 0.552034924 |
| CountryMexico | -0.877 | 0.558 | -1.571 | 0.090 | DG 18:2_18:4 | 0.552485032 |

| | | | | | | |
|---|---|---|---|---|---|---|
| CountryMexico | -0.074 | 0.047 | -1.568 | 0.101 | His | 0.553838794 |
| CountryMexico | -0.621 | 0.396 | -1.569 | 0.064 | TG 20:4_34:0 | 0.553838794 |
| CountryMexico | 0.139 | 0.089 | 1.565 | 0.054 | Phenylpyruvate Synthesis | 0.556883974 |
| CountryMexico | 0.217 | 0.140 | 1.557 | 0.098 | PA 18:1_18:1 | 0.562131883 |
| CountryMexico | -0.160 | 0.103 | -1.558 | 0.088 | LPG 18:0 | 0.562131883 |
| CountryMexico | 0.241 | 0.155 | 1.554 | 0.111 | PA 17:0_18:1 | 0.563478323 |
| CountryMexico | -0.348 | 0.224 | -1.551 | 0.064 | GCDH Deficiency | 0.56494543 |
| CountryMexico | -0.384 | 0.248 | -1.548 | 0.066 | Desamino-Tyr | 0.566969333 |
| CountryMexico | -0.273 | 0.178 | -1.536 | 0.117 | CE 16:1 | 0.574518482 |
| CountryMexico | -0.224 | 0.146 | -1.538 | 0.078 | PC 34:4 | 0.574518482 |
| CountryMexico | 0.118 | 0.077 | 1.529 | 0.058 | FA 3:0-2M | 0.577618839 |
| CountryMexico | -0.866 | 0.569 | -1.523 | 0.075 | 3-EpiDCA | 0.58088135 |
| CountryMexico | 0.311 | 0.204 | 1.523 | 0.075 | PE P-16:0/20:3 | 0.58088135 |
| CountryMexico | 0.138 | 0.091 | 1.519 | 0.067 | PC 36:3 | 0.582728943 |
| CountryMexico | -0.174 | 0.114 | -1.520 | 0.051 | PC O-38:6 | 0.582728943 |
| CountryMexico | -0.250 | 0.165 | -1.516 | 0.101 | PG 16:1_16:1 | 0.582728943 |
| CountryMexico | 0.326 | 0.215 | 1.516 | 0.161 | TG 16:0_36:2 | 0.582728943 |
| CountryMexico | 0.272 | 0.180 | 1.513 | 0.117 | TG 18:2_36:2 | 0.58359756 |
| CountryMexico | 0.219 | 0.145 | 1.508 | 0.037 | PI 18:1_18:1 | 0.585318043 |
| CountryMexico | 0.354 | 0.236 | 1.503 | 0.215 | TG 16:1_36:2 | 0.588193853 |
| CountryMexico | -0.313 | 0.209 | -1.501 | 0.094 | TG 20:4_34:1 | 0.589780816 |
| CountryMexico | -0.110 | 0.073 | -1.499 | 0.077 | Sum of SphoPs | 0.590112266 |
| CountryMexico | -0.401 | 0.269 | -1.494 | 0.075 | TG 20:4_32:0 | 0.592061225 |
| CountryMexico | 0.098 | 0.066 | 1.494 | 0.066 | Ratio of MUFA-LPCs to SFA-LPCs | 0.592061225 |
| CountryMexico | -0.128 | 0.086 | -1.493 | 0.049 | PC O-36:2 | 0.592675102 |
| CountryMexico | 0.347 | 0.233 | 1.491 | 0.082 | c4-OH-Pro | 0.59388093 |
| CountryMexico | 0.133 | 0.090 | 1.489 | 0.073 | Brain Trp Availability | 0.595076706 |
| CountryMexico | 0.167 | 0.112 | 1.487 | 0.051 | Trp | 0.596251964 |
| CountryMexico | -0.585 | 0.394 | -1.487 | 0.046 | TG 20:4_36:5 | 0.596251964 |
| CountryMexico | -0.144 | 0.097 | -1.482 | 0.086 | Hex2Cer d18:1/16:0 | 0.599049007 |
| CountryMexico | 0.112 | 0.076 | 1.477 | 0.060 | Uric acid | 0.600852354 |
| CountryMexico | 0.362 | 0.245 | 1.474 | 0.065 | TG 20:0_32:3 | 0.602663432 |
| CountryMexico | -0.156 | 0.106 | -1.471 | 0.046 | PI 18:2_20:4 | 0.60424403 |
| CountryMexico | 0.236 | 0.161 | 1.466 | 0.074 | C4 | 0.60757359 |
| CountryMexico | -0.237 | 0.162 | -1.466 | 0.134 | 5-HIAA / Quinolinic Acid Ratio | 0.60757359 |
| CountryMexico | -0.360 | 0.246 | -1.464 | 0.070 | Arginino-Suc | 0.608220485 |
| CountryMexico | 0.098 | 0.067 | 1.455 | 0.055 | Met | 0.6086916 |
| CountryMexico | -0.455 | 0.312 | -1.457 | 0.141 | CE 14:0 | 0.6086916 |
| CountryMexico | -0.103 | 0.071 | -1.456 | 0.129 | PA 18:3_18:3 | 0.6086916 |
| CountryMexico | -0.140 | 0.096 | -1.457 | 0.082 | PC O-28:0 | 0.6086916 |
| CountryMexico | -0.187 | 0.128 | -1.456 | 0.057 | Picolinic Acid to Trp Ratio | 0.6086916 |

| | | | | | | |
|---|---|---|---|---|---|---|
| CountryMexico | -0.227 | 0.157 | -1.448 | 0.091 | LPE 18:2 | 0.611656345 |
| CountryMexico | -0.190 | 0.132 | -1.447 | 0.105 | Hex-Cer d18:1/18:1 | 0.611799407 |
| CountryMexico | -0.347 | 0.240 | -1.444 | 0.077 | SPHK activity (8) | 0.611832799 |
| CountryMexico | -0.386 | 0.267 | -1.443 | 0.047 | Cer d16:1/18:0 | 0.612808528 |
| CountryMexico | -0.157 | 0.109 | -1.441 | 0.103 | CE 16:0 | 0.614126863 |
| CountryMexico | 0.110 | 0.077 | 1.441 | 0.033 | Tyr | 0.614226121 |
| CountryMexico | 0.263 | 0.183 | 1.439 | 0.115 | TG 18:1_36:3 | 0.614226121 |
| CountryMexico | -0.164 | 0.115 | -1.423 | 0.066 | LPI 20:1 | 0.619637508 |
| CountryMexico | -0.147 | 0.103 | -1.424 | 0.035 | PG 18:1_20:1 | 0.619637508 |
| CountryMexico | 0.092 | 0.065 | 1.419 | 0.047 | Sum of Aromatic AAs | 0.621264515 |
| CountryMexico | -0.069 | 0.049 | -1.414 | 0.080 | Fischer Ratio | 0.62415714 |
| CountryMexico | 0.192 | 0.136 | 1.413 | 0.036 | C18:2 | 0.62426111 |
| CountryMexico | -0.202 | 0.143 | -1.409 | 0.098 | Z-3-Met-glutaconic acid | 0.625881202 |
| CountryMexico | 0.270 | 0.191 | 1.408 | 0.121 | Bilirubin | 0.625881202 |
| CountryMexico | -0.189 | 0.135 | -1.408 | 0.080 | PE 36:0 | 0.625881202 |
| CountryMexico | -0.612 | 0.437 | -1.402 | 0.156 | CA | 0.627793048 |
| CountryMexico | -0.141 | 0.101 | -1.396 | 0.178 | N8-Ac-Spermidine | 0.631895514 |
| CountryMexico | -0.146 | 0.104 | -1.397 | 0.064 | 2MBG (NBS) | 0.631895514 |
| CountryMexico | 0.302 | 0.217 | 1.393 | 0.092 | C10:2 | 0.633207468 |
| CountryMexico | -0.182 | 0.131 | -1.391 | 0.033 | HCys Synthesis | 0.633878322 |
| CountryMexico | -0.175 | 0.126 | -1.386 | 0.092 | PE P-16:0/22:6 | 0.636062823 |
| CountryMexico | -0.177 | 0.129 | -1.376 | 0.050 | Hex-Cer d18:1/23:0 | 0.641957342 |
| CountryMexico | -0.657 | 0.477 | -1.376 | 0.026 | GLCA Synthesis from CDCA | 0.641998079 |
| CountryMexico | -0.216 | 0.157 | -1.373 | 0.081 | LPE 20:3 | 0.644250121 |
| CountryMexico | -0.149 | 0.108 | -1.372 | 0.043 | Secondary 3MG Aciduria | 0.644250121 |
| CountryMexico | -0.514 | 0.375 | -1.370 | 0.068 | TG 18:3_38:6 | 0.644819909 |
| CountryMexico | -0.255 | 0.187 | -1.369 | 0.082 | MCFA PPAR Modulators | 0.644819909 |
| CountryMexico | 0.181 | 0.133 | 1.359 | 0.056 | alpha-AAA | 0.650670635 |
| CountryMexico | -0.153 | 0.112 | -1.358 | 0.056 | FA 18:0 | 0.650670635 |
| CountryMexico | -0.515 | 0.379 | -1.358 | 0.037 | PC 36:0 | 0.650670635 |
| CountryMexico | -0.278 | 0.204 | -1.362 | 0.029 | TG 18:0_36:4 | 0.650670635 |
| CountryMexico | 0.296 | 0.218 | 1.358 | 0.094 | TG 20:1_34:2 | 0.650670635 |
| CountryMexico | 0.145 | 0.107 | 1.357 | 0.035 | 4-Met-2-oxovaleric acid | 0.651770479 |
| CountryMexico | -0.158 | 0.117 | -1.354 | 0.064 | Hex-Cer d18:1/22:0 | 0.651802516 |
| CountryMexico | 0.319 | 0.235 | 1.355 | 0.029 | 3,5-DiOH-Benzoic acid | 0.651802516 |
| CountryMexico | 0.282 | 0.208 | 1.355 | 0.082 | TG 16:0_38:3 | 0.651802516 |
| CountryMexico | -0.153 | 0.113 | -1.352 | 0.045 | Ratio of MUFA-PGs to SFA-PGs | 0.652798468 |
| CountryMexico | -0.149 | 0.111 | -1.344 | 0.038 | LPG 18:2 | 0.656075836 |
| CountryMexico | 0.157 | 0.117 | 1.344 | 0.046 | Sum of Asym. and Sym. Arg Methylation | 0.656075836 |
| CountryMexico | -0.148 | 0.111 | -1.336 | 0.080 | IDO Activity | 0.66232806 |
| CountryMexico | -0.121 | 0.091 | -1.334 | 0.072 | Ratio of SG to Glucose | 0.662629197 |

| | | | | | | |
|---|---|---|---|---|---|---|
| CountryMexico | -0.391 | 0.294 | -1.330 | 0.038 | Ind-SO4 | 0.665831465 |
| CountryMexico | 0.092 | 0.069 | 1.324 | 0.141 | Glyoxylic acid | 0.670191792 |
| CountryMexico | -0.167 | 0.126 | -1.320 | 0.071 | CE 18:1 | 0.67030417 |
| CountryMexico | 0.183 | 0.138 | 1.322 | 0.076 | PI 16:0_18:2 | 0.67030417 |
| CountryMexico | -0.528 | 0.399 | -1.321 | 0.047 | TG 18:0_32:0 | 0.67030417 |
| CountryMexico | -0.239 | 0.181 | -1.322 | 0.032 | KAT A Activity | 0.67030417 |
| CountryMexico | -0.284 | 0.216 | -1.316 | 0.038 | PG 18:1_20:2 | 0.672065727 |
| CountryMexico | 0.251 | 0.192 | 1.308 | 0.119 | DG 18:1_18:2 | 0.675787164 |
| CountryMexico | -0.302 | 0.231 | -1.307 | 0.069 | TG 16:0_40:6 | 0.675787164 |
| CountryMexico | -0.430 | 0.330 | -1.303 | 0.061 | TG 17:0_32:1 | 0.675819157 |
| CountryMexico | -0.589 | 0.452 | -1.304 | 0.027 | Gly Conjugation of CDCA | 0.675819157 |
| CountryMexico | -0.218 | 0.168 | -1.300 | 0.104 | PG 17:1_18:1 | 0.676619533 |
| CountryMexico | -0.493 | 0.380 | -1.297 | 0.041 | UDCA | 0.676872741 |
| CountryMexico | -0.167 | 0.129 | -1.295 | 0.084 | Hex-Cer d18:2/22:0 | 0.677094997 |
| CountryMexico | 0.602 | 0.469 | 1.285 | 0.035 | CDCA | 0.681552609 |
| CountryMexico | 0.217 | 0.169 | 1.284 | 0.170 | N-Ac-Putrescine to N1- and N8-Ac-Spermidine Ratio | 0.681552609 |
| CountryMexico | 0.139 | 0.108 | 1.278 | 0.088 | PI 18:1_22:1 | 0.683205039 |
| CountryMexico | -0.236 | 0.185 | -1.274 | 0.073 | Glutaric acid | 0.685703126 |
| CountryMexico | -0.100 | 0.079 | -1.271 | 0.139 | PI 16:1_18:1 | 0.686727662 |
| CountryMexico | -0.102 | 0.081 | -1.268 | 0.172 | Ala | 0.688271753 |
| CountryMexico | 0.185 | 0.146 | 1.266 | 0.026 | Cortisol | 0.688688089 |
| CountryMexico | 0.346 | 0.273 | 1.267 | 0.097 | BABA Synthesis | 0.688688089 |
| CountryMexico | -0.419 | 0.332 | -1.264 | 0.045 | TG 16:0_38:7 | 0.689378866 |
| CountryMexico | -0.573 | 0.453 | -1.264 | 0.037 | Taurine Conjugation of CDCA | 0.689378866 |
| CountryMexico | 0.098 | 0.078 | 1.261 | 0.111 | PG 18:2_22:0 | 0.690878506 |
| CountryMexico | -0.885 | 0.704 | -1.257 | 0.077 | 2,5-Furandicarboxylic acid | 0.690901009 |
| CountryMexico | -0.528 | 0.421 | -1.255 | 0.032 | TDCA | 0.691612288 |
| CountryMexico | -0.138 | 0.110 | -1.254 | 0.039 | PI 18:1_22:6 | 0.691807634 |
| CountryMexico | -0.210 | 0.168 | -1.254 | 0.072 | PC 36:6 | 0.691807634 |
| CountryMexico | -0.123 | 0.098 | -1.246 | 0.070 | PG 16:0_20:5 | 0.696243556 |
| CountryMexico | -0.355 | 0.286 | -1.242 | 0.113 | LPE P-22:1 | 0.698071933 |
| CountryMexico | -0.289 | 0.234 | -1.236 | 0.087 | PE P-18:0/18:3 | 0.700948011 |
| CountryMexico | -0.156 | 0.127 | -1.232 | 0.061 | PI 16:0_20:4 | 0.701559351 |
| CountryMexico | -0.131 | 0.106 | -1.232 | 0.027 | PS 38:6 | 0.701559351 |
| CountryMexico | -0.325 | 0.264 | -1.231 | 0.082 | TG 20:3_32:0 | 0.701559351 |
| CountryMexico | -0.186 | 0.151 | -1.230 | 0.056 | PE 35:2 | 0.702233989 |
| CountryMexico | 0.103 | 0.084 | 1.224 | 0.072 | PG 18:2_20:3 | 0.706559354 |
| CountryMexico | -0.233 | 0.191 | -1.221 | 0.044 | TG 17:0_36:4 | 0.70792728 |
| CountryMexico | -0.182 | 0.150 | -1.216 | 0.102 | CE 20:3 | 0.709804529 |
| CountryMexico | 0.227 | 0.188 | 1.207 | 0.116 | TG 18:1_34:2 | 0.717056776 |
| CountryMexico | 0.077 | 0.064 | 1.200 | 0.092 | PG 16:0_18:3 | 0.71897343 |

| | | | | | | |
|---|---|---|---|---|---|---|
| CountryMexico | -0.222 | 0.186 | -1.194 | 0.028 | N-Ac-Tyr | 0.723648112 |
| CountryMexico | -0.149 | 0.126 | -1.189 | 0.040 | LPC 26:0 | 0.725275887 |
| CountryMexico | 0.137 | 0.115 | 1.188 | 0.032 | PC 34:3 | 0.725275887 |
| CountryMexico | -0.265 | 0.223 | -1.189 | 0.063 | TG 18:2_31:0 | 0.725275887 |
| CountryMexico | -0.245 | 0.206 | -1.187 | 0.110 | TG 20:3_34:2 | 0.726702784 |
| CountryMexico | 0.277 | 0.234 | 1.186 | 0.093 | Ethylmalonic acid | 0.726901406 |
| CountryMexico | 0.233 | 0.198 | 1.182 | 0.026 | Ratio Quinolinic Acid to KYNA | 0.730673005 |
| CountryMexico | -0.231 | 0.196 | -1.180 | 0.049 | LPE 20:2 | 0.732068194 |
| CountryMexico | -0.503 | 0.426 | -1.179 | 0.047 | TG 14:0_35:2 | 0.732068194 |
| CountryMexico | 0.238 | 0.203 | 1.171 | 0.075 | TG 18:3_36:2 | 0.734659259 |
| CountryMexico | -0.245 | 0.210 | -1.170 | 0.125 | Hex-Cer d18:1/26:1 | 0.735495638 |
| CountryMexico | -0.181 | 0.156 | -1.165 | 0.124 | GAA to HArg Ratio | 0.737696495 |
| CountryMexico | -0.139 | 0.119 | -1.161 | 0.028 | Met-SO | 0.738515723 |
| CountryMexico | 0.265 | 0.228 | 1.161 | 0.084 | PI 17:0_18:1 | 0.738515723 |
| CountryMexico | -0.253 | 0.219 | -1.158 | 0.041 | Malonic acid | 0.739863384 |
| CountryMexico | 0.231 | 0.202 | 1.144 | 0.095 | DG 16:0_18:1 | 0.747947347 |
| CountryMexico | -0.426 | 0.374 | -1.141 | 0.083 | TG 17:1_32:1 | 0.749595167 |
| CountryMexico | -0.129 | 0.114 | -1.137 | 0.033 | C18 | 0.751252254 |
| CountryMexico | -0.163 | 0.144 | -1.134 | 0.054 | PE P-18:0/18:1 | 0.751252254 |
| CountryMexico | -0.300 | 0.264 | -1.136 | 0.071 | TG 22:4_34:2 | 0.751252254 |
| CountryMexico | -0.141 | 0.124 | -1.133 | 0.088 | Hex-Cer d18:2/24:0 | 0.75148364 |
| CountryMexico | -0.138 | 0.122 | -1.129 | 0.048 | CE 18:2 | 0.752806895 |
| CountryMexico | 0.266 | 0.236 | 1.128 | 0.123 | Anthranilic acid | 0.752847723 |
| CountryMexico | 0.177 | 0.157 | 1.127 | 0.068 | PA 16:1_18:2 | 0.753042455 |
| CountryMexico | -0.221 | 0.197 | -1.121 | 0.046 | Kynurenic acid | 0.755264704 |
| CountryMexico | -0.203 | 0.181 | -1.119 | 0.153 | E-3-Met-glutaconic acid | 0.756364037 |
| CountryMexico | 0.089 | 0.079 | 1.116 | 0.137 | PA 16:0_19:2 | 0.758060999 |
| CountryMexico | 0.287 | 0.257 | 1.116 | 0.053 | C18:1-OH | 0.758569803 |
| CountryMexico | 0.104 | 0.094 | 1.109 | 0.058 | PG 18:1_18:3 | 0.761693161 |
| CountryMexico | 0.104 | 0.094 | 1.108 | 0.025 | Sum of BC-a-ketoacids | 0.762246381 |
| CountryMexico | -0.504 | 0.456 | -1.107 | 0.025 | TG 16:1_28:0 | 0.762811269 |
| CountryMexico | 0.277 | 0.251 | 1.103 | 0.023 | HMGCR Activity | 0.762811269 |
| CountryMexico | -0.249 | 0.226 | -1.100 | 0.027 | N-Ac-Gly | 0.763652624 |
| CountryMexico | -0.276 | 0.253 | -1.093 | 0.019 | N-Ac-Ile | 0.763652624 |
| CountryMexico | -0.579 | 0.530 | -1.094 | 0.022 | GUDCA | 0.763652624 |
| CountryMexico | -0.390 | 0.356 | -1.094 | 0.044 | NorDCA | 0.763652624 |
| CountryMexico | 0.079 | 0.072 | 1.094 | 0.045 | FA 6:0 | 0.763652624 |
| CountryMexico | -0.359 | 0.328 | -1.094 | 0.062 | LPE 19:0 | 0.763652624 |
| CountryMexico | -0.318 | 0.291 | -1.093 | 0.061 | Xanthine | 0.763652624 |
| CountryMexico | -0.130 | 0.119 | -1.091 | 0.019 | PC 38:0 | 0.764021009 |
| CountryMexico | -0.115 | 0.106 | -1.091 | 0.098 | 1-Met-His Synthesis | 0.764021009 |

| | | | | | | |
|---|---|---|---|---|---|---|
| CountryMexico | -0.194 | 0.179 | -1.082 | 0.030 | Citramalic acid | 0.769696796 |
| CountryMexico | -0.281 | 0.260 | -1.081 | 0.019 | TG 20:3_36:4 | 0.770515651 |
| CountryMexico | 0.547 | 0.506 | 1.080 | 0.049 | Taurine Conjugation of CA | 0.770857275 |
| CountryMexico | -0.332 | 0.308 | -1.075 | 0.089 | TG 20:2_34:3 | 0.772981038 |
| CountryMexico | 0.136 | 0.127 | 1.066 | 0.108 | PA 16:0_18:1 | 0.7734465 |
| CountryMexico | 0.208 | 0.195 | 1.068 | 0.059 | PA 18:1_18:3 | 0.7734465 |
| CountryMexico | 0.223 | 0.208 | 1.070 | 0.157 | TG 16:1_36:3 | 0.7734465 |
| CountryMexico | 0.079 | 0.074 | 1.066 | 0.054 | PLA activity (2) | 0.7734465 |
| CountryMexico | 0.171 | 0.161 | 1.061 | 0.056 | PI 18:1_20:2 | 0.776133059 |
| CountryMexico | 0.073 | 0.069 | 1.062 | 0.021 | BCAT Ile Catabolism | 0.776133059 |
| CountryMexico | -0.368 | 0.348 | -1.058 | 0.042 | TUDCA | 0.778530373 |
| CountryMexico | 0.136 | 0.129 | 1.053 | 0.103 | PI 16:0_20:3 | 0.7803233 |
| CountryMexico | -0.120 | 0.115 | -1.049 | 0.023 | PE 32:2 | 0.78317616 |
| CountryMexico | 0.241 | 0.230 | 1.046 | 0.154 | TG 20:1_34:1 | 0.784490321 |
| CountryMexico | 0.146 | 0.140 | 1.043 | 0.177 | N1,N8-Di-Ac-Spermidine | 0.784816986 |
| CountryMexico | -0.439 | 0.420 | -1.044 | 0.071 | TG 14:0_35:1 | 0.784816986 |
| CountryMexico | -0.465 | 0.446 | -1.044 | 0.103 | TG 22:4_32:0 | 0.784816986 |
| CountryMexico | -0.761 | 0.731 | -1.041 | 0.075 | MG 22:4 | 0.786541942 |
| CountryMexico | -0.432 | 0.418 | -1.033 | 0.093 | Trigonelline | 0.789010564 |
| CountryMexico | -0.351 | 0.340 | -1.033 | 0.100 | 7-KetoDCA | 0.789010564 |
| CountryMexico | 0.113 | 0.109 | 1.033 | 0.068 | PG 18:2_18:4 | 0.789010564 |
| CountryMexico | 0.088 | 0.085 | 1.035 | 0.066 | PG 18:2_20:4 | 0.789010564 |
| CountryMexico | -0.195 | 0.189 | -1.030 | 0.076 | Hex-Cer d18:2/18:0 | 0.789447768 |
| CountryMexico | 0.224 | 0.217 | 1.030 | 0.059 | LPI 18:3 | 0.789447768 |
| CountryMexico | 0.170 | 0.166 | 1.027 | 0.085 | DG 18:1_20:1 | 0.789904956 |
| CountryMexico | -0.516 | 0.507 | -1.019 | 0.034 | GDCA | 0.792854484 |
| CountryMexico | 0.248 | 0.243 | 1.020 | 0.142 | Kynureninase A Activity | 0.792854484 |
| CountryMexico | 0.169 | 0.166 | 1.017 | 0.053 | PE P-18:1/18:2 | 0.793046292 |
| CountryMexico | -0.256 | 0.252 | -1.017 | 0.068 | TG 16:0_33:2 | 0.793046292 |
| CountryMexico | 0.144 | 0.142 | 1.014 | 0.044 | PI 16:0_18:1 | 0.793175569 |
| CountryMexico | 0.084 | 0.083 | 1.015 | 0.148 | PG 16:0_22:2 | 0.793175569 |
| CountryMexico | -0.509 | 0.502 | -1.013 | 0.021 | PC O-30:1 | 0.793525266 |
| CountryMexico | -0.104 | 0.103 | -1.012 | 0.079 | PC 40:5 | 0.793644287 |
| CountryMexico | -0.419 | 0.417 | -1.006 | 0.080 | TG 18:0_32:1 | 0.796176473 |
| CountryMexico | 0.185 | 0.184 | 1.004 | 0.093 | TG 16:0_36:3 | 0.796433474 |
| CountryMexico | 0.124 | 0.125 | 1.000 | 0.069 | C3 | 0.799544403 |
| CountryMexico | -0.342 | 0.342 | -1.000 | 0.065 | TG 18:1_33:0 | 0.799544403 |
| CountryMexico | 0.082 | 0.082 | 0.998 | 0.048 | LPI 18:1 | 0.800131331 |
| CountryMexico | 0.113 | 0.114 | 0.995 | 0.097 | PG 18:1_20:5 | 0.800342785 |
| CountryMexico | 0.080 | 0.080 | 0.996 | 0.066 | Mannose | 0.800342785 |
| CountryMexico | 0.202 | 0.203 | 0.993 | 0.091 | LPG 14:0 | 0.800847391 |

| | | | | | | |
|---|---|---|---|---|---|---|
| CountryMexico | -0.534 | 0.540 | -0.988 | 0.018 | TG 18:1_26:0 | 0.805160419 |
| CountryMexico | -0.182 | 0.185 | -0.987 | 0.017 | PE 36:2 | 0.805861192 |
| CountryMexico | -0.290 | 0.294 | -0.986 | 0.052 | Xanthurenic acid | 0.806686742 |
| CountryMexico | -0.248 | 0.253 | -0.979 | 0.018 | TG 18:0_36:5 | 0.811336065 |
| CountryMexico | 0.178 | 0.183 | 0.973 | 0.091 | TG 18:2_34:1 | 0.815633025 |
| CountryMexico | -0.207 | 0.214 | -0.968 | 0.091 | TG 16:0_37:3 | 0.81685067 |
| CountryMexico | -0.211 | 0.219 | -0.965 | 0.096 | Cer d18:0/24:1 | 0.819143037 |
| CountryMexico | -0.308 | 0.321 | -0.960 | 0.060 | DiCA 12:0 | 0.821138352 |
| CountryMexico | -0.160 | 0.168 | -0.957 | 0.058 | Cer d18:2/24:0 | 0.82201371 |
| CountryMexico | -0.109 | 0.115 | -0.953 | 0.055 | PG 22:4_22:6 | 0.822351061 |
| CountryMexico | -0.183 | 0.193 | -0.950 | 0.019 | TG 18:2_38:5 | 0.823101139 |
| CountryMexico | -0.202 | 0.213 | -0.949 | 0.029 | GAMT Deficiency | 0.824005995 |
| CountryMexico | -0.432 | 0.458 | -0.944 | 0.075 | 4-EPS | 0.826362516 |
| CountryMexico | -0.173 | 0.185 | -0.939 | 0.051 | CE 22:6 | 0.826794632 |
| CountryMexico | 0.109 | 0.116 | 0.942 | 0.021 | Orn Synthesis | 0.826794632 |
| CountryMexico | -0.059 | 0.063 | -0.941 | 0.061 | Ratio of PUFA-PIs to SFA-PIs | 0.826794632 |
| CountryMexico | -0.057 | 0.061 | -0.938 | 0.055 | Ratio of UFA-PIs to SFA-PIs | 0.82759926 |
| CountryMexico | -0.104 | 0.111 | -0.937 | 0.033 | Sum of UFA-LPGs | 0.828250934 |
| CountryMexico | 0.853 | 0.919 | 0.928 | 0.068 | SPHK activity (4) | 0.832348024 |
| CountryMexico | -0.853 | 0.919 | -0.928 | 0.068 | LPP3 activity (4) | 0.832348024 |
| CountryMexico | 0.201 | 0.218 | 0.926 | 0.040 | 4-Guanidinobutanoic acid | 0.833477325 |
| CountryMexico | -0.260 | 0.281 | -0.925 | 0.076 | TG 22:5_32:1 | 0.833893495 |
| CountryMexico | 0.201 | 0.218 | 0.920 | 0.098 | t4-OH-Pro | 0.835282342 |
| CountryMexico | -0.104 | 0.113 | -0.921 | 0.019 | Cer d18:1/24:0 | 0.835282342 |
| CountryMexico | -0.137 | 0.149 | -0.921 | 0.210 | Vanillylmandelic acid | 0.835282342 |
| CountryMexico | -0.281 | 0.308 | -0.913 | 0.035 | TG 16:0_32:3 | 0.836014266 |
| CountryMexico | -0.126 | 0.138 | -0.911 | 0.050 | Hex-Cer d18:2/23:0 | 0.836178985 |
| CountryMexico | -0.201 | 0.220 | -0.910 | 0.062 | LPS 20:1 | 0.836178985 |
| CountryMexico | 0.182 | 0.200 | 0.910 | 0.062 | Biliverdin | 0.836178985 |
| CountryMexico | 0.068 | 0.075 | 0.908 | 0.040 | SPBP d14:1 | 0.83670792 |
| CountryMexico | -0.515 | 0.569 | -0.905 | 0.022 | TG 18:0_30:1 | 0.839492953 |
| CountryMexico | 0.195 | 0.216 | 0.904 | 0.033 | TG 18:1_36:5 | 0.839568031 |
| CountryMexico | -0.136 | 0.153 | -0.891 | 0.032 | PI 18:1_20:5 | 0.842057457 |
| CountryMexico | 0.081 | 0.091 | 0.895 | 0.035 | PG 18:2_20:5 | 0.842057457 |
| CountryMexico | 0.254 | 0.285 | 0.892 | 0.050 | TMAO Synthesis (direct) | 0.842057457 |
| CountryMexico | 0.148 | 0.166 | 0.890 | 0.033 | IBD Deficiency (NBS) | 0.842057457 |
| CountryMexico | 0.182 | 0.205 | 0.886 | 0.104 | TG 18:1_34:3 | 0.84391978 |
| CountryMexico | -0.319 | 0.361 | -0.884 | 0.029 | ProBetaine | 0.8449897 |
| CountryMexico | 0.326 | 0.370 | 0.880 | 0.084 | DG 14:1_18:1 | 0.846270213 |
| CountryMexico | -0.104 | 0.118 | -0.882 | 0.076 | FA 24:4n-6 | 0.846270213 |
| CountryMexico | -0.360 | 0.412 | -0.874 | 0.031 | TG 18:0_32:2 | 0.847307863 |

| | | | | | | |
|---|---|---|---|---|---|---|
| CountryMexico | -0.098 | 0.112 | -0.873 | 0.031 | Sum of AAA-derived Lactic Acids | 0.847802961 |
| CountryMexico | 0.105 | 0.122 | 0.865 | 0.083 | PA 18:1_22:1 | 0.851141565 |
| CountryMexico | 0.140 | 0.162 | 0.861 | 0.041 | PA 18:1_18:4 | 0.852316061 |
| CountryMexico | -0.221 | 0.257 | -0.860 | 0.062 | TG 16:1_38:5 | 0.853538645 |
| CountryMexico | 0.158 | 0.185 | 0.853 | 0.047 | FA 4:0-2M | 0.854441147 |
| CountryMexico | -0.056 | 0.066 | -0.845 | 0.042 | FA 4:0 | 0.856858178 |
| CountryMexico | -0.181 | 0.215 | -0.842 | 0.034 | TG 18:0_36:3 | 0.856998891 |
| CountryMexico | 0.080 | 0.095 | 0.837 | 0.023 | 3-Met-2-oxovaleric acid | 0.858411328 |
| CountryMexico | 0.148 | 0.177 | 0.837 | 0.052 | LPI 22:0 | 0.858534902 |
| CountryMexico | 0.099 | 0.119 | 0.836 | 0.029 | HSer to Creatinine Ratio | 0.858981624 |
| CountryMexico | -0.474 | 0.569 | -0.834 | 0.030 | DG 16:1_18:0 | 0.859320896 |
| CountryMexico | -0.177 | 0.213 | -0.834 | 0.091 | LPS 18:2 | 0.859320896 |
| CountryMexico | 0.373 | 0.448 | 0.834 | 0.038 | Sum of Unconjugated Primary BAs | 0.859320896 |
| CountryMexico | -0.288 | 0.347 | -0.830 | 0.139 | IAG | 0.861344099 |
| CountryMexico | -0.166 | 0.200 | -0.829 | 0.068 | TG 16:0_40:7 | 0.861604978 |
| CountryMexico | -0.083 | 0.100 | -0.827 | 0.050 | PA 18:1_20:0 | 0.861979889 |
| CountryMexico | -0.347 | 0.420 | -0.826 | 0.015 | Primary BA Conjugation | 0.862018337 |
| CountryMexico | 0.060 | 0.073 | 0.826 | 0.024 | PC 36:2 | 0.862141302 |
| CountryMexico | -0.097 | 0.117 | -0.825 | 0.077 | Cer d18:1/23:0 | 0.862975487 |
| CountryMexico | -0.296 | 0.360 | -0.822 | 0.023 | TG 17:1_36:5 | 0.864654653 |
| CountryMexico | -0.242 | 0.297 | -0.816 | 0.154 | DHEAS | 0.867480579 |
| CountryMexico | -0.190 | 0.233 | -0.816 | 0.075 | PE 32:1 | 0.867480579 |
| CountryMexico | 0.125 | 0.153 | 0.816 | 0.252 | N-Ac-Putrescine | 0.867480579 |
| CountryMexico | -0.335 | 0.411 | -0.815 | 0.048 | TG 16:1_30:1 | 0.867896696 |
| CountryMexico | 0.109 | 0.135 | 0.813 | 0.036 | Symmetrical Arg Methylation | 0.869277197 |
| CountryMexico | 0.062 | 0.077 | 0.809 | 0.014 | PC 32:3 | 0.869565295 |
| CountryMexico | -0.169 | 0.209 | -0.810 | 0.045 | TG 18:2_33:2 | 0.869565295 |
| CountryMexico | -0.051 | 0.063 | -0.809 | 0.031 | Ratio of MUFA-PIs to SFA-PIs | 0.869565295 |
| CountryMexico | 0.188 | 0.234 | 0.806 | 0.104 | DG 16:1_18:2 | 0.870246055 |
| CountryMexico | 0.067 | 0.083 | 0.801 | 0.122 | PA 14:0_14:1 | 0.871772124 |
| CountryMexico | 0.093 | 0.116 | 0.801 | 0.042 | PA 16:0_18:2 | 0.871772124 |
| CountryMexico | 0.157 | 0.195 | 0.804 | 0.061 | PA 18:1_20:3 | 0.871772124 |
| CountryMexico | 0.079 | 0.098 | 0.800 | 0.012 | PG 18:0_22:1 | 0.871772124 |
| CountryMexico | -0.383 | 0.482 | -0.793 | 0.080 | TG 16:0_38:1 | 0.872661621 |
| CountryMexico | -0.201 | 0.252 | -0.797 | 0.041 | TG 18:0_34:3 | 0.872661621 |
| CountryMexico | 0.175 | 0.221 | 0.794 | 0.028 | TG 18:3_36:3 | 0.872661621 |
| CountryMexico | 0.087 | 0.110 | 0.792 | 0.042 | BVRA Activity | 0.872661621 |
| CountryMexico | -0.111 | 0.141 | -0.788 | 0.023 | PE P-18:1/18:1 | 0.873211377 |
| CountryMexico | 0.259 | 0.330 | 0.787 | 0.070 | TG 17:2_36:4 | 0.873211377 |
| CountryMexico | 0.189 | 0.240 | 0.786 | 0.032 | Creatine to Creatinine Ratio | 0.873516102 |
| CountryMexico | 0.104 | 0.133 | 0.784 | 0.017 | PC 32:2 | 0.874202189 |

| | | | | | | |
|---|---|---|---|---|---|---|
| CountryMexico | -0.249 | 0.318 | -0.783 | 0.023 | TG 14:0_38:4 | 0.874395202 |
| CountryMexico | 0.139 | 0.179 | 0.780 | 0.078 | PE P-18:0/18:2 | 0.875580329 |
| CountryMexico | -0.066 | 0.084 | -0.779 | 0.099 | PG 18:1_22:4 | 0.875580329 |
| CountryMexico | 0.093 | 0.119 | 0.777 | 0.017 | Uridine | 0.875702921 |
| CountryMexico | -0.151 | 0.195 | -0.777 | 0.018 | PI 18:1_22:4 | 0.875702921 |
| CountryMexico | 0.242 | 0.311 | 0.778 | 0.068 | 3-Met-His to 1-Met-His Ratio | 0.875702921 |
| CountryMexico | -0.123 | 0.158 | -0.774 | 0.163 | Cer d18:2/23:0 | 0.877461625 |
| CountryMexico | -0.166 | 0.215 | -0.772 | 0.041 | 5-Amino-4-oxovaleric acid | 0.877871344 |
| CountryMexico | 0.073 | 0.094 | 0.773 | 0.026 | PI 17:1_18:2 | 0.877871344 |
| CountryMexico | -0.443 | 0.578 | -0.767 | 0.047 | Cer d18:1/26:1 | 0.878863772 |
| CountryMexico | -0.124 | 0.161 | -0.768 | 0.098 | PE 38:6 | 0.878863772 |
| CountryMexico | -0.365 | 0.475 | -0.768 | 0.011 | TG 18:2_28:0 | 0.878863772 |
| CountryMexico | -0.330 | 0.428 | -0.770 | 0.014 | Gly Conjugation of Primary BAs | 0.878863772 |
| CountryMexico | -0.324 | 0.424 | -0.764 | 0.037 | TG 18:3_30:0 | 0.880010242 |
| CountryMexico | -0.090 | 0.118 | -0.762 | 0.037 | N-Ac-Glu | 0.88159885 |
| CountryMexico | 0.105 | 0.137 | 0.762 | 0.076 | PA 18:1_22:2 | 0.88159885 |
| CountryMexico | -0.089 | 0.118 | -0.759 | 0.021 | LPG 16:0 | 0.88159885 |
| CountryMexico | -0.105 | 0.138 | -0.760 | 0.050 | LPG 18:1 | 0.88159885 |
| CountryMexico | 0.074 | 0.098 | 0.759 | 0.075 | CPT-2 Deficiency (NBS) | 0.88159885 |
| CountryMexico | -0.150 | 0.199 | -0.753 | 0.054 | TG 16:0_38:5 | 0.885950818 |
| CountryMexico | 0.164 | 0.218 | 0.750 | 0.088 | TG 20:2_34:1 | 0.886103407 |
| CountryMexico | 0.112 | 0.150 | 0.748 | 0.030 | LPI 19:0 | 0.886235944 |
| CountryMexico | -0.158 | 0.212 | -0.746 | 0.138 | CE 18:0 | 0.886402257 |
| CountryMexico | -0.088 | 0.117 | -0.747 | 0.045 | CACT Deficiency (NBS) | 0.886402257 |
| CountryMexico | 0.088 | 0.117 | 0.747 | 0.045 | CPT-1 Deficiency (NBS) | 0.886402257 |
| CountryMexico | -0.337 | 0.454 | -0.743 | 0.026 | Taurine Conjugation of Primary BAs | 0.887834689 |
| CountryMexico | 0.150 | 0.202 | 0.742 | 0.015 | BAIBA | 0.888469436 |
| CountryMexico | -0.119 | 0.161 | -0.741 | 0.056 | N-Ac-Pro | 0.888469436 |
| CountryMexico | 0.197 | 0.266 | 0.741 | 0.035 | PA 18:2_22:4 | 0.888469436 |
| CountryMexico | -0.063 | 0.089 | -0.710 | 0.050 | FA 2:0 | 0.89033088 |
| CountryMexico | -0.293 | 0.407 | -0.719 | 0.045 | 2-OH-Benzoic acid | 0.89033088 |
| CountryMexico | 0.111 | 0.155 | 0.716 | 0.080 | PA 18:2_20:0 | 0.89033088 |
| CountryMexico | 0.126 | 0.177 | 0.711 | 0.058 | LPG 16:1 | 0.89033088 |
| CountryMexico | -0.074 | 0.104 | -0.711 | 0.090 | PC O-34:3 | 0.89033088 |
| CountryMexico | 0.053 | 0.075 | 0.716 | 0.050 | PG 18:2_22:1 | 0.89033088 |
| CountryMexico | -0.302 | 0.421 | -0.716 | 0.050 | TG 14:0_32:2 | 0.89033088 |
| CountryMexico | 0.304 | 0.414 | 0.734 | 0.057 | TG 20:1_32:0 | 0.89033088 |
| CountryMexico | -0.076 | 0.106 | -0.715 | 0.039 | Sum of Carboxylic Acids | 0.89033088 |
| CountryMexico | -0.071 | 0.100 | -0.707 | 0.041 | Sum of VLCFA-Cer | 0.89033088 |
| CountryMexico | 0.056 | 0.078 | 0.722 | 0.057 | Sum of LCFA-LPIs | 0.89033088 |
| CountryMexico | 0.315 | 0.442 | 0.713 | 0.047 | 7a-Dehydroxylation of CA | 0.89033088 |

| | | | | | | |
|---|---|---|---|---|---|---|
| CountryMexico | 0.080 | 0.113 | 0.708 | 0.033 | Urea to Creatinine Ratio | 0.89033088 |
| CountryMexico | -0.055 | 0.075 | -0.730 | 0.060 | FA 16:0 to FA 18:1 Ratio | 0.89033088 |
| CountryMexico | 0.100 | 0.139 | 0.720 | 0.011 | Cortisone Synthesis | 0.89033088 |
| CountryMexico | -0.112 | 0.155 | -0.722 | 0.066 | MC Deficiency (NBS) | 0.89033088 |
| CountryMexico | 0.112 | 0.155 | 0.722 | 0.066 | PA (NBS) | 0.89033088 |
| CountryMexico | 0.156 | 0.222 | 0.703 | 0.055 | C14:2-OH | 0.891499549 |
| CountryMexico | -0.147 | 0.212 | -0.695 | 0.033 | C5-OH (C3-DC-M) | 0.897008532 |
| CountryMexico | 0.128 | 0.185 | 0.694 | 0.066 | PA 16:2_18:1 | 0.89707141 |
| CountryMexico | -0.051 | 0.074 | -0.693 | 0.045 | LPE 22:0 | 0.897545831 |
| CountryMexico | 0.090 | 0.130 | 0.692 | 0.163 | LPI 14:0 | 0.897545831 |
| CountryMexico | 0.283 | 0.408 | 0.693 | 0.012 | p-Cresol-SO4 to p-Cresol Glucuronide | 0.897545831 |
| CountryMexico | -0.073 | 0.106 | -0.691 | 0.024 | PG 22:5_22:6 | 0.89771212 |
| CountryMexico | 0.135 | 0.197 | 0.686 | 0.024 | C16:1-OH | 0.898103103 |
| CountryMexico | 0.183 | 0.267 | 0.684 | 0.026 | DiCA 14:0 | 0.898103103 |
| CountryMexico | 0.074 | 0.108 | 0.684 | 0.155 | PI 18:2_22:0 | 0.898103103 |
| CountryMexico | -0.158 | 0.232 | -0.679 | 0.011 | C5-DC (C6-OH) | 0.898286684 |
| CountryMexico | -0.113 | 0.166 | -0.677 | 0.069 | LPA 14:1 | 0.898286684 |
| CountryMexico | -0.234 | 0.344 | -0.679 | 0.021 | TG 18:2_30:1 | 0.898286684 |
| CountryMexico | 0.127 | 0.188 | 0.674 | 0.051 | TG 18:1_36:4 | 0.898694799 |
| CountryMexico | 0.184 | 0.273 | 0.673 | 0.131 | TG 16:1_34:1 | 0.898824989 |
| CountryMexico | 0.112 | 0.166 | 0.673 | 0.053 | SCAD Deficiency (NBS) | 0.898824989 |
| CountryMexico | -0.156 | 0.234 | -0.665 | 0.019 | Cholesterol Synthesis | 0.902980367 |
| CountryMexico | -0.164 | 0.249 | -0.659 | 0.162 | Sum of Steroid Hormones | 0.904084918 |
| CountryMexico | -0.069 | 0.105 | -0.662 | 0.072 | FA 2:0 to Glucose Ratio | 0.904084918 |
| CountryMexico | -0.170 | 0.259 | -0.656 | 0.012 | C9 | 0.904940218 |
| CountryMexico | -0.160 | 0.244 | -0.656 | 0.061 | DG 18:2_20:4 | 0.904940218 |
| CountryMexico | 0.049 | 0.075 | 0.655 | 0.062 | PG 16:0_22:1 | 0.905123395 |
| CountryMexico | -0.057 | 0.087 | -0.648 | 0.076 | FA 24:1n-9 | 0.905904979 |
| CountryMexico | -0.261 | 0.406 | -0.642 | 0.033 | IsoLCA | 0.907380555 |
| CountryMexico | 0.140 | 0.217 | 0.644 | 0.082 | TG 18:1_34:4 | 0.907380555 |
| CountryMexico | -0.249 | 0.390 | -0.639 | 0.042 | LPE P-14:0 | 0.908928691 |
| CountryMexico | 0.146 | 0.228 | 0.639 | 0.077 | Pro Hydroxylation | 0.90900853 |
| CountryMexico | 0.316 | 0.495 | 0.638 | 0.016 | TG 18:3_34:0 | 0.909091752 |
| CountryMexico | -0.077 | 0.121 | -0.636 | 0.038 | Sum of MUFA-LPGs | 0.909778426 |
| CountryMexico | -0.181 | 0.286 | -0.634 | 0.118 | TG 20:3_32:1 | 0.909786066 |
| CountryMexico | -0.062 | 0.099 | -0.632 | 0.060 | PG 18:1_18:1 | 0.910503371 |
| CountryMexico | 0.083 | 0.131 | 0.632 | 0.046 | FA16:0 to FA 16:1 Ratio | 0.910503371 |
| CountryMexico | -0.084 | 0.134 | -0.628 | 0.013 | HCys | 0.911021528 |
| CountryMexico | -0.109 | 0.173 | -0.628 | 0.045 | IVA (NBS) | 0.911021528 |
| CountryMexico | -0.084 | 0.135 | -0.626 | 0.019 | SBCAD Deficiency (NBS) | 0.911388227 |
| CountryMexico | -0.145 | 0.232 | -0.625 | 0.085 | TG 17:0_34:1 | 0.911954172 |

| | | | | | | |
|---|---|---|---|---|---|---|
| CountryMexico | -0.043 | 0.068 | -0.624 | 0.175 | Glx to N-Ac-Asp Ratio | 0.911954172 |
| CountryMexico | -0.066 | 0.106 | -0.621 | 0.052 | Hex-Cer d18:1/20:0 | 0.912485066 |
| CountryMexico | 0.093 | 0.150 | 0.620 | 0.022 | PE P-18:0/16:0 | 0.912485066 |
| CountryMexico | 0.043 | 0.070 | 0.620 | 0.084 | Ratio of PUFA-PCs O to MUFA-PCs O | 0.912485066 |
| CountryMexico | 0.098 | 0.161 | 0.607 | 0.054 | AABA | 0.913408785 |
| CountryMexico | 0.042 | 0.069 | 0.606 | 0.023 | Val | 0.913408785 |
| CountryMexico | 0.064 | 0.104 | 0.616 | 0.069 | Cer d18:1/24:1 | 0.913408785 |
| CountryMexico | 0.120 | 0.199 | 0.604 | 0.039 | Hex-Cer d18:2/20:0 | 0.913408785 |
| CountryMexico | -0.080 | 0.131 | -0.610 | 0.020 | LPI 20:4 | 0.913408785 |
| CountryMexico | 0.096 | 0.157 | 0.613 | 0.023 | 3-OH-Isobutyric acid | 0.913408785 |
| CountryMexico | 0.090 | 0.146 | 0.614 | 0.036 | PA 18:2_18:2 | 0.913408785 |
| CountryMexico | -0.058 | 0.095 | -0.608 | 0.084 | PI 16:0_18:3 | 0.913408785 |
| CountryMexico | -0.073 | 0.118 | -0.616 | 0.047 | PI 18:2_20:0 | 0.913408785 |
| CountryMexico | -0.114 | 0.189 | -0.604 | 0.042 | TG 18:1_38:6 | 0.913408785 |
| CountryMexico | -0.152 | 0.247 | -0.615 | 0.031 | TG 18:2_32:2 | 0.913408785 |
| CountryMexico | -0.107 | 0.175 | -0.611 | 0.084 | TG 20:3_36:3 | 0.913408785 |
| CountryMexico | 0.046 | 0.076 | 0.603 | 0.055 | Sum of LPIs | 0.913408785 |
| CountryMexico | -0.118 | 0.194 | -0.606 | 0.011 | ACY1 Deficiency | 0.913408785 |
| CountryMexico | -0.075 | 0.123 | -0.608 | 0.060 | LDH Activity | 0.913408785 |
| CountryMexico | -0.315 | 0.527 | -0.597 | 0.042 | CMPF | 0.91401703 |
| CountryMexico | -0.083 | 0.138 | -0.598 | 0.023 | Phenyl-Lac | 0.91401703 |
| CountryMexico | -0.064 | 0.107 | -0.598 | 0.102 | PC 38:3 | 0.91401703 |
| CountryMexico | -0.250 | 0.416 | -0.600 | 0.076 | TG 20:3_36:5 | 0.91401703 |
| CountryMexico | -0.263 | 0.439 | -0.599 | 0.060 | Sum of 12a-OH BAs | 0.91401703 |
| CountryMexico | -0.057 | 0.097 | -0.593 | 0.061 | PI 18:2_22:6 | 0.914423252 |
| CountryMexico | -0.042 | 0.071 | -0.592 | 0.011 | Sum of PUFA-PIs | 0.914423252 |
| CountryMexico | -0.041 | 0.068 | -0.593 | 0.014 | Sum of UFA-PIs | 0.914423252 |
| CountryMexico | 0.251 | 0.425 | 0.590 | 0.019 | 12-KetoDCA to DCA Ratio | 0.914423252 |
| CountryMexico | 0.063 | 0.109 | 0.581 | 0.035 | C0 | 0.914922917 |
| CountryMexico | 0.088 | 0.152 | 0.579 | 0.048 | C2 | 0.914922917 |
| CountryMexico | -0.035 | 0.060 | -0.581 | 0.042 | ADMA | 0.914922917 |
| CountryMexico | -0.069 | 0.119 | -0.579 | 0.032 | Lac | 0.914922917 |
| CountryMexico | -0.064 | 0.110 | -0.584 | 0.101 | CerP d18:1/16:0 | 0.914922917 |
| CountryMexico | 0.219 | 0.376 | 0.583 | 0.072 | 3-OH-Sebacic acid | 0.914922917 |
| CountryMexico | 0.099 | 0.172 | 0.577 | 0.076 | LPI 18:2 | 0.914922917 |
| CountryMexico | 0.054 | 0.094 | 0.573 | 0.030 | PI 18:0_18:1 | 0.914922917 |
| CountryMexico | -0.055 | 0.094 | -0.584 | 0.032 | PI 18:2_20:5 | 0.914922917 |
| CountryMexico | -0.050 | 0.087 | -0.573 | 0.087 | PG 18:2_20:0 | 0.914922917 |
| CountryMexico | -0.115 | 0.199 | -0.577 | 0.066 | TG 17:0_34:2 | 0.914922917 |
| CountryMexico | 0.113 | 0.196 | 0.578 | 0.100 | TG 17:1_36:3 | 0.914922917 |
| CountryMexico | -0.178 | 0.311 | -0.572 | 0.020 | TG 18:2_30:0 | 0.914922917 |

| | | | | | | |
|---|---|---|---|---|---|---|
| CountryMexico | -0.105 | 0.184 | -0.571 | 0.076 | TG 18:2_35:2 | 0.914922917 |
| CountryMexico | 0.092 | 0.159 | 0.578 | 0.054 | Sum of Aminobutyric Acids | 0.914922917 |
| CountryMexico | 0.168 | 0.288 | 0.581 | 0.078 | Xanthine Synthesis | 0.914922917 |
| CountryMexico | 0.061 | 0.105 | 0.583 | 0.038 | MMA (NBS) | 0.914922917 |
| CountryMexico | 0.055 | 0.097 | 0.567 | 0.064 | Pro | 0.916510777 |
| CountryMexico | -0.038 | 0.066 | -0.565 | 0.049 | 4-OH-Phenylpyruvic acid | 0.916729216 |
| CountryMexico | -0.234 | 0.414 | -0.566 | 0.007 | TG 16:0_30:2 | 0.916729216 |
| CountryMexico | -0.297 | 0.532 | -0.559 | 0.057 | DCA | 0.917790309 |
| CountryMexico | 0.094 | 0.168 | 0.558 | 0.041 | PA 18:1_22:3 | 0.917790309 |
| CountryMexico | 0.083 | 0.150 | 0.558 | 0.029 | PG 18:0_18:2 | 0.917790309 |
| CountryMexico | -0.122 | 0.218 | -0.558 | 0.091 | TG 16:0_35:3 | 0.917790309 |
| CountryMexico | 0.143 | 0.257 | 0.557 | 0.067 | TG 18:3_35:2 | 0.917790309 |
| CountryMexico | 0.111 | 0.200 | 0.554 | 0.013 | FA 18:3 | 0.918913652 |
| CountryMexico | 0.053 | 0.097 | 0.549 | 0.063 | PA 18:2_20:1 | 0.91933565 |
| CountryMexico | 0.033 | 0.061 | 0.552 | 0.030 | Sum of SFA-LPIs | 0.91933565 |
| CountryMexico | -0.085 | 0.156 | -0.542 | 0.051 | Cer d18:2/22:0 | 0.9204228 |
| CountryMexico | 0.054 | 0.100 | 0.545 | 0.017 | PG 18:2_22:4 | 0.9204228 |
| CountryMexico | 0.076 | 0.141 | 0.543 | 0.027 | PG 22:6_22:6 | 0.9204228 |
| CountryMexico | 0.133 | 0.244 | 0.545 | 0.054 | TG 18:1_32:0 | 0.9204228 |
| CountryMexico | 0.043 | 0.080 | 0.536 | 0.043 | Muscle Protein Degradation | 0.921613943 |
| CountryMexico | -0.071 | 0.133 | -0.532 | 0.022 | N1-Ac-Spermidine | 0.922372951 |
| CountryMexico | 0.082 | 0.154 | 0.531 | 0.063 | Sum of PUFA-LPIs | 0.922782695 |
| CountryMexico | 0.031 | 0.059 | 0.530 | 0.054 | Valinemia (NBS) | 0.922999129 |
| CountryMexico | 0.157 | 0.297 | 0.529 | 0.101 | TG 14:0_36:2 | 0.92332277 |
| CountryMexico | 0.044 | 0.084 | 0.523 | 0.049 | PI 18:0_18:0 | 0.92450393 |
| CountryMexico | -0.149 | 0.285 | -0.521 | 0.062 | TG 16:0_33:1 | 0.92450393 |
| CountryMexico | -0.151 | 0.290 | -0.521 | 0.045 | TG 18:2_33:0 | 0.92450393 |
| CountryMexico | -0.104 | 0.200 | -0.520 | 0.088 | TG 18:2_33:1 | 0.92450393 |
| CountryMexico | -0.156 | 0.302 | -0.518 | 0.019 | TG 16:0_40:8 | 0.924595921 |
| CountryMexico | 0.108 | 0.210 | 0.518 | 0.007 | BAIBA Synthesis | 0.924595921 |
| CountryMexico | 0.164 | 0.319 | 0.513 | 0.058 | DG 18:1_18:3 | 0.925185866 |
| CountryMexico | -0.184 | 0.359 | -0.512 | 0.074 | TG 14:0_38:5 | 0.925185866 |
| CountryMexico | 0.130 | 0.254 | 0.513 | 0.065 | TG 16:0_34:1 | 0.925185866 |
| CountryMexico | -0.106 | 0.209 | -0.508 | 0.043 | DG 18:1_20:4 | 0.926862898 |
| CountryMexico | -0.034 | 0.067 | -0.509 | 0.024 | Sum of MUFA-PIs | 0.926862898 |
| CountryMexico | 0.170 | 0.335 | 0.507 | 0.063 | TG 17:2_34:2 | 0.928019016 |
| CountryMexico | 0.250 | 0.499 | 0.501 | 0.043 | 3-IPA | 0.929951194 |
| CountryMexico | -0.112 | 0.224 | -0.499 | 0.033 | PE P-18:0/14:0 | 0.929951194 |
| CountryMexico | -0.117 | 0.233 | -0.502 | 0.021 | PE P-18:0/20:2 | 0.929951194 |
| CountryMexico | -0.275 | 0.548 | -0.501 | 0.004 | TG 16:0_28:1 | 0.929951194 |
| CountryMexico | -0.062 | 0.125 | -0.499 | 0.084 | Gly to Ala Ratio | 0.929951194 |

| | | | | | | |
|---|---|---|---|---|---|---|
| CountryMexico | 0.064 | 0.128 | 0.498 | 0.051 | PA 18:1_20:2 | 0.930429319 |
| CountryMexico | -0.033 | 0.066 | -0.497 | 0.016 | Sum of PIs | 0.930575858 |
| CountryMexico | -0.205 | 0.415 | -0.494 | 0.072 | GLCAS | 0.931229276 |
| CountryMexico | -0.161 | 0.326 | -0.494 | 0.036 | TG 16:0_32:2 | 0.931229276 |
| CountryMexico | 0.121 | 0.247 | 0.492 | 0.038 | TG 17:1_36:4 | 0.931229276 |
| CountryMexico | -0.128 | 0.260 | -0.494 | 0.029 | TG 20:1_32:3 | 0.931229276 |
| CountryMexico | 0.104 | 0.212 | 0.492 | 0.091 | TG 20:2_34:2 | 0.931229276 |
| CountryMexico | -0.219 | 0.442 | -0.495 | 0.019 | Gly Conjugation of DCA | 0.931229276 |
| CountryMexico | -0.231 | 0.472 | -0.489 | 0.052 | Taurine Conjugation of DCA | 0.932254227 |
| CountryMexico | 0.105 | 0.215 | 0.488 | 0.053 | DG 18:2_18:2 | 0.933412361 |
| CountryMexico | -0.200 | 0.411 | -0.486 | 0.080 | LPG 14:1 | 0.933935607 |
| CountryMexico | -0.152 | 0.315 | -0.483 | 0.044 | CE 22:2 | 0.934202878 |
| CountryMexico | -0.157 | 0.324 | -0.483 | 0.057 | TG 14:0_34:3 | 0.934202878 |
| CountryMexico | -0.180 | 0.378 | -0.476 | 0.004 | TG 18:0_38:7 | 0.934202878 |
| CountryMexico | -0.100 | 0.207 | -0.484 | 0.042 | Hex3Cer d18:1/24:1 | 0.934202878 |
| CountryMexico | -0.031 | 0.065 | -0.477 | 0.016 | Sum of (L)PIs | 0.934202878 |
| CountryMexico | 0.045 | 0.094 | 0.483 | 0.049 | Sum of UFA-LPIs | 0.934202878 |
| CountryMexico | -0.034 | 0.071 | -0.480 | 0.111 | Ratio of PUFA-LPCs to SFA-LPCs | 0.934202878 |
| CountryMexico | -0.127 | 0.270 | -0.471 | 0.031 | TMAO Synthesis | 0.936181229 |
| CountryMexico | 0.043 | 0.091 | 0.471 | 0.099 | Sum of MUFA-PAs | 0.93628788 |
| CountryMexico | 0.133 | 0.283 | 0.470 | 0.038 | Cer d18:1/26:0 | 0.936700062 |
| CountryMexico | -0.035 | 0.075 | -0.468 | 0.020 | PI 16:1_18:2 | 0.936700062 |
| CountryMexico | 0.126 | 0.270 | 0.467 | 0.113 | TG 18:1_32:1 | 0.936700062 |
| CountryMexico | 0.152 | 0.325 | 0.467 | 0.064 | TG 20:2_32:0 | 0.936700062 |
| CountryMexico | 0.053 | 0.113 | 0.464 | 0.057 | PA 18:1_18:2 | 0.937503096 |
| CountryMexico | 0.053 | 0.115 | 0.464 | 0.102 | PI 18:1_20:0 | 0.937543055 |
| CountryMexico | -0.281 | 0.607 | -0.464 | 0.053 | SPBP d14:0 | 0.937543055 |
| CountryMexico | -0.118 | 0.257 | -0.460 | 0.015 | TG 14:0_36:4 | 0.939125854 |
| CountryMexico | -0.223 | 0.488 | -0.456 | 0.030 | TG 16:0_28:2 | 0.940600234 |
| CountryMexico | -0.077 | 0.170 | -0.453 | 0.071 | PE 33:2 | 0.941316941 |
| CountryMexico | 0.189 | 0.420 | 0.450 | 0.059 | TrpBetaine | 0.942567747 |
| CountryMexico | -0.227 | 0.507 | -0.447 | 0.042 | TG 18:0_30:0 | 0.943367741 |
| CountryMexico | -0.083 | 0.185 | -0.446 | 0.034 | SM 38:3 | 0.943409458 |
| CountryMexico | -0.118 | 0.266 | -0.444 | 0.119 | MG 20:1 | 0.944129441 |
| CountryMexico | 0.095 | 0.218 | 0.437 | 0.045 | 5-AVA | 0.944324442 |
| CountryMexico | -0.102 | 0.233 | -0.437 | 0.055 | PE 28:0 | 0.944324442 |
| CountryMexico | 0.059 | 0.133 | 0.441 | 0.055 | PI 18:1_20:3 | 0.944324442 |
| CountryMexico | 0.070 | 0.160 | 0.438 | 0.026 | PG 16:0_16:0 | 0.944324442 |
| CountryMexico | -0.032 | 0.074 | -0.440 | 0.029 | PG 18:1_18:2 | 0.944324442 |
| CountryMexico | -0.132 | 0.301 | -0.440 | 0.007 | TG 14:0_40:5 | 0.944324442 |
| CountryMexico | 0.093 | 0.209 | 0.443 | 0.080 | TG 18:1_35:3 | 0.944324442 |

| | | | | | | |
|---|---|---|---|---|---|---|
| CountryMexico | 0.078 | 0.176 | 0.440 | 0.098 | Ratio of HArg to SDMA | 0.944324442 |
| CountryMexico | 0.062 | 0.142 | 0.440 | 0.067 | Ratio of PUFA-LPIs to MUFA-LPIs | 0.944324442 |
| CountryMexico | 0.056 | 0.130 | 0.433 | 0.042 | PG 16:2_18:2 | 0.94592227 |
| CountryMexico | 0.108 | 0.248 | 0.434 | 0.036 | TG 22:5_32:0 | 0.94592227 |
| CountryMexico | -0.110 | 0.257 | -0.430 | 0.053 | TG 16:1_36:5 | 0.946050549 |
| CountryMexico | -0.111 | 0.259 | -0.429 | 0.033 | TG 18:0_36:2 | 0.946050549 |
| CountryMexico | 0.088 | 0.205 | 0.429 | 0.039 | TG 18:3_34:1 | 0.946050549 |
| CountryMexico | 0.086 | 0.202 | 0.425 | 0.037 | DG 16:0_18:2 | 0.946902937 |
| CountryMexico | -0.064 | 0.152 | -0.422 | 0.065 | HArg | 0.947227209 |
| CountryMexico | -0.072 | 0.171 | -0.423 | 0.005 | FA 20:3n-6 | 0.947227209 |
| CountryMexico | -0.032 | 0.076 | -0.421 | 0.049 | PI 15:1_16:0 | 0.947555697 |
| CountryMexico | -0.046 | 0.111 | -0.418 | 0.007 | Cer d18:2/16:0 | 0.948379179 |
| CountryMexico | 0.045 | 0.108 | 0.416 | 0.031 | DLD (NBS) | 0.948604678 |
| CountryMexico | -0.275 | 0.663 | -0.415 | 0.049 | SPHK activity (6) | 0.948660588 |
| CountryMexico | -0.106 | 0.257 | -0.414 | 0.063 | TG 18:1_31:0 | 0.948738381 |
| CountryMexico | 0.106 | 0.256 | 0.413 | 0.128 | Phenylglucuronide | 0.949979614 |
| CountryMexico | 0.033 | 0.079 | 0.411 | 0.064 | PI 15:0_16:0 | 0.950306327 |
| CountryMexico | -0.152 | 0.370 | -0.410 | 0.008 | TG 20:1_32:2 | 0.950306327 |
| CountryMexico | -0.077 | 0.190 | -0.405 | 0.012 | N-Ac-Trp | 0.950395586 |
| CountryMexico | -0.030 | 0.073 | -0.409 | 0.077 | PG 16:0_18:2 | 0.950395586 |
| CountryMexico | 0.077 | 0.196 | 0.396 | 0.090 | TG 22:5_34:1 | 0.951141183 |
| CountryMexico | -0.134 | 0.338 | -0.396 | 0.056 | TG 20:3_34:0 | 0.951151408 |
| CountryMexico | 0.076 | 0.194 | 0.393 | 0.043 | TG 18:2_36:3 | 0.951771697 |
| CountryMexico | 0.045 | 0.114 | 0.391 | 0.027 | PG 16:1_18:2 | 0.952790918 |
| CountryMexico | 0.035 | 0.090 | 0.391 | 0.026 | Sum of Sulfur-Containing AAs | 0.952790918 |
| CountryMexico | -0.037 | 0.096 | -0.389 | 0.051 | PS 36:5 | 0.952862176 |
| CountryMexico | -0.094 | 0.244 | -0.386 | 0.095 | TG 16:0_35:2 | 0.952862176 |
| CountryMexico | 0.125 | 0.324 | 0.386 | 0.038 | Taurine Conjugation of UDCA | 0.952862176 |
| CountryMexico | -0.082 | 0.213 | -0.384 | 0.046 | FA 16:1 | 0.953194155 |
| CountryMexico | -0.137 | 0.358 | -0.382 | 0.012 | TG 18:1_30:2 | 0.953506789 |
| CountryMexico | 0.084 | 0.223 | 0.378 | 0.149 | TG 18:1_33:1 | 0.954167936 |
| CountryMexico | -0.065 | 0.173 | -0.373 | 0.018 | FA 20:2 | 0.954169483 |
| CountryMexico | -0.042 | 0.112 | -0.376 | 0.060 | PG 18:1_22:1 | 0.954169483 |
| CountryMexico | 0.039 | 0.105 | 0.368 | 0.013 | PG 18:2_22:3 | 0.954169483 |
| CountryMexico | -0.079 | 0.212 | -0.373 | 0.111 | TG 17:1_34:2 | 0.954169483 |
| CountryMexico | -0.095 | 0.254 | -0.372 | 0.021 | TG 18:3_32:0 | 0.954169483 |
| CountryMexico | 0.115 | 0.309 | 0.372 | 0.140 | TG 20:1_32:1 | 0.954169483 |
| CountryMexico | 0.063 | 0.175 | 0.360 | 0.022 | PE 34:3 | 0.956073178 |
| CountryMexico | -0.065 | 0.179 | -0.361 | 0.091 | TG 17:0_36:3 | 0.956073178 |
| CountryMexico | 0.093 | 0.262 | 0.355 | 0.095 | TG 16:1_34:3 | 0.956770865 |
| CountryMexico | 0.149 | 0.424 | 0.352 | 0.020 | TG 17:2_36:2 | 0.956860885 |

| | | | | | | |
|---|---|---|---|---|---|---|
| CountryMexico | -0.075 | 0.214 | -0.352 | 0.133 | TG 20:3_34:1 | 0.956860885 |
| CountryMexico | 0.086 | 0.245 | 0.353 | 0.006 | CMAMMA | 0.956860885 |
| CountryMexico | 0.027 | 0.080 | 0.338 | 0.006 | a-Ketoisovaleric acid | 0.959319434 |
| CountryMexico | 0.054 | 0.162 | 0.333 | 0.018 | LPI 17:0 | 0.959319434 |
| CountryMexico | -0.064 | 0.191 | -0.337 | 0.006 | PE 36:3 | 0.959319434 |
| CountryMexico | 0.029 | 0.084 | 0.341 | 0.063 | PI 14:0_18:2 | 0.959319434 |
| CountryMexico | 0.058 | 0.170 | 0.339 | 0.066 | PC 32:1 | 0.959319434 |
| CountryMexico | -0.109 | 0.326 | -0.336 | 0.085 | TG 16:1_32:2 | 0.959319434 |
| CountryMexico | 0.076 | 0.225 | 0.339 | 0.065 | TG 16:1_36:4 | 0.959319434 |
| CountryMexico | -0.093 | 0.277 | -0.334 | 0.064 | TG 18:3_32:1 | 0.959319434 |
| CountryMexico | 0.082 | 0.245 | 0.332 | 0.073 | TG 22:6_34:1 | 0.959319434 |
| CountryMexico | -0.086 | 0.254 | -0.338 | 0.045 | TG 22:6_34:2 | 0.959319434 |
| CountryMexico | 0.023 | 0.068 | 0.335 | 0.018 | Sum of BCAAs | 0.959319434 |
| CountryMexico | -0.065 | 0.196 | -0.330 | 0.004 | Cer d18:2/20:0 | 0.959403841 |
| CountryMexico | 0.056 | 0.172 | 0.324 | 0.049 | FA 18:1 | 0.960574279 |
| CountryMexico | 0.067 | 0.208 | 0.321 | 0.117 | PE P-16:0/18:3 | 0.960877313 |
| CountryMexico | 0.049 | 0.152 | 0.319 | 0.038 | Ratio of PUFA-LPIs to SFA-LPIs | 0.960877313 |
| CountryMexico | 0.050 | 0.167 | 0.303 | 0.088 | PE P-18:0/20:3 | 0.965204506 |
| CountryMexico | 0.064 | 0.215 | 0.296 | 0.052 | MG 22:2 | 0.967943463 |
| CountryMexico | -0.077 | 0.258 | -0.297 | 0.030 | TG 22:5_34:3 | 0.967943463 |
| CountryMexico | -0.080 | 0.271 | -0.297 | 0.072 | IsoUDCA to UDCA Ratio | 0.967943463 |
| CountryMexico | 0.018 | 0.062 | 0.289 | 0.077 | Ser | 0.968500369 |
| CountryMexico | 0.057 | 0.194 | 0.293 | 0.126 | TG 18:1_33:2 | 0.968500369 |
| CountryMexico | 0.019 | 0.068 | 0.287 | 0.053 | Sum of MUFA-LPIs | 0.968500369 |
| CountryMexico | 0.069 | 0.240 | 0.288 | 0.075 | Kynurenic Acid to Xanthurenic Acid Ratio | 0.968500369 |
| CountryMexico | 0.028 | 0.101 | 0.274 | 0.027 | Cys | 0.969297514 |
| CountryMexico | 0.046 | 0.165 | 0.280 | 0.117 | LPA 15:0 | 0.969297514 |
| CountryMexico | 0.027 | 0.097 | 0.276 | 0.094 | PA 18:1_20:1 | 0.969297514 |
| CountryMexico | -0.047 | 0.172 | -0.274 | 0.010 | PA 18:2_22:3 | 0.969297514 |
| CountryMexico | 0.038 | 0.138 | 0.277 | 0.011 | PI 18:2_20:1 | 0.969297514 |
| CountryMexico | 0.044 | 0.161 | 0.276 | 0.021 | Isonicotinic acid | 0.969297514 |
| CountryMexico | 0.102 | 0.362 | 0.282 | 0.106 | TG 16:1_38:3 | 0.969297514 |
| CountryMexico | 0.081 | 0.286 | 0.284 | 0.002 | TG 17:2_36:3 | 0.969297514 |
| CountryMexico | 0.066 | 0.239 | 0.275 | 0.014 | TG 18:3_34:3 | 0.969297514 |
| CountryMexico | -0.015 | 0.055 | -0.276 | 0.020 | Sum of Essential AAs | 0.969297514 |
| CountryMexico | 0.101 | 0.369 | 0.274 | 0.077 | Ketone Body Synthesis | 0.969297514 |
| CountryMexico | 0.057 | 0.208 | 0.273 | 0.027 | NAT10 Activity | 0.969297514 |
| CountryMexico | -0.028 | 0.102 | -0.273 | 0.073 | PI 18:1_22:5 | 0.969350525 |
| CountryMexico | -0.032 | 0.117 | -0.271 | 0.031 | Sum of FA 12:0 - FA 16:0 | 0.969533946 |
| CountryMexico | 0.022 | 0.081 | 0.270 | 0.022 | SPBP d17:1 | 0.969780456 |
| CountryMexico | 0.063 | 0.234 | 0.268 | 0.034 | PA 16:0_18:3 | 0.969791157 |

| | | | | | | |
|---|---|---|---|---|---|---|
| CountryMexico | -0.084 | 0.315 | -0.266 | 0.047 | TG 22:6_32:0 | 0.969791157 |
| CountryMexico | -0.096 | 0.362 | -0.265 | 0.044 | DG 16:0_16:1 | 0.970486873 |
| CountryMexico | 0.048 | 0.180 | 0.265 | 0.029 | FA 18:2 | 0.970486873 |
| CountryMexico | 0.024 | 0.092 | 0.265 | 0.058 | PG 18:2_18:2 | 0.970486873 |
| CountryMexico | 0.032 | 0.123 | 0.263 | 0.014 | LPI 16:1 | 0.970542986 |
| CountryMexico | 0.038 | 0.145 | 0.263 | 0.040 | N-Ac-Cytidine | 0.970542986 |
| CountryMexico | -0.026 | 0.100 | -0.262 | 0.014 | PI 18:1_18:3 | 0.970542986 |
| CountryMexico | 0.025 | 0.095 | 0.262 | 0.042 | PG 18:2_18:3 | 0.970542986 |
| CountryMexico | 0.060 | 0.226 | 0.264 | 0.031 | TG 18:2_34:3 | 0.970542986 |
| CountryMexico | 0.058 | 0.223 | 0.261 | 0.052 | TG 18:2_35:3 | 0.970640252 |
| CountryMexico | 0.030 | 0.115 | 0.261 | 0.065 | PA 18:2_22:0 | 0.970822832 |
| CountryMexico | 0.020 | 0.077 | 0.256 | 0.020 | PG 18:0_18:3 | 0.971019688 |
| CountryMexico | 0.041 | 0.160 | 0.255 | 0.027 | PG 17:0_18:2 | 0.971049176 |
| CountryMexico | -0.014 | 0.055 | -0.254 | 0.058 | Ratio of MUFA-LPIs to SFA-LPIs | 0.971120038 |
| CountryMexico | -0.058 | 0.229 | -0.254 | 0.005 | TG 16:0_36:5 | 0.97117192 |
| CountryMexico | -0.066 | 0.264 | -0.250 | 0.026 | C5:1 | 0.972286142 |
| CountryMexico | 0.127 | 0.540 | 0.236 | 0.056 | GCA | 0.972286142 |
| CountryMexico | -0.058 | 0.231 | -0.250 | 0.010 | LPI 17:1 | 0.972286142 |
| CountryMexico | 0.045 | 0.181 | 0.249 | 0.026 | PE 34:2 | 0.972286142 |
| CountryMexico | -0.089 | 0.367 | -0.244 | 0.009 | PS 32:0 | 0.972286142 |
| CountryMexico | -0.023 | 0.097 | -0.237 | 0.053 | PC O-34:2 | 0.972286142 |
| CountryMexico | -0.045 | 0.195 | -0.231 | 0.038 | PG 14:0_16:0 | 0.972286142 |
| CountryMexico | -0.028 | 0.114 | -0.241 | 0.095 | PG 18:1_22:0 | 0.972286142 |
| CountryMexico | 0.057 | 0.232 | 0.245 | 0.064 | TG 16:0_34:2 | 0.972286142 |
| CountryMexico | -0.062 | 0.251 | -0.246 | 0.067 | TG 17:0_34:3 | 0.972286142 |
| CountryMexico | -0.060 | 0.255 | -0.237 | 0.105 | TG 17:1_34:1 | 0.972286142 |
| CountryMexico | -0.022 | 0.095 | -0.231 | 0.057 | Sum BCFA | 0.972286142 |
| CountryMexico | 0.040 | 0.173 | 0.232 | 0.048 | Sum of MUFAs | 0.972286142 |
| CountryMexico | 0.017 | 0.073 | 0.235 | 0.056 | Sum of SFA-PIs | 0.972286142 |
| CountryMexico | -0.035 | 0.141 | -0.250 | 0.030 | Sum of Spha | 0.972286142 |
| CountryMexico | 0.041 | 0.181 | 0.228 | 0.032 | LPG 20:1 | 0.97264658 |
| CountryMexico | 0.059 | 0.260 | 0.226 | 0.042 | TG 16:0_34:4 | 0.97264658 |
| CountryMexico | -0.008 | 0.036 | -0.228 | 0.021 | Ratio of PUFA-PIs to MUFA-PIs | 0.97264658 |
| CountryMexico | -0.078 | 0.350 | -0.221 | 0.099 | DG 18:1_20:2 | 0.973245899 |
| CountryMexico | 0.007 | 0.030 | 0.222 | 0.036 | Indoleacetaldehyde | 0.973245899 |
| CountryMexico | 0.041 | 0.185 | 0.221 | 0.026 | PS 34:2 | 0.973245899 |
| CountryMexico | -0.086 | 0.395 | -0.219 | 0.008 | Gly Conjugation of UDCA | 0.974082151 |
| CountryMexico | -0.033 | 0.151 | -0.218 | 0.060 | AMSDH Saturation | 0.974199342 |
| CountryMexico | 0.082 | 0.379 | 0.215 | 0.021 | C3-DC (C4-OH) | 0.974241494 |
| CountryMexico | -0.044 | 0.202 | -0.216 | 0.020 | Creatine | 0.974241494 |
| CountryMexico | -0.027 | 0.126 | -0.217 | 0.098 | PI 18:0_20:0 | 0.974241494 |

| | | | | | | |
|---|---|---|---|---|---|---|
| CountryMexico | 0.021 | 0.097 | 0.215 | 0.084 | PG 18:1_20:4 | 0.974241494 |
| CountryMexico | -0.015 | 0.068 | -0.216 | 0.032 | BCAT Val Catabolism | 0.974241494 |
| CountryMexico | -0.036 | 0.167 | -0.214 | 0.046 | CE 22:5 | 0.974380169 |
| CountryMexico | -0.024 | 0.115 | -0.210 | 0.029 | PI 18:1_22:0 | 0.975050491 |
| CountryMexico | -0.020 | 0.096 | -0.209 | 0.004 | Picolinic acid | 0.975050491 |
| CountryMexico | 0.054 | 0.256 | 0.211 | 0.047 | TG 14:0_36:3 | 0.975050491 |
| CountryMexico | 0.037 | 0.174 | 0.210 | 0.016 | MA (NBS) | 0.975050491 |
| CountryMexico | 0.016 | 0.075 | 0.206 | 0.083 | PC 34:1 | 0.975414812 |
| CountryMexico | 0.024 | 0.115 | 0.205 | 0.028 | PI 16:0_16:0 | 0.975417945 |
| CountryMexico | 0.096 | 0.469 | 0.205 | 0.052 | GDCA Synthesis from CA | 0.975417945 |
| CountryMexico | -0.043 | 0.218 | -0.198 | 0.023 | TG 18:2_36:4 | 0.977246508 |
| CountryMexico | -0.078 | 0.394 | -0.198 | 0.056 | TG 22:6_34:3 | 0.977246508 |
| CountryMexico | -0.055 | 0.280 | -0.195 | 0.033 | GLCA | 0.977553454 |
| CountryMexico | -0.082 | 0.421 | -0.196 | 0.016 | TG 18:1_28:1 | 0.977553454 |
| CountryMexico | 0.049 | 0.249 | 0.196 | 0.051 | TG 18:1_33:3 | 0.977553454 |
| CountryMexico | 0.023 | 0.121 | 0.192 | 0.013 | Glutaminolysis Rate | 0.978070109 |
| CountryMexico | -0.076 | 0.417 | -0.181 | 0.015 | LPG 17:1 | 0.978081461 |
| CountryMexico | -0.027 | 0.141 | -0.192 | 0.035 | PE P-18:1/22:6 | 0.978081461 |
| CountryMexico | 0.022 | 0.121 | 0.179 | 0.073 | PI 18:1_22:2 | 0.978081461 |
| CountryMexico | -0.025 | 0.133 | -0.186 | 0.016 | PC 38:6 | 0.978081461 |
| CountryMexico | -0.025 | 0.136 | -0.185 | 0.074 | PC 40:6 | 0.978081461 |
| CountryMexico | -0.025 | 0.135 | -0.185 | 0.070 | PG 18:0_18:1 | 0.978081461 |
| CountryMexico | -0.055 | 0.299 | -0.184 | 0.112 | TG 16:1_33:1 | 0.978081461 |
| CountryMexico | -0.067 | 0.350 | -0.191 | 0.043 | TG 18:1_30:0 | 0.978081461 |
| CountryMexico | -0.046 | 0.247 | -0.185 | 0.034 | TG 18:1_32:3 | 0.978081461 |
| CountryMexico | -0.030 | 0.162 | -0.183 | 0.091 | Ratio of HArg to ADMA | 0.978081461 |
| CountryMexico | 0.084 | 0.457 | 0.184 | 0.094 | TDCA Synthesis from CA | 0.978081461 |
| CountryMexico | 0.025 | 0.137 | 0.180 | 0.041 | Ratio of Acetylcarnitine to Carnitine | 0.978081461 |
| CountryMexico | 0.019 | 0.106 | 0.177 | 0.121 | Kynurenine | 0.978946317 |
| CountryMexico | 0.046 | 0.258 | 0.177 | 0.072 | TG 18:1_32:2 | 0.978946317 |
| CountryMexico | 0.043 | 0.244 | 0.175 | 0.010 | TG 18:2_36:5 | 0.979042918 |
| CountryMexico | 0.045 | 0.261 | 0.173 | 0.014 | TG 18:3_36:1 | 0.979042918 |
| CountryMexico | 0.023 | 0.131 | 0.176 | 0.042 | b-Oxidation | 0.979042918 |
| CountryMexico | 0.011 | 0.062 | 0.176 | 0.054 | Ratio of UFA-LPCs to SFA-LPCs | 0.979042918 |
| CountryMexico | 0.010 | 0.059 | 0.172 | 0.060 | Phe | 0.979544567 |
| CountryMexico | -0.017 | 0.104 | -0.168 | 0.065 | Choline to Creatinine Ratio | 0.981505603 |
| CountryMexico | 0.034 | 0.203 | 0.167 | 0.034 | TG 18:2_34:2 | 0.981630677 |
| CountryMexico | 0.042 | 0.256 | 0.165 | 0.095 | TG 16:1_34:2 | 0.982740199 |
| CountryMexico | -0.034 | 0.207 | -0.164 | 0.033 | TG 18:2_38:4 | 0.982881418 |
| CountryMexico | 0.032 | 0.199 | 0.163 | 0.038 | 2-OH-Butyric acid | 0.982939275 |
| CountryMexico | -0.035 | 0.220 | -0.161 | 0.017 | TG 18:3_34:2 | 0.983666899 |

| | | | | | | |
|---|---|---|---|---|---|---|
| CountryMexico | 0.083 | 0.521 | 0.160 | 0.029 | Indole Pathway Activity | 0.983666899 |
| CountryMexico | -0.021 | 0.143 | -0.147 | 0.017 | C5 | 0.984282992 |
| CountryMexico | 0.052 | 0.357 | 0.147 | 0.065 | 3-Met-His | 0.984282992 |
| CountryMexico | 0.021 | 0.137 | 0.151 | 0.090 | PA 18:1_22:0 | 0.984282992 |
| CountryMexico | 0.027 | 0.177 | 0.151 | 0.047 | PG 16:0_18:1 | 0.984282992 |
| CountryMexico | 0.040 | 0.255 | 0.157 | 0.040 | TG 20:2_32:1 | 0.984282992 |
| CountryMexico | -0.074 | 0.495 | -0.149 | 0.013 | Sum of Gly-Conjugated BAs | 0.984282992 |
| CountryMexico | 0.013 | 0.087 | 0.144 | 0.051 | PI 16:0_22:1 | 0.985310026 |
| CountryMexico | -0.046 | 0.334 | -0.139 | 0.081 | 12-KetoDCA | 0.985627802 |
| CountryMexico | 0.026 | 0.187 | 0.137 | 0.043 | PI 18:1_22:3 | 0.985627802 |
| CountryMexico | -0.044 | 0.322 | -0.137 | 0.035 | TG 14:0_34:2 | 0.985627802 |
| CountryMexico | -0.041 | 0.303 | -0.137 | 0.016 | TG 16:0_32:0 | 0.985627802 |
| CountryMexico | -0.054 | 0.381 | -0.141 | 0.038 | TG 18:1_30:1 | 0.985627802 |
| CountryMexico | 0.025 | 0.186 | 0.137 | 0.148 | TG 18:1_35:2 | 0.985627802 |
| CountryMexico | 0.034 | 0.244 | 0.140 | 0.029 | TG 18:2_34:4 | 0.985627802 |
| CountryMexico | 0.066 | 0.483 | 0.136 | 0.017 | Sum of Primary BAs | 0.985627802 |
| CountryMexico | 0.012 | 0.089 | 0.136 | 0.016 | Ratio of UFA-LPIs to SFA-LPIs | 0.985627802 |
| CountryMexico | 0.045 | 0.356 | 0.127 | 0.083 | 3-OH-Butyric acid | 0.986052128 |
| CountryMexico | -0.064 | 0.513 | -0.125 | 0.022 | DG 18:2_20:0 | 0.986052128 |
| CountryMexico | 0.013 | 0.097 | 0.132 | 0.099 | PI 16:0_17:0 | 0.986052128 |
| CountryMexico | 0.020 | 0.152 | 0.129 | 0.022 | PI 18:1_20:1 | 0.986052128 |
| CountryMexico | 0.006 | 0.047 | 0.128 | 0.075 | Glucose | 0.986052128 |
| CountryMexico | -0.033 | 0.260 | -0.127 | 0.001 | TG 16:0_36:6 | 0.986052128 |
| CountryMexico | -0.032 | 0.240 | -0.132 | 0.069 | TG 16:0_38:4 | 0.986052128 |
| CountryMexico | -0.028 | 0.215 | -0.131 | 0.020 | TG 18:2_32:0 | 0.986052128 |
| CountryMexico | -0.008 | 0.059 | -0.129 | 0.078 | MCKAT Deficiency (NBS) | 0.986052128 |
| CountryMexico | 0.042 | 0.343 | 0.124 | 0.051 | TG 16:0_32:1 | 0.986376292 |
| CountryMexico | 0.012 | 0.105 | 0.118 | 0.023 | C16 | 0.986778539 |
| CountryMexico | -0.066 | 0.537 | -0.122 | 0.024 | TCA | 0.986778539 |
| CountryMexico | 0.023 | 0.190 | 0.121 | 0.014 | LPA 14:0 | 0.986778539 |
| CountryMexico | 0.026 | 0.219 | 0.118 | 0.056 | MG 22:1 | 0.986778539 |
| CountryMexico | -0.013 | 0.110 | -0.116 | 0.093 | PI 18:2_18:3 | 0.986778539 |
| CountryMexico | -0.038 | 0.317 | -0.121 | 0.042 | TG 22:6_32:1 | 0.986778539 |
| CountryMexico | 0.014 | 0.128 | 0.112 | 0.095 | HArg Synthesis | 0.988376477 |
| CountryMexico | 0.012 | 0.110 | 0.109 | 0.038 | PI 18:0_22:0 | 0.988734432 |
| CountryMexico | 0.019 | 0.198 | 0.097 | 0.031 | FA 22:6n-3 (DHA) | 0.991000108 |
| CountryMexico | 0.019 | 0.191 | 0.101 | 0.019 | PG 17:0_18:1 | 0.991000108 |
| CountryMexico | -0.009 | 0.095 | -0.099 | 0.100 | PG 18:1_20:3 | 0.991000108 |
| CountryMexico | 0.013 | 0.132 | 0.097 | 0.070 | Quinolinic acid | 0.991000108 |
| CountryMexico | -0.025 | 0.253 | -0.099 | 0.021 | TG 20:0_32:4 | 0.991000108 |
| CountryMexico | 0.007 | 0.074 | 0.091 | 0.023 | Ile | 0.991121731 |

| | | | | | | |
|---|---|---|---|---|---|---|
| CountryMexico | -0.020 | 0.245 | -0.082 | 0.067 | Itaconic acid | 0.991121731 |
| CountryMexico | -0.017 | 0.201 | -0.083 | 0.065 | FA 8:0 | 0.991121731 |
| CountryMexico | -0.019 | 0.235 | -0.080 | 0.040 | Cytidine | 0.991121731 |
| CountryMexico | 0.017 | 0.178 | 0.094 | 0.050 | PE P-16:0/16:0 | 0.991121731 |
| CountryMexico | 0.022 | 0.245 | 0.088 | 0.047 | PE P-18:0/20:1 | 0.991121731 |
| CountryMexico | -0.011 | 0.113 | -0.095 | 0.054 | PI 14:0_18:1 | 0.991121731 |
| CountryMexico | 0.015 | 0.193 | 0.076 | 0.060 | TG 18:2_35:1 | 0.991121731 |
| CountryMexico | -0.026 | 0.274 | -0.095 | 0.013 | TG 18:3_36:4 | 0.991121731 |
| CountryMexico | -0.018 | 0.199 | -0.089 | 0.042 | TG 22:5_34:2 | 0.991121731 |
| CountryMexico | 0.014 | 0.189 | 0.075 | 0.054 | Cer d18:1/25:0 | 0.991386868 |
| CountryMexico | 0.008 | 0.109 | 0.074 | 0.017 | OH-Phenyl-Lac | 0.991392622 |
| CountryMexico | -0.013 | 0.199 | -0.067 | 0.086 | C14 | 0.991588357 |
| CountryMexico | 0.004 | 0.068 | 0.058 | 0.041 | Lys | 0.991588357 |
| CountryMexico | 0.029 | 0.506 | 0.058 | 0.002 | TCDCA | 0.991588357 |
| CountryMexico | 0.008 | 0.157 | 0.049 | 0.041 | Cer d18:2/24:1 | 0.991588357 |
| CountryMexico | 0.004 | 0.078 | 0.056 | 0.068 | Fumaric acid | 0.991588357 |
| CountryMexico | 0.013 | 0.258 | 0.050 | 0.069 | LPE P-22:6 | 0.991588357 |
| CountryMexico | -0.005 | 0.082 | -0.063 | 0.070 | PI 16:0_17:2 | 0.991588357 |
| CountryMexico | 0.008 | 0.140 | 0.055 | 0.016 | PI 16:0_20:0 | 0.991588357 |
| CountryMexico | -0.006 | 0.096 | -0.062 | 0.089 | PG 18:1_22:2 | 0.991588357 |
| CountryMexico | -0.020 | 0.359 | -0.055 | 0.047 | TG 14:0_34:1 | 0.991588357 |
| CountryMexico | -0.017 | 0.329 | -0.053 | 0.079 | TG 16:0_35:1 | 0.991588357 |
| CountryMexico | -0.019 | 0.330 | -0.057 | 0.055 | TG 16:1_32:0 | 0.991588357 |
| CountryMexico | 0.020 | 0.287 | 0.068 | 0.138 | TG 17:1_34:3 | 0.991588357 |
| CountryMexico | 0.011 | 0.234 | 0.049 | 0.067 | TG 18:2_32:1 | 0.991588357 |
| CountryMexico | 0.026 | 0.504 | 0.052 | 0.012 | Sum of Conjugated Primary BAs | 0.991588357 |
| CountryMexico | 0.008 | 0.133 | 0.063 | 0.031 | Sum of SFA-PGs | 0.991588357 |
| CountryMexico | -0.006 | 0.100 | -0.056 | 0.109 | AGAT Deficiency | 0.991588357 |
| CountryMexico | -0.007 | 0.108 | -0.069 | 0.016 | Acetic Acid to Butyric Acid Ratio | 0.991588357 |
| CountryMexico | 0.008 | 0.138 | 0.055 | 0.087 | 5-HIAA to Creatinine Ratio | 0.991588357 |
| CountryMexico | 0.008 | 0.170 | 0.047 | 0.067 | PE 35:3 | 0.991779428 |
| CountryMexico | -0.005 | 0.110 | -0.047 | 0.002 | Cys Synthesis | 0.991779428 |
| CountryMexico | -0.005 | 0.103 | -0.047 | 0.016 | GABR | 0.991785958 |
| CountryMexico | -0.008 | 0.181 | -0.044 | 0.038 | FA 20:1n-9 | 0.991846161 |
| CountryMexico | 0.016 | 0.355 | 0.044 | 0.095 | TG 16:1_32:1 | 0.991846161 |
| CountryMexico | -0.017 | 0.432 | -0.040 | 0.003 | TG 18:1_36:0 | 0.991915638 |
| CountryMexico | -0.005 | 0.147 | -0.037 | 0.000 | Lac to FA 2:0 Ratio | 0.991915638 |
| CountryMexico | -0.007 | 0.199 | -0.033 | 0.003 | SPB d14:0 | 0.992832957 |
| CountryMexico | 0.005 | 0.162 | 0.030 | 0.051 | PE P-20:0/18:2 | 0.992951143 |
| CountryMexico | 0.002 | 0.073 | 0.025 | 0.019 | Leu | 0.993726388 |
| CountryMexico | 0.013 | 0.520 | 0.026 | 0.009 | GCDCA | 0.993726388 |

| | | | | | | |
|---|---|---|---|---|---|---|
| CountryMexico | 0.003 | 0.095 | 0.028 | 0.061 | PC O-36:3 | 0.993726388 |
| CountryMexico | 0.005 | 0.233 | 0.021 | 0.058 | TG 16:0_34:3 | 0.995394412 |
| CountryMexico | 0.005 | 0.271 | 0.017 | 0.031 | TMAO | 0.996431003 |
| CountryMexico | 0.001 | 0.095 | 0.014 | 0.133 | 3-MAG-uria Type 1 | 0.997553117 |
| CountryMexico | 0.002 | 0.165 | 0.012 | 0.102 | TG 18:1_38:5 | 0.997872437 |
| CountryMexico | -0.001 | 0.061 | -0.011 | 0.030 | Sum of Solely Ketogenic AAs | 0.998516427 |
| CountryMexico | 0.001 | 0.119 | 0.008 | 0.033 | FA 16:0 | 0.99908292 |
| CountryMexico | 0.001 | 0.207 | 0.004 | 0.023 | PA 16:1_22:0 | 0.999254492 |
| CountryMexico | 0.000 | 0.200 | 0.002 | 0.026 | TG 16:0_36:4 | 0.999254492 |
| CountryMexico | 0.001 | 0.312 | 0.004 | 0.002 | TG 18:1_36:6 | 0.999254492 |
| CountryMexico | -0.001 | 0.151 | -0.006 | 0.056 | BHMT Activity | 0.999254492 |

### SUPPLEMENTAL DATA 2

| Short hand name | Metabolite name | Analyte class | HMDB ID |
|---|---|---|---|
| C0 | Carnitine | Acylcarnitines | HMDB00062 |
| C2 | Acetylcarnitine | Acylcarnitines | HMDB00201 |
| C3 | Propionylcarnitine | Acylcarnitines | HMDB00824 |
| C3-DC (C4-OH) | Malonylcarnitine (Hydroxybutyrylcarnitine) | Acylcarnitines | HMDB02095 |
| C4 | Butyrylcarnitine | Acylcarnitines | HMDB02013 |
| C4:1 | Butenylcarnitine | Acylcarnitines | HMDB0013126 |
| C5 | Valerylcarnitine | Acylcarnitines | HMDB13128 |
| C5-DC (C6-OH) | Glutarylcarnitine (Hydroxyhexanoylcarnitine) | Acylcarnitines | HMDB0013130 |
| C5-OH (C3-DC-M) | Hydroxyvalerylcarnitine (Methylmalonylcarnitine) | Acylcarnitines | HMDB0013132 |
| C5:1 | Tiglylcarnitine | Acylcarnitines | HMDB0002366 |
| C8 | Octanoylcarnitine | Acylcarnitines | HMDB0000791 |
| C9 | Nonanoylcarnitine | Acylcarnitines | HMDB06320 |
| C10 | Decanoylcarnitine | Acylcarnitines | HMDB00651 |
| C10:1 | Decenoylcarnitine | Acylcarnitines | HMDB13205 |
| C10:2 | Decadienoylcarnitine | Acylcarnitines | HMDB0013325 |
| C12 | Dodecanoylcarnitine | Acylcarnitines | HMDB02250 |
| C12:1 | Dodecenoylcarnitine | Acylcarnitines | HMDB0013326 |
| C14 | Tetradecanoylcarnitine | Acylcarnitines | HMDB05066 |
| C14:1 | Tetradecenoylcarnitine | Acylcarnitines | HMDB0002014 |
| C14:1-OH | Hydroxytetradecenoylcarnitine | Acylcarnitines | HMDB0013330 |
| C14:2 | Tetradecadienoylcarnitine | Acylcarnitines | HMDB13331 |
| C14:2-OH | Hydroxytetradecadienoylcarnitine | Acylcarnitines | NA |
| C16 | Hexadecanoylcarnitine | Acylcarnitines | HMDB00222 |
| C16:1 | Hexadecenoylcarnitine | Acylcarnitines | HMDB06317 |
| C16:1-OH | Hydroxyhexadecenoylcarnitine | Acylcarnitines | HMDB13333 |
| C16:2 | Hexadecadienoylcarnitine | Acylcarnitines | HMDB13334 |
| C18 | Octadecanoylcarnitine | Acylcarnitines | HMDB00848 |
| C18:1 | Octadecenoylcarnitine | Acylcarnitines | HMDB0006351 |
| C18:1-OH | Hydroxyoctadecenoylcarnitine | Acylcarnitines | HMDB13339 |
| C18:2 | Octadecadienoylcarnitine | Acylcarnitines | HMDB0006469 |
| Trigonelline | Trigonelline | Alkaloids | HMDB0000875 |
| TMAO | Trimethylamine N-oxide | Amine oxides | HMDB0000925 |
| 1-Met-His | 1-Methylhistidine | Amino acid-related | HMDB0000001 |
| 3-Met-2-oxovaleric acid | 3-Methyl-2-oxovaleric acid | Amino acid-related | HMDB0000491 |
| 3-Met-His | 3-Methylhistidine | Amino acid-related | HMDB0000479 |
| 4-Guanidinobutanoic acid | 4-Guanidinobutanoic acid | Amino acid-related | HMDB0003464 |
| 4-Met-2-oxovaleric acid | 2-Oxoisocaproic acid | Amino acid-related | HMDB0000695 |
| 5-Amino-4-oxovaleric acid | 5-Amino-4-oxovaleric acid | Amino acid-related | HMDB0001149 |
| 5-AVA | 5-Aminovaleric acid | Amino acid-related | HMDB0003355 |

| | | | |
|---|---|---|---|
| 5-Oxo-Pro | 5-Oxoproline | Amino acid-related | HMDB0000267 |
| a-Ketoisovaleric acid | alpha-Ketoisovaleric acid | Amino acid-related | HMDB0000019 |
| AABA | alpha-Aminobutyric acid | Amino acid-related | HMDB0000452 |
| ADMA | Asymmetric dimethylarginine | Amino acid-related | HMDB0001539 |
| alpha-AAA | alpha-Aminoadipic acid | Amino acid-related | HMDB0000510 |
| Anthranilic acid | Anthranilic acid | Amino acid-related | HMDB0001123 |
| Arginino-Suc | Argininosuccinic acid | Amino acid-related | HMDB0000052 |
| BABA | beta-Aminobutyric acid | Amino acid-related | HMDB0031654 |
| BAIBA | 3-Aminoisobutyric acid | Amino acid-related | HMDB0003911 |
| Betaine | Betaine | Amino acid-related | HMDB0000043 |
| c4-OH-Pro | cis-4-Hydroxyproline | Amino acid-related | HMDB0240251 |
| Carnosine | Carnosine | Amino acid-related | HMDB0000033 |
| Cinnamoyl-Gly | Cinnamoylglycine | Amino acid-related | HMDB0011621 |
| Cit | Citrulline | Amino acid-related | HMDB0000904 |
| Creatine | Creatine | Amino acid-related | HMDB0000064 |
| Creatinine | Creatinine | Amino acid-related | HMDB0000562 |
| Cystine | Cystine | Amino acid-related | HMDB0000192 |
| DMG | Dimethylglycine (DMG) | Amino acid-related | HMDB0000092 |
| Guanidinoacetic acid | Guanidinoacetic acid | Amino acid-related | HMDB0000128 |
| HArg | Homoarginine | Amino acid-related | HMDB0000670 |
| HCit | Homocitrulline | Amino acid-related | HMDB0000679 |
| HCys | Homocysteine | Amino acid-related | HMDB0000742 |
| HSer | Homoserine | Amino acid-related | HMDB0000719 |
| Imidazolepropionic acid | Imidazolepropionic acid | Amino acid-related | HMDB0002271 |
| Kynurenine | Kynurenine | Amino acid-related | HMDB0000684 |
| Met-SO | Methionine sulfoxide | Amino acid-related | HMDB0002005 |
| N-Ac-Ala | N-Acetylalanine | Amino acid-related | HMDB0255053 |
| N-Ac-Arg | N-Acetylarginine | Amino acid-related | HMDB0004620 |
| N-Ac-Asn | N-Acetylasparagine | Amino acid-related | HMDB0006028 |
| N-Ac-Asp | N-Acetylaspartic acid | Amino acid-related | HMDB0000812 |
| N-Ac-Gln | N-Acetylglutamine | Amino acid-related | HMDB0006029 |
| N-Ac-Glu | N-Acetylglutamic acid | Amino acid-related | HMDB0001138 |
| N-Ac-Gly | N-Acetylglycine | Amino acid-related | HMDB0000532 |
| N-Ac-His | N-Acetylhistidine | Amino acid-related | HMDB0032055 |
| N-Ac-Ile | N-Acetylisoleucine | Amino acid-related | HMDB0061684 |
| N-Ac-Leu | N-Acetylleucine | Amino acid-related | HMDB0341345 |
| N-Ac-Met | N-Acetylmethionine | Amino acid-related | HMDB0011745 |
| N-Ac-Pro | N-Acetylproline | Amino acid-related | HMDB0094701 |
| N-Ac-Ser | N-Acetylserine | Amino acid-related | HMDB0002931 |
| N-Ac-Trp | N-Acetyltryptophan | Amino acid-related | HMDB0255052 |
| N-Ac-Tyr | N-Acetyltyrosine | Amino acid-related | HMDB0244966 |

| | | | |
|---|---|---|---|
| N-Ac-Val | N-Acetylvaline | Amino acid-related | HMDB0011757 |
| N-Met-Asp | N-Methylaspartic acid | Amino acid-related | HMDB0002393 |
| N2-Ac-Lys | N2-Acetyllysine | Amino acid-related | HMDB0000446 |
| N6-Ac-Lys | N6-Acetyllysine | Amino acid-related | HMDB0000206 |
| Orn | Ornithine | Amino acid-related | HMDB0000214 |
| Phenylacetylglutamine | Phenylacetylglutamine | Amino acid-related | HMDB0006344 |
| ProBetaine | Proline betaine | Amino acid-related | HMDB0004827 |
| Sarcosine | Sarcosine | Amino acid-related | HMDB0000271 |
| SDMA | Symmetric dimethylarginine | Amino acid-related | HMDB0003334 |
| t4-OH-Pro | trans-4-Hydroxyproline | Amino acid-related | HMDB0000725 |
| Taurine | Taurine | Amino acid-related | HMDB0000251 |
| TrpBetaine | Tryptophan betaine | Amino acid-related | HMDB0061115 |
| Ala | Alanine | Amino acids | HMDB0000161 |
| Arg | Arginine | Amino acids | HMDB0000517 |
| Asn | Asparagine | Amino acids | HMDB0000168 |
| Asp | Aspartic acid | Amino acids | HMDB0000191 |
| Cys | Cysteine | Amino acids | HMDB0000574 |
| Gln | Glutamine | Amino acids | HMDB0000641 |
| Glu | Glutamic acid | Amino acids | HMDB0000148 |
| Gly | Glycine | Amino acids | HMDB0000123 |
| His | Histidine | Amino acids | HMDB0000177 |
| Ile | Isoleucine | Amino acids | HMDB0000172 |
| Leu | Leucine | Amino acids | HMDB0000687 |
| Lys | Lysine | Amino acids | HMDB0000182 |
| Met | Methionine | Amino acids | HMDB0000696 |
| Phe | Phenylalanine | Amino acids | HMDB0000159 |
| Pro | Proline | Amino acids | HMDB0000162 |
| Ser | Serine | Amino acids | HMDB0000187 |
| Thr | Threonine | Amino acids | HMDB0000167 |
| Trp | Tryptophan | Amino acids | HMDB0000929 |
| Tyr | Tyrosine | Amino acids | HMDB0000158 |
| Val | Valine | Amino acids | HMDB0000883 |
| 3-EpiDCA | 3-Epideoxycholic acid | Bile acids | HMDB0000438 |
| 7-KetoDCA | 7-Ketodeoxycholic acid | Bile acids | HMDB0000391 |
| 12-KetoDCA | 12-Ketodeoxycholic acid | Bile acids | HMDB0000328 |
| CA | Cholic acid | Bile acids | HMDB0000619 |
| CDCA | Chenodeoxycholic acid | Bile acids | HMDB0000518 |
| DCA | Deoxycholic acid | Bile acids | HMDB0000626 |
| GCA | Glycocholic acid | Bile acids | HMDB0000138 |
| GCDCA | Glycochenodeoxycholic acid | Bile acids | HMDB0000637 |
| GDCA | Glycodeoxycholic acid | Bile acids | HMDB0000631 |

| | | | |
|---|---|---|---|
| GLCA | Glycolithocholic acid | Bile acids | HMDB0000698 |
| GLCAS | Glycolithocholic acid sulfate | Bile acids | HMDB0002639 |
| GUDCA | Glycoursodeoxycholic acid | Bile acids | HMDB0000708 |
| IsoLCA | Isolithocholic acid | Bile acids | HMDB0000717 |
| IsoUDCA | Isoursodeoxycholic acid | Bile acids | HMDB0000686 |
| NorDCA | Nordeoxycholic acid | Bile acids | HMDB0304947 |
| TCA | Taurocholic acid | Bile acids | HMDB0000036 |
| TCDCA | Taurochenodeoxycholic acid | Bile acids | HMDB0000951 |
| TDCA | Taurodeoxycholic acid | Bile acids | HMDB0000896 |
| THDCA | Taurohyodeoxycholic acid | Bile acids | NA |
| TLCA | Taurolithocholic acid | Bile acids | HMDB0000722 |
| TUDCA | Tauroursodeoxycholic acid | Bile acids | HMDB0000874 |
| UDCA | Ursodeoxycholic acid | Bile acids | HMDB0000946 |
| beta-Ala | beta-Alanine | Biogenic amines | HMDB0000056 |
| GABA | gamma-Aminobutyric acid | Biogenic amines | HMDB0000112 |
| Histamine | Histamine | Biogenic amines | HMDB0000870 |
| Putrescine | Putrescine | Biogenic amines | HMDB0001414 |
| Serotonin | Serotonin | Biogenic amines | HMDB0000259 |
| Spermidine | Spermidine | Biogenic amines | HMDB0001257 |
| Urea | Urea | Biogenic amines | HMDB0000294 |
| 2-OH-Butyric acid | 2-Hydroxybutyric acid | Carboxylic acids | HMDB0000008 |
| 3-OH-Butyric acid | 3-Hydroxybutyric acid | Carboxylic acids | HMDB0000442 |
| Glycolic acid | Glycolic acid | Carboxylic acids | HMDB0000115 |
| Glyoxylic acid | Glyoxylic acid | Carboxylic acids | HMDB0000119 |
| HipAcid | Hippuric acid | Carboxylic acids | HMDB0000714 |
| Lac | Lactic acid | Carboxylic acids | HMDB0000190 |
| Mevalonic acid | Mevalonic acid | Carboxylic acids | HMDB0000227 |
| Pyruvic acid | Pyruvic acid | Carboxylic acids | HMDB0000243 |
| Cer d16:1/18:0 | Ceramide d16:1/18:0 | Ceramides | NA |
| Cer d16:1/20:0 | Ceramide d16:1/20:0 | Ceramides | NA |
| Cer d16:1/22:0 | Ceramide d16:1/22:0 | Ceramides | NA |
| Cer d16:1/23:0 | Ceramide d16:1/23:0 | Ceramides | NA |
| Cer d16:1/24:0 | Ceramide d16:1/24:0 | Ceramides | NA |
| Cer d18:1/16:0 | Ceramide d18:1/16:0 | Ceramides | HMDB0004949 |
| Cer d18:1/18:0 | Ceramide d18:1/18:0 | Ceramides | HMDB0004950 |
| Cer d18:1/20:0 | Ceramide d18:1/20:0 | Ceramides | HMDB0004951 |
| Cer d18:1/22:0 | Ceramide d18:1/22:0 | Ceramides | HMDB0004952 |
| Cer d18:1/23:0 | Ceramide d18:1/23:0 | Ceramides | HMDB0000950 |
| Cer d18:1/24:0 | Ceramide d18:1/24:0 | Ceramides | HMDB0004956 |
| Cer d18:1/24:1 | Ceramide d18:1/24:1 | Ceramides | HMDB0004953 |
| Cer d18:1/25:0 | Ceramide d18:1/25:0 | Ceramides | HMDB0004957 |

| | | | |
|---|---|---|---|
| Cer d18:1/26:0 | Ceramide d18:1/26:0 | Ceramides | HMDB0004955 |
| Cer d18:1/26:1 | Ceramide d18:1/26:1 | Ceramides | HMDB0004954 |
| Cer d18:2/16:0 | Ceramide d18:2/16:0 | Ceramides | NA |
| Cer d18:2/18:0 | Ceramide d18:2/18:0 | Ceramides | NA |
| Cer d18:2/20:0 | Ceramide d18:2/20:0 | Ceramides | NA |
| Cer d18:2/22:0 | Ceramide d18:2/22:0 | Ceramides | NA |
| Cer d18:2/23:0 | Ceramide d18:2/23:0 | Ceramides | NA |
| Cer d18:2/24:0 | Ceramide d18:2/24:0 | Ceramides | NA |
| Cer d18:2/24:1 | Ceramide d18:2/24:1 | Ceramides | NA |
| CerP d18:1/16:0 | Ceramide phosphate d18:1/16:0 | Ceramides | HMDB0010700 |
| CE 14:0 | Cholesteryl ester 14:0 | Cholesteryl esters | HMDB0006725 |
| CE 16:0 | Cholesteryl ester 16:0 | Cholesteryl esters | HMDB00885 |
| CE 16:1 | Cholesteryl ester 16:1 | Cholesteryl esters | HMDB00658 |
| CE 17:1 | Cholesteryl ester 17:1 | Cholesteryl esters | HMDB0060060 |
| CE 18:0 | Cholesteryl ester 18:0 | Cholesteryl esters | HMDB0062461 |
| CE 18:1 | Cholesteryl ester 18:1 | Cholesteryl esters | HMDB00918 |
| CE 18:2 | Cholesteryl ester 18:2 | Cholesteryl esters | HMDB05192 |
| CE 18:3 | Cholesteryl ester 18:3 | Cholesteryl esters | HMDB0010370 |
| CE 20:3 | Cholesteryl ester 20:3 | Cholesteryl esters | HMDB06736 |
| CE 20:4 | Cholesteryl ester 20:4 | Cholesteryl esters | HMDB06726 |
| CE 20:5 | Cholesteryl ester 20:5 | Cholesteryl esters | HMDB06731 |
| CE 22:2 | Cholesteryl ester 22:2 | Cholesteryl esters | HMDB06737 |
| CE 22:5 | Cholesteryl ester 22:5 | Cholesteryl esters | HMDB10374 |
| CE 22:6 | Cholesteryl ester 22:6 | Cholesteryl esters | HMDB06733 |
| p-Cresol glucuronide | p-Cresol glucuronide | Cresols | HMDB0011686 |
| p-Cresol-SO4 | p-Cresol sulfate | Cresols | HMDB0011635 |
| 2-OH-Glutaric acid | 2-Hydroxyglutaric acid | Dicarboxylic acids | HMDB0000694 |
| 3-HMGA | 3-Hydroxymethylglutaric acid | Dicarboxylic acids | HMDB0000355 |
| 3-Met-adipic acid | 3-Methyladipic acid | Dicarboxylic acids | HMDB0000555 |
| 3-Met-glutaric acid | 3-Methylglutaric acid | Dicarboxylic acids | HMDB0000752 |
| 3-OH-Glutaric acid | 3-Hydroxyglutaric acid | Dicarboxylic acids | HMDB0000428 |
| 3-OH-Sebacic acid | 3-Hydroxysebacic acid | Dicarboxylic acids | HMDB0000350 |
| a-Ketoglutaric acid | alpha-Ketoglutaric acid | Dicarboxylic acids | HMDB0000208 |
| Adipic acid | Adipic acid | Dicarboxylic acids | HMDB0000448 |
| Citramalic acid | Citramalic acid | Dicarboxylic acids | HMDB0000426 |
| DiCA 12:0 | Dodecanedioic acid | Dicarboxylic acids | HMDB0000623 |
| DiCA 14:0 | Tetradecanedioic acid | Dicarboxylic acids | HMDB0000872 |
| E-3-Met-glutaconic acid | E-3-methylglutaconic acid | Dicarboxylic acids | HMDB0000522 |
| Ethylmalonic acid | Ethylmalonic acid | Dicarboxylic acids | HMDB0000622 |
| Fumaric acid | Fumaric acid | Dicarboxylic acids | HMDB0000134 |
| Glutaric acid | Glutaric acid | Dicarboxylic acids | HMDB0000661 |

| | | | |
|---|---|---|---|
| Itaconic acid | Itaconic acid | Dicarboxylic acids | HMDB0002092 |
| Maleic acid | Maleic acid | Dicarboxylic acids | HMDB0000176 |
| Malic acid | Malic acid | Dicarboxylic acids | HMDB0000156 |
| Malonic acid | Malonic acid | Dicarboxylic acids | HMDB0000691 |
| Methylmalonic acid | Methylmalonic acid | Dicarboxylic acids | HMDB0000202 |
| Oxalic acid | Oxalic acid | Dicarboxylic acids | HMDB0002329 |
| Oxaloacetic acid | Oxaloacetic acid | Dicarboxylic acids | HMDB0000223 |
| Suberic acid | Suberic acid | Dicarboxylic acids | HMDB0000893 |
| Suc | Succinic acid | Dicarboxylic acids | HMDB0000254 |
| Z-3-Met-glutaconic acid | Z-3-Methylglutaconic acid | Dicarboxylic acids | NA |
| DG 14:1_18:1 | Diacylglyceride 14:1_18:1 | Diglycerides | HMDB0007044 |
| DG 16:0_16:1 | Diacylglyceride 16:0_16:1 | Diglycerides | HMDB0007099 |
| DG 16:0_18:1 | Diacylglyceride 16:0_18:1 | Diglycerides | HMDB0007214 |
| DG 16:0_18:2 | Diacylglyceride 16:0_18:2 | Diglycerides | HMDB0007103 |
| DG 16:1_18:0 | Diacylglyceride 16:1_18:0 | Diglycerides | HMDB0007129 |
| DG 16:1_18:1 | Diacylglyceride 16:1_18:1 | Diglycerides | HMDB0007131 |
| DG 16:1_18:2 | Diacylglyceride 16:1_18:2 | Diglycerides | HMDB0007132 |
| DG 17:0_18:1 | Diacylglyceride 17:0_18:1 | Diglycerides | NA |
| DG 18:1_18:1 | Diacylglyceride 18:1_18:1 | Diglycerides | HMDB0007218 |
| DG 18:1_18:2 | Diacylglyceride 18:1_18:2 | Diglycerides | HMDB0007219 |
| DG 18:1_18:3 | Diacylglyceride 18:1_18:3 | Diglycerides | HMDB0007221 |
| DG 18:1_20:1 | Diacylglyceride 18:1_20:1 | Diglycerides | HMDB0007224 |
| DG 18:1_20:2 | Diacylglyceride 18:1_20:2 | Diglycerides | HMDB0007225 |
| DG 18:1_20:4 | Diacylglyceride 18:1_20:4 | Diglycerides | HMDB0007228 |
| DG 18:2_18:2 | Diacylglyceride 18:2_18:2 | Diglycerides | HMDB0007248 |
| DG 18:2_18:4 | Diacylglyceride 18:2_18:4 | Diglycerides | HMDB0007335 |
| DG 18:2_20:0 | Diacylglyceride 18:2_20:0 | Diglycerides | HMDB0007252 |
| DG 18:2_20:4 | Diacylglyceride 18:2_20:4 | Diglycerides | HMDB0007257 |
| Hex2Cer d18:1/14:0 | Dihexosylceramide d18:1/14:0 | Dihexosylceramides | NA |
| Hex2Cer d18:1/16:0 | Dihexosylceramide d18:1/16:0 | Dihexosylceramides | HMDB0004833 |
| Hex2Cer d18:1/18:0 | Dihexosylceramide d18:1/18:0 | Dihexosylceramides | HMDB0004834 |
| Hex2Cer d18:1/22:0 | Dihexosylceramide d18:1/22:0 | Dihexosylceramides | HMDB0004836 |
| Hex2Cer d18:1/24:0 | Dihexosylceramide d18:1/24:0 | Dihexosylceramides | HMDB0004840 |
| Hex2Cer d18:1/24:1 | Dihexosylceramide d18:1/24:1 | Dihexosylceramides | HMDB0004837 |
| Cer d18:0/22:0 | Ceramide d18:0/22:0 | Dihydroceramides | HMDB0011765 |
| Cer d18:0/24:1 | Ceramide d18:0/24:1 | Dihydroceramides | HMDB0011769 |
| FA 2:0 | Acetic acid | Fatty acids | HMDB0000042 |
| FA 3:0-2M | Isobutyric acid | Fatty acids | HMDB0001873 |
| FA 4:0 | Butyric acid | Fatty acids | HMDB0000039 |
| FA 4:0-2M | 2-Methylbutyric acid | Fatty acids | HMDB0002176 |
| FA 4:0-3M | Isovaleric acid | Fatty acids | HMDB0000718 |

| | | | |
|---|---|---|---|
| FA 5:0-3M | 3-Methylvaleric acid | Fatty acids | HMDB0033774 |
| FA 5:0-4M | Isocaproic acid | Fatty acids | HMDB0000689 |
| FA 6:0 | Caproic acid | Fatty acids | HMDB0000535 |
| FA 7:0 | Heptanoic acid | Fatty acids | HMDB0000666 |
| FA 8:0 | Caprylic acid | Fatty acids | HMDB0000482 |
| FA 9:0 | Nonanoic acid | Fatty acids | HMDB0000847 |
| FA 10:0 | Capric acid | Fatty acids | HMDB0000511 |
| FA 11:0 | Undecanoic acid | Fatty acids | HMDB0000947 |
| FA 12:0 | Lauric acid | Fatty acids | HMDB0000638 |
| FA 14:0 | Myristic acid | Fatty acids | HMDB0000806 |
| FA 14:1n-5 | Myristoleic acid | Fatty acids | HMDB0002000 |
| FA 15:0 | Pentadecanoic acid | Fatty acids | HMDB0000826 |
| FA 16:0 | Palmitic acid | Fatty acids | HMDB0000220 |
| FA 16:1 | Palmitoleic acid | Fatty acids | HMDB0012328 |
| FA 18:0 | Stearic acid | Fatty acids | HMDB0000827 |
| FA 18:1 | Oleic acid | Fatty acids | HMDB0000573 |
| FA 18:2 | Linoleic acid | Fatty acids | HMDB0000673 |
| FA 18:3 | Octadecatrienoic acid | Fatty acids | HMDB0001388 |
| FA 20:0 | Arachidic acid | Fatty acids | HMDB0002212 |
| FA 20:1n-9 | Gondoic acid | Fatty acids | HMDB0002231 |
| FA 20:2 | Eicosadienoic acid | Fatty acids | HMDB0005060 |
| FA 20:3n-6 | Dihomo-gamma-linolenic acid (omega6) | Fatty acids | HMDB0002925 |
| FA 20:3n-9 | Mead acid | Fatty acids | HMDB0010378 |
| FA 20:4n-6 (AA) | Arachidonic acid (AA, omega6) | Fatty acids | HMDB0001043 |
| FA 20:5n-3 (EPA) | Eicosapentaenoic acid (EPA, omega3) | Fatty acids | HMDB0001999 |
| FA 22:4n-6 | Docosatetraenoic acid (omega6) | Fatty acids | HMDB0002226 |
| FA 22:5n-3 (DPA) | Docosapentaenoic acid (DPA, omega3) | Fatty acids | HMDB0001976 |
| FA 22:6n-3 (DHA) | Docosahexaenoic acid (DHA, omega3) | Fatty acids | HMDB0002183 |
| FA 24:0 | Lignoceric acid | Fatty acids | HMDB0002003 |
| FA 24:1n-9 | Nervonic acid | Fatty acids | HMDB0002368 |
| FA 24:4n-6 | Tetracosatetraenoic acid (omega6) | Fatty acids | HMDB0006246 |
| Hex-Cer d16:1/20:0 | Hexosylceramide d16:1/20:0 | Hexosylceramides | NA |
| Hex-Cer d18:1/14:0 | Hexosylceramide d18:1/14:0 | Hexosylceramides | HMDB0012321 |
| Hex-Cer d18:1/16:0 | Hexosylceramide d18:1/16:0 | Hexosylceramides | HMDB0004971 |
| Hex-Cer d18:1/18:0 | Hexosylceramide d18:1/18:0 | Hexosylceramides | HMDB0010709 |
| Hex-Cer d18:1/18:1 | Hexosylceramide d18:1/18:1 | Hexosylceramides | HMDB0010714 |
| Hex-Cer d18:1/20:0 | Hexosylceramide d18:1/20:0 | Hexosylceramides | HMDB0010710 |
| Hex-Cer d18:1/22:0 | Hexosylceramide d18:1/22:0 | Hexosylceramides | HMDB0010711 |
| Hex-Cer d18:1/23:0 | Hexosylceramide d18:1/23:0 | Hexosylceramides | NA |
| Hex-Cer d18:1/24:0 | Hexosylceramide d18:1/24:0 | Hexosylceramides | HMDB0004978 |
| Hex-Cer d18:1/24:1 | Hexosylceramide d18:1/24:1 | Hexosylceramides | HMDB0010712 |

| | | | |
|---|---|---|---|
| Hex-Cer d18:1/26:1 | Hexosylceramide d18:1/26:1 | Hexosylceramides | HMDB0010713 |
| Hex-Cer d18:2/16:0 | Hexosylceramide d18:2/16:0 | Hexosylceramides | NA |
| Hex-Cer d18:2/18:0 | Hexosylceramide d18:2/18:0 | Hexosylceramides | NA |
| Hex-Cer d18:2/20:0 | Hexosylceramide d18:2/20:0 | Hexosylceramides | NA |
| Hex-Cer d18:2/22:0 | Hexosylceramide d18:2/22:0 | Hexosylceramides | NA |
| Hex-Cer d18:2/23:0 | Hexosylceramide d18:2/23:0 | Hexosylceramides | NA |
| Hex-Cer d18:2/24:0 | Hexosylceramide d18:2/24:0 | Hexosylceramides | NA |
| Cortisol | Cortisol | Hormones | HMDB0000063 |
| Cortisone | Cortisone | Hormones | HMDB0002802 |
| DHEAS | Dehydroepiandrosterone sulfate | Hormones | HMDB0001032 |
| T4 | Thyroxine (T4) | Hormones | HMDB0000248 |
| 3-IAA | 3-Indoleacetic acid | Indoles and derivatives | HMDB0000197 |
| 3-IPA | 3-Indolepropionic acid | Indoles and derivatives | HMDB0002302 |
| 5-HIAA | 5-Hydroxyindoleacetic acid | Indoles and derivatives | HMDB0000763 |
| IAG | Indolylacryloylglycine | Indoles and derivatives | HMDB0006005 |
| Ind-SO4 | Indoxyl sulfate | Indoles and derivatives | HMDB0000682 |
| Indole-Lac | Indolelactic acid | Indoles and derivatives | HMDB0000671 |
| Indoleacetaldehyde | Indoleacetaldehyde | Indoles and derivatives | HMDB0001190 |
| Met-IAA | Methylindole-3-acetic acid | Indoles and derivatives | HMDB0029738 |
| LPA 14:0 | Lysophosphatidic acid 14:0 | Lysophosphatidic acids | HMDB0062321 |
| LPA 14:1 | Lysophosphatidic acid 14:1 | Lysophosphatidic acids | HMDB0062311 |
| LPA 15:0 | Lysophosphatidic acid 15:0 | Lysophosphatidic acids | HMDB0062324 |
| LPE 14:0 | Lysophosphatidylethanolamine 14:0 | Lysophosphatidylethanolamines | HMDB0011500 |
| LPE 15:0 | Lysophosphatidylethanolamine 15:0 | Lysophosphatidylethanolamines | HMDB0011502 |
| LPE 16:0 | Lysophosphatidylethanolamine 16:0 | Lysophosphatidylethanolamines | HMDB0011503 |
| LPE 17:0 | Lysophosphatidylethanolamine 17:0 | Lysophosphatidylethanolamines | HMDB0061691 |
| LPE 17:1 | Lysophosphatidylethanolamine 17:1 | Lysophosphatidylethanolamines | NA |
| LPE 18:0 | Lysophosphatidylethanolamine 18:0 | Lysophosphatidylethanolamines | HMDB0011130 |
| LPE 18:1 | Lysophosphatidylethanolamine 18:1 | Lysophosphatidylethanolamines | HMDB0011506 |
| LPE 18:2 | Lysophosphatidylethanolamine 18:2 | Lysophosphatidylethanolamines | HMDB0011507 |
| LPE 18:3 | Lysophosphatidylethanolamine 18:3 | Lysophosphatidylethanolamines | HMDB0011508 |
| LPE 19:0 | Lysophosphatidylethanolamine 19:0 | Lysophosphatidylethanolamines | HMDB0062322 |
| LPE 20:2 | Lysophosphatidylethanolamine 20:2 | Lysophosphatidylethanolamines | HMDB0011513 |
| LPE 20:3 | Lysophosphatidylethanolamine 20:3 | Lysophosphatidylethanolamines | HMDB0011516 |
| LPE 20:4 | Lysophosphatidylethanolamine 20:4 | Lysophosphatidylethanolamines | HMDB0011517 |
| LPE 20:5 | Lysophosphatidylethanolamine 20:5 | Lysophosphatidylethanolamines | HMDB0011519 |
| LPE 22:0 | Lysophosphatidylethanolamine 22:0 | Lysophosphatidylethanolamines | HMDB0011520 |
| LPE 22:4 | Lysophosphatidylethanolamine 22:4 | Lysophosphatidylethanolamines | HMDB0011523 |
| LPE 22:5 | Lysophosphatidylethanolamine 22:5 | Lysophosphatidylethanolamines | HMDB0011494 |
| LPE 22:6 | Lysophosphatidylethanolamine 22:6 | Lysophosphatidylethanolamines | HMDB0011526 |
| LPE P-14:0 | Lysophosphatidylethanolamine P-14:0 | Lysophosphatidylethanolamines | NA |

| | | | |
|---|---|---|---|
| LPE P-16:0 | Lysophosphatidylethanolamine P-16:0 | Lysophosphatidylethanolamines | HMDB0011152 |
| LPE P-17:0 | Lysophosphatidylethanolamine P-17:0 | Lysophosphatidylethanolamines | NA |
| LPE P-18:0 | Lysophosphatidylethanolamine P-18:0 | Lysophosphatidylethanolamines | HMDB0240598 |
| LPE P-18:1 | Lysophosphatidylethanolamine P-18:1 | Lysophosphatidylethanolamines | HMDB0240599 |
| LPE P-20:0 | Lysophosphatidylethanolamine P-20:0 | Lysophosphatidylethanolamines | NA |
| LPE P-22:0 | Lysophosphatidylethanolamine P-22:0 | Lysophosphatidylethanolamines | NA |
| LPE P-22:1 | Lysophosphatidylethanolamine P-22:1 | Lysophosphatidylethanolamines | NA |
| LPE P-22:6 | Lysophosphatidylethanolamine P-22:6 | Lysophosphatidylethanolamines | NA |
| LPS 18:0 | Lysophosphatidylserine 18:0 | Lysophosphatidylserines | HMDB0240606 |
| LPS 18:1 | Lysophosphatidylserine 18:1 | Lysophosphatidylserines | HMDB0240603 |
| LPS 18:2 | Lysophosphatidylserine 18:2 | Lysophosphatidylserines | HMDB0240604 |
| LPS 20:1 | Lysophosphatidylserine 20:1 | Lysophosphatidylserines | NA |
| LPS 20:5 | Lysophosphatidylserine 20:5 | Lysophosphatidylserines | NA |
| LPC 14:0 | Lysophosphatidylcholine 14:0 | Lysophosphatidylcholines | HMDB10379 |
| LPC 16:0 | Lysophosphatidylcholine 16:0 | Lysophosphatidylcholines | HMDB10382 |
| LPC 16:1 | Lysophosphatidylcholine 16:1 | Lysophosphatidylcholines | HMDB0010383 |
| LPC 17:0 | Lysophosphatidylcholine 17:0 | Lysophosphatidylcholines | HMDB12108 |
| LPC 18:0 | Lysophosphatidylcholine 18:0 | Lysophosphatidylcholines | HMDB10384 |
| LPC 18:1 | Lysophosphatidylcholine 18:1 | Lysophosphatidylcholines | HMDB02815 |
| LPC 18:2 | Lysophosphatidylcholine 18:2 | Lysophosphatidylcholines | HMDB10386 |
| LPC 20:3 | Lysophosphatidylcholine 20:3 | Lysophosphatidylcholines | HMDB10394 |
| LPC 20:4 | Lysophosphatidylcholine 20:4 | Lysophosphatidylcholines | HMDB10395 |
| LPC 24:0 | Lysophosphatidylcholine 24:0 | Lysophosphatidylcholines | HMDB10405 |
| LPC 26:0 | Lysophosphatidylcholine 26:0 | Lysophosphatidylcholines | HMDB0029205 |
| LPC 26:1 | Lysophosphatidylcholine 26:1 | Lysophosphatidylcholines | HMDB0029220 |
| LPI 14:0 | Lysophosphatidylinositol 14:0 | Lysophosphatidylinositols | NA |
| LPI 14:1 | Lysophosphatidylinositol 14:1 | Lysophosphatidylinositols | NA |
| LPI 16:0 | Lysophosphatidylinositol 16:0 | Lysophosphatidylinositols | HMDB0061695 |
| LPI 16:1 | Lysophosphatidylinositol 16:1 | Lysophosphatidylinositols | NA |
| LPI 17:0 | Lysophosphatidylinositol 17:0 | Lysophosphatidylinositols | NA |
| LPI 17:1 | Lysophosphatidylinositol 17:1 | Lysophosphatidylinositols | NA |
| LPI 18:0 | Lysophosphatidylinositol 18:0 | Lysophosphatidylinositols | HMDB0240261 |
| LPI 18:1 | Lysophosphatidylinositol 18:1 | Lysophosphatidylinositols | HMDB0061693 |
| LPI 18:2 | Lysophosphatidylinositol 18:2 | Lysophosphatidylinositols | HMDB0240597 |
| LPI 18:3 | Lysophosphatidylinositol 18:3 | Lysophosphatidylinositols | NA |
| LPI 19:0 | Lysophosphatidylinositol 19:0 | Lysophosphatidylinositols | NA |
| LPI 20:1 | Lysophosphatidylinositol 20:1 | Lysophosphatidylinositols | NA |
| LPI 20:4 | Lysophosphatidylinositol 20:4 | Lysophosphatidylinositols | HMDB0061690 |
| LPI 22:0 | Lysophosphatidylinositol 22:0 | Lysophosphatidylinositols | NA |
| LPI 22:1 | Lysophosphatidylinositol 22:1 | Lysophosphatidylinositols | NA |
| MG 20:1 | Monoglyceride 20:1 | Monoglycerides | HMDB0011543 |

| | | | |
|---|---|---|---|
| MG 22:1 | Monoglyceride 22:1 | Monoglycerides | HMDB0011552 |
| MG 22:2 | Monoglyceride 22:2 | Monoglycerides | HMDB0011553 |
| MG 22:4 | Monoglyceride 22:4 | Monoglycerides | HMDB0011554 |
| Adenosine | Adenosine | Nucleobase-related | HMDB0000050 |
| AHCys | S-Adenosylhomocysteine | Nucleobase-related | HMDB0000939 |
| Cytidine | Cytidine | Nucleobase-related | HMDB0000089 |
| Hypoxanthine | Hypoxanthine | Nucleobase-related | HMDB0000157 |
| N-Ac-Cytidine | N4-Acetylcytidine | Nucleobase-related | HMDB0005923 |
| Orotic acid | Orotic acid | Nucleobase-related | HMDB0000226 |
| Uric acid | Uric acid | Nucleobase-related | HMDB0000289 |
| Uridine | Uridine | Nucleobase-related | HMDB0000296 |
| Xanthine | Xanthine | Nucleobase-related | HMDB0000292 |
| cAMP | Adenosine monophosphate, cyclic (cAMP) | Nucleotides | HMDB0000058 |
| cGMP | Guanosine monophosphate, cyclic (cGMP) | Nucleotides | HMDB0001314 |
| 2-OH-2-Met-butyric acid | 2-Hydroxy-2-methylbutyric acid | Organic acids | HMDB0001987 |
| 2-OH-Isobutyric acid | 2-Hydroxyisobutyric acid | Organic acids | HMDB0000729 |
| 2,5-Furandicarboxylic acid | Furan-2,5-dicarboxylic acid | Organic acids | HMDB0004812 |
| 3-OH-Isobutyric acid | 3-Hydroxyisobutyric acid | Organic acids | HMDB0000023 |
| 4-OH-HipAcid | 4-Hydroxyhippuric acid | Organic acids | HMDB0013678 |
| Argininic acid | Argininic acid | Organic acids | HMDB0003148 |
| Bilirubin | Bilirubin | Organic acids | HMDB0000054 |
| Biliverdin | Biliverdin | Organic acids | HMDB0001008 |
| CMPF | 3-Carboxy-4-methyl-5-propyl-2-furanpropionic acid | Organic acids | HMDB0061112 |
| Glyceric acid | Glyceric acid | Organic acids | HMDB0006372 |
| 2-OH-Benzoic acid | 2-Hydroxybenzoic acid | Phenolic acids | HMDB0001895 |
| 2-OH-Phenylacetic acid | 2-Hydroxyphenylacetic acid | Phenolic acids | HMDB0000669 |
| 3,5-DiOH-Benzoic acid | 3,5-Dihydroxybenzoic acid | Phenolic acids | HMDB0013677 |
| 4-OH-Phenylacetic acid | 4-Hydroxyphenylacetic acid | Phenolic acids | HMDB0000020 |
| 4-OH-Phenylpyruvic acid | 4-Hydroxyphenylpyruvic acid | Phenolic acids | HMDB0000707 |
| Benzoic acid | Benzoic acid | Phenolic acids | HMDB0001870 |
| Desamino-Tyr | Desaminotyrosine | Phenolic acids | HMDB0002199 |
| Homovanillic acid | Homovanillic acid | Phenolic acids | HMDB0000118 |
| HPHPA | 3-(3-Hydroxyphenyl)-3-hydroxypropanoic acid | Phenolic acids | HMDB0002643 |
| OH-Phenyl-Lac | Hydroxyphenyllactic acid | Phenolic acids | HMDB0000755 |
| Phenyl-Lac | Phenyllactic acid | Phenolic acids | HMDB0000779 |
| Phenylacetic acid | Phenylacetic acid | Phenolic acids | HMDB0000209 |
| Phenylpyruvic acid | Phenylpyruvic acid | Phenolic acids | HMDB0000205 |
| Vanillylmandelic acid | Vanillylmandelic acid | Phenolic acids | HMDB0000291 |
| 4-EPS | 4-Ethylphenyl sulfate | Phenoxy compounds | HMDB0062551 |
| Phenylglucuronide | Phenylglucuronide | Phenoxy compounds | HMDB0059806 |
| PA 14:0_14:1 | Phosphatidic acid 14:0_14:1 | Phosphatidic acids | HMDB0114795 |

| | | | |
|---|---|---|---|
| PA 16:0_18:1 | Phosphatidic acid 16:0_18:1 | Phosphatidic acids | HMDB0114877 |
| PA 16:0_18:2 | Phosphatidic acid 16:0_18:2 | Phosphatidic acids | HMDB0114949 |
| PA 16:0_18:3 | Phosphatidic acid 16:0_18:3 | Phosphatidic acids | HMDB0114836 |
| PA 16:0_19:2 | Phosphatidic acid 16:0_19:2 | Phosphatidic acids | NA |
| PA 16:1_18:1 | Phosphatidic acid 16:1_18:1 | Phosphatidic acids | HMDB0114900 |
| PA 16:1_18:2 | Phosphatidic acid 16:1_18:2 | Phosphatidic acids | HMDB0114856 |
| PA 16:1_22:0 | Phosphatidic acid 16:1_22:0 | Phosphatidic acids | HMDB0114864 |
| PA 16:2_18:1 | Phosphatidic acid 16:2_18:1 | Phosphatidic acids | NA |
| PA 17:0_18:1 | Phosphatidic acid 17:0_18:1 | Phosphatidic acids | NA |
| PA 17:0_18:2 | Phosphatidic acid 17:0_18:2 | Phosphatidic acids | NA |
| PA 17:1_18:1 | Phosphatidic acid 17:1_18:1 | Phosphatidic acids | NA |
| PA 17:1_18:2 | Phosphatidic acid 17:1_18:2 | Phosphatidic acids | NA |
| PA 17:2_18:1 | Phosphatidic acid 17:2_18:1 | Phosphatidic acids | NA |
| PA 18:0_18:1 | Phosphatidic acid 18:0_18:1 | Phosphatidic acids | HMDB0114901 |
| PA 18:0_18:2 | Phosphatidic acid 18:0_18:2 | Phosphatidic acids | HMDB0114951 |
| PA 18:0_18:3 | Phosphatidic acid 18:0_18:3 | Phosphatidic acids | HMDB0114878 |
| PA 18:1_18:1 | Phosphatidic acid 18:1_18:1 | Phosphatidic acids | HMDB0007862 |
| PA 18:1_18:2 | Phosphatidic acid 18:1_18:2 | Phosphatidic acids | HMDB0114902 |
| PA 18:1_18:3 | Phosphatidic acid 18:1_18:3 | Phosphatidic acids | HMDB0114903 |
| PA 18:1_18:4 | Phosphatidic acid 18:1_18:4 | Phosphatidic acids | HMDB0114905 |
| PA 18:1_20:0 | Phosphatidic acid 18:1_20:0 | Phosphatidic acids | HMDB0114906 |
| PA 18:1_20:1 | Phosphatidic acid 18:1_20:1 | Phosphatidic acids | HMDB0114907 |
| PA 18:1_20:2 | Phosphatidic acid 18:1_20:2 | Phosphatidic acids | HMDB0115516 |
| PA 18:1_20:3 | Phosphatidic acid 18:1_20:3 | Phosphatidic acids | HMDB0114908 |
| PA 18:1_22:0 | Phosphatidic acid 18:1_22:0 | Phosphatidic acids | HMDB0114912 |
| PA 18:1_22:1 | Phosphatidic acid 18:1_22:1 | Phosphatidic acids | HMDB0114913 |
| PA 18:1_22:2 | Phosphatidic acid 18:1_22:2 | Phosphatidic acids | HMDB0114914 |
| PA 18:1_22:3 | Phosphatidic acid 18:1_22:3 | Phosphatidic acids | NA |
| PA 18:2_18:2 | Phosphatidic acid 18:2_18:2 | Phosphatidic acids | HMDB0114954 |
| PA 18:2_18:3 | Phosphatidic acid 18:2_18:3 | Phosphatidic acids | HMDB0114955 |
| PA 18:2_20:0 | Phosphatidic acid 18:2_20:0 | Phosphatidic acids | HMDB0114958 |
| PA 18:2_20:1 | Phosphatidic acid 18:2_20:1 | Phosphatidic acids | HMDB0114959 |
| PA 18:2_20:2 | Phosphatidic acid 18:2_20:2 | Phosphatidic acids | HMDB0115493 |
| PA 18:2_22:0 | Phosphatidic acid 18:2_22:0 | Phosphatidic acids | HMDB0114964 |
| PA 18:2_22:1 | Phosphatidic acid 18:2_22:1 | Phosphatidic acids | HMDB0114965 |
| PA 18:2_22:3 | Phosphatidic acid 18:2_22:3 | Phosphatidic acids | NA |
| PA 18:2_22:4 | Phosphatidic acid 18:2_22:4 | Phosphatidic acids | HMDB0114967 |
| PA 18:3_18:3 | Phosphatidic acid 18:3_18:3 | Phosphatidic acids | HMDB0114982 |
| PA 20:0_20:4 | Phosphatidic acid 20:0_20:4 | Phosphatidic acids | HMDB0115079 |
| LPG 14:0 | Lysophosphatidylglycerol 14:0 | Lysophosphatidylglycerols | NA |
| LPG 14:1 | Lysophosphatidylglycerol 14:1 | Lysophosphatidylglycerols | NA |

| | | | |
|---|---|---|---|
| LPG 16:0 | Lysophosphatidylglycerol 16:0 | Lysophosphatidylglycerols | HMDB0240601 |
| LPG 16:1 | Lysophosphatidylglycerol 16:1 | Lysophosphatidylglycerols | NA |
| LPG 17:1 | Lysophosphatidylglycerol 17:1 | Lysophosphatidylglycerols | NA |
| LPG 18:0 | Lysophosphatidylglycerol 18:0 | Lysophosphatidylglycerols | NA |
| LPG 18:1 | Lysophosphatidylglycerol 18:1 | Lysophosphatidylglycerols | HMDB0240602 |
| LPG 18:2 | Lysophosphatidylglycerol 18:2 | Lysophosphatidylglycerols | HMDB0240600 |
| LPG 20:1 | Lysophosphatidylglycerol 20:1 | Lysophosphatidylglycerols | NA |
| PE 20:0 | Phosphatidylethanolamine 20:0 | Phosphatidylethanolamines | NA |
| PE 28:0 | Phosphatidylethanolamine 28:0 | Phosphatidylethanolamines | HMDB0008821 |
| PE 30:0 | Phosphatidylethanolamine 30:0 | Phosphatidylethanolamines | HMDB0008889 |
| PE 30:1 | Phosphatidylethanolamine 30:1 | Phosphatidylethanolamines | HMDB0008857 |
| PE 32:0 | Phosphatidylethanolamine 32:0 | Phosphatidylethanolamines | HMDB0008923 |
| PE 32:1 | Phosphatidylethanolamine 32:1 | Phosphatidylethanolamines | HMDB0008859 |
| PE 32:2 | Phosphatidylethanolamine 32:2 | Phosphatidylethanolamines | HMDB0008957 |
| PE 33:0 | Phosphatidylethanolamine 33:0 | Phosphatidylethanolamines | HMDB0008988 |
| PE 33:1 | Phosphatidylethanolamine 33:1 | Phosphatidylethanolamines | HMDB0008894 |
| PE 33:2 | Phosphatidylethanolamine 33:2 | Phosphatidylethanolamines | HMDB0009087 |
| PE 34:0 | Phosphatidylethanolamine 34:0 | Phosphatidylethanolamines | HMDB0009217 |
| PE 34:1 | Phosphatidylethanolamine 34:1 | Phosphatidylethanolamines | HMDB0008927 |
| PE 34:2 | Phosphatidylethanolamine 34:2 | Phosphatidylethanolamines | HMDB0008928 |
| PE 34:3 | Phosphatidylethanolamine 34:3 | Phosphatidylethanolamines | HMDB0008930 |
| PE 34:4 | Phosphatidylethanolamine 34:4 | Phosphatidylethanolamines | HMDB0008870 |
| PE 35:1 | Phosphatidylethanolamine 35:1 | Phosphatidylethanolamines | HMDB0008900 |
| PE 35:2 | Phosphatidylethanolamine 35:2 | Phosphatidylethanolamines | HMDB0008901 |
| PE 35:3 | Phosphatidylethanolamine 35:3 | Phosphatidylethanolamines | HMDB0008903 |
| PE 36:0 | Phosphatidylethanolamine 36:0 | Phosphatidylethanolamines | HMDB0008991 |
| PE 36:1 | Phosphatidylethanolamine 36:1 | Phosphatidylethanolamines | HMDB0008993 |
| PE 36:2 | Phosphatidylethanolamine 36:2 | Phosphatidylethanolamines | HMDB0008994 |
| PE 36:3 | Phosphatidylethanolamine 36:3 | Phosphatidylethanolamines | HMDB0009060 |
| PE 36:4 | Phosphatidylethanolamine 36:4 | Phosphatidylethanolamines | HMDB0008937 |
| PE 36:5 | Phosphatidylethanolamine 36:5 | Phosphatidylethanolamines | HMDB0008877 |
| PE 36:6 | Phosphatidylethanolamine 36:6 | Phosphatidylethanolamines | HMDB0008972 |
| PE 38:0 | Phosphatidylethanolamine 38:0 | Phosphatidylethanolamines | HMDB0008998 |
| PE 38:1 | Phosphatidylethanolamine 38:1 | Phosphatidylethanolamines | HMDB0009224 |
| PE 38:2 | Phosphatidylethanolamine 38:2 | Phosphatidylethanolamines | HMDB0009000 |
| PE 38:3 | Phosphatidylethanolamine 38:3 | Phosphatidylethanolamines | HMDB0008975 |
| PE 38:4 | Phosphatidylethanolamine 38:4 | Phosphatidylethanolamines | HMDB0009228 |
| PE 38:5 | Phosphatidylethanolamine 38:5 | Phosphatidylethanolamines | HMDB0008976 |
| PE 38:6 | Phosphatidylethanolamine 38:6 | Phosphatidylethanolamines | HMDB0008946 |
| PE 38:7 | Phosphatidylethanolamine 38:7 | Phosphatidylethanolamines | HMDB0009104 |
| PE 40:1 | Phosphatidylethanolamine 40:1 | Phosphatidylethanolamines | HMDB0009007 |

| | | | |
|---|---|---|---|
| PE 40:3 | Phosphatidylethanolamine 40:3 | Phosphatidylethanolamines | HMDB0009074 |
| PE 40:4 | Phosphatidylethanolamine 40:4 | Phosphatidylethanolamines | HMDB0009107 |
| PE 40:5 | Phosphatidylethanolamine 40:5 | Phosphatidylethanolamines | HMDB0009075 |
| PE 40:6 | Phosphatidylethanolamine 40:6 | Phosphatidylethanolamines | HMDB0009012 |
| PE 40:7 | Phosphatidylethanolamine 40:7 | Phosphatidylethanolamines | HMDB0009141 |
| PE 40:8 | Phosphatidylethanolamine 40:8 | Phosphatidylethanolamines | HMDB0009207 |
| PE 42:7 | Phosphatidylethanolamine 42:7 | Phosphatidylethanolamines | HMDB0009276 |
| PE 42:8 | Phosphatidylethanolamine 42:8 | Phosphatidylethanolamines | HMDB0009309 |
| PE 44:6 | Phosphatidylethanolamine 44:6 | Phosphatidylethanolamines | HMDB0009507 |
| PE 44:7 | Phosphatidylethanolamine 44:7 | Phosphatidylethanolamines | HMDB0009700 |
| PE 44:11 | Phosphatidylethanolamine 44:11 | Phosphatidylethanolamines | HMDB0009639 |
| PE 44:12 | Phosphatidylethanolamine 44:12 | Phosphatidylethanolamines | HMDB0009705 |
| PE P-16:0/16:0 | Phosphatidylethanolamine P-16:0/16:0 | Phosphatidylethanolamines | HMDB0011158 |
| PE P-16:0/18:1 | Phosphatidylethanolamine P-16:0/18:1 | Phosphatidylethanolamines | HMDB0011342 |
| PE P-16:0/18:2 | Phosphatidylethanolamine P-16:0/18:2 | Phosphatidylethanolamines | HMDB0011343 |
| PE P-16:0/18:3 | Phosphatidylethanolamine P-16:0/18:3 | Phosphatidylethanolamines | HMDB0009180 |
| PE P-16:0/20:3 | Phosphatidylethanolamine P-16:0/20:3 | Phosphatidylethanolamines | HMDB0011351 |
| PE P-16:0/20:4 | Phosphatidylethanolamine P-16:0/20:4 | Phosphatidylethanolamines | HMDB0011352 |
| PE P-16:0/20:5 | Phosphatidylethanolamine P-16:0/20:5 | Phosphatidylethanolamines | HMDB0011354 |
| PE P-16:0/22:4 | Phosphatidylethanolamine P-16:0/22:4 | Phosphatidylethanolamines | HMDB0011358 |
| PE P-16:0/22:5 | Phosphatidylethanolamine P-16:0/22:5 | Phosphatidylethanolamines | HMDB0009675 |
| PE P-16:0/22:6 | Phosphatidylethanolamine P-16:0/22:6 | Phosphatidylethanolamines | HMDB0009708 |
| PE P-18:0/14:0 | Phosphatidylethanolamine P-18:0/14:0 | Phosphatidylethanolamines | HMDB0011368 |
| PE P-18:0/16:0 | Phosphatidylethanolamine P-18:0/16:0 | Phosphatidylethanolamines | HMDB0011371 |
| PE P-18:0/16:1 | Phosphatidylethanolamine P-18:0/16:1 | Phosphatidylethanolamines | HMDB0011372 |
| PE P-18:0/18:1 | Phosphatidylethanolamine P-18:0/18:1 | Phosphatidylethanolamines | HMDB0009049 |
| PE P-18:0/18:2 | Phosphatidylethanolamine P-18:0/18:2 | Phosphatidylethanolamines | HMDB0011376 |
| PE P-18:0/18:3 | Phosphatidylethanolamine P-18:0/18:3 | Phosphatidylethanolamines | HMDB0011377 |
| PE P-18:0/20:1 | Phosphatidylethanolamine P-18:0/20:1 | Phosphatidylethanolamines | HMDB0011381 |
| PE P-18:0/20:2 | Phosphatidylethanolamine P-18:0/20:2 | Phosphatidylethanolamines | HMDB0011382 |
| PE P-18:0/20:3 | Phosphatidylethanolamine P-18:0/20:3 | Phosphatidylethanolamines | HMDB0011384 |
| PE P-18:0/20:4 | Phosphatidylethanolamine P-18:0/20:4 | Phosphatidylethanolamines | HMDB0005779 |
| PE P-18:0/20:5 | Phosphatidylethanolamine P-18:0/20:5 | Phosphatidylethanolamines | HMDB0011387 |
| PE P-18:0/22:3 | Phosphatidylethanolamine P-18:0/22:3 | Phosphatidylethanolamines | NA |
| PE P-18:0/22:4 | Phosphatidylethanolamine P-18:0/22:4 | Phosphatidylethanolamines | HMDB0011391 |
| PE P-18:0/22:5 | Phosphatidylethanolamine P-18:0/22:5 | Phosphatidylethanolamines | HMDB0011392 |
| PE P-18:0/22:6 | Phosphatidylethanolamine P-18:0/22:6 | Phosphatidylethanolamines | HMDB0009709 |
| PE P-18:1/18:1 | Phosphatidylethanolamine P-18:1/18:1 | Phosphatidylethanolamines | HMDB0009084 |
| PE P-18:1/18:2 | Phosphatidylethanolamine P-18:1/18:2 | Phosphatidylethanolamines | HMDB0011409 |
| PE P-18:1/20:4 | Phosphatidylethanolamine P-18:1/20:4 | Phosphatidylethanolamines | HMDB0011452 |
| PE P-18:1/20:5 | Phosphatidylethanolamine P-18:1/20:5 | Phosphatidylethanolamines | HMDB0011453 |

| | | | |
|---|---|---|---|
| PE P-18:1/22:6 | Phosphatidylethanolamine P-18:1/22:6 | Phosphatidylethanolamines | HMDB0011460 |
| PE P-20:0/18:2 | Phosphatidylethanolamine P-20:0/18:2 | Phosphatidylethanolamines | NA |
| PE P-20:0/20:4 | Phosphatidylethanolamine P-20:0/20:4 | Phosphatidylethanolamines | NA |
| PE P-20:0/20:5 | Phosphatidylethanolamine P-20:0/20:5 | Phosphatidylethanolamines | NA |
| PI 14:0_18:1 | Phosphatidyl-inositol 14:0_18:1 | Phosphatidylinositols | NA |
| PI 14:0_18:2 | Phosphatidyl-inositol 14:0_18:2 | Phosphatidylinositols | NA |
| PI 15:0_16:0 | Phosphatidyl-inositol 15:0_16:0 | Phosphatidylinositols | NA |
| PI 15:1_16:0 | Phosphatidyl-inositol 15:1_16:0 | Phosphatidylinositols | NA |
| PI 16:0_16:0 | Phosphatidyl-inositol 16:0_16:0 | Phosphatidylinositols | HMDB0009778 |
| PI 16:0_17:0 | Phosphatidyl-inositol 16:0_17:0 | Phosphatidylinositols | NA |
| PI 16:0_17:1 | Phosphatidyl-inositol 16:0_17:1 | Phosphatidylinositols | NA |
| PI 16:0_17:2 | Phosphatidyl-inositol 16:0_17:2 | Phosphatidylinositols | NA |
| PI 16:0_18:1 | Phosphatidyl-inositol 16:0_18:1 | Phosphatidylinositols | HMDB0009782 |
| PI 16:0_18:2 | Phosphatidyl-inositol 16:0_18:2 | Phosphatidylinositols | HMDB0009784 |
| PI 16:0_18:3 | Phosphatidyl-inositol 16:0_18:3 | Phosphatidylinositols | NA |
| PI 16:0_20:0 | Phosphatidyl-inositol 16:0_20:0 | Phosphatidylinositols | HMDB0009785 |
| PI 16:0_20:3 | Phosphatidyl-inositol 16:0_20:3 | Phosphatidylinositols | HMDB0009886 |
| PI 16:0_20:4 | Phosphatidyl-inositol 16:0_20:4 | Phosphatidylinositols | HMDB0009893 |
| PI 16:0_22:1 | Phosphatidyl-inositol 16:0_22:1 | Phosphatidylinositols | NA |
| PI 16:1_18:0 | Phosphatidyl-inositol 16:1_18:0 | Phosphatidylinositols | HMDB0009799 |
| PI 16:1_18:1 | Phosphatidyl-inositol 16:1_18:1 | Phosphatidylinositols | HMDB0009835 |
| PI 16:1_18:2 | Phosphatidyl-inositol 16:1_18:2 | Phosphatidylinositols | NA |
| PI 17:0_18:1 | Phosphatidyl-inositol 17:0_18:1 | Phosphatidylinositols | NA |
| PI 17:1_18:1 | Phosphatidyl-inositol 17:1_18:1 | Phosphatidylinositols | NA |
| PI 17:1_18:2 | Phosphatidyl-inositol 17:1_18:2 | Phosphatidylinositols | NA |
| PI 18:0_18:0 | Phosphatidyl-inositol 18:0_18:0 | Phosphatidylinositols | HMDB0009808 |
| PI 18:0_18:1 | Phosphatidyl-inositol 18:0_18:1 | Phosphatidylinositols | HMDB0240667 |
| PI 18:0_18:2 | Phosphatidyl-inositol 18:0_18:2 | Phosphatidylinositols | HMDB0009847 |
| PI 18:0_18:3 | Phosphatidyl-inositol 18:0_18:3 | Phosphatidylinositols | HMDB0009810 |
| PI 18:0_20:0 | Phosphatidyl-inositol 18:0_20:0 | Phosphatidylinositols | NA |
| PI 18:0_20:3 | Phosphatidyl-inositol 18:0_20:3 | Phosphatidylinositols | HMDB0009887 |
| PI 18:0_20:4 | Phosphatidyl-inositol 18:0_20:4 | Phosphatidylinositols | HMDB0009815 |
| PI 18:0_22:0 | Phosphatidyl-inositol 18:0_22:0 | Phosphatidylinositols | NA |
| PI 18:1_18:1 | Phosphatidyl-inositol 18:1_18:1 | Phosphatidylinositols | HMDB0009824 |
| PI 18:1_18:2 | Phosphatidyl-inositol 18:1_18:2 | Phosphatidylinositols | HMDB0009849 |
| PI 18:1_18:3 | Phosphatidyl-inositol 18:1_18:3 | Phosphatidylinositols | HMDB0009839 |
| PI 18:1_20:0 | Phosphatidyl-inositol 18:1_20:0 | Phosphatidylinositols | NA |
| PI 18:1_20:1 | Phosphatidyl-inositol 18:1_20:1 | Phosphatidylinositols | HMDB0009871 |
| PI 18:1_20:2 | Phosphatidyl-inositol 18:1_20:2 | Phosphatidylinositols | NA |
| PI 18:1_20:3 | Phosphatidyl-inositol 18:1_20:3 | Phosphatidylinositols | HMDB0009843 |
| PI 18:1_20:4 | Phosphatidyl-inositol 18:1_20:4 | Phosphatidylinositols | HMDB0009896 |

| | | | |
|---|---|---|---|
| PI 18:1_20:5 | Phosphatidyl-inositol 18:1_20:5 | Phosphatidylinositols | NA |
| PI 18:1_22:0 | Phosphatidyl-inositol 18:1_22:0 | Phosphatidylinositols | NA |
| PI 18:1_22:1 | Phosphatidyl-inositol 18:1_22:1 | Phosphatidylinositols | NA |
| PI 18:1_22:2 | Phosphatidyl-inositol 18:1_22:2 | Phosphatidylinositols | NA |
| PI 18:1_22:3 | Phosphatidyl-inositol 18:1_22:3 | Phosphatidylinositols | NA |
| PI 18:1_22:4 | Phosphatidyl-inositol 18:1_22:4 | Phosphatidylinositols | NA |
| PI 18:1_22:5 | Phosphatidyl-inositol 18:1_22:5 | Phosphatidylinositols | NA |
| PI 18:1_22:6 | Phosphatidyl-inositol 18:1_22:6 | Phosphatidylinositols | NA |
| PI 18:2_18:3 | Phosphatidyl-inositol 18:2_18:3 | Phosphatidylinositols | NA |
| PI 18:2_20:0 | Phosphatidyl-inositol 18:2_20:0 | Phosphatidylinositols | HMDB0009851 |
| PI 18:2_20:1 | Phosphatidyl-inositol 18:2_20:1 | Phosphatidylinositols | HMDB0009852 |
| PI 18:2_20:4 | Phosphatidyl-inositol 18:2_20:4 | Phosphatidylinositols | NA |
| PI 18:2_20:5 | Phosphatidyl-inositol 18:2_20:5 | Phosphatidylinositols | NA |
| PI 18:2_22:0 | Phosphatidyl-inositol 18:2_22:0 | Phosphatidylinositols | NA |
| PI 18:2_22:1 | Phosphatidyl-inositol 18:2_22:1 | Phosphatidylinositols | NA |
| PI 18:2_22:6 | Phosphatidyl-inositol 18:2_22:6 | Phosphatidylinositols | NA |
| PS 32:0 | Phosphatidyl-serine 32:0 | Phosphatidylserines | HMDB0000614 |
| PS 34:1 | Phosphatidyl-serine 34:1 | Phosphatidylserines | HMDB0112267 |
| PS 34:2 | Phosphatidyl-serine 34:2 | Phosphatidylserines | HMDB0112296 |
| PS 36:1 | Phosphatidyl-serine 36:1 | Phosphatidylserines | HMDB0010163 |
| PS 36:2 | Phosphatidyl-serine 36:2 | Phosphatidylserines | HMDB0012390 |
| PS 36:3 | Phosphatidyl-serine 36:3 | Phosphatidylserines | HMDB0112315 |
| PS 36:4 | Phosphatidyl-serine 36:4 | Phosphatidylserines | HMDB0012402 |
| PS 36:5 | Phosphatidyl-serine 36:5 | Phosphatidylserines | HMDB0112316 |
| PS 38:4 | Phosphatidyl-serine 38:4 | Phosphatidylserines | HMDB0012383 |
| PS 38:5 | Phosphatidyl-serine 38:5 | Phosphatidylserines | HMDB0112367 |
| PS 38:6 | Phosphatidyl-serine 38:6 | Phosphatidylserines | HMDB0012362 |
| PS 38:7 | Phosphatidyl-serine 38:7 | Phosphatidylserines | HMDB0012373 |
| PS 40:4 | Phosphatidyl-serine 40:4 | Phosphatidylserines | HMDB0112437 |
| PS 40:5 | Phosphatidyl-serine 40:5 | Phosphatidylserines | HMDB0112420 |
| PS 40:6 | Phosphatidyl-serine 40:6 | Phosphatidylserines | HMDB0010167 |
| PS 40:7 | Phosphatidyl-serine 40:7 | Phosphatidylserines | HMDB0012395 |
| PS 40:8 | Phosphatidyl-serine 40:8 | Phosphatidylserines | HMDB0112506 |
| PC 24:0 | Phosphatidylcholine 24:0 | Phosphatidylcholines | NA |
| PC 28:1 | Phosphatidylcholine 28:1 | Phosphatidylcholines | HMDB0007867 |
| PC 30:0 | Phosphatidylcholine 30:0 | Phosphatidylcholines | HMDB0007934 |
| PC 32:0 | Phosphatidylcholine 32:0 | Phosphatidylcholines | HMDB00564 |
| PC 32:1 | Phosphatidylcholine 32:1 | Phosphatidylcholines | HMDB0007872 |
| PC 32:2 | Phosphatidylcholine 32:2 | Phosphatidylcholines | HMDB0008002 |
| PC 32:3 | Phosphatidylcholine 32:3 | Phosphatidylcholines | HMDB0007876 |
| PC 34:1 | Phosphatidylcholine 34:1 | Phosphatidylcholines | HMDB0007971 |

| | | | |
|---|---|---|---|
| PC 34:2 | Phosphatidylcholine 34:2 | Phosphatidylcholines | HMDB07973 |
| PC 34:3 | Phosphatidylcholine 34:3 | Phosphatidylcholines | HMDB08006 |
| PC 34:4 | Phosphatidylcholine 34:4 | Phosphatidylcholines | HMDB0007883 |
| PC 36:0 | Phosphatidylcholine 36:0 | Phosphatidylcholines | HMDB0008265 |
| PC 36:1 | Phosphatidylcholine 36:1 | Phosphatidylcholines | HMDB0008037 |
| PC 36:2 | Phosphatidylcholine 36:2 | Phosphatidylcholines | HMDB0008039 |
| PC 36:3 | Phosphatidylcholine 36:3 | Phosphatidylcholines | HMDB0007980 |
| PC 36:4 | Phosphatidylcholine 36:4 | Phosphatidylcholines | HMDB0008042 |
| PC 36:5 | Phosphatidylcholine 36:5 | Phosphatidylcholines | HMDB0007984 |
| PC 36:6 | Phosphatidylcholine 36:6 | Phosphatidylcholines | HMDB0007892 |
| PC 38:0 | Phosphatidylcholine 38:0 | Phosphatidylcholines | HMDB0007893 |
| PC 38:3 | Phosphatidylcholine 38:3 | Phosphatidylcholines | HMDB0008046 |
| PC 38:4 | Phosphatidylcholine 38:4 | Phosphatidylcholines | HMDB0007988 |
| PC 38:5 | Phosphatidylcholine 38:5 | Phosphatidylcholines | HMDB0007989 |
| PC 38:6 | Phosphatidylcholine 38:6 | Phosphatidylcholines | HMDB0008083 |
| PC 40:2 | Phosphatidylcholine 40:2 | Phosphatidylcholines | HMDB0008276 |
| PC 40:3 | Phosphatidylcholine 40:3 | Phosphatidylcholines | HMDB0008119 |
| PC 40:4 | Phosphatidylcholine 40:4 | Phosphatidylcholines | HMDB0008054 |
| PC 40:5 | Phosphatidylcholine 40:5 | Phosphatidylcholines | HMDB0008055 |
| PC 40:6 | Phosphatidylcholine 40:6 | Phosphatidylcholines | HMDB0008057 |
| PC 42:0 | Phosphatidylcholine 42:0 | Phosphatidylcholines | HMDB0008537 |
| PC 42:1 | Phosphatidylcholine 42:1 | Phosphatidylcholines | HMDB0008059 |
| PC 42:2 | Phosphatidylcholine 42:2 | Phosphatidylcholines | HMDB0008157 |
| PC 42:4 | Phosphatidylcholine 42:4 | Phosphatidylcholines | HMDB0008285 |
| PC 42:5 | Phosphatidylcholine 42:5 | Phosphatidylcholines | HMDB0008287 |
| PC 42:6 | Phosphatidylcholine 42:6 | Phosphatidylcholines | HMDB0008288 |
| PC O-28:0 | Phosphatidylcholine O-28:0 | Phosphatidylcholines | NA |
| PC O-28:1 | Phosphatidylcholine O-28:1 | Phosphatidylcholines | NA |
| PC O-30:0 | Phosphatidylcholine O-30:0 | Phosphatidylcholines | HMDB0013341 |
| PC O-30:1 | Phosphatidylcholine O-30:1 | Phosphatidylcholines | HMDB0013402 |
| PC O-30:2 | Phosphatidylcholine O-30:2 | Phosphatidylcholines | HMDB0013410 |
| PC O-32:1 | Phosphatidylcholine O-32:1 | Phosphatidylcholines | HMDB0013404 |
| PC O-32:2 | Phosphatidylcholine O-32:2 | Phosphatidylcholines | HMDB0013411 |
| PC O-34:0 | Phosphatidylcholine O-34:0 | Phosphatidylcholines | HMDB0013405 |
| PC O-34:1 | Phosphatidylcholine O-34:1 | Phosphatidylcholines | HMDB0013426 |
| PC O-34:2 | Phosphatidylcholine O-34:2 | Phosphatidylcholines | HMDB0011151 |
| PC O-34:3 | Phosphatidylcholine O-34:3 | Phosphatidylcholines | HMDB0013413 |
| PC O-36:0 | Phosphatidylcholine O-36:0 | Phosphatidylcholines | HMDB13406 |
| PC O-36:1 | Phosphatidylcholine O-36:1 | Phosphatidylcholines | HMDB0013427 |
| PC O-36:2 | Phosphatidylcholine O-36:2 | Phosphatidylcholines | HMDB0013418 |
| PC O-36:3 | Phosphatidylcholine O-36:3 | Phosphatidylcholines | HMDB0013429 |

| | | | |
|---|---|---|---|
| PC O-36:4 | Phosphatidylcholine O-36:4 | Phosphatidylcholines | HMDB0013435 |
| PC O-36:5 | Phosphatidylcholine O-36:5 | Phosphatidylcholines | HMDB0013415 |
| PC O-38:0 | Phosphatidylcholine O-38:0 | Phosphatidylcholines | HMDB0013408 |
| PC O-38:1 | Phosphatidylcholine O-38:1 | Phosphatidylcholines | HMDB0013416 |
| PC O-38:2 | Phosphatidylcholine O-38:2 | Phosphatidylcholines | HMDB0013431 |
| PC O-38:3 | Phosphatidylcholine O-38:3 | Phosphatidylcholines | HMDB0013439 |
| PC O-38:4 | Phosphatidylcholine O-38:4 | Phosphatidylcholines | HMDB0013420 |
| PC O-38:5 | Phosphatidylcholine O-38:5 | Phosphatidylcholines | HMDB11253 |
| PC O-38:6 | Phosphatidylcholine O-38:6 | Phosphatidylcholines | HMDB0013409 |
| PC O-40:1 | Phosphatidylcholine O-40:1 | Phosphatidylcholines | HMDB0013433 |
| PC O-40:2 | Phosphatidylcholine O-40:2 | Phosphatidylcholines | HMDB0013437 |
| PC O-40:3 | Phosphatidylcholine O-40:3 | Phosphatidylcholines | HMDB0013445 |
| PC O-40:4 | Phosphatidylcholine O-40:4 | Phosphatidylcholines | HMDB0013442 |
| PC O-40:5 | Phosphatidylcholine O-40:5 | Phosphatidylcholines | HMDB0013444 |
| PC O-40:6 | Phosphatidylcholine O-40:6 | Phosphatidylcholines | HMDB0013422 |
| PC O-42:1 | Phosphatidylcholine O-42:1 | Phosphatidylcholines | HMDB0013434 |
| PC O-42:2 | Phosphatidylcholine O-42:2 | Phosphatidylcholines | HMDB0013438 |
| PC O-42:3 | Phosphatidylcholine O-42:3 | Phosphatidylcholines | HMDB0013458 |
| PC O-42:4 | Phosphatidylcholine O-42:4 | Phosphatidylcholines | HMDB0013448 |
| PC O-42:5 | Phosphatidylcholine O-42:5 | Phosphatidylcholines | HMDB0013451 |
| PC O-44:3 | Phosphatidylcholine O-44:3 | Phosphatidylcholines | HMDB0013449 |
| PC O-44:4 | Phosphatidylcholine O-44:4 | Phosphatidylcholines | HMDB0013455 |
| PC O-44:5 | Phosphatidylcholine O-44:5 | Phosphatidylcholines | HMDB0013456 |
| PC O-44:6 | Phosphatidylcholine O-44:6 | Phosphatidylcholines | HMDB0013450 |
| PG 14:0_16:0 | Phosphatidylglycerol 14:0_16:0 | Phosphatidylglycerols | NA |
| PG 15:0_18:1 | Phosphatidylglycerol 15:0_18:1 | Phosphatidylglycerols | NA |
| PG 16:0_16:0 | Phosphatidylglycerol 16:0_16:0 | Phosphatidylglycerols | HMDB0010570 |
| PG 16:0_16:1 | Phosphatidylglycerol 16:0_16:1 | Phosphatidylglycerols | HMDB0010585 |
| PG 16:0_18:1 | Phosphatidylglycerol 16:0_18:1 | Phosphatidylglycerols | HMDB10574 |
| PG 16:0_18:2 | Phosphatidylglycerol 16:0_18:2 | Phosphatidylglycerols | HMDB0010645 |
| PG 16:0_18:3 | Phosphatidylglycerol 16:0_18:3 | Phosphatidylglycerols | HMDB0010660 |
| PG 16:0_19:1 | Phosphatidylglycerol 16:0_19:1 | Phosphatidylglycerols | NA |
| PG 16:0_20:3 | Phosphatidylglycerol 16:0_20:3 | Phosphatidylglycerols | HMDB0010579 |
| PG 16:0_20:4 | Phosphatidylglycerol 16:0_20:4 | Phosphatidylglycerols | HMDB0010580 |
| PG 16:0_20:5 | Phosphatidylglycerol 16:0_20:5 | Phosphatidylglycerols | NA |
| PG 16:0_22:1 | Phosphatidylglycerol 16:0_22:1 | Phosphatidylglycerols | NA |
| PG 16:0_22:2 | Phosphatidylglycerol 16:0_22:2 | Phosphatidylglycerols | NA |
| PG 16:1_16:1 | Phosphatidylglycerol 16:1_16:1 | Phosphatidylglycerols | HMDB0010586 |
| PG 16:1_18:0 | Phosphatidylglycerol 16:1_18:0 | Phosphatidylglycerols | HMDB0010587 |
| PG 16:1_18:1 | Phosphatidylglycerol 16:1_18:1 | Phosphatidylglycerols | HMDB0010631 |
| PG 16:1_18:2 | Phosphatidylglycerol 16:1_18:2 | Phosphatidylglycerols | HMDB0010646 |

| | | | |
|---|---|---|---|
| PG 16:1_20:4 | Phosphatidylglycerol 16:1_20:4 | Phosphatidylglycerols | HMDB0116608 |
| PG 16:1_22:1 | Phosphatidylglycerol 16:1_22:1 | Phosphatidylglycerols | NA |
| PG 16:2_18:1 | Phosphatidylglycerol 16:2_18:1 | Phosphatidylglycerols | NA |
| PG 16:2_18:2 | Phosphatidylglycerol 16:2_18:2 | Phosphatidylglycerols | NA |
| PG 16:3_18:1 | Phosphatidylglycerol 16:3_18:1 | Phosphatidylglycerols | NA |
| PG 17:0_18:1 | Phosphatidylglycerol 17:0_18:1 | Phosphatidylglycerols | NA |
| PG 17:0_18:2 | Phosphatidylglycerol 17:0_18:2 | Phosphatidylglycerols | NA |
| PG 17:1_18:1 | Phosphatidylglycerol 17:1_18:1 | Phosphatidylglycerols | NA |
| PG 18:0_18:1 | Phosphatidylglycerol 18:0_18:1 | Phosphatidylglycerols | HMDB0010617 |
| PG 18:0_18:2 | Phosphatidylglycerol 18:0_18:2 | Phosphatidylglycerols | HMDB0010647 |
| PG 18:0_18:3 | Phosphatidylglycerol 18:0_18:3 | Phosphatidylglycerols | HMDB0010606 |
| PG 18:0_22:1 | Phosphatidylglycerol 18:0_22:1 | Phosphatidylglycerols | NA |
| PG 18:1_18:1 | Phosphatidylglycerol 18:1_18:1 | Phosphatidylglycerols | HMDB0010634 |
| PG 18:1_18:2 | Phosphatidylglycerol 18:1_18:2 | Phosphatidylglycerols | HMDB0010649 |
| PG 18:1_18:3 | Phosphatidylglycerol 18:1_18:3 | Phosphatidylglycerols | HMDB0010636 |
| PG 18:1_20:0 | Phosphatidylglycerol 18:1_20:0 | Phosphatidylglycerols | NA |
| PG 18:1_20:1 | Phosphatidylglycerol 18:1_20:1 | Phosphatidylglycerols | NA |
| PG 18:1_20:2 | Phosphatidylglycerol 18:1_20:2 | Phosphatidylglycerols | NA |
| PG 18:1_20:3 | Phosphatidylglycerol 18:1_20:3 | Phosphatidylglycerols | HMDB0010639 |
| PG 18:1_20:4 | Phosphatidylglycerol 18:1_20:4 | Phosphatidylglycerols | HMDB0116610 |
| PG 18:1_20:5 | Phosphatidylglycerol 18:1_20:5 | Phosphatidylglycerols | NA |
| PG 18:1_22:0 | Phosphatidylglycerol 18:1_22:0 | Phosphatidylglycerols | NA |
| PG 18:1_22:1 | Phosphatidylglycerol 18:1_22:1 | Phosphatidylglycerols | NA |
| PG 18:1_22:2 | Phosphatidylglycerol 18:1_22:2 | Phosphatidylglycerols | NA |
| PG 18:1_22:3 | Phosphatidylglycerol 18:1_22:3 | Phosphatidylglycerols | NA |
| PG 18:1_22:4 | Phosphatidylglycerol 18:1_22:4 | Phosphatidylglycerols | HMDB0010626 |
| PG 18:1_22:5 | Phosphatidylglycerol 18:1_22:5 | Phosphatidylglycerols | HMDB0116616 |
| PG 18:2_18:2 | Phosphatidylglycerol 18:2_18:2 | Phosphatidylglycerols | HMDB0010650 |
| PG 18:2_18:3 | Phosphatidylglycerol 18:2_18:3 | Phosphatidylglycerols | HMDB0010651 |
| PG 18:2_18:4 | Phosphatidylglycerol 18:2_18:4 | Phosphatidylglycerols | NA |
| PG 18:2_20:0 | Phosphatidylglycerol 18:2_20:0 | Phosphatidylglycerols | NA |
| PG 18:2_20:3 | Phosphatidylglycerol 18:2_20:3 | Phosphatidylglycerols | HMDB0010654 |
| PG 18:2_20:4 | Phosphatidylglycerol 18:2_20:4 | Phosphatidylglycerols | HMDB0116584 |
| PG 18:2_20:5 | Phosphatidylglycerol 18:2_20:5 | Phosphatidylglycerols | NA |
| PG 18:2_22:0 | Phosphatidylglycerol 18:2_22:0 | Phosphatidylglycerols | NA |
| PG 18:2_22:1 | Phosphatidylglycerol 18:2_22:1 | Phosphatidylglycerols | NA |
| PG 18:2_22:3 | Phosphatidylglycerol 18:2_22:3 | Phosphatidylglycerols | NA |
| PG 18:2_22:4 | Phosphatidylglycerol 18:2_22:4 | Phosphatidylglycerols | HMDB0010656 |
| PG 20:3_20:4 | Phosphatidylglycerol 20:3_20:4 | Phosphatidylglycerols | HMDB0116588 |
| PG 20:4_20:4 | Phosphatidylglycerol 20:4_20:4 | Phosphatidylglycerols | HMDB0116596 |
| PG 20:4_22:1 | Phosphatidylglycerol 20:4_22:1 | Phosphatidylglycerols | NA |

| | | | |
|---|---|---|---|
| PG 20:4_22:3 | Phosphatidylglycerol 20:4_22:3 | Phosphatidylglycerols | NA |
| PG 20:4_22:4 | Phosphatidylglycerol 20:4_22:4 | Phosphatidylglycerols | NA |
| PG 22:4_22:6 | Phosphatidylglycerol 22:4_22:6 | Phosphatidylglycerols | NA |
| PG 22:5_22:6 | Phosphatidylglycerol 22:5_22:6 | Phosphatidylglycerols | HMDB0116635 |
| PG 22:6_22:6 | Phosphatidylglycerol 22:6_22:6 | Phosphatidylglycerols | HMDB0116605 |
| N-Ac-Putrescine | N-Acetylputrescine | Polyamines | HMDB0002064 |
| N1-Ac-Spermidine | N1-Acetylspermidine | Polyamines | HMDB0001276 |
| N1,N8-Di-Ac-Spermidine | N1,N8-Diacetylspermidine | Polyamines | HMDB0041947 |
| N1,N12-Di-Ac-Spermine | N1,N12-Diacetylspermine | Polyamines | HMDB0002172 |
| N8-Ac-Spermidine | N8-Acetylspermidine | Polyamines | HMDB0002189 |
| Isonicotinic acid | Isonicotinic acid | Pyridinecarboxylic acids | HMDB0060665 |
| Kynurenic acid | Kynurenic acid | Pyridinecarboxylic acids | HMDB0000715 |
| Picolinic acid | Picolinic acid | Pyridinecarboxylic acids | HMDB0002243 |
| Quinaldic acid | Quinaldic acid | Pyridinecarboxylic acids | HMDB0000842 |
| Quinolinic acid | Quinolinic acid | Pyridinecarboxylic acids | HMDB0000232 |
| Xanthurenic acid | Xanthurenic acid | Pyridinecarboxylic acids | HMDB0000881 |
| SPBP d14:0 | C14 Sphinganine-1-phosphate | Sphinganine and sphingosine phosphates | NA |
| SPBP d14:1 | C14 Sphingosine-1-phosphate | Sphinganine and sphingosine phosphates | NA |
| SPBP d16:1 | C16 Sphingosine-1-phosphate | Sphinganine and sphingosine phosphates | HMDB0060061 |
| SPBP d17:0 | C17 Sphinganine-1-phosphate | Sphinganine and sphingosine phosphates | NA |
| SPBP d17:1 | C17 Sphingosine-1-phosphate | Sphinganine and sphingosine phosphates | NA |
| SPBP d18:1 | C18 Sphingosine-1-phosphate | Sphinganine and sphingosine phosphates | HMDB0000277 |
| SPB d14:0 | C14 Sphinganine | Sphinganines and sphingosines | NA |
| SPB d16:0 | C16 Sphinganine | Sphinganines and sphingosines | NA |
| SPB d17:0 | C17 Sphinganine | Sphinganines and sphingosines | NA |
| SPB d18:0 | C18 Sphinganine | Sphinganines and sphingosines | HMDB0000269 |
| SPB d18:1 | C18 Sphingosine | Sphinganines and sphingosines | HMDB0000252 |
| SM 33:1 | Sphingomyelin 33:1 | Sphingomyelins | HMDB0240608 |
| SM 34:1 | Sphingomyelin 34:1 | Sphingomyelins | HMDB0061712 |
| SM 34:2 | Sphingomyelin 34:2 | Sphingomyelins | NA |
| SM 35:1 | Sphingomyelin 35:1 | Sphingomyelins | HMDB0240609 |
| SM 36:1 | Sphingomyelin 36:1 | Sphingomyelins | HMDB01348 |
| SM 36:2 | Sphingomyelin 36:2 | Sphingomyelins | HMDB0012100 |
| SM 38:3 | Sphingomyelin 38:3 | Sphingomyelins | NA |
| SM 41:1 | Sphingomyelin 41:1 | Sphingomyelins | HMDB0012105 |
| SM 41:2 | Sphingomyelin 41:2 | Sphingomyelins | HMDB0240614 |
| SM 42:1 | Sphingomyelin 42:1 | Sphingomyelins | HMDB0011697 |
| SM 42:2 | Sphingomyelin 42:2 | Sphingomyelins | HMDB0012107 |
| SM 43:1 | Sphingomyelin 43:1 | Sphingomyelins | HMDB0240671 |
| SM 44:1 | Sphingomyelin 44:1 | Sphingomyelins | HMDB0011698 |
| SM 44:2 | Sphingomyelin 44:2 | Sphingomyelins | HMDB0013461 |

| | | | |
|---|---|---|---|
| 3-Deoxyglucosone | 3-Deoxyglucosone | Sugars | HMDB0245856 |
| Fructose | Fructose | Sugars | HMDB0000660 |
| Glucose | Glucose | Sugars | HMDB0000122 |
| Mannose | Mannose | Sugars | HMDB0000169 |
| NeuAc | Acetylneuraminic acid | Sugars | HMDB0000230 |
| Ribose | Ribose | Sugars | HMDB0000283 |
| Threonic acid | Threonic acid | Sugars | HMDB0000943 |
| AconAcid | Aconitic acid | Tricarboxylic acids | HMDB0000958 |
| Citric acid | Citric acid | Tricarboxylic acids | HMDB0000094 |
| Isocitric acid | Isocitric acid | Tricarboxylic acids | HMDB0000193 |
| TG 14:0_32:2 | Triacylglyceride 14:0_32:2 | Triglycerides | HMDB0042281 |
| TG 14:0_34:1 | Triacylglyceride 14:0_34:1 | Triglycerides | HMDB0042275 |
| TG 14:0_34:2 | Triacylglyceride 14:0_34:2 | Triglycerides | HMDB0042282 |
| TG 14:0_34:3 | Triacylglyceride 14:0_34:3 | Triglycerides | HMDB0042316 |
| TG 14:0_35:1 | Triacylglyceride 14:0_35:1 | Triglycerides | HMDB0042102 |
| TG 14:0_35:2 | Triacylglyceride 14:0_35:2 | Triglycerides | HMDB0042108 |
| TG 14:0_36:2 | Triacylglyceride 14:0_36:2 | Triglycerides | HMDB0062639 |
| TG 14:0_36:3 | Triacylglyceride 14:0_36:3 | Triglycerides | HMDB0042167 |
| TG 14:0_36:4 | Triacylglyceride 14:0_36:4 | Triglycerides | HMDB0042384 |
| TG 14:0_38:4 | Triacylglyceride 14:0_38:4 | Triglycerides | HMDB0042176 |
| TG 14:0_38:5 | Triacylglyceride 14:0_38:5 | Triglycerides | HMDB0042792 |
| TG 14:0_40:5 | Triacylglyceride 14:0_40:5 | Triglycerides | NA |
| TG 16:0_28:1 | Triacylglyceride 16:0_28:1 | Triglycerides | HMDB0042273 |
| TG 16:0_28:2 | Triacylglyceride 16:0_28:2 | Triglycerides | HMDB0044030 |
| TG 16:0_30:2 | Triacylglyceride 16:0_30:2 | Triglycerides | HMDB0047771 |
| TG 16:0_32:0 | Triacylglyceride 16:0_32:0 | Triglycerides | HMDB0005356 |
| TG 16:0_32:1 | Triacylglyceride 16:0_32:1 | Triglycerides | HMDB05359 |
| TG 16:0_32:2 | Triacylglyceride 16:0_32:2 | Triglycerides | HMDB05376 |
| TG 16:0_32:3 | Triacylglyceride 16:0_32:3 | Triglycerides | HMDB0047778 |
| TG 16:0_33:1 | Triacylglyceride 16:0_33:1 | Triglycerides | HMDB0043026 |
| TG 16:0_33:2 | Triacylglyceride 16:0_33:2 | Triglycerides | HMDB0043032 |
| TG 16:0_34:1 | Triacylglyceride 16:0_34:1 | Triglycerides | HMDB05374 |
| TG 16:0_34:2 | Triacylglyceride 16:0_34:2 | Triglycerides | HMDB05362 |
| TG 16:0_34:3 | Triacylglyceride 16:0_34:3 | Triglycerides | HMDB05379 |
| TG 16:0_34:4 | Triacylglyceride 16:0_34:4 | Triglycerides | HMDB0044074 |
| TG 16:0_35:1 | Triacylglyceride 16:0_35:1 | Triglycerides | HMDB0043028 |
| TG 16:0_35:2 | Triacylglyceride 16:0_35:2 | Triglycerides | HMDB0011700 |
| TG 16:0_35:3 | Triacylglyceride 16:0_35:3 | Triglycerides | HMDB0011701 |
| TG 16:0_36:2 | Triacylglyceride 16:0_36:2 | Triglycerides | HMDB05382 |
| TG 16:0_36:3 | Triacylglyceride 16:0_36:3 | Triglycerides | HMDB05384 |
| TG 16:0_36:4 | Triacylglyceride 16:0_36:4 | Triglycerides | HMDB05390 |

| | | | |
|---|---|---|---|
| TG 16:0_36:5 | Triacylglyceride 16:0_36:5 | Triglycerides | HMDB05380 |
| TG 16:0_36:6 | Triacylglyceride 16:0_36:6 | Triglycerides | HMDB0044298 |
| TG 16:0_37:3 | Triacylglyceride 16:0_37:3 | Triglycerides | HMDB0043867 |
| TG 16:0_38:1 | Triacylglyceride 16:0_38:1 | Triglycerides | HMDB05381 |
| TG 16:0_38:2 | Triacylglyceride 16:0_38:2 | Triglycerides | HMDB05383 |
| TG 16:0_38:3 | Triacylglyceride 16:0_38:3 | Triglycerides | HMDB05389 |
| TG 16:0_38:4 | Triacylglyceride 16:0_38:4 | Triglycerides | HMDB05370 |
| TG 16:0_38:5 | Triacylglyceride 16:0_38:5 | Triglycerides | HMDB05385 |
| TG 16:0_38:6 | Triacylglyceride 16:0_38:6 | Triglycerides | HMDB10418 |
| TG 16:0_38:7 | Triacylglyceride 16:0_38:7 | Triglycerides | HMDB0067816 |
| TG 16:0_40:6 | Triacylglyceride 16:0_40:6 | Triglycerides | HMDB0044134 |
| TG 16:0_40:7 | Triacylglyceride 16:0_40:7 | Triglycerides | HMDB0067085 |
| TG 16:0_40:8 | Triacylglyceride 16:0_40:8 | Triglycerides | HMDB05392 |
| TG 16:1_28:0 | Triacylglyceride 16:1_28:0 | Triglycerides | HMDB0042069 |
| TG 16:1_30:1 | Triacylglyceride 16:1_30:1 | Triglycerides | HMDB0042309 |
| TG 16:1_32:0 | Triacylglyceride 16:1_32:0 | Triglycerides | HMDB05359 |
| TG 16:1_32:1 | Triacylglyceride 16:1_32:1 | Triglycerides | HMDB0044888 |
| TG 16:1_32:2 | Triacylglyceride 16:1_32:2 | Triglycerides | HMDB0005432 |
| TG 16:1_33:1 | Triacylglyceride 16:1_33:1 | Triglycerides | HMDB0043200 |
| TG 16:1_34:1 | Triacylglyceride 16:1_34:1 | Triglycerides | HMDB0044889 |
| TG 16:1_34:2 | Triacylglyceride 16:1_34:2 | Triglycerides | HMDB05433 |
| TG 16:1_34:3 | Triacylglyceride 16:1_34:3 | Triglycerides | HMDB05435 |
| TG 16:1_36:2 | Triacylglyceride 16:1_36:2 | Triglycerides | HMDB05438 |
| TG 16:1_36:3 | Triacylglyceride 16:1_36:3 | Triglycerides | HMDB05440 |
| TG 16:1_36:4 | Triacylglyceride 16:1_36:4 | Triglycerides | HMDB05446 |
| TG 16:1_36:5 | Triacylglyceride 16:1_36:5 | Triglycerides | HMDB05436 |
| TG 16:1_38:3 | Triacylglyceride 16:1_38:3 | Triglycerides | HMDB0046079 |
| TG 16:1_38:4 | Triacylglyceride 16:1_38:4 | Triglycerides | HMDB0046105 |
| TG 16:1_38:5 | Triacylglyceride 16:1_38:5 | Triglycerides | HMDB05441 |
| TG 17:0_32:1 | Triacylglyceride 17:0_32:1 | Triglycerides | NA |
| TG 17:0_34:1 | Triacylglyceride 17:0_34:1 | Triglycerides | NA |
| TG 17:0_34:2 | Triacylglyceride 17:0_34:2 | Triglycerides | NA |
| TG 17:0_34:3 | Triacylglyceride 17:0_34:3 | Triglycerides | NA |
| TG 17:0_36:3 | Triacylglyceride 17:0_36:3 | Triglycerides | NA |
| TG 17:0_36:4 | Triacylglyceride 17:0_36:4 | Triglycerides | NA |
| TG 17:1_32:1 | Triacylglyceride 17:1_32:1 | Triglycerides | NA |
| TG 17:1_34:1 | Triacylglyceride 17:1_34:1 | Triglycerides | NA |
| TG 17:1_34:2 | Triacylglyceride 17:1_34:2 | Triglycerides | NA |
| TG 17:1_34:3 | Triacylglyceride 17:1_34:3 | Triglycerides | NA |
| TG 17:1_36:3 | Triacylglyceride 17:1_36:3 | Triglycerides | NA |
| TG 17:1_36:4 | Triacylglyceride 17:1_36:4 | Triglycerides | NA |

| | | | |
|---|---|---|---|
| TG 17:1_36:5 | Triacylglyceride 17:1_36:5 | Triglycerides | NA |
| TG 17:2_34:2 | Triacylglyceride 17:2_34:2 | Triglycerides | NA |
| TG 17:2_36:2 | Triacylglyceride 17:2_36:2 | Triglycerides | NA |
| TG 17:2_36:3 | Triacylglyceride 17:2_36:3 | Triglycerides | NA |
| TG 17:2_36:4 | Triacylglyceride 17:2_36:4 | Triglycerides | NA |
| TG 18:0_30:0 | Triacylglyceride 18:0_30:0 | Triglycerides | HMDB0108000 |
| TG 18:0_30:1 | Triacylglyceride 18:0_30:1 | Triglycerides | HMDB0044726 |
| TG 18:0_32:0 | Triacylglyceride 18:0_32:0 | Triglycerides | HMDB0068957 |
| TG 18:0_32:1 | Triacylglyceride 18:0_32:1 | Triglycerides | HMDB0044753 |
| TG 18:0_32:2 | Triacylglyceride 18:0_32:2 | Triglycerides | HMDB0044889 |
| TG 18:0_34:3 | Triacylglyceride 18:0_34:3 | Triglycerides | HMDB05425 |
| TG 18:0_36:2 | Triacylglyceride 18:0_36:2 | Triglycerides | HMDB05403 |
| TG 18:0_36:3 | Triacylglyceride 18:0_36:3 | Triglycerides | HMDB05405 |
| TG 18:0_36:4 | Triacylglyceride 18:0_36:4 | Triglycerides | HMDB05370 |
| TG 18:0_36:5 | Triacylglyceride 18:0_36:5 | Triglycerides | HMDB05426 |
| TG 18:0_38:6 | Triacylglyceride 18:0_38:6 | Triglycerides | HMDB05412 |
| TG 18:0_38:7 | Triacylglyceride 18:0_38:7 | Triglycerides | HMDB0063278 |
| TG 18:1_26:0 | Triacylglyceride 18:1_26:0 | Triglycerides | NA |
| TG 18:1_28:1 | Triacylglyceride 18:1_28:1 | Triglycerides | HMDB0042280 |
| TG 18:1_30:0 | Triacylglyceride 18:1_30:0 | Triglycerides | HMDB0042130 |
| TG 18:1_30:1 | Triacylglyceride 18:1_30:1 | Triglycerides | HMDB0047773 |
| TG 18:1_30:2 | Triacylglyceride 18:1_30:2 | Triglycerides | HMDB0047910 |
| TG 18:1_31:0 | Triacylglyceride 18:1_31:0 | Triglycerides | HMDB0043026 |
| TG 18:1_32:0 | Triacylglyceride 18:1_32:0 | Triglycerides | HMDB05360 |
| TG 18:1_32:1 | Triacylglyceride 18:1_32:1 | Triglycerides | HMDB0062639 |
| TG 18:1_32:2 | Triacylglyceride 18:1_32:2 | Triglycerides | HMDB05433 |
| TG 18:1_32:3 | Triacylglyceride 18:1_32:3 | Triglycerides | HMDB0042354 |
| TG 18:1_33:0 | Triacylglyceride 18:1_33:0 | Triglycerides | HMDB0043055 |
| TG 18:1_33:1 | Triacylglyceride 18:1_33:1 | Triglycerides | HMDB0043229 |
| TG 18:1_33:2 | Triacylglyceride 18:1_33:2 | Triglycerides | HMDB0043235 |
| TG 18:1_33:3 | Triacylglyceride 18:1_33:3 | Triglycerides | HMDB0043243 |
| TG 18:1_34:1 | Triacylglyceride 18:1_34:1 | Triglycerides | HMDB05382 |
| TG 18:1_34:2 | Triacylglyceride 18:1_34:2 | Triglycerides | HMDB05384 |
| TG 18:1_34:3 | Triacylglyceride 18:1_34:3 | Triglycerides | HMDB05440 |
| TG 18:1_34:4 | Triacylglyceride 18:1_34:4 | Triglycerides | HMDB0048626 |
| TG 18:1_35:2 | Triacylglyceride 18:1_35:2 | Triglycerides | HMDB0043266 |
| TG 18:1_35:3 | Triacylglyceride 18:1_35:3 | Triglycerides | HMDB0043267 |
| TG 18:1_36:0 | Triacylglyceride 18:1_36:0 | Triglycerides | HMDB05395 |
| TG 18:1_36:2 | Triacylglyceride 18:1_36:2 | Triglycerides | HMDB05439 |
| TG 18:1_36:3 | Triacylglyceride 18:1_36:3 | Triglycerides | HMDB05455 |
| TG 18:1_36:4 | Triacylglyceride 18:1_36:4 | Triglycerides | HMDB05385 |

| | | | |
|---|---|---|---|
| TG 18:1_36:5 | Triacylglyceride 18:1_36:5 | Triglycerides | HMDB05441 |
| TG 18:1_36:6 | Triacylglyceride 18:1_36:6 | Triglycerides | HMDB0042359 |
| TG 18:1_38:5 | Triacylglyceride 18:1_38:5 | Triglycerides | HMDB05456 |
| TG 18:1_38:6 | Triacylglyceride 18:1_38:6 | Triglycerides | HMDB05462 |
| TG 18:1_38:7 | Triacylglyceride 18:1_38:7 | Triglycerides | HMDB0050036 |
| TG 18:2_28:0 | Triacylglyceride 18:2_28:0 | Triglycerides | HMDB0042076 |
| TG 18:2_30:0 | Triacylglyceride 18:2_30:0 | Triglycerides | HMDB0042136 |
| TG 18:2_30:1 | Triacylglyceride 18:2_30:1 | Triglycerides | HMDB0042316 |
| TG 18:2_31:0 | Triacylglyceride 18:2_31:0 | Triglycerides | HMDB0043032 |
| TG 18:2_32:0 | Triacylglyceride 18:2_32:0 | Triglycerides | HMDB05362 |
| TG 18:2_32:1 | Triacylglyceride 18:2_32:1 | Triglycerides | HMDB05379 |
| TG 18:2_32:2 | Triacylglyceride 18:2_32:2 | Triglycerides | HMDB05435 |
| TG 18:2_33:0 | Triacylglyceride 18:2_33:0 | Triglycerides | HMDB0011703 |
| TG 18:2_33:1 | Triacylglyceride 18:2_33:1 | Triglycerides | HMDB0043235 |
| TG 18:2_33:2 | Triacylglyceride 18:2_33:2 | Triglycerides | HMDB0011711 |
| TG 18:2_34:1 | Triacylglyceride 18:2_34:1 | Triglycerides | HMDB0045870 |
| TG 18:2_34:2 | Triacylglyceride 18:2_34:2 | Triglycerides | HMDB05390 |
| TG 18:2_34:3 | Triacylglyceride 18:2_34:3 | Triglycerides | HMDB05446 |
| TG 18:2_34:4 | Triacylglyceride 18:2_34:4 | Triglycerides | HMDB0048758 |
| TG 18:2_35:1 | Triacylglyceride 18:2_35:1 | Triglycerides | HMDB0052765 |
| TG 18:2_35:2 | Triacylglyceride 18:2_35:2 | Triglycerides | HMDB0052413 |
| TG 18:2_35:3 | Triacylglyceride 18:2_35:3 | Triglycerides | HMDB0043406 |
| TG 18:2_36:1 | Triacylglyceride 18:2_36:1 | Triglycerides | HMDB05389 |
| TG 18:2_36:2 | Triacylglyceride 18:2_36:2 | Triglycerides | HMDB05455 |
| TG 18:2_36:3 | Triacylglyceride 18:2_36:3 | Triglycerides | HMDB05461 |
| TG 18:2_36:4 | Triacylglyceride 18:2_36:4 | Triglycerides | HMDB0005474 |
| TG 18:2_36:5 | Triacylglyceride 18:2_36:5 | Triglycerides | HMDB05447 |
| TG 18:2_38:4 | Triacylglyceride 18:2_38:4 | Triglycerides | HMDB05412 |
| TG 18:2_38:5 | Triacylglyceride 18:2_38:5 | Triglycerides | HMDB0050760 |
| TG 18:2_38:6 | Triacylglyceride 18:2_38:6 | Triglycerides | HMDB05475 |
| TG 18:3_30:0 | Triacylglyceride 18:3_30:0 | Triglycerides | HMDB0043011 |
| TG 18:3_32:0 | Triacylglyceride 18:3_32:0 | Triglycerides | HMDB10417 |
| TG 18:3_32:1 | Triacylglyceride 18:3_32:1 | Triglycerides | HMDB0047809 |
| TG 18:3_34:0 | Triacylglyceride 18:3_34:0 | Triglycerides | HMDB0043934 |
| TG 18:3_34:1 | Triacylglyceride 18:3_34:1 | Triglycerides | HMDB0045896 |
| TG 18:3_34:2 | Triacylglyceride 18:3_34:2 | Triglycerides | HMDB0048096 |
| TG 18:3_34:3 | Triacylglyceride 18:3_34:3 | Triglycerides | HMDB0048758 |
| TG 18:3_35:2 | Triacylglyceride 18:3_35:2 | Triglycerides | HMDB0043643 |
| TG 18:3_36:1 | Triacylglyceride 18:3_36:1 | Triglycerides | HMDB0046079 |
| TG 18:3_36:2 | Triacylglyceride 18:3_36:2 | Triglycerides | HMDB10460 |
| TG 18:3_36:3 | Triacylglyceride 18:3_36:3 | Triglycerides | HMDB0044287 |

| | | | |
|---|---|---|---|
| TG 18:3_36:4 | Triacylglyceride 18:3_36:4 | Triglycerides | HMDB10490 |
| TG 18:3_38:5 | Triacylglyceride 18:3_38:5 | Triglycerides | HMDB0050768 |
| TG 18:3_38:6 | Triacylglyceride 18:3_38:6 | Triglycerides | HMDB0053582 |
| TG 20:0_32:3 | Triacylglyceride 20:0_32:3 | Triglycerides | HMDB0045870 |
| TG 20:0_32:4 | Triacylglyceride 20:0_32:4 | Triglycerides | HMDB0045896 |
| TG 20:0_34:1 | Triacylglyceride 20:0_34:1 | Triglycerides | HMDB0045584 |
| TG 20:1_32:0 | Triacylglyceride 20:1_32:0 | Triglycerides | NA |
| TG 20:1_32:1 | Triacylglyceride 20:1_32:1 | Triglycerides | HMDB05378 |
| TG 20:1_32:2 | Triacylglyceride 20:1_32:2 | Triglycerides | HMDB05434 |
| TG 20:1_32:3 | Triacylglyceride 20:1_32:3 | Triglycerides | HMDB0048073 |
| TG 20:1_34:1 | Triacylglyceride 20:1_34:1 | Triglycerides | HMDB05424 |
| TG 20:1_34:2 | Triacylglyceride 20:1_34:2 | Triglycerides | HMDB05439 |
| TG 20:1_34:3 | Triacylglyceride 20:1_34:3 | Triglycerides | HMDB05445 |
| TG 20:2_32:0 | Triacylglyceride 20:2_32:0 | Triglycerides | HMDB0043900 |
| TG 20:2_32:1 | Triacylglyceride 20:2_32:1 | Triglycerides | HMDB0044068 |
| TG 20:2_34:1 | Triacylglyceride 20:2_34:1 | Triglycerides | HMDB0044096 |
| TG 20:2_34:2 | Triacylglyceride 20:2_34:2 | Triglycerides | HMDB0042588 |
| TG 20:2_34:3 | Triacylglyceride 20:2_34:3 | Triglycerides | HMDB0048125 |
| TG 20:3_32:0 | Triacylglyceride 20:3_32:0 | Triglycerides | HMDB0042169 |
| TG 20:3_32:1 | Triacylglyceride 20:3_32:1 | Triglycerides | HMDB0047798 |
| TG 20:3_32:2 | Triacylglyceride 20:3_32:2 | Triglycerides | HMDB0048599 |
| TG 20:3_34:0 | Triacylglyceride 20:3_34:0 | Triglycerides | HMDB0043929 |
| TG 20:3_34:1 | Triacylglyceride 20:3_34:1 | Triglycerides | HMDB0044091 |
| TG 20:3_34:2 | Triacylglyceride 20:3_34:2 | Triglycerides | HMDB0042589 |
| TG 20:3_34:3 | Triacylglyceride 20:3_34:3 | Triglycerides | HMDB0048126 |
| TG 20:3_36:3 | Triacylglyceride 20:3_36:3 | Triglycerides | HMDB0049390 |
| TG 20:3_36:4 | Triacylglyceride 20:3_36:4 | Triglycerides | HMDB10492 |
| TG 20:3_36:5 | Triacylglyceride 20:3_36:5 | Triglycerides | HMDB0044357 |
| TG 20:4_30:0 | Triacylglyceride 20:4_30:0 | Triglycerides | HMDB0042146 |
| TG 20:4_32:0 | Triacylglyceride 20:4_32:0 | Triglycerides | HMDB05363 |
| TG 20:4_32:1 | Triacylglyceride 20:4_32:1 | Triglycerides | HMDB05380 |
| TG 20:4_32:2 | Triacylglyceride 20:4_32:2 | Triglycerides | HMDB05436 |
| TG 20:4_33:2 | Triacylglyceride 20:4_33:2 | Triglycerides | HMDB0043419 |
| TG 20:4_34:0 | Triacylglyceride 20:4_34:0 | Triglycerides | HMDB05370 |
| TG 20:4_34:1 | Triacylglyceride 20:4_34:1 | Triglycerides | HMDB05385 |
| TG 20:4_34:2 | Triacylglyceride 20:4_34:2 | Triglycerides | HMDB05391 |
| TG 20:4_34:3 | Triacylglyceride 20:4_34:3 | Triglycerides | HMDB05447 |
| TG 20:4_36:2 | Triacylglyceride 20:4_36:2 | Triglycerides | HMDB05412 |
| TG 20:4_36:3 | Triacylglyceride 20:4_36:3 | Triglycerides | HMDB05462 |
| TG 20:4_36:4 | Triacylglyceride 20:4_36:4 | Triglycerides | HMDB05475 |
| TG 20:4_36:5 | Triacylglyceride 20:4_36:5 | Triglycerides | HMDB05448 |

| | | | |
|---|---|---|---|
| TG 20:5_34:1 | Triacylglyceride 20:5_34:1 | Triglycerides | HMDB0047835 |
| TG 20:5_34:2 | Triacylglyceride 20:5_34:2 | Triglycerides | HMDB0048629 |
| TG 20:5_36:2 | Triacylglyceride 20:5_36:2 | Triglycerides | HMDB10464 |
| TG 20:5_36:3 | Triacylglyceride 20:5_36:3 | Triglycerides | HMDB0044357 |
| TG 22:3_30:2 | Triacylglyceride 22:3_30:2 | Triglycerides | NA |
| TG 22:4_32:0 | Triacylglyceride 22:4_32:0 | Triglycerides | HMDB0042172 |
| TG 22:4_34:2 | Triacylglyceride 22:4_34:2 | Triglycerides | HMDB0042592 |
| TG 22:5_32:0 | Triacylglyceride 22:5_32:0 | Triglycerides | HMDB0043910 |
| TG 22:5_32:1 | Triacylglyceride 22:5_32:1 | Triglycerides | HMDB0047813 |
| TG 22:5_34:1 | Triacylglyceride 22:5_34:1 | Triglycerides | HMDB0047836 |
| TG 22:5_34:2 | Triacylglyceride 22:5_34:2 | Triglycerides | HMDB0047997 |
| TG 22:5_34:3 | Triacylglyceride 22:5_34:3 | Triglycerides | HMDB0044498 |
| TG 22:6_32:0 | Triacylglyceride 22:6_32:0 | Triglycerides | HMDB10418 |
| TG 22:6_32:1 | Triacylglyceride 22:6_32:1 | Triglycerides | HMDB0042359 |
| TG 22:6_34:1 | Triacylglyceride 22:6_34:1 | Triglycerides | HMDB0044107 |
| TG 22:6_34:2 | Triacylglyceride 22:6_34:2 | Triglycerides | HMDB0048631 |
| TG 22:6_34:3 | Triacylglyceride 22:6_34:3 | Triglycerides | HMDB0048136 |
| Hex3Cer d18:1/16:0 | Trihexosylceramide d18:1/16:0 | Trihexosylceramides | HMDB0004879 |
| Hex3Cer d18:1/18:0 | Trihexosylceramide d18:1/18:0 | Trihexosylceramides | HMDB0004880 |
| Hex3Cer d18:1/22:0 | Trihexosylceramide d18:1/22:0 | Trihexosylceramides | HMDB0004882 |
| Hex3Cer d18:1/24:1 | Trihexosylceramide d18:1/24:1 | Trihexosylceramides | HMDB0004883 |
| Biotin (B7) | Biotin (B7) | Vitamins and cofactors | HMDB0000030 |
| Choline | Choline | Vitamins and cofactors | HMDB0000097 |
| Nicotinamide (B3) | Nicotinamide (B3) | Vitamins and cofactors | HMDB0001406 |
| Pantothenic acid (B5) | Pantothenic acid (B5) | Vitamins and cofactors | HMDB0250782 |

**SUPPLEMENTAL DATA 3**

| Short name | Formula |
|---|---|
| TMAO Synthesis | TMAO / (Betaine + C0 + Choline) |
| TMAO Synthesis (direct) | TMAO / Choline |
| AGAT Deficiency | Arg / Guanidinoacetic acid |
| Asn Synthesis | Asn / Asp |
| Brain Trp Availability | Trp / (Leu + Ile + Val + Tyr + Phe + Met) |
| Cys Synthesis | Cys / (Ser + Met) |
| DLD (NBS) | Pro / Phe |
| Fischer Ratio | (Ile + Leu + Val) / (Phe + Trp + Tyr) |
| GABR | Arg / (Orn + Cit) |
| Glutaminase Activity | Glu / Gln |
| Glutaminolysis Rate | (Ala + Asp + Glu + Lac + Suc) / Gln |
| GSH Constituents | Glu + Gly + Cys |
| Gly Synthesis | Gly / Ser |
| MTHFR Deficiency (NBS) | Met / Phe |
| PKU (NBS) | Tyr / Phe |
| Glx to N-Ac-Asp Ratio | (Gln + Glu) / N-Ac-Asp |
| Glu to a-Ketoglutarate Ratio | Glu / a-Ketoglutaric acid |
| Gly to Ala Ratio | Gly / Ala |
| Ratio of His to HCys+Phe+Sar | His / (HCys + Phe + Sarcosine) |
| Ratio of Non-Essential to Essential AAs | (Ala + Arg + Asn + Asp + Cys + Gln + Glu + Gly + Pro + Ser + Tyr) / (His + Ile + Leu + Lys + Met + Phe + Thr + Trp + Val) |
| Ratio of Pro to Cit | Pro / Cit |
| Ratio of SG to Glucose | (Gly + Ser) / Glucose |
| Ratio of SGA to Glucose | (Gly + Ser + Ala) / Glucose |
| Sum of AAs | Ala + Arg + Asn + Asp + Cys + Gln + Glu + Gly + His + Ile + Leu + Lys + Met + Phe + Pro + Ser + Thr + Trp + Tyr + Val |
| Sum of Aromatic AAs | Phe + Trp + Tyr |
| Sum of BCAAs | Ile + Leu + Val |
| Sum of Essential AAs | His + Ile + Leu + Lys + Met + Phe + Thr + Trp + Val |
| Sum of Non-Essential AAs | Ala + Arg + Asn + Asp + Cys + Gln + Glu + Gly + Pro + Ser + Tyr |
| Sum of Solely Glucogenic AAs | Ala + Arg + Asn + Asp + Cys + Gln + Glu + Gly + His + Met + Pro + Ser + Thr + Val |
| Sum of Solely Ketogenic AAs | Leu + Lys |
| Sum of Sulfur-Containing AAs | Met + Cys |
| Valinemia (NBS) | Val / Phe |
| 1-Met-His Synthesis | 1-Met-His / His |
| AABA Synthesis | AABA / Thr |
| ACY1 Deficiency | (N-Ac-Met + N-Ac-Glu + N-Ac-Gly + N-Ac-Ala + N-Ac-Leu + N-Ac-Ile + N-Ac-Val) / (Met + Glu + Gly + Ala + Leu + Ile + Val) |
| ACY2 Deficiency | N-Ac-Asp / Asp |
| Asp Methylation | N-Met-Asp / Asp |

| | |
|---|---|
| Asymmetrical Arg Methylation | ADMA / Arg |
| BABA Synthesis | BABA / Glu |
| BAIBA Synthesis | BAIBA / Val |
| Betaine Synthesis | Betaine / Choline |
| BHMT Activity | DMG / Betaine |
| CPS Deficiency (NBS) | Cit / Phe |
| Carnosine Synthesis | Carnosine / His |
| Cit Synthesis | Cit / Orn |
| Citrullinemia | Cit / (Orn + Arg) |
| Cystine Synthesis | Cystine / Cys |
| Glu Acetylation | N-Ac-Glu / Glu |
| GAMT Deficiency | Guanidinoacetic acid / Creatine |
| HArg Synthesis | HArg / (Arg + Lys) |
| HCys Synthesis | HCys / Met |
| Imidazolepropionic Acid Synthesis | Imidazolepropionic acid / His |
| IDO Activity | Kynurenine / Trp |
| BCAT Ile Catabolism | 3-Met-2-oxovaleric acid / Ile |
| Kynureninase A Activity | Anthranilic acid / Kynurenine |
| BCAT Leu Catabolism | 4-Met-2-oxovaleric acid / Leu |
| Lys Carbamylation | HCit / Lys |
| Met Oxidation | Met-SO / Met |
| Muscle Protein Degradation | 1-Met-His / Creatinine |
| NOS activity | Cit / Arg |
| Orn Synthesis | Orn / Arg |
| OTC Deficiency (NBS) | Orn / Cit |
| Pro Hydroxylation | (c4-OH-Pro + t4-OH-Pro) / Pro |
| 3-Met-His to 1-Met-His Ratio | 3-Met-His / 1-Met-His |
| Creatine to Creatinine Ratio | Creatine / Creatinine |
| Creatinine to Bilirubin Ratio | Creatinine / Bilirubin |
| GAA to HArg Ratio | Guanidinoacetic acid / HArg |
| Ratio of HArg to ADMA | HArg / ADMA |
| Ratio of HArg to SDMA | HArg / SDMA |
| HSer to Creatinine Ratio | HSer / Creatinine |
| Kynurenine to Serotonin Ratio | Kynurenine / Serotonin |
| Sarcosine Synthesis from Choline | Sarcosine / Choline |
| Sarcosine Synthesis from Gly | Sarcosine / Gly |
| Sum of Aminobutyric Acids | AABA + BABA + GABA |
| Sum of Asym. and Sym. Arg Methylation | (ADMA + SDMA) / Arg |
| Sum of BC-a-ketoacids | 3-Met-2-oxovaleric acid + a-Ketoisovaleric acid + 4-Met-2-oxovaleric acid |
| Sum of Dimethylated Arg | ADMA + SDMA |
| Symmetrical Arg Methylation | SDMA / Arg |

| | |
|---|---|
| Taurine Synthesis | Taurine / Cys |
| BCAT Val Catabolism | a-Ketoisovaleric acid / Val |
| 12-KetoDCA to DCA Ratio | 12-KetoDCA / DCA |
| 7a-Dehydroxylation of CA | DCA / CA |
| Gly Conjugation of CDCA | GCDCA / CDCA |
| Gly Conjugation of CA | GCA / CA |
| Gly Conjugation of DCA | GDCA / DCA |
| Gly Conjugation of Primary BAs | (GCA + GCDCA) / (CA + CDCA) |
| Gly Conjugation of UDCA | GUDCA / UDCA |
| GDCA Synthesis from CA | GDCA / CA |
| GLCA Synthesis from CDCA | GLCA / CDCA |
| Ratio of CDCA to CA | CDCA / CA |
| Primary BA Conjugation | (GCA + GCDCA + TCA + TCDCA) / (CA + CDCA) |
| IsoUDCA to UDCA Ratio | IsoUDCA / UDCA |
| Sum of 12a-OH BAs | 3-EpiDCA + 7-KetoDCA + CA + DCA + GCA + GDCA + NorDCA + TCA + TDCA + THDCA |
| Sum of Conjugated Primary BAs | GCA + GCDCA + TCA + TCDCA |
| Sum of Gly-Conjugated BAs | GCA + GCDCA + GDCA + GLCA + GUDCA |
| Sum of Primary BAs | CA + CDCA + GCA + GCDCA + TCA + TCDCA |
| Sum of Unconjugated Primary BAs | CA + CDCA |
| Taurine Conjugation of CDCA | TCDCA / CDCA |
| Taurine Conjugation of CA | TCA / CA |
| Taurine Conjugation of DCA | TDCA / DCA |
| Taurine Conjugation of Primary BAs | (TCA + TCDCA) / (CA + CDCA) |
| Taurine Conjugation of UDCA | TUDCA / UDCA |
| TDCA Synthesis from CA | TDCA / CA |
| TLCA Synthesis from CDCA | TLCA / CDCA |
| UDCA Synthesis from CDCA | UDCA / CDCA |
| b-Ala Synthesis | beta-Ala / Carnosine |
| GABA Synthesis | GABA / Glu |
| Histamine Synthesis | Histamine / His |
| Putrescine Synthesis | Putrescine / Orn |
| Urea to Creatinine Ratio | Urea / Creatinine |
| Serotonin Synthesis | Serotonin / Trp |
| Spermidine Synthesis | Spermidine / Putrescine |
| Benzoic Acid Conjugation | HipAcid / Benzoic acid |
| Cholesterol Synthesis | Mevalonic acid / Creatinine |
| HipAcid Synthesis | HipAcid / Gly |
| HMGCR Activity | Mevalonic acid / 3-HMGA |
| Ketone Body Synthesis | 3-OH-Butyric acid / FA 4:0 |
| LDH Activity | Lac / Glucose |
| MRC Disorders | Lac / Pyruvic acid |

| | |
|---|---|
| Pyruvate Carboxylase Deficiency | Pyruvic acid / Oxaloacetic acid |
| Glycolic Acid to Oxalic Acid Ratio | Glycolic acid / Oxalic acid |
| Lac to FA 2:0 Ratio | Lac / FA 2:0 |
| Pyruvate to Glucose Ratio | Pyruvic acid / Glucose |
| Sum of Carboxylic Acids | 2-OH-Butyric acid + 3-OH-Butyric acid + Glycolic acid + Glyoxylic acid + HipAcid + Lac + Mevalonic acid + Pyruvic acid |
| p-Cresol-SO4 Synthesis | p-Cresol-SO4 / Tyr |
| p-Cresol-SO4 to p-Cresol Glucuronide | p-Cresol-SO4 / p-Cresol glucuronide |
| Total p-Cresol Derivatives | p-Cresol-SO4 + p-Cresol glucuronide |
| 2-OH-Glutarate Synthesis | 2-OH-Glutaric acid / a-Ketoglutaric acid |
| 3-MAG-uria Type 1 | Z-3-Met-glutaconic acid / E-3-Met-glutaconic acid |
| ACOD1 Activity | Itaconic acid / AconAcid |
| CMAMMA | Malonic acid / Methylmalonic acid |
| Ethylmalonic Aciduria | Ethylmalonic acid / Suc |
| Fumarase Activity | Malic acid / Fumaric acid |
| GCDH Deficiency | 3-OH-Glutaric acid / Glutaric acid |
| MDH1 Activity | Malic acid / Oxaloacetic acid |
| MDH2 Activity | Oxaloacetic acid / Malic acid |
| a-Ketoglutarate to Citrate Ratio | a-Ketoglutaric acid / Citric acid |
| Suc to a-Ketoglutarate Ratio | Suc / a-Ketoglutaric acid |
| Secondary 3MG Aciduria | 3-Met-glutaric acid / (Z-3-Met-glutaconic acid + E-3-Met-glutaconic acid) |
| SDH Activity | Suc / Fumaric acid |
| FA 2:0 to Glucose Ratio | FA 2:0 / Glucose |
| Acetic Acid to Butyric Acid Ratio | FA 2:0 / FA 4:0 |
| Acetic Acid to Isobutyric Acid Ratio | FA 2:0 / FA 3:0-2M |
| Butyric Acid to Isobutyric Acid Ratio | FA 4:0 / FA 3:0-2M |
| FA 8:0 to FA 10:0 Ratio | FA 8:0 / FA 10:0 |
| FA 16:0 to FA 18:1 Ratio | FA 16:0 / FA 18:1 |
| FA16:0 to FA 16:1 Ratio | FA 16:0 / FA 16:1 |
| SCD-1 index | FA 18:1 / FA 18:0 |
| Sum BCFA | FA 3:0-2M + FA 4:0-2M + FA 4:0-3M + FA 5:0-3M + FA 5:0-4M |
| Sum MCFA | FA 7:0 + FA 8:0 + FA 9:0 + FA 10:0 + FA 11:0 + FA 12:0 |
| Sum of MUFAs | FA 14:1n-5 + FA 16:1 + FA 18:1 + FA 20:1n-9 + FA 24:1n-9 |
| MCFA PPAR Modulators | FA 8:0 + FA 9:0 + FA 10:0 |
| Sum of FA 12:0 - FA 16:0 | FA 12:0 + FA 14:0 + FA 15:0 + FA 16:0 |
| Cortisone Synthesis | Cortisone / Cortisol |
| Sum of Steroid Hormones | Cortisol + Cortisone + DHEAS |
| Indole Pathway Activity | 3-IPA / Trp |
| 5-HIAA to Creatinine Ratio | 5-HIAA / Creatinine |
| 5-HIAA / Quinolinic Acid Ratio | 5-HIAA / Quinolinic acid |
| Serotonin Catabolism | 5-HIAA / Serotonin |

| | |
|---|---|
| Serotonin Pathway Activity | 5-HIAA / Trp |
| Sum of AAA-derived Lactic Acids | Indole-Lac + Phenyl-Lac + OH-Phenyl-Lac |
| NAT10 Activity | N-Ac-Cytidine / Cytidine |
| AHCys to Leu Ratio | AHCys / Leu |
| Uric Acid to Creatinine Ratio | Uric acid / Creatinine |
| AHCY Deficiency | AHCys / (Adenosine + HCys) |
| Xanthine Synthesis | Xanthine / Hypoxanthine |
| Cyclic Nucleotides Ratio | cAMP / cGMP |
| BVRA Activity | Bilirubin / Biliverdin |
| Phenylpyruvate Synthesis | Phenylpyruvic acid / Phe |
| TAT Activity | 4-OH-Phenylpyruvic acid / Tyr |
| Homovanillate to Vanillylmandelate Ratio | Homovanillic acid / Vanillylmandelic acid |
| Phenylacetate to PAGln Ratio | Phenylacetic acid / Phenylacetylglutamine |
| Phenylpyruvate to Citrate Ratio | Phenylpyruvic acid / Citric acid |
| N-Ac-Putrescine to N1- and N8-Ac-Spermidine Ratio | N-Ac-Putrescine / (N1-Ac-Spermidine + N8-Ac-Spermidine) |
| Spermidine Acetylation | (N1-Ac-Spermidine + N8-Ac-Spermidine + N1,N8-Di-Ac-Spermidine) / Spermidine |
| AMSDH Saturation | Picolinic acid / Quinolinic acid |
| KYNA DH | Quinaldic acid / Kynurenic acid |
| KAT A Activity | Kynurenic acid / Kynurenine |
| Ratio Quinolinic Acid to KYNA | Quinolinic acid / Kynurenic acid |
| Kynurenic Acid to Xanthurenic Acid Ratio | Kynurenic acid / Xanthurenic acid |
| Picolinic Acid to Trp Ratio | Picolinic acid / Trp |
| Fructose to Glucose Ratio | Fructose / Glucose |
| Aconitase Activity | Isocitric acid / Citric acid |
| CS Activity | Citric acid / Oxaloacetic acid |
| Aconitate to Citrate Ratio | AconAcid / Citric acid |
| Choline to Creatinine Ratio | Choline / Creatinine |
| 2MBG (NBS) | C5 / C3 |
| b-Oxidation | (C2 + C3) / C0 |
| CACT Deficiency (NBS) | (C16 + C18) / C0 |
| CPT-1 Deficiency (NBS) | C0 / (C16 + C18) |
| CPT-2 Deficiency (NBS) | (C16 + C18:1) / C2 |
| Carnitine Uptake Defect (NBS) | (C0 + C2 + C3 + C16 + C18 + C18:1) / Cit |
| EMA (NBS) | C4 / C8 |
| IBD Deficiency (NBS) | C4 / C2 |
| IVA (NBS) | C5 / C2 |
| MA (NBS) | C3 / C2 |
| MCAD Deficiency (NBS) | C8 / C2 |
| MCKAT Deficiency (NBS) | C8 / C10 |
| MMA (NBS) | C3 / C0 |
| MC Deficiency (NBS) | C16 / C3 |

| | |
|---|---|
| PA (NBS) | C3 / C16 |
| Ratio of Acetylcarnitine to Carnitine | C2 / C0 |
| Carnitine to Creatinine Ratio | C0 / Creatinine |
| SBCAD Deficiency (NBS) | C5 / C0 |
| SCAD Deficiency (NBS) | C4 / C3 |
| VLCAD Deficiency (NBS) | C14:1 / C16 |
| Sum of VLCFA-Cer | Cer d16:1/22:0 + Cer d16:1/23:0 + Cer d16:1/24:0 + Cer d18:1/22:0 + Cer d18:1/23:0 + Cer d18:1/24:0 + Cer d18:1/24:1 + Cer d18:1/25:0 + Cer d18:1/26:0 + Cer d18:1/26:1 + Cer d18:2/22:0 + Cer d18:2/23:0 + Cer d18:2/24:0 + Cer d18:2/24:1 |
| Sum of PUFA-CEs | CE 18:2 + CE 18:3 + CE 20:3 + CE 20:4 + CE 20:5 + CE 22:2 + CE 22:5 + CE 22:6 |
| Protection against MDD | DG 18:1_18:2 / DG 18:2_20:4 |
| Ratio of MUFA-LPCs to SFA-LPCs | (LPC 26:1 + LPC 18:1 + LPC 16:1) / (LPC 26:0 + LPC 24:0 + LPC 18:0 + LPC 17:0 + LPC 16:0) |
| Ratio of PUFA-LPCs to MUFA-LPCs | (LPC 20:4 + LPC 20:3 + LPC 18:2) / (LPC 26:1 + LPC 18:1 + LPC 16:1) |
| Ratio of PUFA-LPCs to SFA-LPCs | (LPC 20:4 + LPC 20:3 + LPC 18:2) / (LPC 26:0 + LPC 24:0 + LPC 18:0 + LPC 17:0 + LPC 16:0) |
| Ratio of UFA-LPCs to SFA-LPCs | (LPC 26:1 + LPC 20:4 + LPC 20:3 + LPC 18:2 + LPC 18:1 + LPC 16:1) / (LPC 26:0 + LPC 24:0 + LPC 18:0 + LPC 17:0 + LPC 16:0) |
| Sum of LCFA-LPCs | LPC 14:0 + LPC 16:0 + LPC 16:1 + LPC 17:0 + LPC 18:0 + LPC 18:1 + LPC 18:2 + LPC 20:3 + LPC 20:4 |
| Sum of LPCs | LPC 14:0 + LPC 16:0 + LPC 16:1 + LPC 17:0 + LPC 18:0 + LPC 18:1 + LPC 18:2 + LPC 20:3 + LPC 20:4 + LPC 24:0 + LPC 26:0 + LPC 26:1 |
| Sum of MUFA-LPCs | LPC 16:1 + LPC 18:1 + LPC 26:1 |
| Sum of PUFA-LPCs | LPC 18:2 + LPC 20:3 + LPC 20:4 |
| Sum of SFA-LPCs | LPC 14:0 + LPC 16:0 + LPC 17:0 + LPC 18:0 + LPC 24:0 + LPC 26:0 |
| Sum of UFA-LPCs | LPC 16:1 + LPC 18:1 + LPC 18:2 + LPC 20:3 + LPC 20:4 + LPC 26:1 |
| Sum of VLCFA-LPCs | LPC 24:0 + LPC 26:0 + LPC 26:1 |
| Sum of MUFA-LPGs | LPG 14:1 + LPG 16:1 + LPG 17:1 + LPG 18:1 + LPG 20:1 |
| Sum of UFA-LPGs | LPG 14:1 + LPG 16:1 + LPG 17:1 + LPG 18:1 + LPG 18:2 + LPG 20:1 |
| PLA activity (2) | (LPI 22:1 + LPI 22:0 + LPI 20:4 + LPI 20:1 + LPI 19:0 + LPI 18:3 + LPI 18:2 + LPI 18:1 + LPI 18:0 + LPI 17:1 + LPI 17:0 + LPI 16:1 + LPI 16:0 + LPI 14:1 + LPI 14:0) / (PI 18:2_22:6 + PI 18:2_22:1 + PI 18:2_22:0 + PI 18:2_20:5 + PI 18:2_20:4 + PI 18:2_20:1 + PI 18:2_20:0 + PI 18:2_18:3 + PI 18:1_22:6 + PI 18:1_22:5 + PI 18:1_22:4 + PI 18:1_22:3 + PI 18:1_22:2 + PI 18:1_22:1 + PI 18:1_22:0 + PI 18:1_20:5 + PI 18:1_20:4 + PI 18:1_20:3 + PI 18:1_20:2 + PI 18:1_20:1 + PI 18:1_20:0 + PI 18:1_18:3 + PI 18:1_18:2 + PI 18:1_18:1 + PI 18:0_22:0 + PI 18:0_20:4 + PI 18:0_20:3 + PI 18:0_20:0 + PI 18:0_18:3 + PI 18:0_18:2 + PI 18:0_18:1 + PI 18:0_18:0 + PI 17:1_18:2 + PI 17:1_18:1 + PI 17:0_18:1 + PI 16:1_18:2 + PI 16:1_18:1 + PI 16:1_18:0 + PI 16:0_22:1 + PI 16:0_20:4 + PI 16:0_20:3 + PI 16:0_20:0 + PI 16:0_18:3 + PI 16:0_18:2 + PI 16:0_18:1 + PI 16:0_17:2 + PI 16:0_17:1 + PI 16:0_17:0 + PI 16:0_16:0 + PI 15:1_16:0 + PI 15:0_16:0 + PI 14:0_18:2 + PI 14:0_18:1) |

| | |
|---|---|
| Ratio of MUFA-LPIs to SFA-LPIs | (LPI 14:1 + LPI 16:1 + LPI 17:1 + LPI 18:1 + LPI 20:1 + LPI 22:1) / (LPI 14:0 + LPI 16:0 + LPI 17:0 + LPI 18:0 + LPI 19:0 + LPI 22:0) |
| Ratio of PUFA-LPIs to MUFA-LPIs | (LPI 18:2 + LPI 18:3 + LPI 20:4) / (LPI 14:1 + LPI 16:1 + LPI 17:1 + LPI 18:1 + LPI 20:1 + LPI 22:1) |
| Ratio of PUFA-LPIs to SFA-LPIs | (LPI 18:2 + LPI 18:3 + LPI 20:4) / (LPI 14:0 + LPI 16:0 + LPI 17:0 + LPI 18:0 + LPI 19:0 + LPI 22:0) |
| Ratio of UFA-LPIs to SFA-LPIs | (LPI 14:1 + LPI 16:1 + LPI 17:1 + LPI 18:1 + LPI 18:2 + LPI 18:3 + LPI 20:1 + LPI 20:4 + LPI 22:1) / (LPI 14:0 + LPI 16:0 + LPI 17:0 + LPI 18:0 + LPI 19:0 + LPI 22:0) |
| Sum of LCFA-LPIs | LPI 14:0 + LPI 14:1 + LPI 16:0 + LPI 16:1 + LPI 17:0 + LPI 17:1 + LPI 18:0 + LPI 18:1 + LPI 18:2 + LPI 18:3 + LPI 19:0 + LPI 20:1 + LPI 20:4 |
| Sum of LPIs | LPI 14:0 + LPI 14:1 + LPI 16:0 + LPI 16:1 + LPI 17:0 + LPI 17:1 + LPI 18:0 + LPI 18:1 + LPI 18:2 + LPI 18:3 + LPI 19:0 + LPI 20:1 + LPI 20:4 + LPI 22:0 + LPI 22:1 |
| Sum of MUFA-LPIs | LPI 14:1 + LPI 16:1 + LPI 17:1 + LPI 18:1 + LPI 20:1 + LPI 22:1 |
| Sum of (L)PIs | LPI 14:0 + LPI 14:1 + LPI 16:0 + LPI 16:1 + LPI 17:0 + LPI 17:1 + LPI 18:0 + LPI 18:1 + LPI 18:2 + LPI 18:3 + LPI 19:0 + LPI 20:1 + LPI 20:4 + LPI 22:0 + LPI 22:1 + PI 14:0_18:1 + PI 14:0_18:2 + PI 15:0_16:0 + PI 15:1_16:0 + PI 16:0_16:0 + PI 16:0_17:0 + PI 16:0_17:1 + PI 16:0_17:2 + PI 16:0_18:1 + PI 16:0_18:2 + PI 16:0_18:3 + PI 16:0_20:0 + PI 16:0_20:3 + PI 16:0_20:4 + PI 16:0_22:1 + PI 16:1_18:0 + PI 16:1_18:1 + PI 16:1_18:2 + PI 17:0_18:1 + PI 17:1_18:1 + PI 17:1_18:2 + PI 18:0_18:0 + PI 18:0_18:1 + PI 18:0_18:2 + PI 18:0_18:3 + PI 18:0_20:0 + PI 18:0_20:3 + PI 18:0_20:4 + PI 18:0_22:0 + PI 18:1_18:1 + PI 18:1_18:2 + PI 18:1_18:3 + PI 18:1_20:0 + PI 18:1_20:1 + PI 18:1_20:2 + PI 18:1_20:3 + PI 18:1_20:4 + PI 18:1_20:5 + PI 18:1_22:0 + PI 18:1_22:1 + PI 18:1_22:2 + PI 18:1_22:3 + PI 18:1_22:4 + PI 18:1_22:5 + PI 18:1_22:6 + PI 18:2_18:3 + PI 18:2_20:0 + PI 18:2_20:1 + PI 18:2_20:4 + PI 18:2_20:5 + PI 18:2_22:0 + PI 18:2_22:1 + PI 18:2_22:6 |
| Sum of PUFA-LPIs | LPI 18:2 + LPI 18:3 + LPI 20:4 |
| Sum of SFA-LPIs | LPI 14:0 + LPI 16:0 + LPI 17:0 + LPI 18:0 + LPI 19:0 + LPI 22:0 |
| Sum of UFA-LPIs | LPI 14:1 + LPI 16:1 + LPI 17:1 + LPI 18:1 + LPI 18:2 + LPI 18:3 + LPI 20:1 + LPI 20:4 + LPI 22:1 |
| Sum of VLCFA-LPIs | LPI 22:0 + LPI 22:1 |
| Sum of MUFA-PAs | PA 14:0_14:1 + PA 16:0_18:1 + PA 16:1_18:1 + PA 16:1_22:0 + PA 17:0_18:1 + PA 17:1_18:1 + PA 18:0_18:1 + PA 18:1_18:1 + PA 18:1_20:0 + PA 18:1_20:1 + PA 18:1_22:0 + PA 18:1_22:1 |
| Ratio of PUFA-PCs O to MUFA-PCs O | (PC O-34:3 + PC O-36:3 + PC O-36:4 + PC O-36:5 + PC O-38:3 + PC O-38:4 + PC O-38:5 + PC O-38:6 + PC O-40:3 + PC O-40:4 + PC O-40:5 + PC O-40:6 + PC O-42:3 + PC O-42:4 + PC O-42:5 + PC O-44:3 + PC O-44:4 + PC O-44:5 + PC O-44:6) / (PC O-28:1 + PC O-30:1 + PC O-32:1 + PC O-34:1 + PC O-36:1 + PC O-38:1 + PC O-40:1 + PC O-42:1) |
| Sum of MUFA-PCs O | PC O-28:1 + PC O-30:1 + PC O-32:1 + PC O-34:1 + PC O-36:1 + PC O-38:1 + PC O-40:1 + PC O-42:1 |
| Sum of PUFA-PCs O | PC O-34:3 + PC O-36:3 + PC O-36:4 + PC O-36:5 + PC O-38:3 + PC O-38:4 + PC O-38:5 + PC O-38:6 + PC O-40:3 + PC O-40:4 + PC O-40:5 + PC O-40:6 + PC O-42:3 + PC O-42:4 + PC O-42:5 + PC O-44:3 + PC O-44:4 + PC O-44:5 + PC O-44:6 |
| Sum of PUFA-PCs | PC 32:3 + PC 34:3 + PC 34:4 + PC 36:3 + PC 36:4 + PC 36:5 + PC 36:6 + PC 38:3 + PC 38:4 + PC 38:5 + PC 38:6 + PC 40:3 + PC 40:4 + PC 40:5 + PC 40:6 + PC 42:4 + PC 42:5 + PC 42:6 |

| | |
|---|---|
| Sum of PUFA-PC (O)s | PC 32:3 + PC 34:3 + PC 34:4 + PC 36:3 + PC 36:4 + PC 36:5 + PC 36:6 + PC 38:3 + PC 38:4 + PC 38:5 + PC 38:6 + PC 40:3 + PC 40:4 + PC 40:5 + PC 40:6 + PC 42:4 + PC 42:5 + PC 42:6 + PC O-34:3 + PC O-36:3 + PC O-36:4 + PC O-36:5 + PC O-38:3 + PC O-38:4 + PC O-38:5 + PC O-38:6 + PC O-40:3 + PC O-40:4 + PC O-40:5 + PC O-40:6 + PC O-42:3 + PC O-42:4 + PC O-42:5 + PC O-44:3 + PC O-44:4 + PC O-44:5 + PC O-44:6 |
| Sum of UFA-PCs O | PC O-28:1 + PC O-30:1 + PC O-30:2 + PC O-32:1 + PC O-32:2 + PC O-34:1 + PC O-34:2 + PC O-34:3 + PC O-36:1 + PC O-36:2 + PC O-36:3 + PC O-36:4 + PC O-36:5 + PC O-38:1 + PC O-38:2 + PC O-38:3 + PC O-38:4 + PC O-38:5 + PC O-38:6 + PC O-40:1 + PC O-40:2 + PC O-40:3 + PC O-40:4 + PC O-40:5 + PC O-40:6 + PC O-42:1 + PC O-42:2 + PC O-42:3 + PC O-42:4 + PC O-42:5 + PC O-44:3 + PC O-44:4 + PC O-44:5 + PC O-44:6 |
| Sum of PUFA-PEs | PE 34:3 + PE 34:4 + PE 35:3 + PE 36:3 + PE 36:4 + PE 36:5 + PE 36:6 + PE 38:3 + PE 38:4 + PE 38:5 + PE 38:6 + PE 38:7 + PE 40:3 + PE 40:4 + PE 40:5 + PE 40:6 + PE 40:7 + PE 40:8 + PE 42:7 + PE 42:8 + PE 44:6 + PE 44:7 + PE 44:11 + PE 44:12 |
| Ratio of MUFA-PGs to SFA-PGs | (PG 15:0_18:1 + PG 16:0_16:1 + PG 16:0_18:1 + PG 16:0_19:1 + PG 16:0_22:1 + PG 16:1_16:1 + PG 16:1_18:0 + PG 16:1_18:1 + PG 16:1_22:1 + PG 17:0_18:1 + PG 17:1_18:1 + PG 18:0_18:1 + PG 18:0_22:1 + PG 18:1_18:1 + PG 18:1_20:0 + PG 18:1_20:1 + PG 18:1_22:0 + PG 18:1_22:1) / (PG 14:0_16:0 + PG 16:0_16:0) |
| Sum of MUFA-PGs | PG 16:1_16:1 + PG 18:1_20:1 + PG 16:0_19:1 + PG 16:0_22:1 + PG 18:0_18:1 + PG 18:0_22:1 + PG 15:0_18:1 + PG 16:1_18:1 + PG 16:0_16:1 + PG 16:1_22:1 + PG 17:0_18:1 + PG 17:1_18:1 + PG 18:1_18:1 + PG 16:0_18:1 + PG 18:1_22:1 + PG 16:1_18:0 + PG 18:1_20:0 + PG 18:1_22:0 |
| Sum of SFA-PGs | PG 16:0_16:0 + PG 14:0_16:0 |
| Ratio of MUFA-PIs to SFA-PIs | (PI 14:0_18:1 + PI 15:1_16:0 + PI 16:0_17:1 + PI 16:0_18:1 + PI 16:0_22:1 + PI 16:1_18:0 + PI 17:0_18:1 + PI 18:0_18:1 + PI 18:1_20:0 + PI 18:1_22:0 + PI 18:1_22:1 + PI 18:1_20:1 + PI 17:1_18:1 + PI 18:1_18:1 + PI 16:1_18:1) / (PI 15:0_16:0 + PI 16:0_16:0 + PI 16:0_17:0 + PI 16:0_20:0 + PI 18:0_18:0 + PI 18:0_20:0 + PI 18:0_22:0) |
| Ratio of PUFA-PIs to MUFA-PIs | (PI 14:0_18:2 + PI 16:0_17:2 + PI 16:0_18:2 + PI 16:0_18:3 + PI 16:0_20:3 + PI 16:0_20:4 + PI 16:1_18:2 + PI 17:1_18:2 + PI 18:0_18:2 + PI 18:0_18:3 + PI 18:0_20:3 + PI 18:0_20:4 + PI 18:1_18:2 + PI 18:1_18:3 + PI 18:1_20:2 + PI 18:1_20:3 + PI 18:1_20:4 + PI 18:1_20:5 + PI 18:1_22:2 + PI 18:1_22:3 + PI 18:1_22:4 + PI 18:1_22:5 + PI 18:1_22:6 + PI 18:2_18:3 + PI 18:2_20:0 + PI 18:2_20:1 + PI 18:2_20:4 + PI 18:2_20:5 + PI 18:2_22:0 + PI 18:2_22:1 + PI 18:2_22:6) / (PI 14:0_18:1 + PI 15:1_16:0 + PI 16:0_17:1 + PI 16:0_18:1 + PI 16:0_22:1 + PI 16:1_18:0 + PI 17:0_18:1 + PI 18:0_18:1 + PI 18:1_20:0 + PI 18:1_22:0 + PI 18:1_22:1 + PI 18:1_20:1 + PI 18:1_18:1 + PI 17:1_18:1 + PI 16:1_18:1) |
| Ratio of PUFA-PIs to SFA-PIs | (PI 14:0_18:2 + PI 16:0_17:2 + PI 16:0_18:2 + PI 16:0_18:3 + PI 16:0_20:3 + PI 16:0_20:4 + PI 16:1_18:2 + PI 17:1_18:2 + PI 18:0_18:2 + PI 18:0_18:3 + PI 18:0_20:3 + PI 18:0_20:4 + PI 18:1_18:2 + PI 18:1_18:3 + PI 18:1_20:2 + PI 18:1_20:3 + PI 18:1_20:4 + PI 18:1_20:5 + PI 18:1_22:2 + PI 18:1_22:3 + PI 18:1_22:4 + PI 18:1_22:5 + PI 18:1_22:6 + PI 18:2_18:3 + PI 18:2_20:0 + PI 18:2_20:1 + PI 18:2_20:4 + PI 18:2_20:5 + PI 18:2_22:0 + PI 18:2_22:1 + PI 18:2_22:6) / (PI 15:0_16:0 + PI 16:0_16:0 + PI 16:0_17:0 + PI 16:0_20:0 + PI 18:0_18:0 + PI 18:0_20:0 + PI 18:0_22:0) |

| | |
|---|---|
| Ratio of UFA-PIs to SFA-PIs | (PI 14:0_18:1 + PI 14:0_18:2 + PI 15:1_16:0 + PI 16:0_17:1 + PI 16:0_17:2 + PI 16:0_18:1 + PI 16:0_18:2 + PI 16:0_18:3 + PI 16:0_20:3 + PI 16:0_20:4 + PI 16:0_22:1 + PI 16:1_18:0 + PI 16:1_18:1 + PI 16:1_18:2 + PI 17:0_18:1 + PI 17:1_18:1 + PI 17:1_18:2 + PI 18:0_18:1 + PI 18:0_18:2 + PI 18:0_18:3 + PI 18:0_20:3 + PI 18:0_20:4 + PI 18:1_18:1 + PI 18:1_18:2 + PI 18:1_18:3 + PI 18:1_20:0 + PI 18:1_20:1 + PI 18:1_20:2 + PI 18:1_20:3 + PI 18:1_20:4 + PI 18:1_20:5 + PI 18:1_22:0 + PI 18:1_22:1 + PI 18:1_22:2 + PI 18:1_22:3 + PI 18:1_22:4 + PI 18:1_22:5 + PI 18:1_22:6 + PI 18:2_18:3 + PI 18:2_20:0 + PI 18:2_20:1 + PI 18:2_20:4 + PI 18:2_20:5 + PI 18:2_22:0 + PI 18:2_22:1 + PI 18:2_22:6) / (PI 15:0_16:0 + PI 16:0_16:0 + PI 16:0_17:0 + PI 16:0_20:0 + PI 18:0_18:0 + PI 18:0_20:0 + PI 18:0_22:0) |
| Ratio of PI 18:0_20:4 to PIs | PI 18:0_20:4 / (PI 14:0_18:1 + PI 14:0_18:2 + PI 15:0_16:0 + PI 15:1_16:0 + PI 16:0_16:0 + PI 16:0_17:0 + PI 16:0_17:1 + PI 16:0_17:2 + PI 16:0_18:1 + PI 16:0_18:2 + PI 16:0_18:3 + PI 16:0_20:0 + PI 16:0_20:3 + PI 16:0_20:4 + PI 16:0_22:1 + PI 16:1_18:0 + PI 16:1_18:1 + PI 16:1_18:2 + PI 17:0_18:1 + PI 17:1_18:1 + PI 17:1_18:2 + PI 18:0_18:0 + PI 18:0_18:1 + PI 18:0_18:2 + PI 18:0_18:3 + PI 18:0_20:0 + PI 18:0_20:3 + PI 18:0_20:4 + PI 18:0_22:0 + PI 18:1_18:1 + PI 18:1_18:2 + PI 18:1_18:3 + PI 18:1_20:0 + PI 18:1_20:1 + PI 18:1_20:2 + PI 18:1_20:3 + PI 18:1_20:4 + PI 18:1_20:5 + PI 18:1_22:0 + PI 18:1_22:1 + PI 18:1_22:2 + PI 18:1_22:3 + PI 18:1_22:4 + PI 18:1_22:5 + PI 18:1_22:6 + PI 18:2_18:3 + PI 18:2_20:0 + PI 18:2_20:1 + PI 18:2_20:4 + PI 18:2_20:5 + PI 18:2_22:0 + PI 18:2_22:1 + PI 18:2_22:6) |
| Sum of MUFA-PIs | PI 14:0_18:1 + PI 15:1_16:0 + PI 16:0_17:1 + PI 16:0_18:1 + PI 16:0_22:1 + PI 16:1_18:0 + PI 17:0_18:1 + PI 18:0_18:1 + PI 18:1_20:0 + PI 18:1_22:0 + PI 16:1_18:1 + PI 17:1_18:1 + PI 18:1_18:1 + PI 18:1_20:1 + PI 18:1_22:1 |
| Sum of PIs | PI 14:0_18:1 + PI 14:0_18:2 + PI 15:0_16:0 + PI 15:1_16:0 + PI 16:0_16:0 + PI 16:0_17:0 + PI 16:0_17:1 + PI 16:0_17:2 + PI 16:0_18:1 + PI 16:0_18:2 + PI 16:0_18:3 + PI 16:0_20:0 + PI 16:0_20:3 + PI 16:0_20:4 + PI 16:0_22:1 + PI 16:1_18:0 + PI 16:1_18:1 + PI 16:1_18:2 + PI 17:0_18:1 + PI 17:1_18:1 + PI 17:1_18:2 + PI 18:0_18:0 + PI 18:0_18:1 + PI 18:0_18:2 + PI 18:0_18:3 + PI 18:0_20:0 + PI 18:0_20:3 + PI 18:0_20:4 + PI 18:0_22:0 + PI 18:1_18:1 + PI 18:1_18:2 + PI 18:1_18:3 + PI 18:1_20:0 + PI 18:1_20:1 + PI 18:1_20:2 + PI 18:1_20:3 + PI 18:1_20:4 + PI 18:1_20:5 + PI 18:1_22:0 + PI 18:1_22:1 + PI 18:1_22:2 + PI 18:1_22:3 + PI 18:1_22:4 + PI 18:1_22:5 + PI 18:1_22:6 + PI 18:2_18:3 + PI 18:2_20:0 + PI 18:2_20:1 + PI 18:2_20:4 + PI 18:2_20:5 + PI 18:2_22:0 + PI 18:2_22:1 + PI 18:2_22:6 |
| Sum of PUFA-PIs | PI 14:0_18:2 + PI 16:0_17:2 + PI 16:0_18:2 + PI 16:0_18:3 + PI 16:0_20:3 + PI 16:0_20:4 + PI 16:1_18:2 + PI 17:1_18:2 + PI 18:0_18:2 + PI 18:0_18:3 + PI 18:0_20:3 + PI 18:0_20:4 + PI 18:1_18:2 + PI 18:1_18:3 + PI 18:1_20:2 + PI 18:1_20:3 + PI 18:1_20:4 + PI 18:1_20:5 + PI 18:1_22:2 + PI 18:1_22:3 + PI 18:1_22:4 + PI 18:1_22:5 + PI 18:1_22:6 + PI 18:2_18:3 + PI 18:2_20:0 + PI 18:2_20:1 + PI 18:2_20:4 + PI 18:2_20:5 + PI 18:2_22:0 + PI 18:2_22:1 + PI 18:2_22:6 |
| Sum of SFA-PIs | PI 15:0_16:0 + PI 16:0_16:0 + PI 16:0_17:0 + PI 16:0_20:0 + PI 18:0_18:0 + PI 18:0_20:0 + PI 18:0_22:0 |

| | |
|---|---|
| Sum of UFA-PIs | PI 14:0_18:1 + PI 14:0_18:2 + PI 15:1_16:0 + PI 16:0_17:1 + PI 16:0_17:2 + PI 16:0_18:1 + PI 16:0_18:2 + PI 16:0_18:3 + PI 16:0_20:3 + PI 16:0_20:4 + PI 16:0_22:1 + PI 16:1_18:0 + PI 16:1_18:1 + PI 16:1_18:2 + PI 17:0_18:1 + PI 17:1_18:1 + PI 17:1_18:2 + PI 18:0_18:1 + PI 18:0_18:2 + PI 18:0_18:3 + PI 18:0_20:3 + PI 18:0_20:4 + PI 18:1_18:1 + PI 18:1_18:2 + PI 18:1_18:3 + PI 18:1_20:1 + PI 18:1_20:2 + PI 18:1_20:3 + PI 18:1_20:4 + PI 18:1_20:5 + PI 18:1_22:0 + PI 18:1_22:1 + PI 18:1_22:2 + PI 18:1_22:3 + PI 18:1_22:4 + PI 18:1_22:5 + PI 18:1_22:6 + PI 18:2_18:3 + PI 18:2_20:0 + PI 18:2_20:1 + PI 18:2_20:4 + PI 18:2_20:5 + PI 18:2_22:0 + PI 18:2_22:1 + PI 18:2_22:6 |
| Ratio of PUFA-PSs to MUFA-PSs | (PS 36:3 + PS 36:4 + PS 36:5 + PS 38:4 + PS 38:5 + PS 38:6 + PS 38:7 + PS 40:4 + PS 40:5 + PS 40:6 + PS 40:7 + PS 40:8) / (PS 34:1 + PS 36:1) |
| Sum of MUFA-PSs | PS 34:1 + PS 36:1 |
| Sum of PUFA-PSs | PS 36:3 + PS 36:4 + PS 36:5 + PS 38:4 + PS 38:5 + PS 38:6 + PS 38:7 + PS 40:4 + PS 40:5 + PS 40:6 + PS 40:7 + PS 40:8 |
| Sum of UFA-PSs | PS 34:1 + PS 34:2 + PS 36:1 + PS 36:2 + PS 36:3 + PS 36:4 + PS 36:5 + PS 38:4 + PS 38:5 + PS 38:6 + PS 38:7 + PS 40:4 + PS 40:5 + PS 40:6 + PS 40:7 + PS 40:8 |
| SPBP d18:1 to Cer d18:1/24:1 Ratio | SPBP d18:1 / Cer d18:1/24:1 |
| SPHK activity (4) | SPBP d18:1 / SPB d18:1 |
| SPHK activity (6) | SPBP d14:0 / SPB d14:0 |
| SPHK activity (8) | SPBP d17:0 / SPB d17:0 |
| Sum of SphoPs | SPBP d14:1 + SPBP d16:1 + SPBP d17:1 + SPBP d18:1 |
| LPP3 activity (4) | SPB d18:1 / SPBP d18:1 |
| Sum of Spha | SPB d14:0 + SPB d16:0 + SPB d17:0 + SPB d18:0 |
| Ratio of OC-FA SMs to EC-FA SMs | (SM 33:1 + SM 35:1 + SM 41:1 + SM 41:2 + SM 43:1) / (SM 34:1 + SM 34:2 + SM 36:1 + SM 36:2 + SM 38:3 + SM 42:1 + SM 42:2 + SM 44:1 + SM 44:2) |
| Sum of EC-FA SMs | SM 34:1 + SM 34:2 + SM 36:1 + SM 36:2 + SM 38:3 + SM 42:1 + SM 42:2 + SM 44:1 + SM 44:2 |
| Sum of LCFA-SMs | SM 33:1 + SM 35:1 + SM 34:1 + SM 34:2 + SM 36:1 + SM 36:2 |
| Sum of OC-FA SMs | SM 33:1 + SM 35:1 + SM 41:1 + SM 41:2 + SM 43:1 |
| Sum of SMs | SM 33:1 + SM 35:1 + SM 41:1 + SM 41:2 + SM 43:1 + SM 34:1 + SM 34:2 + SM 36:1 + SM 36:2 + SM 38:3 + SM 42:1 + SM 42:2 + SM 44:1 + SM 44:2 |
| Sum of VLCFA-SMs | SM 43:1 + SM 42:1 + SM 42:2 + SM 44:1 + SM 44:2 |

**SUPPLEMENTAL DATA 4**

| Analyte class | Number of molecules |
|---|---|
| Acylcarnitines | 40 |
| Alkaloids | 2 |
| Amine oxides | 1 |
| Amino acid-related | 77 |
| Amino acids | 20 |
| Bile acids | 24 |
| Biogenic amines | 10 |
| Carboxylic acids | 8 |
| Catechols | 3 |
| Ceramides | 29 |
| Cholesteryl esters | 22 |
| Cresols | 2 |
| Dicarboxylic acids | 25 |
| Diglycerides | 41 |
| Dihexosylceramides | 9 |
| Dihydroceramides | 8 |
| Fatty acids | 39 |
| Hexosylceramides | 20 |
| Hormones | 5 |
| Indoles and derivatives | 17 |
| Lysophosphatidic acids | 8 |
| Lysophosphatidylcholines | 12 |
| Lysophosphatidylethanolamines | 43 |
| Lysophosphatidylglycerols | 10 |
| Lysophosphatidylinositols | 15 |
| Lysophosphatidylserines | 12 |
| Monoglycerides | 12 |
| Nucleobase-related | 14 |
| Nucleobases | 5 |
| Nucleotides | 2 |
| Organic acids | 16 |
| Phenolic acids | 22 |
| Phenoxy compounds | 2 |
| Phosphatidic acids | 41 |
| Phosphatidylcholines | 76 |
| Phosphatidylethanolamines | 95 |
| Phosphatidylglycerols | 64 |
| Phosphatidylinositols | 53 |
| Phosphatidylserines | 18 |

| | |
|---|---|
| Polyamines | 7 |
| Pyridinecarboxylic acids | 6 |
| Sphinganine and sphingosine phosphates | 8 |
| Sphinganines and sphingosines | 8 |
| Sphingomyelins | 14 |
| Sugars | 7 |
| Tricarboxylic acids | 3 |
| Triglycerides | 242 |
| Trihexosylceramides | 6 |
| Vitamins and cofactors | 8 |
